# Live-cell imaging of enhancer-promoter dynamics reveals transient contact-driven gene activation

**DOI:** 10.64898/2026.06.21.733446

**Authors:** Jin H. Yang, Henrik D. Pinholt, Jack Toppen, Miles K. Huseyin, James M. Jusuf, Christos C. Katsifis, Jacob Kæstel-Hansen, Leonid A. Mirny, Christoph Zechner, Anders S. Hansen

## Abstract

Enhancers are key regulators of mammalian gene expression, yet how they interact with promoters in space (contact vs. action-at-a-distance) and in time (transient vs. stable) remains poorly understood. Recent studies suggest that enhancers can activate promoters across distances exceeding 200 nanometers, challenging classical contact models, but limited spatiotemporal resolution has obscured the mechanistic details of enhancer-promoter (E–P) interactions and their link to transcription. Here, we engineered a synthetic biology platform optimized for the simultaneous visualization of E–P 3D distance and nascent transcription using super-resolution live-cell imaging. By applying five complementary approaches integrating imaging, 3D genomics, and gene expression data across cell lines, we estimate that transcriptional activation is mediated by ∼25-42 nanometer contacts on the seconds timescale. Our results support a transient contact mechanism for E–P-mediated gene activation.

## Introduction

Fine-tuned regulation of gene expression is essential for cellular function. Enhancers are primary units of gene control in mammals (*1–4*). Enhancers are frequently located distally and are thought to loop to their target genes to activate them. For example, the key enhancer for the oncogene *MYC* in T cells is located 1.8 Mb downstream of *MYC* and activates *MYC* through enhancer-promoter (E-P) looping (*5*). Although mammalian enhancers were first discovered in 1981 (*6, 7*) and despite their central importance in gene regulation, the mechanisms of E-P looping in time and space remain debated (*1, 2, 8–11*).

The classic textbook model for E-P looping is the “stable contact model” (*12*) (**Fig. 1A**), wherein the loop brings transcriptional activators bound at the enhancer into stable contact with the promoter, thus enabling transcription activation. However, several lines of evidence argue against stable E-P loops. First, live-cell imaging studies have shown that the structural loops formed by CTCF and cohesin are short-lived ( ∼10–30 min lifetimes) (*13–15*). Because E-P loops tend to be weaker than CTCF loops, E-P loops may be even more transient (*16–18*). More directly, imaging of E-P looping in live cells has also failed to detect stable contact (*19–27*). Together, these observations would in principle be more consistent with a transient contact model (“hit-and-run”, **Fig. 1A**).

**Figure 1:**
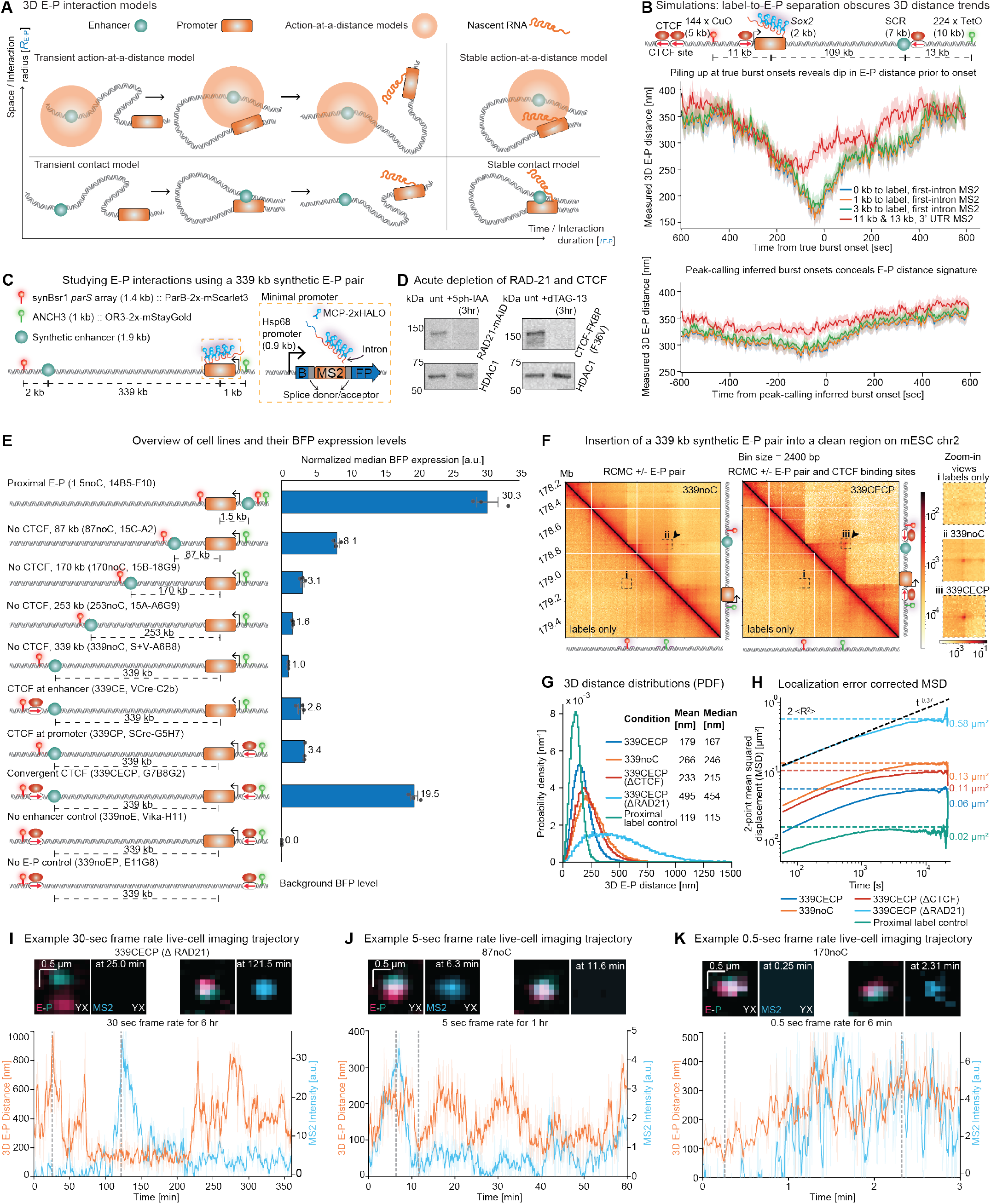
A synthetic bottom-up approach to mechanistically dissect enhancer-promoter interactions. (**A**) Threedimensional models of E–P interactions. Along the interaction radius axis, 3D E–P interaction models can be divided into contact type and action-at-a-distance type; along the interaction duration axis, 3D E–P interaction models are classified into transient and stable categories. (**B**) Simulation of the effect of label-to-E-P separation, MS2 placement, and burst onset calling methods on the ability to detect the correlation between E-P spatial proximity and transcription. The schematic illustrates the simulation setup for the *Sox2* locus, where the simulated 3D conformations of *Sox2* and SCR were coupled with a simple two-state promoter model, with E-P fluorescent probe placement similar to (*19*). Below we compared the E-P distance pile-up at the precise moments where the two-state promoter transitioned from the OFF to the ON state (middle panel) versus at the burst onsets determined by peak-calling (lower panel), with different label-to-EP separations and MS2 placements. (**C**) Schematic of the 339 kb synthetic E-P pair integrated into a ‘vacant and clean’ region on chromosome 2 of mESCs, featuring fluorescent labeling of the enhancer (synBsr1 *parS*) and promoter (ANCH3), alongside an intronic MS2 cassette for nascent transcription labeling and a tagBFP reporter for quantification of expression using flow cytometry. (**D**) Western blots confirming the acute depletion of RAD21 (via 5ph-IAA) and CTCF (via dTAG-13) using endogenous mAID and FKBP(F36V) degron tags, with HDAC1 serving as a loading control. (**E**) Overview of engineered cell lines with varying E-P configurations and CTCF binding site (CBS) placement, alongside normalized median BFP expression levels (normalized to 339noC; **Fig. S4**). Error bars represent the standard error of the mean (SEM) across three biological replicates. (**F**) Region Capture Micro-C (RCMC) contact maps of the 339 kb synthetic E-P pair demonstrating focal enrichment (left panel) that is further strengthened by convergent CBSs (right panel). An 80 kb zoom-in window centered around the E-P coordinates is shown on the right. (**G**) PDF of measured 3D E-P distances across various 339 kb E-P configurations along with the proximal label control cell line and cohesin/CTCF depletion conditions. (**H**) Localization error-corrected two-point mean squared displacement (MSD) of the cell lines in (G). Solid lines represent the empirical two-point MSDs. Horizontal dashed lines indicate the theoretical MSD plateau at 2 ⟨*R*^2^⟩ (⟨*R*^2^⟩ is the mean squared 3D distance between the E-P pair). The black dashed line indicates a reference power-law scaling of *t*^0.37^. (**I-K**) Representative live-cell imaging trajectories demonstrating the temporal dynamics of 3D E-P distance (orange) and corresponding MS2 intensity (cyan) across 30-second (I), 5-second (J), and 0.5-second (K) frame rates. The faint background lines represent the raw trajectories, and the bold solid lines represent the smoothed trajectories with a 5-frame rolling average.

However, mounting evidence argues against E-P contact models altogether. In *Drosophila*, E-P pairs were found to be separated by distances of ∼200 nm during transcription activation (*20, 28, 29*), which challenges the notion of contact (the largest transcriptional complexes are ∼20–40 nm (*10, 30–32*)) and instead argues for enhancer “actionat-a-distance” models (*2, 11, 33*) (**Fig. 1A**). Similarly, a pioneering study that simultaneously visualized *Sox2* E-P looping and nascent transcription in mouse embryonic stem cells (mESCs) failed to detect E-P contact (*19*). In fact, the study seemingly ruled out contact models by reporting no relationship between E-P proximity and transcription initiation (*19*). Subsequent work on *Sox2* found condensates to predict bursting better than E-P distance (*34*). Consistently, a study of the *Shh* E-P pair during mESC differentiation found that *Shh* activation was associated with increased 3D distances (*35*), the exact opposite of what the E-P contact model would predict. Similar observations were made for estrogen-responsive genes (*36*).

Notably, despite studying a variety of different E-P pairs in different organisms including *Drosophila*, mouse, and human cells, prior imaging studies of E-P looping have either failed to detect E-P contact altogether or found that 3D distances of ≥200 nm separate the E-P pair during transcription activation (*2, 11, 33*). Clearly, ≥200 nm is not compatible with the classic textbook model of contact (**Fig. 1A**). Instead, the observed absence of contact has led to the concept of an “enhancer sphere of influence” and motivated the development of mechanistic models that can explain the observed enhancer “action-at-a-distance” (*2*). These ‘action-at-a-distance’ models include 1) condensate or hub models, where ≥200 nm-sized condensates bridge between the E-P pair (*3, 33, 34, 37, 38*); 2) gradient models, where the enhancer establishes a gradient of acetylated transcription factors to activate the promoter without direct contact (*11*) or RNA gradient models (*39*); 3) polymerization models, where for example poly-ADP-ribosylation bridges the E-P pair (*35*). Regardless of the molecular mechanism, both stable and transient “action-at-a-distance” models have been proposed (*1, 8*) (**Fig. 1A**).

Despite the strong evidence for “action-at-a-distance” models, if the “hit-and-run transient contact” model were true and E-P contact events were very transient, it remains possible that they might have been missed by prior studies due to technical biases inherent to imaging-based studies of E-P looping (*1, 2, 8, 10, 40, 41*) (**SI Note 1**).

Given the foundational importance of understanding the mechanisms of E-P-mediated transcription activation (**Fig. 1A**), here we took a bottom-up synthetic biology approach to investigate spatiotemporal E-P mechanisms. Specifically, we engineered into a “vacant” genomic region a synthetic E-P pair specifically optimized to enable superresolution live-cell imaging of the spatiotemporal dynamics of E-P-mediated gene activation. We applied five complementary approaches that all consistently point to a transient contact model for E-P interactions.

## Results

### Design considerations for studying E-P interactions

To design an optimal experimental system for studying E-P looping (*1,8,40*), we began by revisiting the landmark study of the *Sox2* E-P pair in mESCs (*19*). This work achieved the first simultaneous 3-color live-cell imaging of E-P looping and nascent transcription in mammalian cells (*19*) after earlier reports in *Drosophila* (*20*). By using “MS2 burst pileup analysis” (*19, 20*), they reported the complete absence of correlation between E-P proximity and transcription burst initiation (*19*), seemingly ruling out contact models. Of the four E-P models (**Fig. 1A**), the transient E-P contact “hit-and-run” model is by far the most difficult to detect and exclude. To understand if transient E-P contact can be ruled out based on the MS2 burst pile-up analysis, we performed 3D polymer simulations of the *Sox2* locus coupled to a “hit-and-run” transient contact model for transcription activation (**SI Note 1**). We found that a true causal relationship between transient E-P contact (**Fig. 1B, top**) can nevertheless completely disappear (**Fig. 1B, bottom**), due to a combination of localization uncertainty, a stochastic delay between E-P contact and MS2-signal appearance, and the challenge of identifying true burst onsets. This analysis (**SI Note 1**) pinpointed eight key criteria for investigating E-P interactions in a manner that allows distinction between all four mechanistic models outlined in **Fig. 1A**: 1) Use small ( ≤2 kb) fluorescent DNA labels; 2) Minimize separation between DNA label and E/P; 3) Use small E/P ( ≤2 kb); 4) Place the nascent RNA MS2 array near the 5’-UTR to minimize time-delay between E-P proximity and nascent RNA detection; 5) Ensure promoter transcription is entirely dependent on a single enhancer; 6) Maximize E-P separation (*>*100 kb) to enable distinction between “looping” and “no looping”; 7) Image fast enough and for long enough to detect E-P looping events; 8) Account for noise and time-delays in the analysis. To simultaneously achieve all eight criteria, we took a synthetic biology approach and engineered an E-P pair into a large “vacant” genomic region in mESCs (**Fig. 1C, Fig. S1A**). The purpose of the synthetic system is not to mimic every endogenous locus, but to create a controlled physical system in which contact and action-at-a-distance models make separable predictions.

### A bottom-up approach using a synthetic E-P pair

To engineer a synthetic enhancer that is both strong and sufficiently compact for high-fidelity live-cell imaging, we fused three previously characterized ∼0.6 kb superenhancer fragments (*42*) into a 1.9 kb construct (**Fig. 1C, Fig. S1B**). For the promoter, we chose the 0.9 kb *Hsp68* promoter fragment, which lacks activity on its own in mESCs and is responsive to distal enhancers (*43, 44*). We placed 128xMS2 (*45*) in the first and only intron (*46*) such that binding of the MCP-2x-Halo proteins to the MS2 RNA hairpins allows for the direct imaging of nascent transcription. We also encoded tagBFP in the two exons so that we can quantify transcription indirectly with flow cytometry measurement of BFP fluorescence (**Fig. 1C**). The synthetic E-P pair was placed 339 kb apart to maximize the dynamic range of 3D E-P distance (**Fig. 1C**). We labeled the promoter and enhancer with ANCH3 (*13, 47*) and synBsr1 *parS* arrays (a novel DNA labeling system developed in this study, **Fig. S1C**), respectively, which enables live-cell imaging of the E-P pair with the fluorescently tagged binding proteins OR3-2x-mStayGold and synBsr1ParB-2x-mScarlet3 (*48*) (**Fig. 1C**). To elucidate the role of cohesin and CTCF in mediating E-P interactions, we endogenously tagged RAD21 (a cohesin subunit) with mAID (*49*) and CTCF with FKBP(F36V) (*50*), enabling acute depletion with the addition of dTAG-13 and 5ph-IAA, respectively (**Fig. 1D, Fig. S2A,B**).

Because gene expression is expected to depend on genomic distance between the E-P pair as a function of EP interaction radius, *R*_E−P_ (the distance threshold for productive E-P interactions) and the E-P interaction duration (*τ*_E−P_), we engineered four additional cell lines that progressively shortened the E-P genomic distance to 253 kb, 170 kb, 87 kb, and 1.5 kb to help constrain the estimates of *R*_E−P_ and *τ*_E−P_ (**Fig. 1E, Figs. S3 and S4**). As expected (*51*), BFP expression increases monotonically with decreasing genomic distance, with the 1.5 kb E-P pair exhibiting a ∼30-fold increase in expression compared with the 339 kb pair (**Fig. 1E**). Additionally, because CTCF’s role as a facilitator of E-P interactions is expected to be highly dependent on *R*_E−P_ (*1*), we engineered three additional cell lines with 3x CTCF binding sites (CBSs) (**Fig. S2C,D**) either at the promoter, the enhancer, or both (**Fig. 1E, Figs. S3 and S4**), where convergent CBSs conferred a ∼20-fold increase in expression compared to the E-P pair lacking CBSs entirely. The control cell line with the synthetic *Hsp68* promoter alone (339noE) had no BFP expression (**Fig. 1E, Fig. S4**), thus establishing our synthetic promoter as a faithful reporter of the activity of the synthetic enhancer. To test if the insertion led to an observable E-P loop, we performed Region Capture Micro-C (RCMC) (*52*). The insertion of the 339 kb synthetic E-P pair led to focal enrichment between the E-P pair (**Fig. 1F**), which was substantially strengthened with convergent CBSs. Notably, synBsr1 *parS* and ANCH3 insertion alone does not lead to the same focal enrichment, validating that the “loop dots” on the RCMC maps indeed correspond to E-P interactions (**Fig. 1F, Fig. S5**).

To directly observe the relationship between E-P 3D distance and nascent transcription, we optimized and performed lattice light-sheet based Super-Resolution Live-Cell Imaging (llsSRLCI) and the movies were subsequently converted to 3D E-P distance trajectories and MS2 signal trajectories (**Fig. 1G-K, Figs. S6–S10**). llsSRLCI enables gentle, large field-of-view imaging with ∼40 nm isotropic super-resolution (**Tables S6 and S7**) and across almost 5 orders of magnitude in time, a substantial improvement over our past study (*13*). To avoid the confounding effects (*8, 13*) of replicated sister chromatids which result in E-P trajectories that cannot be precisely localized and contain two copies of the gene, we filtered these out (**Fig. S10**).

We validated our trajectory analysis with a series of controls. As expected, adding 3xCBSs to the 339noC line strongly decreased the mean E-P distance from 266 to 179 nm (**Fig. 1G**), while cohesin depletion dramatically increased it to 495 nm, consistent with prior work (*13–15*). As a co-localization control (**Fig. S3**), we placed the ANCH3 and synBsr1 labels only 0.4 kb apart, which strongly reduced the mean 3D distance to 119 nm as expected (**Fig. 1G**; as previously discussed (*1, 8*), due to localization uncertainty, co-localization will result in non-zero measured distances). Furthermore, we observed a strong correlation between our BFPand MS2-based measurements of transcription (**Fig. S11A**). The mean-squared displacement (MSD) of the relative 3D positions of the E-P pair showed a scaling of MSD ∼ *t*^0.37^ across different cell lines (**Fig. 1H, Fig. S12**), consistent with recent MINFLUX measurements in mESCs (*53*). To capture dynamics across transcriptionally relevant time-scales, we acquired live-cell movies at frame rates of 30, 5, and 0.5 seconds, for a total duration of 6 hours, 1 hour, and 6 minutes, respectively (**Fig. 1I, Movies S1-S7**). 3D E-P distance measurements were consistent across frame rates for each condition (**Fig. S11B**). Using Bayesian Inference of Looping Dynamics (BILD) (*13*), we estimated a looping probability of (8.9 ± 0.8)% and a median lifetime of 11 min for the 339CECP cell line (**Fig. S13**), in line with prior studies (*13–15*). Likely due to residual cohesin (∼ 5%) remaining after the RAD21 depletion (**Fig. 1C, Fig. S2A**), we observed an extremely rare *>*2 hour presumably CTCF-CTCF loop in a 6-hour trajectory (**Fig. 1I**, 339CECP ΔRAD21, lasting from ∼75–225 min), which resulted in a large transcriptional burst. In an illustrative 5-second resolution trajectory of the 87noC line (**Fig. 1J**), we observed ‘dips’ in distances preceding increases in MS2 signal. In our 0.5second resolution trajectories, we could capture even faster dynamics and observed an example of a ‘dip’ in the E-P 3D distance lasting a few tens of seconds (**Fig. 1K**) before a transcriptional burst about 60 seconds later.

Having validated our synthetic biology platform, we next sought to elucidate how close enhancers and promoters need to be to initiate transcription (*R*_E−P_) and how long these physical interactions last (*τ*_E−P_) (**Fig. 1A**) by deploying five complementary approaches across our 3D genomics, gene expression, and live-cell imaging data. Despite each approach having distinct limitations, they all consistently support a transient E-P contact model.

### Estimate 1: *R*_E−P_ from 3D structure-function relationships

We first sought to estimate *R*_E−P_, the 3D E-P distance threshold for functional interactions, by quantifying how EP interaction strength (measured using RCMC; **Fig. 2A**) scales with gene expression (measured using BFP or MS2; **Fig. 1E**). RCMC, like all other Chromosome Conformation Capture methods, captures genomic interactions through proximity ligation if the two DNA ends are below a distance threshold, *R*_RCMC_ (*1, 54, 55*). Analogously, E-P interactions induce expression below a distance threshold, *R*_E−P_. The activity-by-contact model (*56*) assumes that expression is proportional to E-P loop strength as also supported by prior work (*57*). Under this assumption (which will be relaxed below), a linear relationship between expression and RCMC-measured E-P loop strength for stabilized loop dots visible in RCMC should only occur if *R*_RCMC_ ≈ *R*_E−P_ (**Fig. 2B, SI Note 2**). In contrast, if *R*_RCMC_ *> R*_E−P_ or *R*_RCMC_ *< R*_E−P_, a concave or convex non-linear relationship would be expected, respectively (**SI Note 2**). Strikingly, expression levels across all E-P configurations, whether quantified by flow cytometry (median BFP level) or live-cell imaging (mean MS2 intensity), exhibit a linear relationship with the RCMC-quantified E-P interaction strength (**Fig. 2A,C**). While the *>*1 slope implies a non-productive interaction threshold (explored later in the form of time-gating), the preservation of linearity across cell lines with different E-P and CBS configuration mathematically requires that both metrics probe the same spatial scales (**SI Note 2**). Notably, this linear relationship was robust to substantially different RCMC quantification methods (**Fig. S14A**; *R*^2^ ∈ [0.94; 0.98] across different quantification resolutions and mask sizes). This indicates that *R*_RCMC_ ≈ *R*_E−P_. *R*_RCMC_ has been previously estimated at ∼42 nm (*58*) (**Fig. 2D**). We independently estimated an optimal *R*_E−P_ of ∼25 nm (95% CI: 23–36 nm) using 3D polymer simulations constrained by these measurements (**Fig. 2D, Fig. S14B**). Integrating this simulation-based estimate with prior experimental estimates of the RCMC capture radius (*58*) and our observation that *R*_RCMC_ ≈ *R*_E−P_ (**Fig. 2B,C**), we thus arrive at our first estimate: *R*_E−P_ ≈ 25–42 nm.

**Figure 2:**
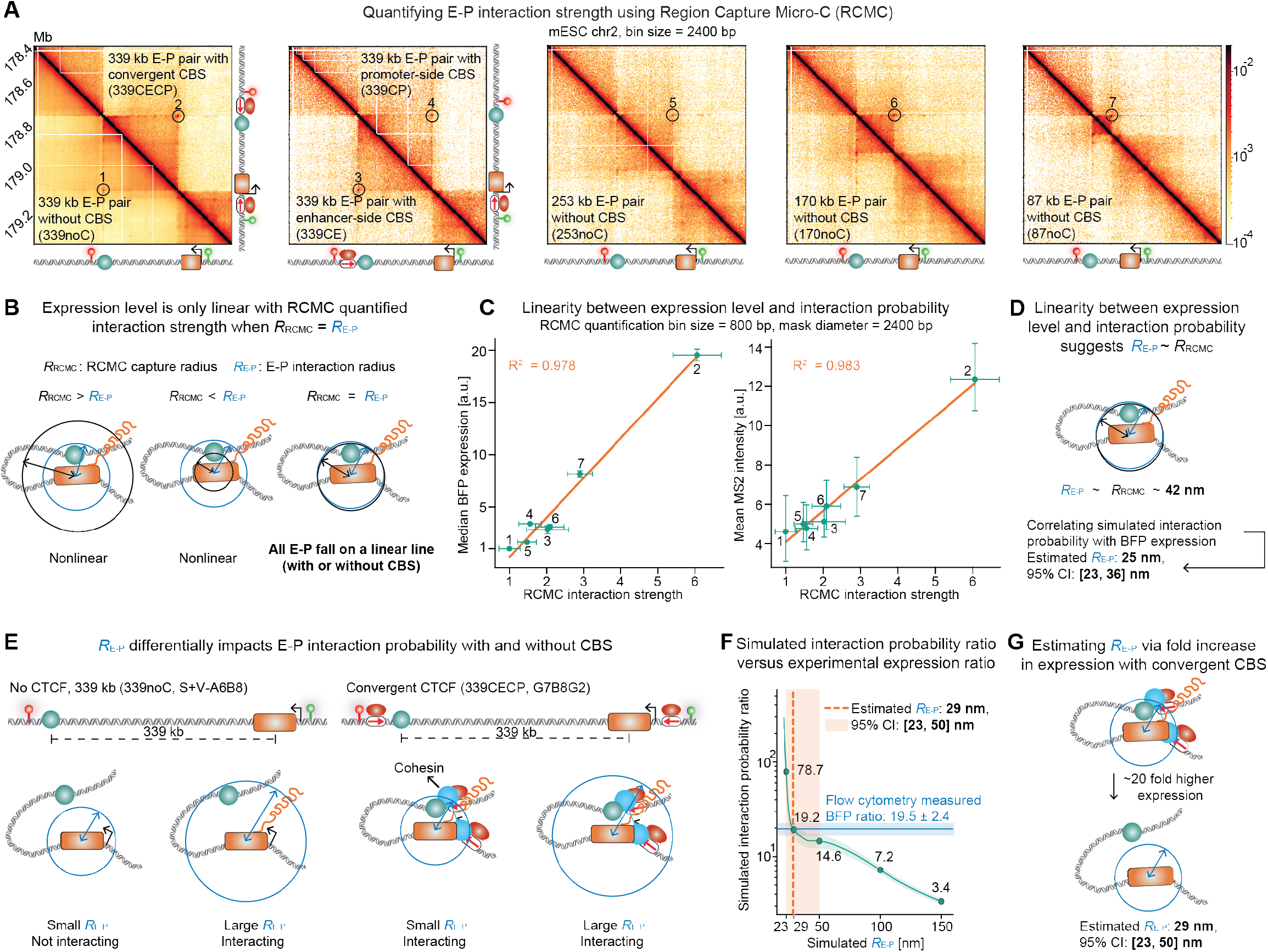
Estimating E-P interaction radius using structure-function relationships. (**A**) Region Capture Micro-C (RCMC) contact maps across different E-P configurations, with the focal E-P interactions highlighted in circles. (**B**) Schematic illustrating that expression level scales linearly with RCMC-quantified interaction strength only when the RCMC capture radius (*R*_RCMC_) approximates the E-P interaction radius (*R*_E−P_). (**C**) Linear correlation between RCMC interaction strength and transcription levels, quantified by median BFP expression (left) and the mean of the per-trajectory average MS2 intensities (right). The *x*-axis represents the contact frequency normalized to the 339noC condition. For both plots, the *x*-axis error bars represent the standard error of the mean (SEM) across two biological replicates. The *y*-axis error bars represent the SEM of median BFP expression across three biological replicates for the left plot, and the 95% confidence interval of the MS2 intensity across trajectories. The numbers next to each data point correspond to the numbered circles in (A). (**D**) Estimation of *R*_E−P_ given the linearity between expression and interaction probability. (**E**) Schematic illustrating how *R*_E−P_ differentially impacts the interaction probabilities of E-P pairs with and without convergent CBSs. (**F**) Simulated interaction probability ratio of 339CECP and 339noC (teal curve), assumed here to be equivalent to their BFP expression ratio, across different *R*_E−P_ values (five values highlighted as points; shaded teal ribbon represents 95% confidence intervals calculated using temporal block bootstrapping). Comparing the simulation curve with the flow cytometry-measured BFP expression ratio (solid horizontal blue line; shaded blue region represents the SEM of median BFP expression across three biological replicates) yields an estimated *R*_E−P_ of ∼29 nm (vertical dashed orange line; 95% CI: 23–50 nm, shaded orange region). (**G**) Schematic summary of the *R*_E−P_ estimate based on the roughly 20-fold expression increase conferred by convergent CBSs.

### Estimate 2: *R*_E−P_ from CTCF-mediated facilitation

As our second estimate of *R*_E−P_, we studied how adding convergent CBSs near the enhancer and promoter affected gene expression (**Fig. 2E**, 339noC vs. 339CECP). The increase in gene expression due to CTCF-mediated facilitation can be used to quantitatively constrain the possible values of *R*_E−P_ (*1*). For example, if *R*_E−P_ were ≈ 250 nm, given the median distance of 246 nm for the E-P pair in the 339noC condition (**Fig. 1G**, 339noC), this E-P pair would be interacting 50% of the time. However, since interaction frequency cannot exceed 100%, CTCF-mediated facilitation could only result in maximally a twofold increase in interaction frequency and expression. This contrasts with our observed 19.5-fold increase in expression upon adding convergent CBSs, which can only be accounted for by a much smaller *R*_E−P_. To understand how much smaller *R*_E−P_ would need to be, we performed experimentally constrained 3D polymer simulations under the assumption that expression is proportional to E-P interaction probability (**Fig. 2E, Figs. S14 and S15**). Only a very small *R*_E−P_ ≈ 29 nm (95% CI: 23 to 50 nm) reproduces the experimentally observed 19.5-fold increase in BFP expression with convergent CBSs (**Fig. 2F**). This simulation-derived estimate is further validated by a direct analytical calculation using the Gaussian chain approximation, which applies the 8.9% looped fraction derived from our BILD analysis (**Fig. S13E**) to independently yield *R*_E−P_ ≈ 28.9 nm (**SI Note 2** section 12.6). Therefore, the ∼20-fold higher expression driven by convergent CBSs provides a second complementary estimate of *R*_E−P_ ≈ 29 nm (**Fig. 2G**).

### Estimate 3: *R*_E−P_ and *τ*_E−P_ from machine learning based analysis of live-cell trajectories

For our third estimate of *R*_E−P_ and our first estimate of *τ*_E−P_ we turned directly to our live-cell trajectories of E-P 3D distance and nascent transcription (**Fig. 1I**). Because prior work did not detect a relationship between E-P proximity and transcriptional burst onset (*19*), we investigated if such a relationship existed in our data using a data-driven machine-learning approach robust to noise (**SI Note 1**). We developed a two-step approach to isolate E-P distance-dependent burst onsets: 1) we applied change-point detection (CPD) to generate a preliminary set of putative burst labels; and 2) we trained a Long Short-Term Memory (LSTM) neural network using exclusively 3D E-P distance-derived features as input to predict these CPD-derived binary labels (**Fig. 3A**). An LSTM model is a type of recurrent neural network that is ideally suited for this prediction task because it is designed to learn relationships in time-ordered data, where an event now (E-P proximity below *R*_E−P_) influences an outcome later (appearance of nascent MS2 burst) (*59*).

**Figure 3:**
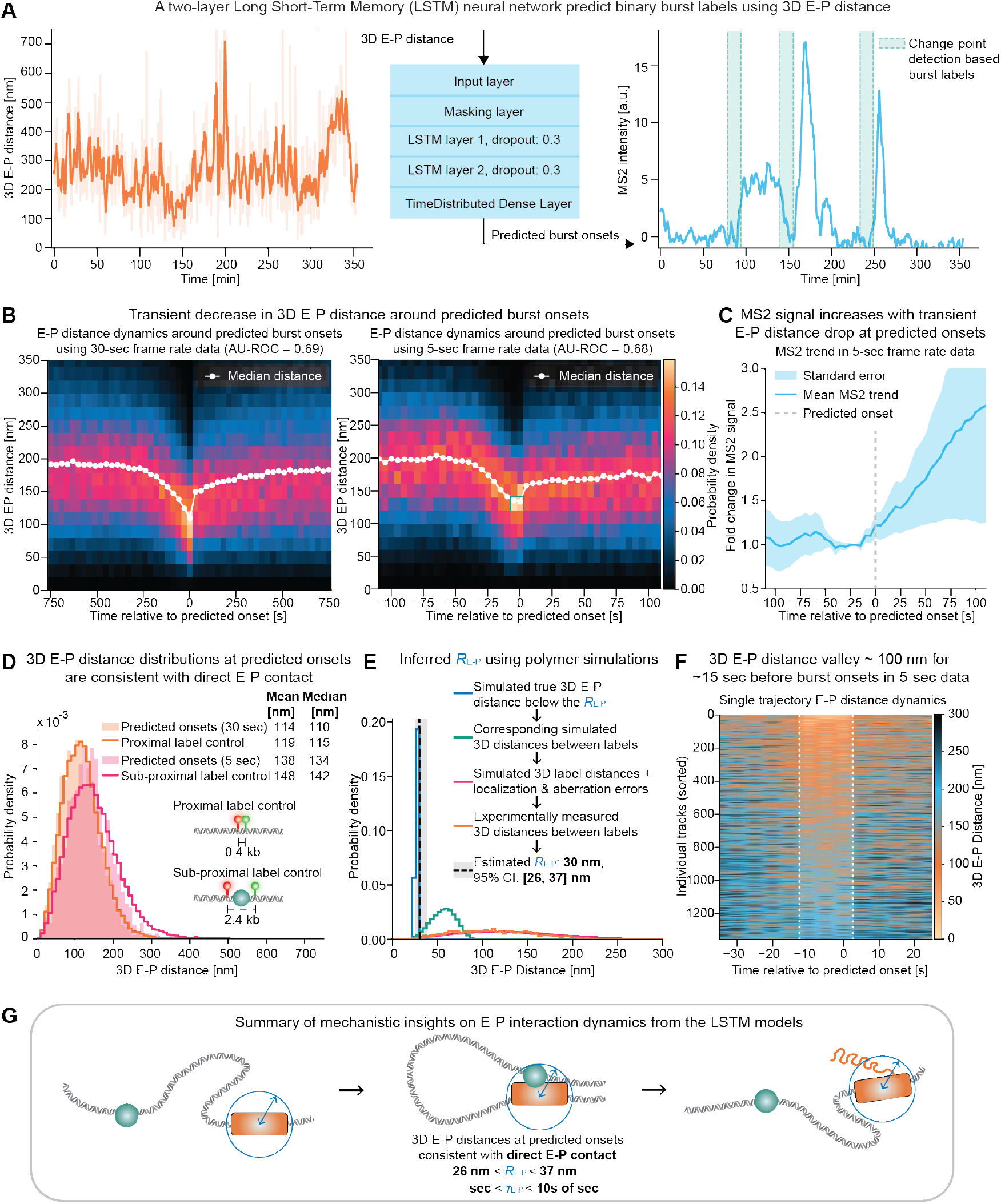
Estimating E-P interaction radius and duration using Long Short-Term Memory (LSTM) models. (**A**) Architecture of the two-layer LSTM neural network designed to predict binary transcriptional burst labels using 3D E-P distance as input. (**B**) Analysis of 30-second frame rate (left) and 5-second frame rate (right) data reveals a transient decrease in 3D E-P distance at predicted burst onsets, with the decrease in E-P distance in 5-second data lasting 10 seconds (2 frames, highlighted in a teal box). All live-cell trajectories from both the untreated and ΔRAD21 conditions of 339noC, 253noC, 170noC, 87noC, the untreated condition of 339CE and 339CP, the ΔRAD21 and ΔCTCF conditions of 339CECP, were used for both the 30-second and 5-second (if available for 5-second) data analysis. (**C**) Mean fold-change in MS2 signal relative to pre-onset baseline level aligned to the predicted burst onset time in the 5-second dataset. (**D**) The 3D E-P distance distributions at predicted burst onsets closely mirror those of the proximal (400 bp DNA separation) and sub-proximal (2400 bp DNA separation) label controls for the 30-second and 5-second datasets, respectively. (**E**) Convolution of simulated true E-P distances with localization and aberration errors, performed by scanning a fine grid of hypothetical *R*_E−P_ values ranging from 20 to 200 nm, yields an optimal *R*_E−P_ estimate of ∼30 nm. Displayed here is an illustration of the convolution process for the estimated *R*_E−P_ of ∼30 nm. (**F**) A heatmap of *N* = 1, 355 individual 5-second frame rate trajectories, sorted by mean 3D E-P distance during the 15-second window prior to the predicted burst onset (including the onset frame), highlights a distinct ∼100 nm E-P distance ‘dip’ for roughly 15 seconds prior to predicted burst onsets. (**G**) Summary of *R*_E−P_ and *τ*_E−P_ estimates derived from the LSTM analysis.

Applying the trained LSTM model to live-cell imaging trajectories, we achieved an Area Under the Receiver Operating Characteristic (AU-ROC) of ≈0.69 for both the 30-second and 5-second data (**Fig. S16A**), and found a pronounced transient decrease in the 3D E-P distance precisely at the predicted burst onsets (**Fig. 3B**), validating that the identified bursts are coupled to 3D E-P distance dynamics. Further confirming this relationship, aligning the corresponding MS2 trajectories to these predicted onsets reveals a sharp, corresponding increase in the mean MS2 signal (**Fig. 3C, Fig. S16B**). As additional validation, the predicted burst onset frequency showed a positive linear correlation with the BFP expression levels across conditions (*R*^2^ ≈ 0.9, **Fig. S16C**). To test the robustness of our temporal alignment and confirm feature dependence, we introduced a synthetic misalignment positive control. Artificially shifting the input 3D E-P distance by -4 frames resulted in an identical shift in the transient E-P distance dip relative to the predicted burst onsets, while maintaining predictive power with an AU-ROC of 0.68 (**Fig. S16D**). Conversely, as a negative control, feeding the LSTM scrambled, randomized E-P distances completely abolished the model’s ability to predict burst onsets (**Fig. S16E**), confirming that E-P distance dynamics, rather than random distance fluctuations or alignment artifacts, underlie the model’s predictions. We further validated our LSTM neural network against simulated ground truth *R*_E−P_ of 40 nm, 100 nm, and 150 nm, respectively, and demonstrated its ability to distinguish a small *R*_E−P_ of 40 nm from hypothetical larger *R*_E−P_ of 100 nm and beyond (**Fig. S17**).

Next, we took two approaches to estimate *R*_E−P_ from the observed dip in 3D E-P distances before burst onsets (**Fig. 3B**). We note that measured 3D distances will always be substantially greater than true distances because of the offset between the E-P pair and DNA labels, localization uncertainty (**SI Note 1**; (*1, 8, 40*)), and other noise sources. To account for this, we turned to our two co-localization control cell lines, a proximal label control (0.4 kb label separation) and a sub-proximal label control (2.4 kb label separation, **Fig. S3**). Comparing the 3D distance distributions at predicted burst onsets to these co-localization controls revealed them to be almost indistinguishable (**Fig. 3D**). Specifically, the 30-second predicted onsets mirrored the proximal label control (median distances of 110 nm and 115 nm, respectively), while the 5-second predicted onsets aligned with the sub-proximal label control (median distances of 134 nm and 142 nm, respectively). Thus, if we interpret the co-localization controls as direct contact, the observed dip in the 3D distances before burst onsets effectively estimates *R*_E−P_ ≈ contact.

As a second approach, we attempted to “deconvolve” the measured 3D distances from known noise sources. Specifically, we isolated the interacting sub-populations of E-P conformations within our polymer simulations by scanning different *R*_E−P_ values to serve as the upper cut-off of the true E-P distance distribution. After extracting the corresponding simulated label distance distribution and convolving it with localization errors (estimated via MSD fitting, **Table S6**) and aberration correction errors, we found that an *R*_E−P_ of ∼ 30 nm (95% CI: 26 to 37 nm) best reproduces the experimentally observed label distance distribution at the 30-second predicted burst onsets (**Fig. 3E**).

To also estimate *τ*_E−P_, we used the higher temporal resolution 5-sec trajectories where we observed a clear E-P distance ‘dip’ to ∼120 nm for approximately 10 seconds prior to the predicted burst onset (**Fig. 3B**). To confirm that this distance dip is not merely an effect of population averaging, we visualized the E-P distance over time for individual trajectories as a sorted heatmap, revealing a ∼15 seconds E-P distance dip at the single-trajectory level (**Fig. 3F**). The observed E-P contact duration of ∼15 seconds is much longer than the duration expected for purely diffusive contacts (*53*) and instead points to affinity-mediated stabilization of E-P interactions resulting in extended E-P interaction durations, which is also consistent with the observed ‘E-P loop dot’ in the RCMC maps even in the absence of CBSs (**Figs. 1F and 2A**).

In summary, our third approach estimates *R*_E−P_ ≈ contact based on the experimental co-localization controls whereas simulation-based deconvolution of noise estimates *R*_E−P_ ≈ 30 nm (95% CI: 26 to 37 nm). Comparing against simulations where the ground truth is known, we find that the LSTM tends to slightly overestimate *R*_E−P_ (**Fig. S17B**) suggesting that *R*_E−P_ ≈ 30 nm is more likely to be an upper bound. Simple dip quantification estimates this observed interaction duration to be ∼15 seconds. Given the significant uncertainty inherent to this analysis, we conservatively estimate the E-P interaction duration *τ*_E−P_ to fall within the range of seconds to tens of seconds (**Fig. 3G**).

### Estimate 4: *τ*_E−P_ through a minimal model of E-P-mediated activation with VEPI

To complement our ML-based analysis of live-cell trajectories and provide another estimate of *τ*_E−P_, we employed Bayesian inference to infer a minimal mechanistic model of E-P-mediated transcription activation. We developed Variational Enhancer-Promoter Inference (VEPI), a method that infers transcription events from E-P and MS2 trajectories, while parameterizing an end-to-end model of E-P-driven transcription ensembles (**Fig. 4A**). In contrast to previous methods (*60, 61*), VEPI relies on a path-space variational inference-based approach to MS2 inference, which infers promoter states concurrently with deconvolving MS2 into RNA Polymerase II (Pol II) loading events. The method explicitly models the E-P localization noise, MS2 noise, and any constant offsets present in the MS2 data.

**Figure 4:**
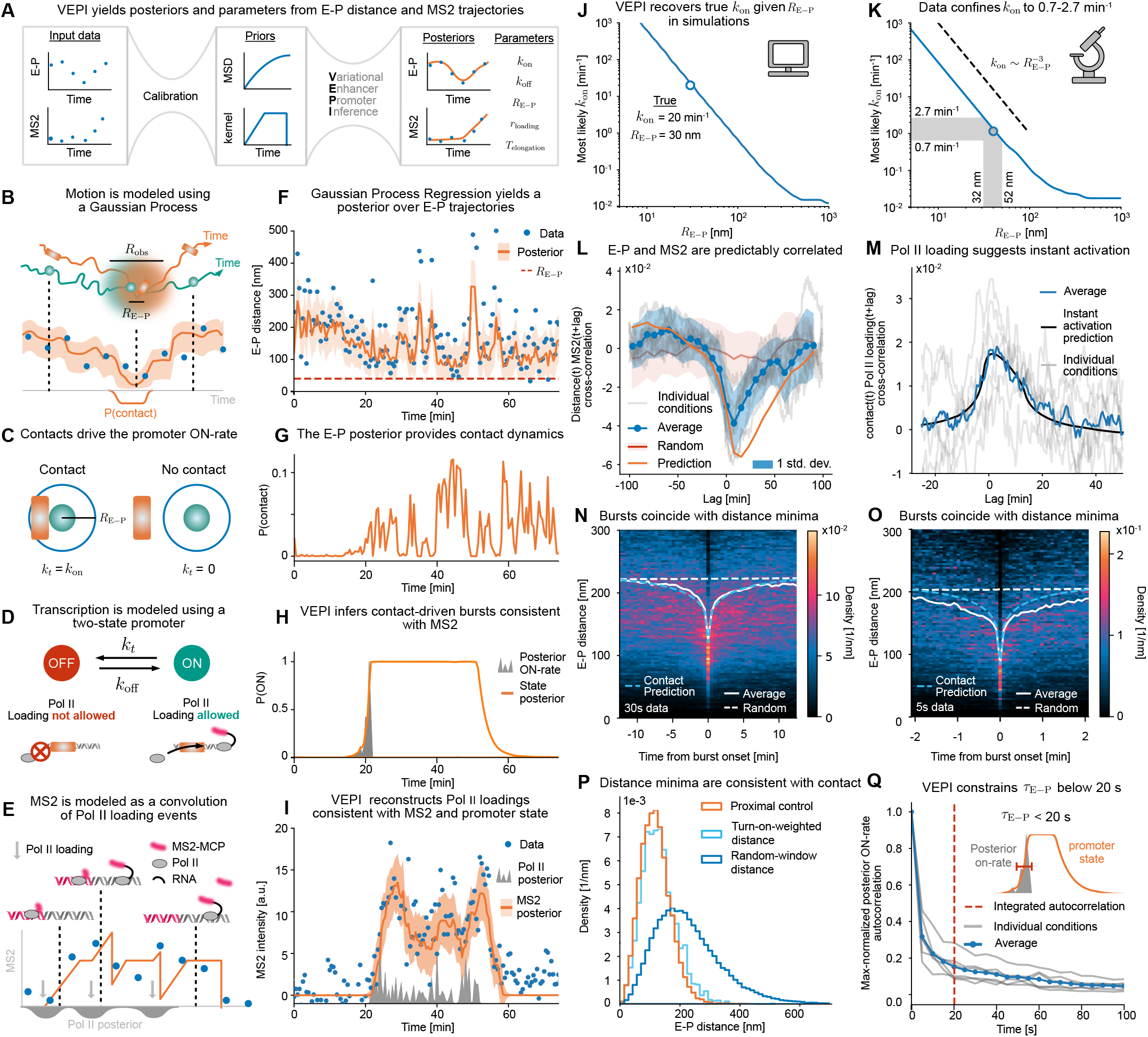
Inferring E-P interaction duration using Variational Enhancer-Promoter Inference (VEPI) (**A**) VEPI workflow (**B**-**E**) Model assumptions of VEPI. (**F**) Gaussian Process Regression (GPR) posterior. Shaded orange region and line represent 95% confidence intervals and median across the posterior ensemble of trajectories, respectively. (**G**) Contact probability from the GPR posterior and *R*_E−P_ = 40 nm. (**H**) Posterior distribution of promoter state. (**I**) Posterior distribution of Pol II loadings and MS2. Shaded gray represents the density of Pol II loadings across the posterior ensemble. Average MS2 (orange curve) and ± 1 standard deviation (shaded orange) were obtained from 10,000 posterior Pol II loading samples. (**J**) Most likely *k*_on_ for a given *R*_E−P_ in simulated data. **K**) Most likely *k*_on_ for a given *R*_E−P_ using 30 s imaging of 87noC, 170noC, 253noC, 339noC, 339CE, 339CP, and 339CECP. (**L**) Cross-correlation of E-P distance and MS2. Random curve (red) was obtained by pairing each E-P track with a new MS2 that is closest in duration to itself. Conditions as in (K) except 339CECP. Prediction (orange curve) was obtained as an average of per-condition simulations containing 1000 trajectories each using inferred *k*_on_, *k*_off_, per-track loading rates and noise, kernel parameters, and MSD parameters. (**M**) Cross-correlation of contact and Pol II loading. Conditions as (K) excluding 339CECP. Instant activation prediction was obtained as an average of per-condition predictions by convolving the contact and p(on) autocorrelation function shifted by the inferred loading time. (**N, O**) Distribution of E-P distance windows weighted by posterior rate of turning on at the center. Random is the same as in (L). (**P**) Histogram of weighted distances at the center time in 30 s data of N. (**Q**) Posterior ON-rate autocorrelation. *τ*_E−P_ was estimated as the integral of the curve from 0 to 3500 s. Conditions were as (K) excluding 339CECP. All cross-correlations were done on per-track mean-centered and per-track standard deviation-scaled data. For autocorrelations, no scaling was applied, and the per-condition mean was subtracted for each track.

We modeled E-P dynamics as a Gaussian Process (GP), allowing us to infer the hidden motion of the E-P pair between frames, below the localization error (**Fig. 4B**). For modeling transcription, we sought the simplest possible model consistent with bursty transcription amenable to a contact-driven activation. We therefore chose a two-state random telegraph process with a contact-driven ON-rate (**Fig. 4C**) where polymerases could only load in the ON-state (**Fig. 4D**). Following previous work, MS2 was modeled as a sum of deterministic contributions from each Pol II loading in the trajectory (*60, 62*) (**Fig. 4E**).

VEPI models the link between E-P distance and transcription through an effective time-dependent ON-rate of the promoter given by *k*_on_*P* (contact), where *P* (contact) can be computed using Gaussian Process Regression on the E-P trajectories (**Fig. 4F, G**) (*63*). We then fed this upstream drive into our MS2 inference method to infer promoter state transitions and polymerase loading events (**Fig. 4H, I**). To validate VEPI, we ran simulations across a range of signal-to-noise ratios, transition rates, measurement intervals, and number of trajectories. We found that we are able to call bursts with an AUC consistently above 0.9, Pol II loadings with an average error of less than one polymerase, and infer transition rates to within 80% of their true value (**Figs. S18 and S19**).

We found that VEPI robustly recovers the functional relationship between *k*_on_ and *R*_E−P_ in simulations (**Fig. 4J**). However, we found that *R*_E−P_ and *k*_on_ could not be identified independently in the regime of small contact radii, likely due to the time-scale separation employed during inference (**Fig. S20B**). For this reason, we constrained *R*_E−P_ based on our previous estimates and inferred the remaining parameters using VEPI. Our analysis revealed *k*_on_ to lie in the range 0.7-2.7 min^-1^. (**Fig. 4K, Table S9**). To validate the resulting parameterized model, we plotted the crosscorrelation between E-P distance and MS2 intensity. Strikingly, the data display a clear dip in cross-correlation, confirming spatiotemporal coupling between E-P proximity and transcription (**Fig. 4L**). We tested the ability of our minimal model to reproduce this relationship by simulating from the inferred parameters. While the rise after contact was predicted slightly broader than the data (within 2*σ*), the negative lag cross-correlation and magnitude of the dip at lag 0 were predicted to within 1*σ* of the data (**Fig. 4L**).

Having parameterized the model, we next analyzed the posterior reconstructions of E-P-driven transcription activation. The cross-correlation between contact and posterior rate of Pol II loading was consistent with activation starting immediately after contact, given the temporal resolution of our data (**Fig. 4M**). Further investigating the dynamics around activation, we computed E-P distance windows from the trajectories weighted by the posterior probability of a burst onset at their center. We found E-P dips consistent with single-frame contact in 30 s data while 5 s data displayed a single-frame dip surrounded by a broader region of lower distances (**Fig. 4N,O,P**). We hypothesized that this broadening might be due to extended contacts that were averaged out due to the uncertainty in burst onset detection. To investigate this more directly, we reasoned that the width of peaks in the posterior ON-rate would provide an upper bound on the duration of the activating interaction. Plotting the autocorrelation of the posterior ON-rate, we found a narrow window of correlation with a characteristic timescale of around 20 s (**Fig. 4Q**). This provides an upper bound on *τ*_E−P_ as the autocorrelation of posterior ON-rate will be broadened by localization error, finite timeresolution, and uncertainty in Pol II loadings coming from noise in MS2.

In summary, using VEPI and our *R*_E−P_-estimates, we find the E-P trajectories to be consistent with a two-state promoter model driven by contact-gated activation with a rate of activation when contacting in the interval 0.7-2.7 min^-1^, consistent with transient contact-driven promoter activation. We stress that while agreement with data was found to be satisfactory, the model omits several properties of the system, including the possibility of multiple promoter states, randomness in splicing and Pol II dynamics, and attraction or loop extrusion in the polymer dynamics. Despite its shortcomings, the model captures the cross-correlation seen in the data and allows us to obtain an orthogonal estimate of *τ*_E−P_, constraining it to lie below 20 s.

### Estimate 5: *R*_E−P_ and *τ*_GATE_ from cohesin and CTCF perturbation experiments

Our complementary estimates consistently place *R*_E−P_ within the ∼20–50 nm range, which matches the ∼40– 50 nm molecular size of the cohesin complex (*64*). This physical alignment predicts that E-P-mediated transcription might be particularly sensitive to perturbations of cohesin-extrusion (*65–69*) mediated interactions. To test this, we acutely depleted cohesin subunit RAD21 (**Fig. 1D and 5A**) and performed llsSRLCI. Remarkably, cohesin depletion reduced mean nascent transcription (MS2) by ∼80% to 97% across all distal E-P pairs after just 2 hours of depletion (BFP quantification not feasible due to the long half-life of BFP). Strikingly, transcription was abolished for the 339 kb E-P pairs (both with and without convergent CBSs): their post-depletion MS2 signals matched the control cell line lacking the synthetic promoter (**Fig. 5A**).

**Figure 5:**
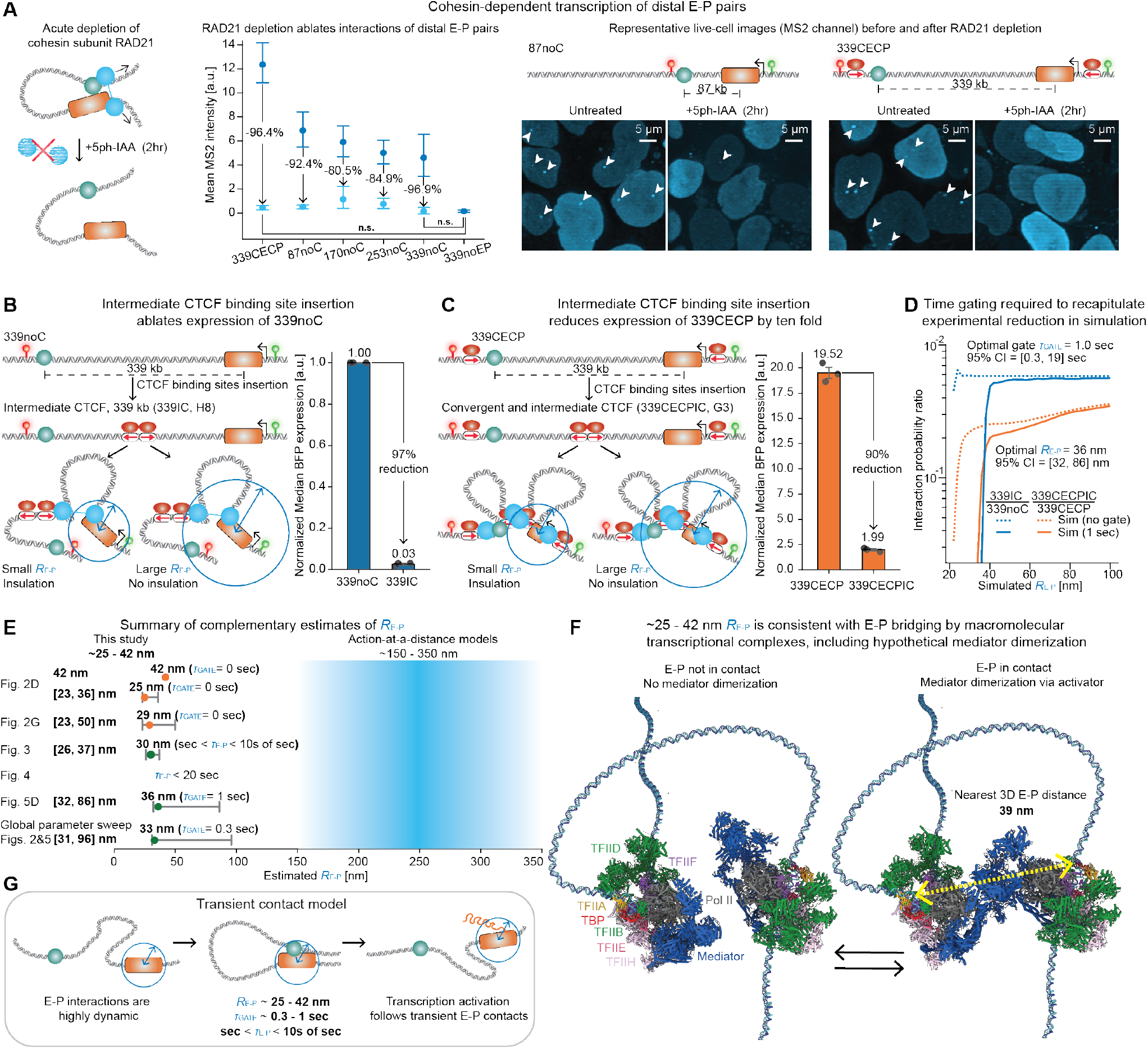
Complementary estimates converge to an E-P interaction radius of tens of nanometers. (**A**) Acute depletion of RAD21 via 5ph-IAA (left panel) ablates transcription of distal E-P pairs, resulting in a near-complete reduction (80.5% to 96.9%) in mean MS2 intensity across multiple E-P configurations (middle panel). Representative live-cell MS2 (MCP-2x-Halo-JFX650) images are shown before and after depletion (right panel). (**B**) Insulation via the insertion of a pair of intermediate, divergently oriented 3x CBSs drastically reduces BFP expression (97% reduction) for 339noC, consistent with a small *R*_E−P_. Error bars represent the standard error of the mean (SEM) across three biological replicates. (**C**) A similar intermediate CBS insertion in 339CECP also strongly reduces expression (90% reduction), implying small *R*_E−P_ underlying the observed insulation. The displayed median BFP expression values were also normalized to 339noC (**Fig. S4**) for consistency. Error bars represent the SEM across three biological replicates. (**D**) Comparing the simulated time-gated interaction probability ratios (339IC/339noC and 339CECPIC/339CECP) to their experimental expression ratios in (B) and (C) yields an optimal *τ*_GATE_ ≈ 1.0 second (95% CI: 0.3–19 seconds) and an optimal *R*_E−P_ ≈ 36 nm (95% CI: 32–86 nm; **Fig. S25A**). (**E**) Summary comparison of the complementary *R*_E−P_ estimates derived in this study, with an average *R*_E−P_ of ∼ 32.5 ± 2.4 nm (mean ± s.e.m.), markedly smaller than previous models suggesting action-at-a-distance. (**F**) Structural modeling indicates that an estimated *R*_E−P_ of ∼ 25–42 nm is biochemically consistent with direct enhancer-promoter contact. The schematic illustrates a hypothetical model of E-P contact achieved by Mediator dimerization via activator. This composite model utilizes human preinitiation complex (PIC) structures (*31*), with the central dimerization interface modeled by aligning RNA Polymerase II and replacing the human Mediator with an activator-bound yeast Mediator dimer (*109*). In the contact state, the nearest 3D distance between the DNA segments traversing the two PICs measures ~39 nm (yellow dashed arrow). (**G**) An emergent transient contact model of transcription activation, demonstrating that E-P interactions are highly transient and trigger transcription only when the physical distance reduces to within a compact interaction radius (*R*_E−P_ ≈ 25–42 nm) for a minimum duration (*τ*_GATE_ ≈ 0.3–1 s).

Consistent with the observed ∼10-20 second E-P distance dip (**Figs. 3B,F and 4N,O,Q**), the near-complete ablation of transcription in the 339 kb E-P pairs following RAD21 depletion further points to a potential “activation timescale” or “time gate”, where E-P encounters briefer than the time gate are filtered out, akin to what has been observed for inducible budding yeast promoters (*70, 71*). Specifically, the post-depletion median E-P distance of ∼454 nm (**Fig. 1G**) combined with an MSD exponent of ∼0.37 and estimated prefactor Γ ≈ 0.015 *µ*m^2^*/*s^0.37^ (**Tables S6 and S7**), predicts a median first passage time (FPT) on the order of minutes (*53*). The presumably relatively frequent brief encounters leading to little transcription suggest these sub-diffusive encounters might be too brief to trigger transcription activation, compared with cohesin-mediated encounters which may last seconds or longer (*68, 72*).

To simultaneously test the time gating and small *R*_E−P_ hypotheses, we first inserted a pair of 3xCBSs in divergent orientations in the middle of the 339 kb E-P pairs without convergent CBSs (339noC) (**Fig. 5B**). In terms of *R*_E−P_, this insertion should only affect transcription if *R*_E−P_ is very small; if *R*_E−P_ was large, the E-P pair would still be brought within the *R*_E−P_ sphere despite cohesin stalling at the intermediate CBSs (**Fig. 5B**). In terms of time-gating, the inserted pair of 3xCBSs would be expected to largely eliminate all cohesin-bridged E-P interactions (*68*) but not diffusive encounters. Strikingly, we observed an almost complete loss of transcription (97% as measured by BFP level) upon 3xCBS pair insertion. This can only be explained if *R*_E−P_ is very small and is consistent with time-gating.

Next, we performed a similar 3xCBS pair insertion in the 339CECP cell line, which harbors convergent CBSs proximal to the E-P pair (**Fig. 5C**). Prior studies have suggested a “loop stacking” model, predicting that the formation of a CTCF rosette by such CBS insertion would increase expression (*73*). On the other hand, a CTCF rosette would prevent direct single cohesin-bridged E-P interactions and enforce larger E-P distances, thereby decreasing expression if *R*_E−P_ is very small and if productive interactions are time-gated (**Fig. 5C**). We observed strongly reduced expression (90% as measured by BFP level). These results argue against the loop stacking model (*73*) and are consistent with *R*_E−P_ being very small.

To quantitatively explore these effects, we turned to experimentally constrained 3D polymer simulations. In the simplest model without time-gating, where expression is proportional to interaction probability, CBS insertion could maximally confer a ∼65% reduction in interaction probability for the 339 kb E-P pair without convergent CBSs, much lower than the 97% experimental reduction in BFP expression (**Fig. 5D**). We could only match the experimentally observed reduction by incorporating a time gate, *τ*_GATE_, where interactions briefer than *τ*_GATE_ cannot activate transcription (**Fig. 5D**). We performed a systematic sweep (**Fig. S25A**) of simulation parameters that could explain the data. The best fit parameters yielded a minimum required interaction duration, or time gate (*τ*_GATE_), of ∼1.0 second (95% CI: 0.3 to 19 seconds) and an *R*_E−P_ ≈ 36 nm (95% CI: 32 to 86 nm) for transcription activation. The requirement for time-gating also led us to re-visit the polymer simulations from our first and second estimates (**Fig. 2**) of *R*_E−P_. By combining all nine cell lines with different E-P genomic distances and CBS configurations from **Figs. 2 and 5**, we found a globally optimal *τ*_GATE_ of ∼0.3 second (95% CI: 0.2 to 19 seconds) and a globally optimal *R*_E−P_ of ∼33 nm (95% CI: 31 to 96 nm) (**Fig. S25B, Fig. 5E**).

In summary, our perturbation experiments can only be explained by incorporating time-gating, and our fifth and final estimate of *R*_E−P_ is ∼36 nm (**Fig. 5D**).

## Discussion

Despite the discovery of mammalian enhancers in 1981 (*6, 7*) and their central role in gene regulation, the mechanisms of E-P looping interactions in space and time remain heavily debated (*1,2,8–11*). This lack of consensus is partly for technical reasons: it is extremely challenging to simultaneously observe E-P interaction dynamics and nascent transcription in real time in living cells with nanometer precision, and even modest noise can easily obscure molecular contact (**Fig. 1B; SI Note 1**).

To overcome these limitations, we have engineered a synthetic biology platform specifically designed to dissect E-P interaction mechanisms. Using this system, we developed five complementary approaches to estimate *R*_E−P_, *τ*_E−P_, and/or *τ*_GATE_ (**Fig. 5E**). Although these methods vary in their directness and carry distinct limitations, their underlying technical assumptions are largely orthogonal. Strikingly, however, all estimates converge on the same conclusion: functional E-P interactions occur at the scale of *R*_E−P_ ≈ 25–42 nm and therefore likely involve direct contact. Specifically, we found an average *R*_E−P_ of 32.5 ± 2.4 nm (mean ± s.e.m.) for point estimates of *R*_E−P_ (**Fig. 5E**).

While the specific protein complexes involved in E-P looping remain to be fully defined, to put our *R*_E−P_ ≈ 25–42 nm estimates into molecular context, we performed structural modeling of a putative Mediator-Pre-initiation Complex (PIC) dimerization model of E-P looping (*30–32, 74*), which arrives at a very similar nearest DNA-DNA distance of ∼39 nm (**Fig. 5F**). However, we emphasize that although our estimates (**Fig. 5E**) provide strong evidence of direct physical bridging between E-P bound complexes, they do not allow us to determine the molecular components, and Mediator dimerization represents just one of many plausible molecular models. Indeed, an *R*_E−P_ of ∼25–42 nm also matches the size of active transcription foci visualized by electron microscopy (*75*), the size of clusters of transcription regulatory factors (*76, 77*), and the size of the cohesin complex (*64*).

Although action-at-a-distance models appear supported by several observations (*1–3,8–11,19,20,28,29,33–36,38*), they are challenging to reconcile with selectivity in gene regulation in the nucleus. First, if enhancers can activate promoters across distances of ∼200–300 nm, CTCFmediated insulation would be largely ineffective, which directly contradicts our observations here (**Fig. 5B-D**) and many prior studies (*44, 57, 78–82*). Second, CTCF-mediated E-P facilitation similarly cannot be explained by simple isotropic action-at-a-distance models (*1*). Insertion of convergent CBSs proximal to the E-P pair (from 339noC to 339CECP) only modestly reduces the median 3D distance from 246 to 167 nm (**Fig. 1G**), but dramatically increases expression by ∼20-fold (**Figs. 1E and 2E-F**). CTCF-mediated facilitation of E-P interactions has also been reported in prior studies (*5, 68, 83–85*). Third, the gene density in the mammalian nucleus is high: ∼300–400 genes per µm^3^ (*1*). Thus, if hypothetically *R*_E−P_ ≈ 300 nm, the activation sphere of a single enhancer (volume ≈ 0.11 µm^3^) would simultaneously encompass ∼40 genes, presumably leading to widespread off-target activation. Reciprocally, the human genome’s ∼1.8 million candidate enhancers (*86*) results in an astonishing density of ∼22,000– 29,000 enhancers per µm^3^. This means that any given promoter would have roughly 3,000 candidate enhancers residing within its 300 nm radius sphere. Even if only a small fraction of these enhancers are active in a given cell type, every promoter would be constantly enveloped by dozens to hundreds of functional enhancers, making selectivity challenging to explain.

While we propose a transient contact model based on short-lived physical bridging between macromolecular complexes as a general mechanism for E-P interactions (**Fig. 5G**), we note that condensate models were originally proposed specifically for super-enhancers (*3*), large clusters of enhancers that often span 10–30 kb (*42*), where cooperative interactions among enhancer segments eventually lead to phase separation (*87*). Indeed, we have for technical reasons (we cannot reliably assign a single (*x,y,z*)-coordinate to a ∼25-kb sized enhancer) studied a compact *<*2kb-sized enhancer here and our findings thus remain consistent with action-at-a-distance for large super-enhancers as well as contact-scale E-P proximity within a condensate.

Along with a stringent *R*_E−P_ ≈ 25–42 nm in space, our results also point to short-lived stabilization of E-P interactions in time. We observe that these functional E-P interactions last for ∼15 seconds (**Figs. 3B,F and 4N,O,Q**), though we emphasize that this observed duration is associated with substantially higher uncertainty and we conservatively place *τ*_E−P_ between seconds and tens of seconds (**Fig. 3G**). Thus, our *τ*_E−P_ estimate is consistent with a transient contact model (“hit-and-run”) for E-P interactions, and inconsistent with very stable E-P loops in the absence of CBSs. Nevertheless, our *τ*_E−P_ estimate points to stabilization above and beyond purely diffusive contacts which would be much more transient (*53*). Presumably, this stabilization is due to affinity-mediated interactions between transcriptional proteins bound at both the enhancer and the promoter (**Fig. 5F**). In this context, we note that our *τ*_E−P_estimate matches the residence time of many transcription factors and components of the general transcriptional machinery, which is also on the order of seconds to tens of seconds (*88–96*). Stabilized but short-lived E-P interactions make two further predictions that are corroborated by 3D genomics data. First, most E-P interactions should be visible as a loop/”dot” in high-resolution 3D genomics experiments, as observed here (**Figs. 1F and 2A**) and previously (*17,18,52,97–101*). Second, stabilized but short-lived E-P interactions predict that most E-P loops without nearby CBSs should be largely robust to acute depletion of CTCF and cohesin, as indeed also observed here (**Fig. S5**) and previously (*52, 97, 98, 102*).

Finally, only by adding time-gating to our E-P model can we explain the perturbation experiments in **Fig. 5B-D**. By time-gating, we refer to the ability of the promoter to filter out E-P interactions that are too brief and only respond to sufficiently long-duration E-P interactions. Time-gating has previously been observed for inducible budding yeast promoters and explained through multi-state promoter models (*70, 71, 103*) and has been hypothesized in the context of mammalian E-P regulation (*1, 51, 68, 72, 104*). Our direct observation of stabilized but short-lived E-P interactions on the order of ∼15 seconds thus provides evidence consistent with time-gating models. Through modeling, we estimate *τ*_GATE_ ≈ 0.3-1 second for our synthetic E-P pair, though we emphasize that this estimate is associated with substantial uncertainty. This implies that more transient E-P encounters are filtered out and transcriptionally unproductive. Indeed, recent chromatin dynamics studies have suggested that frequent subdiffusive contacts between enhancers and promoters within hundreds of kb and hundreds of nanometers are both frequent and unavoidable (*53*). Thus, time-gating may be essential for E-P selectivity: if non-cognate E-P contact events are more transient (≪1 second) they would be filtered out and only stabilized (≥ 1 second) cognate E-P contacts would result in transcriptional activation. Time-gating also explains the unique gene regulatory importance of cohesin (**Fig. 5A**). Recent work (*68*) proposed that cohesin-bridged E–P encounters are primary drivers of transcription. Because such bridges can have a small contact radius and longer duration (*72*), they provide attractive candidates for the contacts selected by promoter time-gating. We suggest that time-gating, and perhaps kinetic proofreading as previously proposed (*105–107*), are likely important to understanding E-P-mediated gene regulation and will be important to further explore in future work.

Lastly, we end by noting key limitations of our study. First, we have studied only one specific E-P pair in mESCs. Future studies will be required to test the generality of our findings for other E-P pairs and in other cell types. Second, we have taken a synthetic biology approach. This enabled us to comprehensively dissect spatiotemporal E-P interaction mechanisms in a ‘clean’ genomic region, without redundancy between dozens of enhancers and promoters nearby. However, many endogenous genes are regulated by multiple enhancers and future studies will be required to understand how multiple enhancers and promoters interact. Indeed, while expression was strongly cohesin-dependent in our system, genomic regions with redundant enhancers tend to have reduced cohesin-dependence (*52, 69, 102, 108*). Third, although our five estimates are all consistent with the transient contact model for E-P interactions, we note that our estimates of *R*_E−P_, *τ*_E−P_, and *τ*_GATE_ are all approximate. We anticipate that the experimental and computational methodologies developed in this study may facilitate the study of other E-P pairs in the future.

## Acknowledgments

We thank Seychelle Vos for discussions, advice, and help with the structural biology modeling for Figure 5F. We thank Luke Lavis for generously providing Janelia Fluor dyes. We thank the Hansen, Zechner, and Mirny labs for discussions throughout this project. We thank Edouard Bertrand for kindly sharing the MS2 array plasmid used here. We thank Gerd Blobel for feedback on the manuscript. We thank Nadezda Fursova and Daniel Larson for discussions about MS2 imaging. We thank Neil Blackledge for discussions about intron selection. We thank Andrea Perry for early contributions to image processing.

## Author contributions

J.H.Y., H.D.P., J.T., M.K.H., L.M., C.Z., A.S.H. contributed to project design. L.M., C.Z., and A.S.H. supervised the project. A.S.H. coordinated the overall project. J.H.Y., M.K.H., A.S.H. designed the synthetic biology platform. M.K.H. identified the genomic region, designed the synthetic reporter, and tagged RAD21 with the mAID degron. J.H.Y. performed 27 gene edits to generate the 15 final cell lines. J.H.Y. designed and engineered the synBsr1 DNA labeling system, and C.C.K. validated the labeling system and built the fluorescent synBsr1-ParB construct. J.H.Y. designed and validated the synthetic enhancer and the engineered CTCF binding sites. J.H.Y. performed time course Western blot experiments to validate acute depletion of RAD21 and CTCF. J.H.Y. conducted FACS experiments to optimize expression levels of fluorescent binding proteins, and performed flow cytometry experiments to quantify tagBFP expression across cell lines. J.H.Y. and J.M.J. performed RCMC experiments. J.H.Y. developed the RCMC data analysis pipeline to quantify the relationship between expression and E-P interaction. J.H.Y. performed polymer simulations and developed the analysis pipeline. J.H.Y. and H.D.P. developed mathematical derivation for comparing RCMC and flow cytometry measurements. J.H.Y. developed the MS2 intensity correction method. J.H.Y. and H.D.P. performed Bayesian MSD fitting. J.H.Y. developed the CPD burst calling pipeline and the deep learning framework to predict burst onset using E-P distances. J.H.Y. developed four complementary analyses to infer E-P interaction radius and duration. A.S.H. provided supervision.

J.H.Y. cultured cells and performed all live-cell imaging experiments. J.T. developed the Fyrtarn library and applied these computational methods for processing the livecell imaging acquisitions to trajectories of distances and intensities. J.T. and J.H.Y. performed manual quality control on all live-cell trajectories using a user interface developed by J.T. A.S.H. provided supervision. J.T. developed the methodology for MS2 correlation function calibration and H.D.P. implemented it. Both were done under the supervision of C.Z. J.M.J. performed BILD analysis to estimate looping probability and lifetime. A.S.H. provided supervision. H.D.P. developed, implemented, and validated VEPI under supervision of C.Z. with input from L.M. H.D.P. performed cross-correlation analyses and MS2 correlation function fits. H.D.P. performed Rouse model and VEPI parameterized data simulations and analyzed inference outputs.

J.H.Y., H.D.P., J.T., and A.S.H. drafted the manuscript and all authors edited the manuscript.

## Data and materials availability

The raw and processed RCMC data generated in this study can be found at NCBI Gene Expression Omnibus under accession number GSE335363 at https://www.ncbi.nlm.nih.gov/geo/query/acc.cgi?acc=GSE335363. Trajectory data can be found on Zenodo at https://doi.org/10.5281/zenodo.20293143. The Fyrtarn library for yielding trajectories from lattice light-sheet acquisitions can be found at https://github.com/ahansenlab/Fyrtarn. The code for RCMC data analysis, polymer simulations and analysis, LSTM burst onset prediction, and Bayesian MSD fitting is available under different modules within the synEP repository at https://github.com/ahansenlab/synEP. The code to run Bayesian Inference of Looping Dynamics (BILD) on the data produced in this manuscript is available at https://github.com/ahansenlab/synEP_bild. The code to run VEPI is available at https://github.com/henrik-dahl-pinholt/VEPI Yang et al 2026.

## Materials and Methods

Materials and Methods are provided in the Supplementary Information.

