## Supplementary Information for "Live-cell imaging of enhancer-promoter dynamics reveals transient contact-driven gene activation"

#### Materials and Methods

**1 Data and code availability 4**

**2 Cell line generation, culture, and treatment conditions 4**

**14 Supplementary Figures** **66**

**15 Supplementary Tables** **92**

**16 Movie Captions** **98**

### 1 Data and code availability

The raw and processed RCMC data generated in this study can be found at NCBI Gene Expression Omnibus under accession number GSE335363 at https://www.ncbi.nlm.nih.gov/geo/query/acc.cgi?acc=GSE335363. Trajectory data can be found on Zenodo at https://doi.org/10.5281/zenodo.20293143. The Fyrtarn library for yielding trajectories from lattice light-sheet acquisitions can be found at https://github.com/ahansenlab/Fyrtarn. The code for RCMC data analysis, polymer simulations and analysis, LSTM burst onset prediction, and Bayesian MSD fitting is available under different modules within the `synEP` repository at https://github.com/ahansenlab/synEP. The code to run Bayesian Inference of Looping Dynamics (BILD) on the data produced in this manuscript is available at https://github.com/ahansenlab/synEP_bild. The code to run VEPI is available at https://github.com/henrik-dahl-pinholt/VEPI_Yang_et_al_2026.

### 2 Cell line generation, culture, and treatment conditions

#### 2.1 Cell culture

Mouse embryonic stem cells (JM8.N4 mESCs) [1] were cultured on plates pre-coated with 0.1% sterile gelatin solution (Sigma-Aldrich, G1890-100G) under feeder free conditions in a base medium consisting of KnockOut DMEM (Thermo Fisher, 10829-018) with 15% FBS (HyClone, SH30396.03, Lot. No. AE28209315) and 1000U/mL LIF (Cell Guidance Systems, GFM200-1000), 1 mM MEM Non-Essential Amino Acid Solution (Thermo Fisher, 11140-050), 2 mM GlutaMAX (Thermo Fisher, 35050-061), 100 µg/ml Penicillin-Streptomycin (Thermo Fisher, 15140-122), 100 µg/ml Primocin (Fisher Scientific, NC9392943) and 0.1 mM 2-mercaptoethanol (Sigma, M-3148) supplemented with 2i (10 µM MEK inhibitor (Tocris, PD0325901) and 3 µM GSK inhibitor (Sigma, SML1046-25MG)), as well as 2 µM GSK-3484862 (TargetMol, T11469). mESCs were fed daily by replacing half the medium and passaged every two days using TrypLE Express Enzyme (1X, Thermo Fisher Scientific, 12605028) for dissociation.

#### 2.2 Design of synBsr1 system for labeling the synthetic enhancer

To enable fluorescent labeling of the enhancer, we repurposed the Bsr1-a plasmid partition (Par) system in *Burkholderia cenocepacia* J2315 [2]. In the endogenous context of Par systems, the DNA-binding protein (ParB) interacts with the centromere (*parS*) to create a partition complex to facilitate plasmid copy segregation ahead of cell division [2]. Local spreading of ParB proteins after binding to *parS* sequences achieves high level of ParB enrichment [3, 4]. This property enables high signal-to-noise fluorescent labeling of DNA locus tagged by *parS* sites by expressing ParB protein conjugated with fluorophores. We identified four Bsr1-a *parS* sites on *Burkholderia cenocepacia* J2315 plasmid pBCJ2315 (NC_011003) using the palindromic 14 bp sequence TTGGCTCGAGCCAA [2]. We extracted 7 bp upstream and 7 bp downstream flanking the 14 bp palindromic sequence (for a total of 28 bp) for all four sites: 92,279 - 92,306 bp (site 1), 92,379 - 92,406 bp (site 2), 92,432 - 92,459 bp (site 3), 804 - 831 bp (site 4). To engineer the synBsr1 *parS* array, we tiled each of these sites sequentially 4-5 times (5 times for site 1 and site 2, 4 times for site 3 and site 4), with 47 bp buffer sequence between adjacent sites, making up a total of 1331 bp (including 14 bp PCR overhangs on the two ends). The 47 bp buffer sequences between adjacent sites were unique and generated by SiteOut [5] to exclude potential transcription factor binding sites (**Fig. S1C**).

The 290 amino acid Bsr1-a ParB protein sequence was extracted from the UniProtKB database [6] using the identifier B4QN6_BURCJ, converted to DNA sequence, and then codon optimized using the GENEWIZ codon optimization tool for expression in *Mus musculus* (Mouse). The codon optimized coding sequence of Bsr1-a ParB, which we named synBsr1-ParB, was cloned into pASH400 expressing synBsr1-ParB-2x-mScarlet3-NLS from an L30 promoter, enabling fluorescent labeling of the enhancer tagged with synBsr1 *parS* array (**Fig. S1C**).

##### 2.2.1 DNA sequence of the 1331 bp synBsr1 array

The 1331 bp synBsr1 array developed in this study is composed of 14 bp PCR overhangs at two ends, 18 sites of the 14 bp conserved Bsr1-a *parS* and their variable 7 bp upstream and downstream flanking sequences, and the 47 bp spacer sequence without TF binding sites between the *parS* sites (**Fig. S1C**):

GTAATCTATTTGGT GCCGGTCTTGGCTCGAGCCAAGATCGGC TTTCATAGTTCCAATGCGCTGCGATTTTTGTCCAAACCTCATCAAAG T
GGGATCTTGGCTCGAGCCAACCCTTGG GGAATCGTAAATCCAAAGTGATTGTAAGGATGCCTAAGCACTTACAT ACCGGTCTTGGCTCGAGCCA
AGACTGTT AGTGTTACTAACGATAAAGACACAAACGTCGAAACGGATGCGATAAG GGAGAGGTTGGCTCGAGCCAAGGTGCCA TTCCCACATC
ATTTCACATATAGTGCTTCATCGGGCAGTACTCTGAG GCCGGTCTTGGCTCGAGCCAAGATCGGC TCCAGGATGGTAGCTGTGGGCTCCGTTTT
GGGCTAAGTTAAGTCCAG TGGGATCTTGGCTCGAGCCAACCCTTGG CCTAGTGGGACCGCTGTACAGGATAAACGTTAAGGCACGGTCTGCGT
ACCGGTCTTGGCTCGAGCCAAGACTGTT TAGATTTCTACAAGCCAGTCAGATTGCAAGTTGAGAAGTTATGCAGT GGAGAGGTTGGCTCGAGCC
AAGGTGCCA CCGGTGACAATCTTTAACATAGCATCCTCTAAGGCCTTATCCAATTC GCCGGTCTTGGCTCGAGCCAAGATCGGC TGAAAGACC
TGTAAATGCCCGATAAGCTGACTCGAGGTATTTGTGAG TGGGATCTTGGCTCGAGCCAACCCTTGG AGTGTATCAACGTAATGCTACTGATAGC
GCTCATACACATTTTGCAA ACCGGTCTTGGCTCGAGCCAAGACTGTT GGAACGTACAGATTGCTTTACCGGTGGACCGCTTTTTATCGAAACAA

GGAGAGGTTGGCTCGAGCCAAGGTGCCA CTAATAAAATAGGTAAATATTTATTAAAAATTGACGGGAATCCCCCG GCCGGTCTTGGCTCGAGCC
AAGATCGGC ATAACCGCTCGCGTTACGGTTTTCGTGTAAAGGTGAAGATTCAGCCT TGGGATCTTGGCTCGAGCCAACCCTTGG GGCGTGCTC
AATGGCCATAACGTTCCACGAGTTAATCGAACAGTTCA ACCGGTCTTGGCTCGAGCCAAGACTGTT CTCGCTAGCTACGAGCACTAGGGTTCGT
GTGTCTGAGTTCAATATTT GGAGAGGTTGGCTCGAGCCAAGGTGCCA GTATTCGTGAGTGACGGCGAGATCCTGATTATAGACTTAATTCACCT
GCCGGTCTTGGCTCGAGCCAAGATCGGC TGGTAATACACATTCATGGATTCTCCCACGCTTCGGAATCGCGGTAT TGGGATCTTGGCTCGAGCC
AACCCTTGG GACGTACAACGGTA

##### 2.2.2 The 870 bp synBsr1-ParB coding sequence

The 870 bp synBsr1-ParB coding sequence has been codon optimized for expression in *Mus musculus* (Mouse):

ATGGCCCCTAGTCTGGCGACTATCTCTCAGCTGAGGGGCGGCGAAGTGGGCAGCAGCCAAGGCGCCACCAAGTACGAGATCGGACAGACCTAC
GAGGTGCCTATCGGCAAGATCAAGAGCAACAGCGTGAACCCTAGGGCCATCTACACCGCGTCGGCCGTCAGCGAAATGGCCGAGTCCCTCACCGCT
AGGGGACAAGGACAGAGCGCTAGCGCCTACGTGGACGAGGCCGGCGACATTGTGCTGATCGATGGCGAGAGGAGGCTGAGGGGTGCTAGGGCCGCC
GGCCTGCCTACCCTGAGGGTGGAGATTAGGCCTAAGCCTGCTAGCGAGAGGGAGCTGTACGAGGAGGCTAGGGCCGCCAACGTGGAGAGGAAGGAT
CAGAGCCCTCTGGACGACGCCCTGAAGTGGAAGGAGCTCCTGTCTAGGAAGATCTACCCTACCCAAGTGGCCCTGGCCAAGGCCCTGAACCTGGGC
GAGGACCACGTGTCTAGGACCCTGAGCCTGGCTCAGCTGCCTTCTAGGATCGTGCAAGCCGCCGCCGAGTACCCTGAGCTGCTGAGCCTGAAGATG
CTGAACGCCATTAGGGAGTTCTGGGAGGTGAAGGGCGAGGAGGAGACCCTGGAGCTGGTGCTGGATGCGGCGAAAACCGGCATCGGCTACCGAGAT
GTTGCGGCACGCCGGAAGGCCGCCGCCAAGGGCACCGTGAAGAGGCCTAGGAGCACAAGGGAGCAGCTGAGCTTTAGGGGCGCCAAGGGCGAGTTC
AAGAGCTTCGAGGAGGACGGAAGGATCGAGCTGAAGCTGAAGGGCCTGGCCCCTGACGTGGCCGCCGAAATAAGCGAGAAGATCCTGGCCCTGTTC
CCTAAGGAG

#### 2.3 Genome-editing and cell line generation

Genome-editing was performed in JM8.N4 mESCs using CRISPR/Cas9 largely according to published procedures [7, 8], but with modifications. We designed sgRNAs using the CRISPOR tool [9]. For each insertion we designed 2-3 individual sgRNAs and cloned them into the Cas9 plasmid. We co-transfected a Cas9 plasmid (pSpCas9(BB)-2A-Puro (PX459) V2.0 [7]) encoding Cas9, the sgRNA, and a puromycin resistance gene, and a repair plasmid (DNA segment to be inserted typically flanked by ∼1000 bp of homology on the left and on the right) using Lipofectamine 2000 (Thermo Fisher Scientific, 11668030) according to a published protocol [10] with some modifications. Specifically, we dissociated the mESCs from the culture dish and counted 80,000 cells into an Eppendorf tube and centrifuged at 300 × g for 3 minutes. While the cells were being spun down, we prepared 1.06 µg of HDR template and sgRNA vector (with a molar ratio 6:1) in 5 µL of Opti-MEM (Thermo Fisher Scientific 31985062), mixed with 2.12 µL of Lipofectamine 2000 pre-mixed with 5 µL of Opti-MEM, and incubated at room temperature for 5 minutes. Then we resuspended the cell pellet directly in the DNA-Lipofectamine 2000 complexes to ensure a single-cell suspension, followed by a 5-minute incubation at room temperature. We then plated 50,000 cells onto a 150 mm dish and the remaining 30,000 cells onto a 100 mm dish. 24 hours after plating, we added puromycin to both plates to a concentration of 1 µg/mL. 24 hours after puromycin addition, the medium in both plates was completely replaced with fresh medium without puromycin. Cells were fed every 2 days thereafter. ∼7–8 days after transfection, cells in the 100 mm plates were dissociated using the TrypLE Express Enzyme (1X) and the genomic DNA was extracted using Quick-DNA Miniprep (Zymo Research, D3025) to validate genotyping primers generated using NCBI Primer-BLAST [11]. We genotyped using one primer external to one homology arm and the other primer either external or within the other homology arm to prevent amplification of any residual repair vector. ∼10–14 days after transfection, single colonies were picked inside the biosafety cabinet under a Lynx EVO stereomicroscope (Vision Engineering) using a 20 µL pipette tip, passaged into two 96-well plates. One of the two 96-well plates was used to extract crude DNA by dissociating cells with 50 µL squishing buffer (10mM TrisHCl pH 8, 1mM EDTA, 25mM NaCl) per well and incubating at 60 ℃ for 1 hour followed by 95 ℃ for 10 minutes after transferring the cell-buffer mixture to a 96-well PCR plate. Desired clones identified using a validated genotyping primer pair were further expanded, high-quality genomic DNA collected (Quick-DNA Kit, Zymo Research), clones were further verified using PCR with multiple primer combinations on high-quality genomic DNA, frozen down and used for downstream experiments after additional clone-specific validation. Periodic pathogen testing was performed and no contamination detected.

To study enhancer-promoter interactions in living cells, we chose a region on chromosome 2 from 178,200,000 to 179,600,000 (mm39) in mESCs since it satisfies the following criteria: 1) clean, without genes expressed in mESCs and containing very few CTCF binding sites (some were later deleted); 2) simple and lacks complex structures in Micro-C [12]; 3) large enough to accommodate an E-P pair with sufficiently large genomic separation (∼340 kb); 4) permissive to gene expression (**Fig. S1A**).

First, we tagged the C-terminus of RAD21 with mAID degron [8, 13, 14] with repair plasmid pASH267 (see **Table S1** for plasmids, associated sgRNA sequences and full details). Clone H1 that was homozygously edited was chosen to proceed with subsequent edits. Next, we tagged the C-terminus of CTCF with FKBP(F36V) degron [8, 14, 15] with repair plasmid pASH326 (see **Table S1** for plasmids, associated sgRNA sequences and full details). After identifying homozygously edited clones using genotyping primers P779 and P780 (**Table S2**), we performed western blotting of CTCF, and clone D12 was chosen for the next edit because it had the lowest "leaky" depletion of CTCF when no dTAG-13 (Fisher Scientific, 66-055) was added and the most complete depletion 3 hours after 500 nM of dTAG-13 was added.

Next, DNA encoding the fluorescently-tagged OR3 proteins was inserted via PiggyBac transposition using plasmid pASH323 (see **Table S1** for plasmids and full details: OR3 and ANCHOR3 systems were obtained from NeoVirTech), where OR3-$(GS)_3$-mStayGoldx2-P2A-T2A-BSD is expressed from an L30 promoter [8]. OR3 protein [8, 16] was C-terminally tagged with 2 copies of mStayGold [17], and the blasticidin resistance gene was expressed using the P2A-T2A linker [18]. After co-transfection of the payload plasmid pASH323 and a plasmid encoding Super PiggyBac Transposase with a molar ratio of 4:1 using Lipofectamine 2000, cells were plated onto 150 mm dishes at low density. 24 hours after plating, 4.8 µg/mL of blasticidin S (Sigma-Aldrich, 15205) [19] was added to select clones with stable genomic integration. 7 days later, the medium was replenished without blasticidin S, and single colonies were picked 3 days after drug removal. Clone F3 was chosen for subsequent edits.

To enable acute depletion of RAD21 with the inserted mAID degron tag, a DNA construct encoding OsTIR(F74G) [20] expressed from an EF1$\alpha$ promoter was next inserted into the TIGRE safe harbor locus [21] using repair plasmid pASH329 (see **Table S1** for plasmids, associated sgRNA sequences and full details). After identifying homozygously edited clones using genotyping primers P978 and P981 (**Table S2**), we performed western blotting to identify the clone that had the least depletion of RAD21 when no 5-Phenyl-1H-indole-3-acetic acid (5ph-IAA, BioAcademia, 30-003-10) was added and the most complete depletion 3 hours after 100 µM of 5ph-IAA was added. The clone H10 was chosen for the next round of genome editing.

Having enabled acute depletion of both RAD21 and CTCF, we next inserted the synthetic promoter construct (**Fig. 1C**) to $\sim$179,050,000 (mm39) on chr2, along with 3x CTCF binding sites (L2, L3 from [22] and L1 from [8]) pointing upstream, as well as the ANCH3 array [16] (obtained from NeoVirTech) to allow fluorescent labeling of the synthetic promoter. The synthetic promoter encoded a tagBFP in the exons (the first 132 bp in the first exon and the remaining 567 bp in the second exon), and 128xMS2 [23] in the middle of the first and only intron, utilizing the intron sequence from [24] to reduce the likelihood of transgene silencing by the HUSH complex. SV40 polyA [25] was added downstream of the coding sequence. The 3x CTCF binding sites were flanked by VloxP sites so that they could be removed later via expression of the VCre recombinase [26]. The synthetic promoter, coding sequence and SV40 polyA were flanked by rox sites, 32-bp DNA sequences recognized by the Dre recombinase for site-specific recombination, so that they can be later removed upon expression of Dre recombinase [27]. The direction of transcription pointed upstream away from the 3x CTCF binding site and ANCH3 array. These elements were encoded on the repair plasmid pASH449 (see **Table S1** for plasmids, associated sgRNA sequences and full details). Homozygously edited clone F4 was chosen using genotyping primers P1444 and P1445 (see **Table S2**).

To ensure activation of transcription from a large genomic distance, we sought to engineer a strong synthetic enhancer. We combined three previously characterized super-enhancer fragments [28] from the enhancers of *Esrrb*, *Klf4*, and *miR-290-295*, to build a 1933 bp synthetic enhancer (**Fig. 1C** and **Fig. S1B**; the DNA sequence is provided at the end of this section). We inserted the synthetic enhancer construct to $\sim$178,714,000 on chr2 ($\sim$339 kb upstream of the synthetic promoter), along with 3x CTCF binding sites upstream (R1, R2, R3 from [22]) pointing downstream, as well as the synBsr1 *parS* array (**Fig. S1C**) to allow fluorescent labeling of the synthetic enhancer. The 3x CTCF binding sites were flanked by SloxP sites so that they can be later removed via expression of the SCre recombinase [26]. The synthetic enhancer was flanked by vox sites so they can be later removed upon expression of Vika recombinase [29]. All these elements were encoded on the repair plasmid pASH450 (see **Table S1** for plasmids, associated sgRNA sequences and full details). Homozygously edited clone A3 was chosen using genotyping primers P1490 and P1491 (see **Table S2**).

Having inserted the E-P pair, the fluorescently-tagged binding proteins were next inserted using PiggyBac transposition. Though we had already expressed OR3-$(GS)_3$-mStayGoldx2-P2A-T2A-BSD as described above, we found the mostly cytoplasmic OR3-$(GS)_3$-mStayGoldx2 proteins do not always robustly label the synthetic promoter in every single cell, and decided to additionally express another OR3 construct with a nuclear localization signal (NLS) using the same fluorophore. Briefly, OR3 was cloned into plasmid pASH349 expressing OR3-NLS-2x-mStayGold-NLS from an L30 promoter, and the NLSs are both SV40 NLSs. OR3 was C-terminally tagged with 2 copies of mStayGold, connected by EV linkers [30]. synBsr1-ParB was used to label the synthetic enhancer and was cloned into pASH400 expressing synBsr1-ParB-2x-mScarlet3-NLS from an L30 promoter. MS2 coat proteins (MCP) were used to label nascent RNA. MCP-2x-Halo was cloned into pASH380 and expressed from an EF1$\alpha$ promoter. Expression of each of these binding proteins was established sequentially in clone A3 by repeating the following step: the plasmid encoding the binding protein was co-transfected with a plasmid encoding Super PiggyBac Transposase (500 ng each per 6-well, 4:1 molar ratio of binding protein plasmid vs. PiggyBac Transposase plasmid) using Lipofectamine 2000. Cells were then grown and passaged for 2 weeks so that most expression was from genomic integration of the cassette expressing the binding protein instead of from transfected plasmids. Then transfected cells were isolated using fluorescence-activated cell sorting (FACS) as detailed in the section below. Cells with above-background expression level (untransfected parental cells were used as negative control) were sorted into 10 bins and plated into appropriate dishes depending on cell count. Each of the 10 populations was imaged under the lattice light-sheet microscope to select for the population with the highest signal-to-noise ratio (SNR). The selected population was then plated at low density, individual clones/colonies were picked and plated on 96-well plates. The 96-well plates were then first imaged under the EVOS M5000 microscope to assess cell-to-cell variation in the expression level (except for Halo), and the most homogeneous 24 clones were then passaged and imaged under the lattice light-sheet microscope again to select the clone with the highest SNR and the least cell-to-cell variation. Clone G1 was used for further genome editing.

We next deleted a 2331 bp DNA segment containing an endogenous CTCF binding site between the 339 kb E-P pair at ∼178,835,000 bp (mm39) on chr2. We built a plasmid that can simultaneously express two guide RNAs, pASH375, to improve deletion efficiency. We also provided a repair template by cloning ∼1 kb DNA immediately upstream and ∼1 kb DNA immediately downstream of the 2331 bp deleted segment into pASH470 to improve the probability of a precise deletion and reduce the probability of random insertions or deletions [31,32]. Homozygously edited clone H3 was chosen using genotyping primers P1591 and P1592 (see **Table S2**).

Having obtained clone H3, we found that the expression level of the binding proteins reduced over time, leading to suboptimal SNR. Further, we found a sub-population of the cells with reduced expression from the synthetic promoter also started to emerge. We hypothesized DNA methylation could be responsible for the silencing we observed, and tried different concentrations of DNMT1 inhibitor GSK-3484862 [33, 34] in the culture media, and found the addition of 2 µM GSK-3484862 is well tolerated by the cells for long-term culture and maintained a stable expression profile for both of the binding proteins and the tagBFP from the synthetic promoter, consistent with prior findings [34]. Through multiple rounds of subcloning we were able to identify the clone G7B8G2-clone8-clone7 (referred to as G7B8G2 in this study for simplicity) that recovered the original expression level for different binding proteins and the tagBFP from the synthetic promoter, quantified by flow cytometry and compared to the original quantification. We kept 2 µM GSK-3484862 in culture media across all cell lines from this point on for all downstream experiments, which helped maintain long-term stable expression verified by periodic flow cytometry.

Having established the cell line G7B8G2 with stable expression, harboring the 339 kb E-P pair with convergent CTCF binding sites, we continued further gene editing to generate control cell lines as well as cell lines with different E-P configurations. To generate the control cell line without the synthetic enhancer, we transfected G7B8G2 with pASH447, which encoded both the Vika recombinase and the puromycin resistance gene (see **Table S1**). The expression of Vika recombinase allows the excision of the synthetic enhancer flanked by vox sites [29]. We followed the same protocol as the Cas9-based gene editing protocol described above, except in this case the entire 1.06 µg of DNA used for transfection was pASH447. The expression of the puromycin resistance gene allowed the same selection protocol used in the Cas9-based gene editing protocol described above for enrichment of the transfected cells. We picked colonies grown from single cells into 96-well plates, and homozygously edited clone Vika-H11 was chosen using genotyping primers P1858 and P1859 (see **Table S2**).

To generate the control cell line without the synthetic E-P pair, we transfected clone Vika-H11 with pASH448, which encoded both the Dre recombinase and the puromycin resistance gene (see **Table S1**) to excise the synthetic promoter flanked by rox sites [27]. We followed the same protocol as the Cas9-based gene editing protocol described above with the entire 1.06 µg of DNA used for transfection being pASH448. We picked colonies grown from single cells into 96-well plates, and homozygously edited clone E11G8 was chosen using genotyping primers P1911 and P1912 (see **Table S2**).

To generate the cell line with engineered 3x CTCF binding sites only on the promoter side (339CP; **Fig. S3**), we transfected clone G7B8G2 with pASH445, which encoded both the SCre recombinase and the puromycin resistance gene (see **Table S1**) to excise the engineered 3x CTCF binding sites pointing downstream that were directly upstream of the synthetic enhancer and flanked by SloxP sites [26]. We followed the same protocol as the Cas9-based gene editing protocol described above with the entire 1.06 µg of DNA used for transfection being pASH445. We picked colonies grown from single cells into 96-well plates, and homozygously edited clone S-G5 was chosen using genotyping primers P1860 and P1861 (see **Table S2**). Clone S-G5 was further edited to delete a 1414 bp fragment upstream of the enhancer containing a weak endogenous CTCF binding site, by transfecting pASH375 modified to encode two guide RNAs, GTTACGATGTGCGCTCCAGT and TGGTTGTACTTGTTGCGTCC. Homozygously edited clone S-G5H7 was chosen using genotyping primers P1964 and P1965 (see **Table S2**).

To generate the cell line with engineered 3x CTCF binding sites only on the enhancer side (339CE; **Fig. S3**), we transfected clone G7B8G2 with pASH446, which encoded both the VCre recombinase and the puromycin resistance gene (see **Table S1**) to excise the engineered 3x CTCF binding sites pointing upstream that were directly downstream of the synthetic promoter and flanked by VloxP sites [26]. We followed the same protocol as the Cas9-based gene editing protocol described above with the entire 1.06 µg of DNA used for transfection being pASH446. We picked colonies grown from single cells into 96-well plates, and homozygously edited clone V-C2b was chosen using genotyping primers P1862 and P1863 (see **Table S2**).

To generate the cell line without engineered CTCF binding sites next to the 339 kb E-P pair (339noC; **Fig. S3**), we co-transfected clone G7B8G2 with pASH445 and pASH446, and followed the same protocol as the Cas9-based gene editing protocol described above with the 1.06 µg transfected DNA evenly split between pASH445 and pASH446. We picked colonies grown from single cells into 96-well plates, and homozygously edited clone S+V-A6 was chosen using genotyping primers P1860 and P1861 (for excision of the engineered 3x CTCF binding sites upstream of the synthetic enhancer) as well as genotyping primers P1862 and P1863 (for excision of the engineered 3x CTCF binding sites downstream of the synthetic promoter). Clone S+V-A6 was further edited to delete a 1243 bp fragment upstream of the enhancer containing a weak endogenous CTCF binding site, by transfecting pASH375 modified to encode two guide RNAs, GTTACGATGTGCGCTCCAGT and TGGTTGTACTTGTTGCGTCC. Homozygously edited clone S+V-A6B8 was chosen using genotyping primers P1964 and P1965.

To generate the control cell line with only the ANCH3 and synBsr1 fluorescent labels (without the synthetic E-P pair and without engineered CTCF binding sites: Labels only control; **Fig. S3**), we co-transfected clone S+V-A6B8 with pASH447 and pASH448, and followed the same protocol as the Cas9-based gene editing protocol described above with the 1.06 µg transfected DNA evenly split between pASH447 and pASH448. We picked colonies grown from single cells into 96-well plates, and homozygously edited clone Dre+Vika-S+V-A6B8-E11 was chosen using genotyping primers P1911 and P1912 (for excision of the synthetic promoter) as well as genotyping primers P1858 and P1859 (for excision of the synthetic enhancer).

To generate the cell line with a 253 kb synthetic E-P pair without engineered CTCF binding sites (253noC; **Fig. S3**), we deleted 86 kb between the 339 kb E-P pair in clone S+V-A6. We co-transfected clone S+V-A6 with pASH375 modified to encode two guide RNAs, CACTCCAGGTGTCCCGTCAT and TATCGATCACCTCCAGCCAG, along with the HDR template pASH472 encoding homology arms spanning the deletion junction (see **Table S1**). We followed the same protocol as the Cas9-based gene editing protocol described above, and homozygously edited clone 15A-A6 was chosen using genotyping primers P1897 and P1898 (see **Table S2**). Clone 15A-A6 was further edited to delete a 1243 bp fragment upstream of the enhancer containing a weak endogenous CTCF binding site, by transfecting pASH375 modified to encode two guide RNAs, GTTACGATGTGCGCTCCAGT and TGGTTGTACTTGTTGCGTCC. Homozygously edited clone 15A-A6G9 was chosen using genotyping primers P1964 and P1965.

To generate the cell line with a 170 kb synthetic E-P pair without engineered CTCF binding sites (170noC; **Fig. S3**), we deleted 169 kb between the 339 kb E-P pair in clone S+V-A6. We co-transfected clone S+V-A6 with pASH375 modified to encode two guide RNAs, CACTCCAGGTGTCCCGTCAT and TAGTCCCTTGTAGAATCGAA, along with the HDR template pASH473 encoding homology arms spanning the deletion junction (see **Table S1**). We followed the same protocol as the Cas9-based gene editing protocol described above, and homozygously edited clone 15B-18 was chosen using genotyping primers P1903 and P1904 (see **Table S2**). Clone 15B-18 was further edited to delete a 1243 bp fragment upstream of the enhancer containing a weak endogenous CTCF binding site, by transfecting pASH375 modified to encode two guide RNAs, GTTACGATGTGCGCTCCAGT and TGGTTGTACTTGTTGCGTCC. Homozygously edited clone 15B-18G9 was chosen using genotyping primers P1964 and P1965.

To generate the cell line with a 87 kb synthetic E-P pair without engineered CTCF binding sites (87noC; **Fig. S3**), we deleted 252 kb between the 339 kb E-P pair in clone S+V-A6. We co-transfected clone S+V-A6 with pASH375 modified to encode two guide RNAs, CACTCCAGGTGTCCCGTCAT and CTCCTACAGAAGTCGATGTG, along with the HDR template pASH474 encoding homology arms spanning the deletion junction (see **Table S1**). We followed the same protocol as the Cas9-based gene editing protocol described above, and homozygously edited clone 15C-E11 was chosen using genotyping primers P1907 and P1908 (see **Table S2**). Clone 15C-E11 was further edited to delete a 1243 bp fragment upstream of the enhancer containing a weak endogenous CTCF binding site, by transfecting pASH375 modified to encode two guide RNAs, GTTACGATGTGCGCTCCAGT and TGGTTGTACTTGTTGCGTCC. Homozygously edited clone 15C-A2 was chosen using genotyping primers P1964 and P1965.

To generate the control cell line with the E-P fluorescent labels close to each other (Sub-proximal label control; **Fig. S3**), we deleted $\sim$84 kb DNA between the two fluorescent labels in 15C-A2. We co-transfected clone 15C-A2 with pASH375 modified to encode two guide RNAs, ATTGTGTTCAAGTCCCGATC and ACCGTGAAGGCGCGAGAGTC, along with the HDR template pASH471 encoding homology arms spanning the deletion junction (see **Table S1**). We followed the same protocol as the Cas9-based gene editing protocol described above, and homozygously edited clone 14A-A11 was chosen using genotyping primers P1864 and P1865 (see **Table S2**). This leaves 2362 bp between the ANCH3 and synBsr1 arrays. To further reduce the distance between the 2 arrays and generate the control cell line with the E-P fluorescent labels virtually next to each other (Proximal label control; **Fig. S3**), we deleted the synthetic enhancer remaining in the 2362 bp by transfecting clone 14A-A11 with pASH447 encoding both the Vika recombinase and the puromycin resistance gene (see **Table S1**) to excise the synthetic enhancer flanked by vox sites [29]. Homozygously edited clone 14A-A11E6 was chosen using genotyping primers P1858 and P1859 (see **Table S2**).

To generate the control cell line with the synthetic promoter and synthetic enhancer next to each other (1.5noC; **Fig. S3**), we deleted $\sim$81 kb DNA between the 87 kb synthetic E-P pair in 15C-A2. We co-transfected clone 15C-A2 with pASH375 modified to encode two guide RNAs, ATTGTGTTCAAGTCCCGATC and CAGGTGGCACTCCCGTATGG, along with the HDR template pASH475 encoding homology arms spanning the deletion junction (see **Table S1**). We followed the same protocol as the Cas9-based gene editing protocol described above, and homozygously edited clone 14B-F8 was chosen using genotyping primers P1866 and P1867 (see **Table S2**). Due to the gene body remaining between the synthetic E-P pair, the center-to-center synthetic E-P pair distance was $\sim$5 kb. To further reduce the distance between the synthetic E-P pair, we first deleted the synthetic enhancer by transfecting clone 14B-F8 with pASH447 encoding both the Vika recombinase and the puromycin resistance gene (see

**Table S1**) to excise the synthetic enhancer flanked by vox sites [29]. Homozygously edited clone C6 was chosen using genotyping primers P1858 and P1859 (see **Table S2**). Then we inserted the synthetic enhancer downstream and right next to the synthetic promoter using the repair plasmid pASH476 (see **Table S1**), shortening the center-to-center distance between the synthetic E-P to ∼1.5 kb. Homozygously edited clone 14B5-F10 was chosen using genotyping primers P2001 and P2002 (see **Table S2**).

To generate the cell line with insulating CTCF binding sites between the 339 kb E-P pair with 3x convergent CTCF binding sites (339CECPIC; **Fig. S3**), we inserted 3x CTCF binding sites roughly in the middle of the E-P pair pointing upstream of the synthetic enhancer, and another set of 3x CTCF binding sites next to the first set pointing downstream of the synthetic promoter. Using the same protocol as the Cas9-based gene editing protocol described above, we transfected clone G7B8G2 with the repair plasmid pASH477 (see **Table S1**) and the plasmid encoding Cas9, puromycin resistant gene, and guide RNA. Homozygously edited clone G3 was chosen using genotyping primers P2031 and P2032 (see **Table S2**).

To generate the cell line with insulating CTCF binding sites between the 339 kb E-P pair without engineered CTCF binding sites (339IC; **Fig. S3**), similar to how we generated clone G3, we transfected clone S+V-A6B8 with the repair plasmid pASH477 (see **Table S1**) and the plasmid encoding Cas9, puromycin-resistant gene, and guide RNA. Homozygously edited clone H8 was chosen using genotyping primers P2031 and P2032 (see **Table S2**).

##### 2.3.1 DNA sequence of the 1.9 kb synthetic enhancer

The 1.9 kb synthetic enhancer we engineered is composed of three previously characterized super-enhancer fragments [28] from the enhancers of *Esrrb*, *Klf4*, and *miR-290-295*, respectively (**Fig. 1C** and **Fig. S1B**):

TCTCTCCCCAGGTTTGAAGTCATATTCTAATTTAGAAGTAATTGTCTATTGTATCAGTCAGTAGGGATAACTCTTTACTGCCACACCCTTTGA
AAATGGAGATGTTTACCCCCTCTGTCCCTAGTAGCCCAGACGAGCCTCAAACTTGGCTGTATAGCTGAGCATGACCTTGAACTGTTGCTCCTTCTG
TCCCGGATTCCCAAGGGCTGGCATTACCGGCTGGTATCACCTGATTTACGAGGTTGCTTTCTTTTGCTGGTGGTATTCAACTGCAAAGTTGAGCTA
TCAAGTCATTGGCAAAGAGGACAAAGCCTTTGGATTTGGAGGTGGCCCACGAATTCCACAGATCCGGGAAGCACAGGCCCACGTCCTCCACTCCAA
AGGATCATTAACAGATCCTGACTGTAGTGGTGCTGACGGCAGGGAGTATAGAACCCACAACGTGACCGGAACAAGCCTGTGGTGATTGCCCGGAAA
GCAGCCAGGCCACCTGCAGCATCCCTGCTTCAAGGTCAACTGAAAAGCCTGGGCTCCAGCAGAGTGGCTTTGACCACTGCTGCCTCCTTTAGCCAG
AGGGTGACACTGGAGCGTTGACCCACCTCCTCCAAGGGTCAGGAGTTAGATCAGCCTGAACAGGACCACTGGGTCCTCATAGGCTTTGTAGGTTTG
AATGGGACAGGAGTTTCGAATGAGAACAGACCCATTTTGTACCAGTCCTTGATGTGCAATCCTTGATTAGTTCCA CGTCTCAGAGGAAGGGACTC
CTAAAATGTCTATTGCAAAATTCATAGGTACTAGCCAGGCAGGAAAGAGAAGGTGAGCGGTGGGTATAGACTAGAAGGCCTTGGGAGAGGCAGAGG
AGCTTCGGGTACAAAGCCTGGAATCTGGGGAGAGTGGAGTCAGTGAGAGGAGCAGTGAGTGATTCTAGATAGCTAAAGCAGAGGGTGGGAAGGGCT
AGGATTGCTTTTGACCTTGGAGGCCCCAGGGCACTGTTATTTTCCAAGGCCCCACTCCCACATCCTATCATACACATTGAAATTCACCCACTTTGC
ATATCAAATGAGTTATATATAGCTAACTGGGGAGGCCAGTTGCAAAGACAGTTGACATAATGTTACCTTTTGTAGACATTTAATTACACACTCATC
AATTTCAACTTGGCAACCTCCTCATTTTTGGGAAATGTGGGCGCGGACAGAGCAGCTGTAGAAAACAAATGTGGTAGCTCGGCTTACAATACCTGG
TCTGCAGTCAGTAGTGGAGAGTGGGTCGCTGTAGGAAGTGAGCCCCAGGGCCATCTAAGGGCCTCCACCAAGGGCTTCTGGTGCCTAG CTGTGTA
TGATTCGGGAGGCTGTAAGTGAATCGGTTAAGGCCAGACCCTGGGTCCAGCAGCCCGAAACAGGTGAGACTGGCTACGTGTCTAAAGTAAGGTAAC
CAAACCAGCAGAGTAGCCAAATTACAAAAGGTGTGATCCCCACCTCTCTAACCTATCTGGTCACCTTGGCTCCCAGCAAGGTGACCAGAGCAGTTT
GGAATCTCTTCCTAGGGGACAGGTTCCCTCCTCTGGCTGACACCAGGCTCCAGCCTTTCCCCTTTGCCCAGCATGGGATTCCTGAGGAAATTCCAT
TTCCTTGAGAACCATTGTCTCAATAATCTGTTAGCCCTCGGGAAGGGCGCAGGTGTCTTTAGAATCTCTTCCTAGGGATCCGGATTGTCACAATGC
TCTGCCCTGGATGGGGCAAACCTCCCAGTGTGTTGGGGGGCAGGGCGGGGAGCAGCACAGCCCCAGTGTTCTGGGCAATGGAATTTATCTAAGGAG
CTGGCAATTTCAAGGAGGAATTCAGGTCCCCAAAGTGAGCTGGAATATTGTGACTCCCACCTGGAATCCCAGCATTCTGGAAACAGAGGCAGTAGC
CTCTGCAGTTCAAGGCCA

#### 2.4 Protein depletion

For depletion of RAD21 tagged with mAID, cells were incubated in media containing 100 µM 5ph-IAA (BioAcademia, 30-003-10). For depletion of CTCF tagged with FKBP(F36V), cells were incubated in media containing 500 nM dTAG-13 (Fisher Scientific, 66-055). The same concentration of drug was kept in PBS during wash steps prior to collection of cell pellets for RCMC or western blotting. The same concentration of drug was kept in media during imaging experiments involving depletion of either or both of RAD21 and CTCF.

#### 2.5 Western blotting

Protein extracts were first prepared using nuclei isolated from cultured cells. To isolate nuclei, pelleted cells in Eppendorf tubes were suspended and incubated on ice with buffer A that was 10 times the volume of the cell pellets, and buffer A consisted of 10 mM HEPES pH7.9, 1.5 mM MgCl2, 10 mM KCl, 0.5 mM dithiothreitol, 0.5 mM phenylmethylsulfonyl fluoride, and 1X cOmplete protease inhibitor (Roche, 11873580001). The mixture was centrifuged at 1500 g, 4 ℃ for 5 minutes, and the supernatant was removed carefully with a pipette. Then the pellet was resuspended in 3 volumes of buffer A supplemented with 0.1% IGEPAL CA-630 (Sigma-Aldrich, I8896-100ML). The tube containing the mixture was inverted 10 times and centrifuged at 1500 g, 4 ℃ for 5 minutes. The pellet was then resuspended in 1 volume of buffer B containing 5 mM HEPES pH7.9, 26% glycerol, 1.5 mM MgCl2, 0.2 mM

EDTA, 0.5 mM dithiothreitol, 1X cOmplete protease inhibitor, and 250 mM NaCl. The NaCl concentration was raised to 400 mM by adding the appropriate volume of 5 M NaCl to the side of the Eppendorf tube without contacting the mixture at the bottom of the tube, and the Eppendorf tube was quickly vortexed to achieve a final NaCl concentration of 400 mM. The solution was then incubated on ice for 1 hour with occasional agitation, followed by centrifugation at 2500 g, 4 ℃ for 20 minutes. The supernatant was taken as the nuclear protein extract, and its concentration was determined using the Qubit protein assay kit (Thermo Fisher, Q33212), with 10 µg of protein extract used for western blotting.

1/3 of the volume of protein extract worth of loading buffer, containing 2% SDS, 100 mM Tris pH 6.8, 100 mM dithiothreitol, 10% glycerol and 0.1% bromophenol blue, was added to the protein extract and boiled at 95 ℃ for 10 minutes. The extract was then loaded on a 4–15% Mini-PROTEAN TGX™ precast protein gel (Bio-Rad, 4561085) with the Precision Plus Protein dual color standards (Bio-Rad, 1610374S). The gel was run in 1X Tris/Glycine/SDS running buffer at 200 V for $\sim$1 hour. Then the proteins separated on the gel were transferred to a 0.2 µm nitrocellulose membrane (GenScript, L00732) using the eBlot L1 fast wet transfer system (GenScript, L00686) with the standard transfer setting. Post transfer, the nitrocellulose membrane was blocked in 5% nonfat dry milk (Genesee Scientific, 20-241) in PBS-T (PBS with 0.1% Tween 20 (VWR, M147-1L)) for 30 minutes, and then cut with a razor blade right below the 75 kD band of the standards, separating the membrane into two sections: the top section would be blotted for RAD21 or CTCF, whereas the bottom section would be blotted for HDAC1 as loading control. The cut membranes were then incubated for 16 hours at 4 ℃ with the following primary antibodies suspended in PBS-T with 5% nonfat dry milk: CTCF 1:1000 (Millipore, 07-729), RAD21 1:1000 (Abcam, ab154769), HDAC1 1:1000 (Abcam, ab109411). The membranes were briefly washed in PBS, followed by three washes in PBS-T, each for 2 minutes. Then the membranes were incubated for an hour at room temperature with the secondary antibody, anti-rabbit IgG, HRP-linked (Cytiva, NA934-1ML), suspended 1:5000 in PBS-T with 5% nonfat dry milk. The blots were imaged by chemiluminescence with the Pierce™ ECL Western Blotting Substrate (Thermo Fisher Scientific, catalog number: 32209) and measured with the ChemiDoc MP imaging system. Protein quantification was performed using Image Studio Lite.

#### 3 Fluorescence-activated cell sorting and flow cytometry

##### 3.1 Fluorescence-activated cell sorting (FACS)

To isolate clones with optimized expression levels of binding proteins, fluorescence-activated cell sorting (FACS) was performed using a Sony MA900 multi-application cell sorter (Sony Biotechnology) equipped with a collinear four-laser system (405, 488, 561, and 638 nm). Sony Cell Sorter Software v3.0 was used for instrument control and data acquisition. To exclude non-cellular debris and background noise, the event trigger threshold was set on forward scatter (FSC) at 5%. The sensor gain for FSC was set to 3, and the sensor gain for back scatter (BSC) was set to 23.5%. A sequential hierarchical gating strategy was used to isolate single target cells. Primary cellular events (”cells”) were first identified and gated using an FSC-Area (FSC-A) versus BSC-Area (BSC-A) plot to exclude subcellular debris. To eliminate multi-cell aggregates and coincident events, a two-step doublet discrimination strategy was applied to the primary cell population. First, a primary singlet population (”singlets-1”) was isolated using an FSC-Width (FSC-W) versus FSC-Height (FSC-H) plot. This population was then further refined using a BSC-Width (BSC-W) versus BSC-Height (BSC-H) plot to establish the final purified single-cell gate (”singlets-2”). Only events falling within the ”singlets-2” gate were evaluated for fluorescence and subsequent sorting.

Fluorophore detection and integrated photomultiplier tube (PMT) sensor gains were configured by mapping each fluorophore to its corresponding excitation laser and emission filter channel: mStayGold was detected on the FL1 channel (FITC; 488 nm excitation, 525/50 nm emission) with a gain set to 35.0%; mScarlet3 was detected on the FL2 channel (PE; 561 nm excitation, 585/30 nm emission) with a gain set to 40.0%; and HaloTag® (labeled by incubating cells with 50 nM JFX650 dye for at least 24 hours prior to sorting) was detected on the FL10 channel (APC; 638 nm excitation, 665/30 nm emission) with a gain set to 52.0%. Cells exhibiting above-background fluorescence relative to untransfected parental control cells were sorted into 10 distinct expression bins and subsequently plated for downstream lattice light-sheet screening.

##### 3.2 Flow cytometry quantification of tagBFP expression

For quantification of tagBFP expression levels in the established clones, analytical flow cytometry was performed on the same Sony MA900 instrument. The exact optical configurations and hierarchical gating strategy described for cell sorting above were maintained. Only events falling within the final ”singlets-2” gate were utilized for tagBFP quantification. Target tagBFP excitation was achieved utilizing the 405 nm violet laser, and emission was captured using the FL6 detector (Brilliant Violet 421 channel, 450/50 nm emission). The FL6 PMT sensor gain was set to 40.0%, while the FL1, FL2, and FL10 detector gains remained unchanged as above (35.0%, 40.0%, and 52.0%, respectively). Sample acquisition was performed at a flow rate of approximately 1,000–3,000 events per second, with recording configured to capture a minimum of 100,000 ”singlets-2” events per condition. The distributions of measured single-cell tagBFP expression level for each condition are shown in **Fig. S4**.

#### 4 Microscopy experiments and analysis

##### 4.1 Live-Cell Imaging of the dynamics of the synthetic E-P pair and transcription

For live-cell imaging, mESCs were grown for one (seeding ∼100,000 cells) or two days (seeding ∼50,000 cells) on one well of the Nunc™ Lab-Tek™ 8-well chambered coverglass (no 1.5H, Thermo Fisher Scientific, 155409) coated with Geltrex™ (Thermo Fisher Scientific, A1413301), according to manufacturer's instructions. The same medium for cell culture supplemented with 50 nM JFX650-HaloTag® dye [35,36] was used. For conditions requiring RAD21 depletion, 5ph-IAA (BioAcademia 30-003) was added to a final concentration of 100 µM 2 hours prior to the start of the first movie. For conditions requiring CTCF depletion, dTAG-13 (Fisher Scientific, 66-055) was added to a final concentration of 500 nM 2 hours prior to the start of the first movie. Acquisition was performed on a ZEISS Lattice light-sheet 7 (LLS7) microscope equipped with two objectives, the 3.3x/0.4NA excitation objective (placed at 30° relative to the coverglass) and the 44.83x/1.0 NA detection objective (60° relative to the coverglass). ZEN software v3.12 was used for acquisition control and image deskewing and deconvolution. Excitation was achieved using three laser lines: 488 nm, 561 nm, and 640 nm. Data were recorded using two cameras (Hamamatsu ORCA-Fusion sCMOS). To achieve three-color imaging, the 488 nm (exciting the mStayGold fluorophores tagging the synthetic promoter) and 640 nm (exciting the JFX650 dye labeling nascent RNA) lasers were fired simultaneously. Their emissions were split by a secondary beam splitter dichroic mirror (SBS LP 640). The nascent RNA signal was directed to camera 1 (equipped with a BP 570-620 IR+ emission filter), while the synthetic promoter signal was directed to camera 2 (equipped with a BP 495-550/BP 570-620 emission filter). The 561 nm laser (exciting the mScarlet3 fluorophores tagging the synthetic enhancer) was fired sequentially and its emission was likewise captured on camera 2, ensuring that both DNA labels were recorded on the same camera for 3D E-P distance calculation. The stage was equipped with a humidified incubation chamber at 37°C supplied with 5.5% $CO_2$. Samples were loaded on the meniscus lens (served as the relay of the optics to the coverglass) with water immersion. For acquisition without auto-focus (0.5-second and 5-second frame rate movies), samples were allowed to equilibrate for at least an hour to prevent shifting in z during acquisition. For all acquisitions, we used the “Sinc3 65 × 1500” light sheet and a step size (stage displacement step between consecutive slices) of 0.3 µm.

For 30-second frame rate movies, we recorded 208 slices per frame with an exposure time of 30 ms, with autofocus (using an extended range of 100 µm), for a total duration of 6 hours, with the power of each laser set to: 488 nm (2%), 561 nm (3%), and 640 nm (10%). While minor variations in auto-focusing time introduced slight fluctuations in the interval between consecutive frames (standard deviation of 1.9 s), the average frame rate across the acquisitions remained strictly at the 30-second target. For 5-second frame rate movies, we recorded 70 slices per frame with an exposure time of 30 ms, without auto-focus, for a total duration of 1 hour, with the power of each laser set to: 488 nm (2%), 561 nm (3%), and 640 nm (10%). For 0.5-second frame rate movies, we recorded 13 slices per frame with an exposure time of 10 ms, for a total duration of 6 minutes, with the power of each laser set to: 488 nm (6%), 561 nm (9%), and 640 nm (20%).

To correct for chromatic aberrations, we recorded the chromatic offset frequently with 0.2 µm TetraSpeck microspheres (Thermo Fisher Scientific, T7280) plated on mESCs. 50,000 mESCs were plated on one well of the Nunc™ Lab-Tek™ 8-well chambered coverglass coated with Geltrex™. One day later, the original vial of 0.2 µm TetraSpeck microspheres was sonicated with a VEVOR Ultrasonic Cleaner for 5 minutes, and 10 µL microspheres suspension was diluted in 990 µL media containing 50 nM JFX650-HaloTag® dye. The 1:100 diluted microsphere suspension was sonicated for another 10 minutes, and the media in chambered coverglass well plated with mESCs was replaced with 200 µL of the 1:100 diluted microsphere suspension in media. One day after, the media was replaced with 4% formaldehyde in PBS (diluted from Pierce™ 16% Formaldehyde, Thermo Fisher Scientific, 28908), and incubated at room temperature for 15 minutes, followed by 3 PBS washes, 5 minutes each. After the last PBS wash, 200 µL of PBS was added to the chambered coverglass well containing the mESCs with microspheres on top. To record the chromatic offset, we recorded 210 slices per frame with an exposure time of 10 ms, with auto-focus (saved focus from imaging the synthetic E-P pair), for a total of 5 frames, with the power set to 100% for all three lasers.

Deskewed and deconvolved versions of live-cell movies were first generated using the ZEN software v3.6. For deskewing, the interpolation was set to ”Linear”, and processing method was set to ”Cover Glass Transformation”. The raw effective pixel size was 145 nm × 145 nm in $x$ and $y$, with a 300 nm step size in $z$. Following deskewing and cover glass transformation, the resultant pixel size was 145 nm × 145 nm × 145 nm. The same deskew settings were also selected for deconvolution. For deconvolution, the algorithm was set to ”Constrained Iterative”, normalization was set to ”Clip” with a factor of 1.00, and strength was set manually to 5.0. No correction options were selected, and all advanced settings were set to default options.

Across conditions with transcriptional bursting, we observed a progressive decay in the MS2 signal even after background subtraction and photobleaching correction. We hypothesize this signal attenuation is due to the continuous sequestration of the finite nuclear pool of MCP-2x-Halo proteins by accumulating transcribed RNAs, as supported by prior studies [37–39]. To account for this effect, we developed an empirical, data-driven correction pipeline to reverse the observed MS2 signal decay (**Supplementary Note 3**).

#### 4.2 Fyrtarn framework to yield distance-intensity trajectories

We developed Fyrtarn as an efficient computational framework for detecting, tracking, and measuring cellular dynamics within terabyte-scale lattice light-sheet microscopy (see **Fig. S6A** for an overview). This editable Python library leverages just-in-time (JIT) compilation and CPU/GPU parallelization through Numba [40] and CuPy [41] to efficiently process light-sheet volumes. Further, we make use of nuclear segmentation to drastically reduce the search space for detecting features of interest, allowing for computationally intensive methods to accurately measure cellular dynamics. This open source library is available on GitHub: https://github.com/ahansenlab/Fyrtarn.

##### 4.2.1 Deskewing and cover-slip transformation

To minimize the reading and writing of acquired and processed volumes, we later added efficient GPU-based deskewing to Fyrtarn, including a cover-slip transform to yield isotropic volumes. Below, we derive the deskewing operation which can be described by a single affine transformation. Our implementation aims to match the coordinate system of the ZEN software v3.6 and utilizes the CuPy library [41] to make the transformation computationally tractable (see **Fig. S6A**).

$$A = \underbrace{\begin{bmatrix} 1 & 0 & 0 & -\frac{a}{2} \\ 0 & 1 & 0 & -\frac{b}{2} \\ 0 & 0 & 1 & -\frac{c}{2} \\ 0 & 0 & 0 & 1 \end{bmatrix}}_{\text{Shift}} \quad B = \underbrace{\begin{bmatrix} 1 & 0 & 0 & 0 \\ \frac{\cos\theta}{r} & 1 & 0 & 0 \\ 0 & 0 & 1 & 0 \\ 0 & 0 & 0 & 1 \end{bmatrix}}_{\text{Shear}} \quad C = \underbrace{\begin{bmatrix} \frac{\sin\theta}{r} & 0 & 0 & 0 \\ 0 & 1 & 0 & 0 \\ 0 & 0 & 1 & 0 \\ 0 & 0 & 0 & 1 \end{bmatrix}}_{\text{Scale}}$$

$$D = \underbrace{\begin{bmatrix} \cos\theta & -\sin\theta & 0 & 0 \\ \sin\theta & \cos\theta & 0 & 0 \\ 0 & 0 & 1 & 0 \\ 0 & 0 & 0 & 1 \end{bmatrix}}_{\text{Rotate}} \quad E = \underbrace{\begin{bmatrix} 1 & 0 & 0 & 0 \\ 0 & 0 & 1 & 0 \\ 0 & -1 & 0 & 0 \\ 0 & 0 & 0 & 1 \end{bmatrix}}_{\text{Transpose/Flip}} \quad F = \underbrace{\begin{bmatrix} 1 & 0 & 0 & \frac{b\sin\theta}{2} \\ 0 & 1 & 0 & \frac{c}{2} \\ 0 & 0 & 1 & \frac{a}{2r} + \frac{b\cos\theta}{2} \\ 0 & 0 & 0 & 1 \end{bmatrix}}_{\text{Shift}}$$

where:

$a =$ Number of slices in the stack

$b =$ Height of slice

$c =$ Width of slice

$\theta =$ Angle of the slice (to coverslip)

$r = \frac{\text{pixel size}}{\text{slice step}}$

Let $\boldsymbol{p}_{\text{raw}}$ and $\boldsymbol{p}_{\text{deskew}}$ be positions in the coordinate systems of the raw acquisition and the deskewed volume. We can compute both the transformation matrix and its inverse, which is preferred in affine transformation implementations.

$$\boldsymbol{p}_{\text{deskew}} = \underbrace{F \cdot E \cdot D \cdot C \cdot B \cdot A}_{M} \cdot \boldsymbol{p}_{\text{raw}} \qquad \boldsymbol{p}_{\text{raw}} = \underbrace{A^{-1} \cdot B^{-1} \cdot C^{-1} \cdot D^{-1} \cdot E^{-1} \cdot F^{-1}}_{M^{-1}} \cdot \boldsymbol{p}_{\text{deskew}}$$

$$M = \begin{bmatrix} 0 & -\sin\theta & 0 & y\sin\theta \\ 0 & 0 & 1 & 0 \\ -\frac{1}{r} & -\cos\theta & 0 & y\cos\theta + \frac{a}{r} \\ 0 & 0 & 0 & 1 \end{bmatrix} \qquad M^{-1} = \begin{bmatrix} r\cot\theta & 0 & -r & a \\ -\csc\theta & 0 & 0 & b \\ 0 & 1 & 0 & 0 \\ 0 & 0 & 0 & 1 \end{bmatrix}$$

##### 4.2.2 Richardson-Lucy Deconvolution with Total Variation Regularization

The movies in this study were deconvolved using a proprietary algorithm implemented in Zeiss ZEN v3.6. To independently test the robustness of the Zeiss deconvolution algorithm, we implemented the Richardson-Lucy deconvolution algorithm [42] in the Fyrtarn library, which we found to yield similar results. Using volumes of size $15 \times 15 \times 15$ of bead localizations, we computed empirical PSF kernels for the 488 nm, 561 nm, and 640 nm channels (see

**Fig. S8A** and **Fig. S9A**) to enable deconvolution of acquired volumes after deskewing. To facilitate nuclei segmentation and feature detection within Fyrtarn, we utilize Richardson-Lucy deconvolution [42] with edge-preserving total variation (TV) regularization, as described below. We then can apply Richardson-Lucy deconvolution separately to each deskewed channel for a fixed number of iterations leveraging a CuPy-based GPU implementation [41] (see **Fig. S6A**).

$$f_{k+1} = f_k \cdot \left[ \frac{\left(\frac{f_0}{f_k \otimes h}\right) \otimes \hat{h}}{1 - \lambda \cdot \text{div}\left(\frac{\nabla f_k}{|\nabla f_k|}\right)} \right] \quad (1)$$

where:

$h =$ Point spread function (PSF)

$\lambda =$ TV regularization parameter

$\hat{h} =$ Adjoint PSF

$f_k =$ Deconvolved volume after $k$ iterations

###### 4.2.3 Nuclei segmentation and tracking

To reduce the search space for E-P loci and improve the quality of the measured trajectories, we performed nuclear segmentation on all acquired movies. Since there could be up to four localizations of enhancer or promoter loci per nucleus at any time during the cell cycle of the diploid mESC, nuclear segmentation helped reduce false positive detections. We implemented an efficient GPU-based 3D Laplacian of Gaussian (LoG) filter method, using CuPy [41] functions, that leverages watershed-like morphological gray erosion/dilation to separate nearby nuclei.

Deconvolved volumes, from ZEN processing, of synBsr1-ParB-2x-mScarlet3 signal (**Fig. S7A**), which had the most homogeneous nuclear-localized fluorescence, were first padded with five pixels of zeros in each dimension, background subtracted, and then filtered with a median filter of size $3 \times 3 \times 3$ and a Gaussian filter with scale $\sigma = 4.0$ (**Fig. S7B**). We applied zero-crossing detection using a $3 \times 3 \times 3$ neighborhood following a Laplacian filter of the smoothed volume then filled such shells to yield a mask of detected nuclei (**Fig. S7C**). To separate nuclei, the mask was morphologically eroded using an ellipsoidal structuring element with principal axes of $15 \times 15 \times 5$, to account for nuclei shapes. The resulting masks were then labeled by connected components with a neighborhood of $3 \times 3 \times 3$ (**Fig. S7D**). Labeled nuclei were then recovered by morphological gray dilation using the same ellipsoidal structuring element as the erosion operation and filtered for volumes of $5 \times 10^4$ to $3 \times 10^5$ pixels (or about $152$ to $915$ µm$^3$) (**Fig. S7E**). Bounding boxes of the deskewed and deconvolved channels, along with nucleus masks, were saved for each frame to enable efficient, parallelized downstream processing.

Centroids for each segmented nucleus were computed from the mean position of all pixels contained within each mask. Nuclei centroids were then linked between frames (**Fig. S6A**) without memory using TrackPy [43] with a search radius of 25 pixels (∼3.6 µm) and filtered for a minimum track length of 60 frames.

###### 4.2.4 Dot detection and super-resolved subpixel localization

To assist in localizing enhancer-promoter (E-P) dots, we first computed PSFs for the 488 nm and 561 nm channels **Fig. S8A**. We found scale-space maxima detection [44] to be the most robust method for finding initial integer positions of diffraction-limited dots in the deconvolved volumes. This method constructs a 4D representation of the 3D signal by applying normalized Laplacian of Gaussian (LoG) filters across a range of scales, or $\sigma$, then marks candidate detections based on local maxima in normalized LoG response using a $3 \times 3 \times 3 \times 3$ kernel (see **Fig. S8B**). To select for dots with approximate diameters of 1.5 to 3.0 pixels (∼0.2 to 0.4 µm), we chose ten scales, $\sigma \in \{\frac{0.75}{\sqrt{2}} \cdot 2^{k/9} : 0 \leq k \leq 9\}$. Candidate detections were then first filtered by their responses using a threshold based on selected scalar multiplied by the median of masked nucleus signal and capped at a maximum number five detections. Nearby detections at integer positions were removed within 10 pixels (∼1.5 µm) based on the pairwise weaker normalized LoG response and dots within 3 pixels (∼0.4 µm) were flagged as replicated for subsequent filtering.

We found the nuclei to have structure in all three imaging channels that would otherwise make accurate estimations of distances and intensities difficult. Using ridge regression with a polynomial basis of order two, we fit $15 \times 15 \times 15$ volumes of deskewed intensities with an ellipsoidal mask constructed an ellipsoidal measurement mask based on an anisotropic Gaussian fit of the respective PSF.

$$B_{\text{model}}(x,y,z) = \beta_0 + \beta_1 x + \beta_2 y + \beta_3 z + \beta_4 x^2 + \beta_5 y^2 + \beta_6 z^2 + \beta_7 xy + \beta_8 xz + \beta_9 yz \quad (2)$$

$$\min_{\beta} \sum_{(i,j,k)\notin M} [I_{\text{volume}}(i,j,k) - B_{\text{model}}(i,j,k)]^2 + \lambda ||\beta||^2 \quad (3)$$

To prevent overfitting of the background intensity estimate, we performed ridge regression with an L2 weight penalty of $\lambda = 0.001$. The calibrated model then was used to estimate and subtract the local background of the spot **Fig. S8C**.

To ”super-resolve” the 3D positions of the enhancer and the promoter by determining their subpixel localizations, we ran the FracShift algorithm [45] on the background subtracted, deskewed volumes. The FracShift algorithm overcomes key limitations of other approaches. Fixed intensity-weighted centroid estimates induce pixel bias when performing subpixel localization of diffraction-limited particles, reducing the precision of the positions. While PSF fitting methods are commonly applied in fixed-cell super-resolution microscopy, a limitation of PSF-fitting methods is their lack of robustness to motion blur which makes PSF-fitting less appropriate for live-cell imaging as previously shown [46].

The FracShift algorithm [45, 47] robustly localizes and super-resolves particles while largely eliminating pixel bias. FracShift works by iteratively refining localizations by resampling pixel intensities after applying the intensity-weighted centroid estimated shift with a PSF-like mask. We created a centroid mask by fitting an anisotropic 3D Gaussian function, along the 30° objective angle, of the 488 nm bead-estimated PSF kernel and positions in the ellipsoid defined by twice the principle axes of the covariance matrix (see **Fig. S8A**). We tested a range of scalings of the ellipsoid mask and found about twice the length of principle axes to yield the best localization precision as evaluated by mean-squared displacement (MSD) fitting of E-P trajectories.

For all localizations, we initialized the FracShift algorithm on the initial pixel coordinates from the scale-space detections from **Fig. S8B** and iterative scheme for 30 iterations (see **Fig. S8C**), which was determined to be a sufficient number iterations to converge to tolerances of $10^{-4}$ pixels. We observed a small number of localizations moving toward the boundary of the $15 \times 15 \times 15$ volume and if the position moved outside the space, the detection and initial localization were removed. We then computed residual pixel bias for all tracked E-P pair positions following localization with the FracShift algorithm and observed that the center-bias had largely been corrected with only a slight residual bias remaining (see **Fig. S8D**).

###### 4.2.5 Enhancer-promoter (E-P) channel linking and pairing

Enhancer and promoter detections in the 561 nm (synBsr1-ParB-2x-mScarlet3) and 488 nm (OR3-2x-mStayGold) channels were linked between frames first separately by channel then the linked channel detections were then joined together. Channel detections were first centered based on the nucleus mask centroid then linked using TrackPy [43] with a search radius of 10 pixels (∼1.5 µm), memory of five frames, and a minimum trajectory length of 15 frames (see **Fig. S8E**). Localization of E-P dots within track gaps was attempted using linearly-interpolated positions with the procedure described above and poorly localizing dots were not saved. We chose a Hungarian Algorithm-like approach to pair channel trajectories [48]. Each E-P pair was first scored by counting the number of frames the localizations were within 5 pixels (∼0.7 µm). After scoring, E-P trajectories were iteratively constructed by joining the next highest score with the addition that a single channel trajectory to link multiple trajectories in the other channel (as represented in **Fig. S8E**).

###### 4.2.6 Estimating and correcting chromatic aberrations

To measure accurate 3D distances, we needed to correct unavoidable chromatic aberrations between the 561 nm (synBsr1-ParB-2x-mScarlet3) and 488 nm (OR3-2x-mStayGold) channels [49]. We measured chromatic aberrations between paired bead localizations in these channels then fit a function to estimate and evaluate these corrections on E-P trajectories. We found that the local maxima, with thresholding, of the Laplacian of Gaussian filter response at a single scale efficiently and effectively detected initial bead localizations. FracShift was first run on each bead detection to yield a subpixel localization then, as a filtering step for computing point spread functions (PSFs), we removed bead localizations that were within (∼1.5 µm) of another localization (see **Fig. S8F**). Bead localizations between channels were paired with a search radius of three pixels (∼0.4 µm) and the offset vector between the channel positions was recorded.

To account for registration drift over time and maximize the accuracy of the bead-based registration error correction, we grouped beads collected from adjacent days (using a 15-day moving average window) to compute a unique correction function for each imaging day. We computed the median absolute deviation (MAD) of the position offset vectors along each axis and removed channel bead pairs that were greater than ±2 MADs. After filtering measured offsets, we calibrated a second-order polynomial to estimate the chromatic offset in 3D space, allowing for an approximate mapping between the 488 nm and 561 nm channels.

$$A_{\text{model}}(x, y, z) = \beta_0 + \beta_1 x + \beta_2 y + \beta_3 z + \beta_4 x^2 + \beta_5 y^2 + \beta_6 z^2 + \beta_7 xy + \beta_8 xz + \beta_9 yz \tag{4}$$

$$\min_{\boldsymbol{\beta}} \sum_{i=1}^{N} ||\boldsymbol{I}_i - \boldsymbol{A}_{\text{model}}(x_i, y_i, z_i)||_2^2 + \lambda ||\boldsymbol{\beta}||_2^2 \tag{5}$$

To reduce overfitting to the measured bead offsets, we performed ridge regression with an L2 weight penalty of $\lambda = 0.001$. We evaluated our chromatic aberration corrections based on the position offsets between all enhancer-promoter (E-P) pair localizations (see **Fig. S8G**).

###### 4.2.7 Intensity measurement localization

In order to quantify MS2 intensities, we needed to compute a measurement position in the 640 nm (MCP-2x-Halo) channel. We performed this procedure in two steps. The first step is to estimate the position of the promoter in the 640 nm channel and the second step is to improve the localization of the measurement if an MS2 burst is detected nearby. We applied 3D image registration, using phase cross-correlation [50], between the 561 nm and 640 nm channel bounding boxes to estimate an effective chromatic and camera shift (example in **Fig. S9C**). To efficiently compute the phase cross-correlations, we reimplemented `skimage.registration.phase_cross_correlation` from [51] to perform the computations on a GPU. Using this effective shift and the bead-estimated chromatic shift between 488 nm and 561 nm channels, we then calculated the position of the promoter in the 640 nm channel.

We observed that the promoter position may not always accurately localize on MS2 dots due to errors in localization, registration, and chromatic aberration corrections. To improve the quantification of MS2 intensities, we chose to use the scale-space extrema detection and subpixel localization for enhancer-promoter (E-P) dots on MS2 dots (shown in **Fig. S9B**). If an MS2 dot was detected within a three pixel (∼0.4 µm) radius of the estimated promoter position in the 640 nm channel, we chose to localize the intensity measurement using the MS2 dot position. Otherwise, intensities were measured using the adjusted promoter localization (see **Fig. S9D**).

###### 4.2.8 Nonlinear background estimation for intensity measurements

To accurately measure MS2 dot intensities, we first averaged $15 \times 15 \times 15$ volumes of bead intensities to yield an empirical 640 nm point spread function (PSF) then constructed an ellipsoidal measurement mask based on an anisotropic Gaussian fit of the PSF (see **Fig. S9A**). Let $M$ be the set of volume positions within the mask. For each MS2 localization, we made a $15 \times 15 \times 15$ volume centered on the position within the bounding box and marked pixels inside and outside the dot mask $M$ such that we could subtract an estimate of the background from the observed MS2 dot intensity. We observed structure within the nuclei that often biased the dot intensity measurements, so we chose to calibrate a second-order polynomial model (shown below) within each volume to improve the estimate of the background intensities.

$$B_{\text{model}}(x, y, z) = \beta_0 + \beta_1 x + \beta_2 y + \beta_3 z + \beta_4 x^2 + \beta_5 y^2 + \beta_6 z^2 + \beta_7 xy + \beta_8 xz + \beta_9 yz \tag{6}$$

To prevent overfitting of the background intensity estimate, we performed ridge regression with an L2 weight penalty of $\lambda = 0.001$ which minimized the variance of sampled intensity measurements among a set of $\lambda$. We then calibrated $B_{\text{model}}$ with the closed-form ridge regression solution and evaluated the fitted model to subtract a background estimate from the summed intensities (see **Fig. S9C**).

$$\min_{\beta} \sum_{(i,j,k)\notin M} [I_{\text{volume}}(i, j, k) - B_{\text{model}}(i, j, k)]^2 + \lambda ||\beta||^2 \tag{7}$$

$$I_{\text{measured}}(x, y, z) = \sum_{(i,j,k)\in M} I_{\text{volume}}(i, j, k) - B_{\text{model}}(i, j, k) \tag{8}$$

###### 4.2.9 MS2 intensity corrections

To yield consistent MS2 intensity trajectories, we applied photobleaching corrections for the JFX650™ dye in addition to normalization for the effective 640 nm laser power. We found that movie acquisitions often had different apparent photobleaching of the MS2 intensity. For this reason, we computed a per-movie ensemble photobleaching curve from individual, normalized intensity curves for each nucleus. Using the respective nuclei bounding boxes, we computed the mean intensity from the nucleus segmentation mask and normalized the curve by the computed intensity in the first frame of the nucleus track. We filtered nuclei photobleaching curves to ensure that the MCP-2x-Halo intensity was approximately monotonically decreasing. We computed the medians of the first and last five frames of each nucleus photobleaching curve as estimates of the intensity range. Any curve in which the end intensity estimate was greater than the start estimate or if the curve maximum or minimum was not within 10% of the start or end intensity estimates, respectively, was removed. An ensemble photobleaching curve was computed from the normalized nuclei intensity curves, ensuring at least five samples at each frame (see **Fig. S9G**).

$$I_{\text{model}}(t) = A_0 + A_1 e^{k_1 t} + A_2 e^{k_2 t} \tag{9}$$

The double exponential model above (9) was calibrated with a log-parameterization to the ensemble curve by minimizing the sum of squared errors (SSE) using `scipy.optimize.minimize` with the BFGS method, initial parameters of $\{A_0 = 0.01, A_1 = 0.5, k_1 = 0.001, A_2 = 0.5, k_2 = 0.0001\}$, and a maximum of $10^4$ iterations (movie correction examples shown in **Fig. S9G**). Over the period of imaging acquisitions, we noticed intensity fluctuations in the 640 nm channel. To correct for these fluctuations, a time-windowed PSF kernel in the 640 nm channel was computed by averaging $15 \times 15 \times 15$ pixel volumes centered on bead localizations using the same window of fifteen days as the chromatic aberration corrections. An absolute intensity factor $I_{640\text{ nm}}$ was computed by measuring the maximum intensity of the time-windowed 640 nm PSF kernel (see **Fig. S9F**), such that the resulting track intensity correction is as follows.

$$I'_{\text{track}}(t) = \frac{I_{\text{track}}(t)}{I_{\text{model}}(t) \cdot I_{640\text{ nm}}} \tag{10}$$

###### 4.2.10 Filtering trajectories for replicated and false positive dots

To ensure the quality of the trajectories for our analysis, we created a user interface (UI) as part of Fyrtarn to visualize each trajectory alongside 3D E-P distance and MS2 intensity plots (see **Fig. S10A** for an example). We applied the nuclear segmentation masks to the corresponding bounding boxes with deconvolution for each trajectory then generated time-lapse maximum intensity projections (MIPs) along each axis with a trace of the localizations used for computing enhancer-promoter (E-P) 3D distances and measuring MS2 intensities. The UI developed in this work allows for trajectories to be subsetted or entirely removed due to linking errors and signs of chromosome replication (see **Fig. S10B**).

We first estimated the onset of replicated loci by using E-P dot detections (from **Fig. S8B**). With a search radius of three pixels (∼0.4 µm) used to mark dots with neighboring detections, we found the number of neighboring detections shared between the pair E-P localizations over a sliding window to reasonably estimate replication as confirmed by visual inspection. In the distance and MS2 intensity plots, we marked the first occurrence (shown in yellow on **Fig. S10A**) of a trajectory having at least three neighboring detections within a five-frame window, which could either be accepted or refined in the interface. Two people performed trajectory filtering on all processed movies using the aforementioned interface.

We noticed that E-P pairs residing close to the edge of the nucleus often exhibited attenuated spatial dynamics. Specifically, we attribute this observation to insufficient background correction when FracShift is applied to localize E-P dots near the nuclear periphery, leading to a slight localization bias away from the edge. With the cells being measurably flatter along the Z-axis relative to the X/Y-axes, this bias mainly affected the dynamics in the Z-axis. To ensure the quality of the measured dynamics, we computed 1D mean-squared displacements (MSDs) for each axis (shown in **Fig. S10A**) and rejected trajectories in which the Z-axis MSD plateaued at a substantially lower time lag than the X-axis and Y-axis MSDs. We note that this procedure reduced the plateau of the ensemble Z-axis MSDs but some residual attenuation of the Z-axis dynamics was still observed. To a similar effect, we ensured the MS2 intensity measurements approached zero at the baseline level and that burst signal did not substantially invade the background measurements, which was evaluated visually and with the background quantification (**Fig. S10A**).

Post manual-QC, we also implemented a statistical ensemble filtering algorithm to remove single-frame jumps and extreme localization anomalies. Specifically, the squared Mahalanobis distances of the inter-frame displacements were modeled using a chi-squared distribution with three degrees of freedom. To control for false positives across all tracked events, we applied a Bonferroni correction to our baseline significance level ($\alpha$ = 1e-7) to determine a dynamic cutoff threshold. When an inter-frame displacement exceeded this adjusted threshold, it was flagged as an anomalous jump, and the spatial coordinates of the locus at the destination frame (the latter of the two frames) were masked as NaN in the dataset.

#### 5 Bayesian MSD fitting

The code for running Bayesian MSD fitting on live-cell trajectories produced in this study is available at https://github.com/ahansenlab/synEP/tree/main/MSDfits. Mean squared displacement (MSD) analysis of live-cell imaging trajectories was performed using the Bayesian inference framework implemented in the `bayesmsd` package [52], with custom modifications to improve numerical stability and integrate imaging data acquired with different frame rates, as described below.

##### 5.1 Two-step fitting

Given that all experimental conditions involved the same genomic locus and therefore share a similar local nuclear environment, the anomalous exponent $\alpha$ was expected to be consistent across conditions. To leverage this physical premise and robustly extract the underlying polymer dynamics, trajectories were modeled using a two-step fitting approach. Initially, Bayesian MSD fitting [52] was performed with $\alpha$ allowed to vary freely between 0.3 and 0.5 [53], which revealed a median $\alpha$ of 0.37 across conditions. Subsequent Bayesian MSD fitting was then carried out with $\alpha$ fixed to 0.37 to infer the localization error alongside the remaining spatial and temporal parameters (crossover time $\tau$ and steady state variance $J$). Although we report an apparent median mESC $\alpha$ of 0.37, we emphasize that chromatin dynamics does not appear to follow a power law behavior across time in mESCs [53] and this $\alpha$-exponent should therefore be interpreted as the local slope rather than being reflective of a true power law.

##### 5.2 Integration of different frame rates

To account for varying temporal resolutions, data acquired at 30-second and 5-second frame rates were integrated where possible. For each experimental condition, if the 5-second frame rate dataset contained more than 40 valid trajectories, the 30-second and 5-second datasets were modeled simultaneously using a joint fitting approach (`bayesmsd.FitGroup`). For 170noC, since 0.5-second frame rate data were also available, the 30-second, 5-second, and 0.5-second datasets were jointly fit. In these joint fits, the underlying physical parameters—including the crossover time scale $\tau$, the steady state variance $J$, the derived scaling prefactor $\Gamma$, and the anomalous exponent $\alpha$—were forced to be shared globally across both temporal resolutions as well as across all three spatial dimensions (assuming isotropic motion). In contrast, the localization errors for each of the three spatial axes were fit independently for the 5- and 30-second (and 0.5-second for 170noC) frame rates. Only if there were $\geq$40 trajectories for a given condition and a given frame rate, the dataset was included in the Bayesian MSD fit.

Parameter optimization was performed using the L-BFGS-B algorithm [54]. Numerical safeguards were integrated directly into the MSD evaluation function to prevent optimizer divergence. This included bounded clipping of exponential arguments and complementary error function (`erfc`) inputs, along with a numerical penalty to restrict the optimizer from exploring mathematically undefined parameter spaces.

#### 6 Bayesian inference of looping dynamics (BILD) to estimate looping probability and lifetime

We applied Bayesian inference of looping dynamics (BILD) [8] on the 339 kb E–P pair with convergent CBSs (339CECP) to quantify the looping probability and lifetime of this CTCF–CTCF loop. Briefly, BILD defines two states ("looped" and "unlooped") from control experiments and then segments the inputted 3D distance trajectories into those states using a hierarchical Bayesian model.

##### 6.1 Calibration of Rouse model

To calibrate the Rouse model underlying BILD, we performed MSD fitting on the experimental data for different conditions; these were performed separately from previous MSD fits because the Rouse model requires $\alpha = 0.5$. Similarly to previous MSD fits, we integrated 5-second frame rate and 30-second frame rate data using `bayesmsd.FitGroup`; in these joint fits, a single $J$ and $\Gamma$ were fitted for each condition, while the localization errors $(\sigma_x, \sigma_y, \sigma_z)$ were allowed to vary between the 5-second and 30-second datasets. The 339noC condition had MSD parameters of $\Gamma = 0.000815\,\mu\mathrm{m}^2\,\mathrm{s}^{-0.5}$ and $J = 0.0229\,\mu\mathrm{m}^2$, which were taken to represent the unlooped state. To parametrize the looped state, we took the parameter $J = 0.0894\,\mu\mathrm{m}^2$ from the 339CECP ΔRAD21 fit and rescaled it to estimate the $J$ parameter for the 1.8-kb average tether between the centers of the fluorophore binding sites when the loop is formed. The localization-error corrected experimental MSDs and the fitted model MSDs are shown in **Fig. S13A**. The fitted MSD parameters were then used to calculate the Rouse model parameters (**Fig. S13B**) following the derivation in previous work [8].

##### 6.2 Inferring looping events using BILD

Looping events were inferred from the live imaging trajectories of the 339CECP cell line as well as the two control cell lines (339noC and 339CECP ΔRAD21). To prevent biases from varying frame rate, we analyzed all trajectories taken from the 30-second movies, while trajectories from 5-second movies were downsampled by every six frames to obtain 30-second movies. Trajectories containing at least 100 frames were used for BILD analysis. To prevent biases from varying trajectory lengths, we ran BILD on 100-frame segments of trajectories starting every 50 frames (e.g., 0 to 100, 50 to 150, 100 to 200, etc.); this resulted in two overlapping BILD inferences at some timepoints. If any inferences at a particular timepoint did not agree, we chose the inference that was closer to the center (or equivalently, further from the beginning and end) of the segment from which it originated. Example trajectories and BILD inferences for all three conditions are shown in **Fig. S13C**. All BILD runs were performed with an evidence bias of $\Delta E = 2$, following previous studies [8, 12].

We next used the BILD inference results to estimate the loop lifetime of the 339CECP loop. In estimating the loop lifetime, we encounter the problem of censoring, which refers to the systematic underestimation of loop lifetimes that arise from looping events that start before the beginning of a trajectory or finish after the end, and are thus not observed in full. The Kaplan-Meier estimator for the survival function is designed to handle this specific problem [55], so we apply it here, following the procedures described in our previous work that were implemented in the accompanying `tracklib` library [8]. The Kaplan-Meier survival curve had a median of 11.0 min, with a 95% confidence interval ranging from 9.5-12.0 min (**Fig. S13D**). A second method to estimate the median loop lifetime is to assume a model of exponentially distributed lifetimes and obtain the maximum likelihood estimate (MLE) of the mean lifetime $\hat{\tau}$ given the data, treating censored and uncensored observations differently (full derivation in [8]).

These exponential models exhibited survival curves that aligned well with the Kaplan-Meier survival curves; the mean lifetime of the 339CECP condition was estimated to be $\hat{\tau} = 16.0$ min, with a 95% confidence interval ranging from 14.5-17.6 min. This would correspond to a median lifetime of $\hat{\tau} \ln 2 = 11.1$ min, with a 95% confidence interval ranging from 10.0-12.2 min.

Lastly, to estimate the looping probability of the 339CECP loop, we calculated the fraction of timepoints at which BILD predicted a loop, and performed false positive correction using the previously established formula corresponding to $\Delta E = 2$ [8]:

$$f_{\text{true}} = (f_{\text{inferred}} - 2.3\%)/1.34.$$

Then, we used the previously described bootstrapping method [8] to estimate the uncertainty, resulting in a mean looping probability of 8.9% and a standard deviation of 0.8 percentage points (**Fig. S13E**).

The code to run the analyses described above is available at https://github.com/ahansenlab/synEP_bild.

### 7 Region Capture Micro-C and analysis

#### 7.1 Experimental procedure and drug treatments

To quantify the interaction probability between the synthetic E-P pair, we performed Region Capture Micro-C (RCMC) [56, 57] in the cell lines with different E-P configurations. Two biological replicates were prepared for each condition (four biological replicates for 339noEP and 339noC(ΔCTCF)). RCMC was performed as previously described [57]. For conditions with RAD21 or CTCF depletion, the cells were treated with 100 µM 5ph-IAA or 500 nM dTAG-13 respectively for three hours before harvest. During harvest, the same concentration of drugs was maintained in PBS to prevent reversal of protein degradation.

#### 7.2 RCMC data analysis pipeline

The code for analyzing the RCMC data produced in this study is available at https://github.com/ahansenlab/synEP/tree/main/RCMCanalysis. The pipeline for RCMC data analysis was modified from a previously written snakemake workflow [58], as described in more detail below.

##### 7.2.1 MS2 array read rescue

The 128x MS2 array is composed of repeats of B-A-A-B-A-B-B-A pattern, where A and B are 353 bp and 351 bp respectively. Reads mapped to the MS2 arrays were multiply mapped due to the presence of A and B repeats, and were rescued and uniformly redistributed among the four copies of the repeats after the initial alignment step.

##### 7.2.2 Two-pass empirical read redistribution

To accurately resolve multi-mapped reads originating from a 1632 bp fragment 400 bp downstream of the synthetic enhancer heterozygously duplicated to 1618 bp upstream of the synBsr1 array (external to the synthetic E-P pair) in all cell lines in this study, we implemented a two-pass empirical redistribution algorithm. In the first pass, baseline alignments were generated and deduplicated using pairtools [59]. Rather than relying on genome-wide distance-decay approximations, we utilized these first-pass baseline alignments to calculate local empirical interaction ratios. Specifically, we isolated uniquely mapped contacts anchored within the 5 kb sequences immediately flanking the duplicated loci. By quantifying interactions in 2 kb bins across the chromosome, we generated a virtual 4C profile that inherently accounted for local interaction features such as CTCF insulation.

In the second pass, multi-mapped reads mapping to the 1632 bp fragment were evaluated and reassigned using a probabilistic model. The assignment probability was calculated by convolving the empirically derived local interaction ratios with the underlying genetics of the locus. To reflect the heterozygosity of the upstream duplicated copy, an assignment weight of 0.5 was applied to the upstream copy, while the homozygous downstream copy received a weight of 1.0. To normalize the visual density of the heterozygous copy for downstream analysis, another copy of the reads assigned to the upstream copy was created by shifting the original reads +1 bp prior to sorting to evade the deduplication algorithm.

##### 7.2.3 Data aggregation and filtering

Following read redistribution, biological replicates and conditions processed across multiple sequencing runs were aggregated into pooled datasets. To remove unligated background artifacts prior to normalization, inward-facing read pairs were subtracted from the aggregated contact matrices using the `remove-inward` function of the neighbor-balance toolkit [60].

###### 7.2.4 Two-step matrix balancing

Finally, the filtered contact data was compiled into multi-resolution matrices from a base 400 bp resolution using `cooler zoomify` [61]. To normalize the data while accurately preserving biological variation in contact density, we employed a two-step balancing approach. First, the matrices were normalized using iterative correction and eigenvector decomposition (ICE) [62]. Subsequently, we applied the neighbor-balance algorithm [60] to restore true variations in physical density by re-normalizing the intermediate ICE-balanced contact map against local interaction patterns—specifically, the average contact frequency of a genomic bin with its immediate neighbors.

##### 7.3 Quantification of E-P interactions and correlation with expression data

To correlate BFP expression measured by flow cytometry (or MS2 signal measured in live-cell imaging) with the 3D E-P interaction probabilities quantified by RCMC, mean interaction scores between the synthetic E-P pair were extracted from the normalized 800 bp resolution contact maps. To ensure the measurements reflected absolute spatial proximity rather than distance-normalized enrichment, we utilized the observed contact frequencies without doing observed-over-expected (O/E) transformations. Quantification was performed by fetching the local contact sub-matrix surrounding the exact E-P coordinates and applying a computed geometric boolean mask to isolate a 2.4 kb by 2.4 kb circular footprint. The RCMC interaction strength was then calculated as the arithmetic mean of the pixels falling within this isolated radial boundary.

To robustly capture biological variance, the mean interaction score for each condition was derived from the fully aggregated contact map combining reads from both biological replicates, while the standard error of the mean was independently calculated by parallel quantification of individual biological replicates. Both the mean interaction scores and their associated standard errors were subsequently scaled relative to the 340 kb E-P pair without CTCF binding sites. Finally, to quantitatively evaluate the structure-function relationship, these normalized 3D interaction scores were correlated with their corresponding BFP reporter gene expression levels using linear regression analysis.

#### 8 Loop extrusion and 3D polymer simulations

The code for running polymer simulations performed in this study is available at https://github.com/ahansenlab/synEP/tree/main/PolymerSimulation. The code for analyzing polymer simulation results produced in this study and for polymer simulation parameter estimation is available at https://github.com/ahansenlab/synEP/tree/main/PolymerSimAnalysis.

##### 8.1 Time steps and lattice set-up

We used a fixed-time-step Monte Carlo algorithm for 1D simulations as described in previous work [63], where each lattice site corresponded to 1 kb of DNA. To perform 1D loop extrusion simulations for estimating cohesin parameters, the chromosome was defined in two distinct ways depending on the condition being simulated. To simulate the CTCF-depleted condition (for estimating cohesin processivity and density), we used a lattice of $G = 2034$ sites with no CBSs. To estimate CBS boundary strengths (defined as the probability of a cohesin motor subunit stalling upon encountering a CBS) and cohesin lifetime boosts when stalled, we simulated an 1100 kb region with the following CTCF boundary locations to emulate the CBSs in the 339CECP cell line:

$$\text{Left-pointing CTCF positions} = [810] \tag{11}$$

$$\text{Right-pointing CTCFs positions} = [469] \tag{12}$$

The CBSs in our simulation were directional and could only stall cohesin movement if the cohesin motor subunit's extrusion direction was convergent with the direction of the CBS. Upon an encounter between a CBS and a cohesin motor subunit, the cohesin motor subunit could be stalled by the CBS with a probability $s = 0.875$, or would pass the CBS with a probability $(1 - s)$. Each CBS in our simulation occupies one monomer and corresponds to the 3x engineered CBSs in our cell lines (**Fig. S2C**). The estimation of $s$ is detailed below. Once one of a cohesin's two motor subunits was stalled by a CBS, no further movement of that subunit was allowed until the cohesin dissociated from the locus and re-associated elsewhere on the DNA. However, the other motor subunit of cohesin was allowed to continue extruding independently until it was either blocked by another CBS or the cohesin dissociated from the DNA.

##### 8.2 Cohesin association and dissociation rates

All 1D loop extrusion simulations were performed with a fixed number of cohesins, determined by the ratio of chromosome length $G$ and cohesin separation $d$ (i.e., the inverse of the density). When a cohesin dissociated from the genome, it immediately reloaded at another pair of adjacent lattice positions with uniform probability, provided that the lattice position was not already occupied by existing cohesin motor subunits. The cohesin dissociation rate was governed by the cohesin processivity $\lambda$ (i.e., the average length of DNA extruded by an unobstructed cohesin before it dissociates) and any additional fold-increase in lifetime, $b$, gained by stabilization at a CBS [8, 64, 65].

#### 8.3 Identifying loop extrusion parameters

We aimed to identify a total of 6 parameters: ① cohesin processivity, ② cohesin separation (density), ③ cohesin extrusion rate, ④ boundary strength (probability that the CBSs stall cohesin), ⑤ fold-increase (boost) of cohesin lifetime due to CTCF, and ⑥ enhancer-promoter stickiness.

①-② Cohesin processivity and separation were jointly estimated by fitting the contact probability decay $P_c(s)$ curve to experimental data. We performed 1D loop extrusion simulations across a grid of processivity and separation values, simulating the CTCF-depleted condition. By minimizing the root-mean-square error (RMSE) between the simulated and experimental $\log(P_c(s))$ curves, we identified the best-fit parameters to be a processivity of 254 kb and a cohesin separation of 200 kb (**Fig. S15A**).

③ For the cohesin extrusion rate, we used an extrusion speed of 125 bp/s per motor unit [8], resulting in a speed of 250 bp/s for one cohesin with two motors. Given our estimated processivity, this yields a total cohesin residence time of $\sim$20 minutes, which is consistent with prior experimental estimates [66]. We note that *in vitro* estimates of the cohesin extrusion speed place it closer to 500 - 2,000 bp/s [67, 68]. A likely explanation for this seeming discrepancy is that *in vitro* experiments measure the instantaneous extrusion speed, whereas what we use is the average speed over a full cycle. In cells, there are likely lots of "mini-barriers" to cohesin which temporarily will slow it down or transiently pause it. Consistent with this, the processivity of cohesin appears much larger in inactive heterochromatin than in active euchromatin, presumably due to fewer extrusion barriers [69]. Thus, the speed used here refers to the average speed over a full cycle, not the instantaneous speed measured *in vitro*.

④ The boundary strength, or the probability $s$ that a CBS stalls a cohesin motor upon encounter, was determined based on CTCF occupancy. Each CBS in this study consists of 3 CTCF binding sites pointing in the same direction. Using a genome-wide average individual CBS occupancy of 50% [70, 71], the probability that at least one of the three sites is occupied (and thus acts as a boundary) is calculated as:

$$s = 1 - (1 - 0.5)^3 = 0.875 \tag{13}$$

Hence, we estimated the boundary strength to be $s = 0.875$. As a consequence of the 3xCBS array, this boundary strength estimate is robust and relatively insensitive to variations in individual CBS occupancy (**Fig. S15B**).

⑤ The fold-increase (boost) of cohesin lifetime upon stalling at a CBS was estimated by comparing simulated and experimental insulation profiles. Using the estimated processivity (254 kb), separation (200 kb), and boundary strength ($s = 0.875$), we swept various boost factors. We found that a boost factor of $b = 5$ was most consistent with the experimental insulation profile (**Fig. S15C**).

⑥ Finally, the EP stickiness (selective attraction energy) was estimated to match the experimentally observed focal enrichment between the E-P pair on the Region Capture Micro-C (RCMC) map for the 339noC cell line. By comparing the simulated focal enrichment between the E-P pair with the RCMC-quantified enrichment, we determined the optimal EP attraction energy to be $3k_BT$ (**Fig. S15D, E**).

#### 8.4 3D polymer simulations via OpenMM

Next, we coupled the 1D loop extrusion dynamics to a 3D polymer model and performed molecular dynamics simulations using Polychrom [72], a package that wraps the molecular simulation toolkit OpenMM [73]. In this coupled model, cohesins act as harmonic bonds between two polymer monomers. These bonds are dynamically updated depending on the position of cohesins on the chromosome. The underlying 1D simulations were first run for 10,000 translocation steps to reach a steady state before being coupled with the 3D polymer simulations.

For the 3D coupled simulations, the 1D cohesin extrusion was set up on a chromosome defined as a lattice of $G = 70,000$ sites consisting of seven repeats of a 10,000 kb region. We simulated ten different conditions to mirror the nine experimental cell lines in this study, as well as the *Sox2* locus shown in **Fig. 1B**. Across simulated conditions, a selective attractive energy of $3k_BT$ was applied between the E and P monomers to emulate experimentally observed "stickiness" between the E-P pair (**Fig. S15**).

Within each 10,000 kb repeat, the specific monomer coordinates for the E-P pairs and CTCF-binding sites (CBSs) were defined as follows:

- **339CECP:** The E-P pair was placed at coordinates $4830$ and $5169$ (yielding a 339 kb E-P distance). The left-pointing and right-pointing CBSs (each corresponding to the 3x engineered CBSs in our cell lines) were placed at $5170$ and $4829$ (yielding a 341 kb CBS-CBS distance).

- **339noC:** The E-P pair was placed at coordinates $4830$ and $5169$ (no CBSs).

- **339IC:** The E-P pair was placed at coordinates $4830$ and $5169$. Intermediate CBSs were placed between the E-P pair at coordinates $4999$ (left-pointing) and $5094$ (right-pointing).
- **339CECPIC:** The E-P pair was placed at coordinates $4830$ and $5169$. The convergent CBSs were placed at $5170$ (right-pointing) and $4829$ (left-pointing). Intermediate CBSs were placed between the E-P pair at coordinates $4999$ (left-pointing) and $5094$ (right-pointing).
- **339CE and 339CP:** The E-P pair was placed at coordinates $4830$ and $5169$, with right-pointing CBS at $4829$. Since the 3x engineered CBSs for 339CE and 339CP is symmetrical with respect to the E-P pair, the same chromosome setup was used for 339CE and 339CP.
- **87noC:** Enhancer at $4956$, Promoter at $5043$ (87 kb E-P distance).
- **170noC:** Enhancer at $4915$, Promoter at $5085$ (170 kb E-P distance).
- **253noC:** Enhancer at $4873$, Promoter at $5126$ (253 kb E-P distance).
- **Sox2 locus configuration:** To emulate the endogenous *Sox2* topology (**Fig. 1B**), the enhancer (SCR) and the promoter (*Sox2*) were placed at coordinates $5061$ and $4952$, respectively (yielding a 109 kb E-P distance). Right-pointing CBS was placed at $4950$ with boundary strength $s$ = 0.5, and left-pointing CBSs were placed at $4936$, $4938$, $5036$, with boundary strength $s$ of 0.5, 0.25, and 0.5, respectively.

Chromosomal polymers were constructed of $G$=70,000 consecutive monomers bonded via the pairwise potential:

$$U_{\text{bonds}}(r) = \frac{k}{2}(r - b_o)^2 \,, \tag{14}$$

where $k = 2k_BT/\delta^2$ is the spring constant ($k_B$: Boltzmann constant, $T$: temperature, and $\delta = 0.1$ monomers), $r = |r_i - r_j|$ is the 3D distance between adjacent monomers, and $b_o = 1$ is one monomer size (the mean 3D distance between adjacent monomers). Monomers connected by a cohesin were held together by the same potential. To account for excluded volume interactions between monomers, we added a weak polynomial repulsive potential:

$$U_{\text{excl}}(r) = \frac{\epsilon_{\text{exc}}}{\epsilon_{\text{m}}}\left(\frac{r}{\sigma}r_{\text{m}}\right)^{12}\left((\frac{r}{\sigma}r_{\text{m}})^2 - 1\right) + \epsilon_{\text{exc}} \,, \tag{15}$$

defined for $r < \sigma = 1.05$, where $r_{\text{m}} = \sqrt{6/7}$, $\epsilon_{\text{m}} = 46656/823543$, and $\epsilon_{\text{exc}} = 50k_BT$. Additionally, a selective attraction energy of $3k_BT$ was applied between the monomers corresponding to the E-P pair.

For each simulation run, the polymer was initialized as a compact conformation on a cubic lattice with normally distributed velocities. The error tolerance for the variable time-step Langevin integrator was set to 0.01, and the collision rate was set to 1. Simulations were performed with spherical confinement to achieve a DNA volume fraction of 20% per simulation volume. We calibrated the simulation time and distances to real time and distances using MSDs calculated from the live-cell imaging data. Using a maximum likelihood estimation procedure to infer the anomalous diffusion parameters from our experimental MSDs, and a least-squares fitting procedure to calibrate our 3D polymer simulation MSDs, we inferred that each stored simulation time step corresponded to approximately $\sim$0.01 second intervals, and monomers corresponded to $\sim$21 nm in diameter. Polymer conformations sampled at approximately $\sim$0.01 second intervals corresponded to 236 Langevin dynamics polymer integration steps using the conditions above. Simulations were run for a total of $\sim$2,500,000 saved blocks. From the resulting conformations, we calculated 3D distance distributions for the exact E-P anchors, as well as adjacent fluorescent probes (e.g., $\pm 1 - 2$ kb offsets from the E-P pair). We also extracted time-course data and simulated RCMC maps (estimating E-P stickiness).

#### 8.5 Implementation of time-gated E-P interactions and parameter search

To evaluate the minimum duration for productive E-P interactions that trigger transcription activation, we implemented a rolling-window time gate ($\tau_{\text{GATE}}$) on the simulated E-P distance trajectories. First, continuous 3D polymer conformations were calibrated to physical units, with each simulation time step corresponding to $\sim$0.01 seconds and each monomer corresponding to $\sim$21 nm. For any given interaction radius $R_{\text{E-P}}$, a binary contact mask was generated, assigning a value of 1 when the 3D E-P distance fell below $R_{\text{E-P}}$, and 0 otherwise.

To account for the dynamic flickering of the polymer, we applied a uniform rolling average filter over a temporal window defined by $\tau_{\text{GATE}}$. An interaction was considered ”productive” only if the E-P pair remained within the distance threshold $R_{\text{E-P}}$ for a fraction of the time gate window exceeding a defined interaction tolerance (e.g., $\geq$ 55%). This approach ensured that interactions too brief were filtered out, while sustained interactions were captured.

To identify the optimal parameters that recapitulate the reduction in tagBFP expression in 339noC and 339CECP upon CBS insertion between the E-P pair (**Fig. 5B,C**), we performed a 3D grid search spanning $R_{\text{E-P}}$ (22–150 nm), $\tau_{\text{GATE}}$ (0.0–30.0 seconds), and interaction tolerance (30%–100%). For each parameter combination, we calculated the time-gated interaction probabilities across different E-P configurations. The optimal parameters were identified by minimizing the sum of squared logarithmic errors between the simulated time-gated interaction ratios and the experimentally measured tagBFP expression ratios.

##### 8.6 Simulation of transcription and burst pileup

To create the E-P distance pile-up at burst onsets identified by simple peak calling (**Fig. 1B**), we implemented a two-state telegraph model of transcription [74] coupled strictly to 3D E-P distance. The promoter transitions from the inactive (OFF) to the active (ON) state at a rate $k_{\text{on}}$ of 5 min$^{-1}$ [75, 76] exclusively during periods when the 3D E-P distance falls below an interaction radius equivalent to 1.5 monomer lengths. The promoter transitions back to the OFF state at a constant rate of $k_{\text{off}} = 0.5$ min$^{-1}$ [23, 77, 78].

While in the ON state, RNA polymerases are loaded onto the gene stochastically following a Poisson process with a mean loading rate of $k_{\text{load}} = 10.0$ min$^{-1}$ [23, 79, 80]. The polymerases subsequently elongate along a 3400 bp gene at a constant speed of 33 bp/s. To evaluate the impact of 5' UTR / first-intron versus 3' UTR labeling, we generated simulated MS2 signals by convolving the polymerase elongation trajectories with position-specific kernels reflecting the binding of fluorescently labeled MCP proteins to the MS2 stem loops. Specifically, for the first-intron MS2 labeling, we implemented a transient "shark fin" kernel profile that rises rapidly upon Pol II loading but drops to zero immediately after the MS2 cassette is fully transcribed, mimicking the excision and release of the intron due to co-transcriptional splicing [81–83]. Conversely, the 3' MS2 labeling was modeled with a kernel that remains zero until the polymerase reaches the MS2 cassette in the 3' UTR, introducing a temporal delay between the initial enhancer-promoter contact and the appearance of MS2 signal.

To recapitulate localization uncertainty in live-cell imaging, we added Gaussian localization noise estimated from Bayesian MSD fitting (same as described above, with $\alpha$ as a freely fitted parameter) to the simulated 3D E-P coordinates ($\sigma_x = 21.1$ nm, $\sigma_y = 23.7$ nm, $\sigma_z = 47.3$ nm) [84]. We then identified transcription bursts by calling peaks on the simulated MS2 signals using the `scipy.signal.find_peaks` algorithm [54]. To isolate distinct bursting events, we applied a minimum peak height threshold of 1.0 and required a minimum temporal separation of 5 simulation frames between consecutive peaks. Using these burst annotations, we performed burst pileup analysis by aligning the simulated E-P distance traces relative to the burst onset.

#### 9 Deep learning-based inference of burst onsets and E-P interaction dynamics

Because prior work did not detect any relationship between E-P proximity and transcription initiation [84], we wanted to take a maximally unbiased and assumption-free approach to test if such a relationship existed in our data. We therefore turned to a deep learning approach, specifically a Long Short-Term Memory (LSTM) [85] recurrent neural network approach, as described in more detail below. The code for change-point detection based burst labeling and for predicting transcription burst onsets using 3D E-P distances with LSTM neural network is available at https://github.com/ahansenlab/synEP/tree/main/LSTM.

##### 9.1 Change-point detection pipeline for initial burst labeling

To isolate transcription burst onsets from confounding fluctuations in MS2 signal caused by stochastic Polymerase II (Pol II) loading, we developed a change-point detection (CPD) pipeline to generate a preliminary set of binary burst labels. Prior to CPD, raw MS2 intensity trajectories were pre-processed to account for outliers and cell-to-cell variations in baseline signal. First, to prevent extreme measurement outliers from skewing the temporal segmentation, the MS2 signals were computationally clipped, capping extreme values at the 1% and 99% percentiles. Next, to account for cell-to-cell variability in baseline fluorescence, each trajectory was normalized against its median intensity, effectively converting raw MS2 intensities into comparable relative fold-changes. To prevent mathematical instability in dim trajectories where dividing by a near-zero median would artificially inflate noise, we set the noise floor to 20% of the global standard deviation. Finally, short gaps (up to 3 frames) in the MS2 trajectories were linearly interpolated to minimize temporal discontinuity.

Following this pre-processing, we applied a penalization-based segmentation algorithm to identify abrupt shifts in the MS2 signal that may correspond to burst onsets. Rather than relying on simple intensity thresholds which are highly susceptible to transient noise, this approach partitions the noisy, continuous trajectories into distinct temporal windows corresponding to bursting and non-bursting states. To optimize the sensitivity and specificity of this binary burst calling, we calibrated the segmentation parameters by evaluating the integrated MS2 signals for frames categorized as bursting frames (area under the curve for the fraction of bursting frames) against normalized median tagBFP expression across different conditions. We selected the parameters that maximized the linear correlation between the integrated MS2 signals and tagBFP expression. Using the Pruned Exact Linear Time (PELT) search algorithm implemented via the Python `ruptures` library [86], this yielded an optimal change-point penalty of 60.0, and a burst classification threshold of 0.85, for both the 30-second and 5-second frame rate datasets.

#### 9.2 LSTM ensemble architecture and training strategy

Because these initial CPD-derived labels inevitably include false-positive transcription onsets, such as secondary MS2 peaks triggered by stochastic Pol II loading rather than genuine promoter state transitions, we used them as "baits" to train an ensemble of Long Short-Term Memory (LSTM) recurrent neural networks. By restricting the models' input to 3D E-P distance, the imperfect CPD labels force the LSTM to learn and extract only the burst onsets truly driven by underlying E-P distance dynamics.

Prior to training, we curated the dataset to exclude specific experimental conditions that lacked a dynamic distance-transcription relationship. First, control conditions missing either the synthetic enhancer or the synthetic promoter (e.g., 339noE, 339noEP) were excluded, as no transcription is expected or observed in these configurations (**Fig. S4**). Second, conditions exhibiting sustained E-P proximity—specifically, the 1.5noC condition (due to its extremely short genomic separation) and the untreated 339CECP condition (due to stable CTCF-CTCF loop formation)—were excluded. In these configurations, the sustained E-P proximity, coupled with the inherent stochasticity of promoter state transitions, effectively decouples transient distance fluctuations from transcription burst onsets, making it challenging for the model to learn a meaningful distance-transcription relationship.

For the 30-second frame rate trajectories, the 3D E-P distance was used as the sole input feature. For the 5-second frame rate trajectories, the higher frequency sampling and shorter acquisition duration required additional transformation of the 3D E-P distance into two additional input features: the discrete temporal derivative (velocity) of 3D E-P distance, and the trajectory-specific localization error estimated via mean-squared displacement (MSD) fitting. E-P distances and velocities were scaled using a MinMaxScaler, while localization errors were standardized.

The neural network architecture began with a Masking layer (mask value = -1.0) to handle variable trajectory lengths, padded to a maximum sequence length of 721 frames for the 30-second frame rate trajectories and 724 frames for the 5-second frame rate trajectories. This was followed by two stacked LSTM layers consisting of 128 and 64 hidden units, respectively, both with a dropout rate of 0.3 to prevent overfitting. The output layer comprised a time-distributed dense layer with a softmax activation to predict the probability of the promoter occupying an active bursting state at each frame.

To mitigate the stochasticity and noise inherent in neural network initialization and training, an ensemble of 10 independent LSTM models was trained using different random seeds. To prevent data leakage, the dataset was split strictly at the trajectory level into 80% training and 20% validation sets, stratified by the density of CPD labels to ensure representative distributions across splits. No individual frames or time-points from the same cell trajectory were shared between the training and validation sets, ensuring that temporal autocorrelation could not artificially inflate model performance. Furthermore, because E-P distances and velocities were scaled, the model was forced to learn event-level temporal dynamics (e.g., local distance dips) rather than confounding condition-specific absolute distances. Prior to training, the raw CPD burst labels were computationally "anchored" to the steepest increase in the smoothed MS2 signal following a local valley, with a fixed burst length of 32 frames to maximize the probability that the label encompasses the true burst onset and to account for the uncertainty of the exact timing of the onset. A 24-frame backward shift was implemented for the 30-second burst labels because E-P interactions likely precede the detectable increase in MS2 signal, particularly given the lower temporal resolution of the 30-second trajectories.

Models were optimized with the Adam optimizer (learning rate = 0.0005, gradient clipvalue = 1.0) and a temporal focal loss function ($\gamma = 1.0$) configured with class weights inversely proportional to the training class frequencies to handle class imbalance, since burst onsets are relatively rare events. The focal loss applied across unmasked time steps $t$ and classes $c \in \{0, 1\}$ was calculated as:

$$\mathcal{L} = -\sum_{t=1}^{T}\sum_{c=0}^{1} \alpha_c (1 - \hat{y}_{t,c})^{\gamma} y_{t,c} \log(\hat{y}_{t,c})$$

where $y_{t,c}$ is the ground-truth binary label, $\hat{y}_{t,c}$ is the predicted probability, and $\alpha_c$ is the frequency-derived class weight. Training was carried out for up to 40 epochs with a batch size of 32, using dynamic learning rate reduction to enable finer weight convergence as validation loss plateaued, and employing early stopping mechanisms to stop training and restore optimal model weights if validation performance stagnated, thereby systematically preventing overfitting.

#### 9.3 Burst onset prediction, positive and negative control

The fundamental prediction task was a frame-by-frame binary classification compared against the CPD labels. Model performance was evaluated independently for every frame. Final predicted burst probabilities were obtained by averaging these frame-by-frame predictions across the 10-model ensemble. To eliminate initial recurrent state artifacts, the first 30 frames (burn-in period) of each trajectory were excluded from analysis.

Analysis was performed using the optimal intersection threshold derived from the precision-recall curve (0.509 for 30-second frame rate trajectories, 0.529 for 5-second frame rate trajectories). Using this threshold, the continuous ensemble predictions were binarized. Discrete burst onsets were then defined as the frames where the prediction probability crossed the threshold from below to above.

To validate the robustness of our temporal alignment and confirm feature dependence, we introduced a synthetic misalignment positive control. For this control, the input 3D E-P distance trajectories were artificially shifted backwards by 4 frames (equivalent to 120 seconds for the 30-second frame rate data) relative to the CPD-derived burst labels prior to training. An identical LSTM neural network was trained on these shifted data. As expected, the model retained its predictive power, but the learned transient dip in the 3D E-P distance pile-up was exactly shifted by -4 frames relative to the predicted burst onsets (**Fig. S16D**). This confirms that the model relies on the temporal alignment of the E-P distance dynamics to make its predictions.

Conversely, to validate that the LSTM predictions were fundamentally driven by E-P spatial dynamics rather than random distance fluctuations, we also trained a negative control model. An identical 10-model LSTM ensemble was trained on the 30-second frame rate trajectories, but with the 3D E-P distance inputs randomly scrambled across time for each trajectory, while preserving the original sequence lengths and target MS2 labels. This scrambled-input ensemble completely failed to predict burst onsets, yielding an AU-ROC of 0.50 (same as the random baseline AU-ROC of 0.5; **Fig. S16E**).

##### 9.4 Inference of the true E-P interaction distance ($R_{\mathrm{E-P}}$)

To infer the true interaction radius ($R_{\mathrm{E-P}}$) from the experimentally measured label distance distribution exactly at the predicted burst onsets, we employed a computational forward-modeling approach. First, we generated extensive ensembles of simulated spatial conformations using the polymer simulations discussed above. From these simulations, we isolated interacting sub-populations by systematically scanning different upper bound thresholds for the physical distance between the enhancer and promoter nodes ($R_{\mathrm{E-P}}$). For each candidate $R_{\mathrm{E-P}}$ threshold, we extracted the corresponding 3D distance distributions between the monomers representing the adjacent fluorescent arrays. We then computationally convolved these simulated label distances with our condition-specific localization errors (estimated via MSD fitting) and measured chromatic aberration residuals. Finally, the resulting simulated 3D label distance distributions were statistically evaluated against the experimental label distance distribution at the predicted burst onsets. The candidate threshold that minimized the discrepancy between the simulated and experimental distributions, an $R_{\mathrm{E-P}}$ of 30 nm (95% CI: 26 to 37 nm), was determined as the most accurate estimate of the true E-P interaction radius underlying transcription activation (**Fig. 3E**).

#### 10 VEPI: Variational Enhancer-Promoter Inference

##### 10.1 List of symbols

| | |
|---|---|
| $s_t$ | Promoter state at time $t$ |
| $s_{0:T}$ | Promoter state trajectory between time $0$ and time $T$ |
| $\boldsymbol{R}_t$ | 3D enhancer-promoter separation vector at time $t$ |
| $R_t$ | 3D enhancer-promoter distance at time $t$ |
| $\boldsymbol{R}_t^{\mathrm{obs}}$ | Observed 3D enhancer-promoter separation vector at time $t$ |
| $\boldsymbol{R}_{0:T}^{\mathrm{obs}}$ | Observed 3D enhancer-promoter separation trajectory between time $0$ and time $T$ |
| $\boldsymbol{R}_{0:T}$ | 3D enhancer-promoter separation trajectory between time $0$ and time $T$ |
| $R_{\mathrm{E-P}}$ | E-P contact radius used to define functional contact |
| $x_t$ | Component of the E-P separation vector at time $t$ |
| $x_t^{\mathrm{obs}}$ | Observed component of the E-P separation vector at time $t$ |
| $\tau_i$ | Time of the $i$-th polymerase loading event in a trajectory |
| $\tau_{0:T}$ | The sequence of polymerase loading times between time $0$ and time $T$ |
| $I_n$ | True MS2 signal at time-point $n$ in a trajectory |
| $I(t)$ | Continuous-time MS2 signal at time $t$ |
| $I_n^{\mathrm{obs}}$ | Observed MS2 signal at time-point $n$ in a trajectory |
| $I^{\mathrm{obs}}(t)$ | Observed continuous-time MS2 signal at time $t$ |
| $I_{0:T}^{\mathrm{obs}}$ | Observed MS2 signal trajectory between time $0$ and time $T$ |
| $K$ | Number of promoter states |
| $J$ | Promoter-state transition-rate matrix |
| $J_{ij}$ | Transition rate to promoter state $i$ from promoter state $j$ ($J_{ii} = 0$) |
| $J_j$ | Exit rate from promoter state $j$ defined as $\sum_i J_{ij}$ |
| $r_i$ | Polymerase loading rate in promoter state $i$ |
| $\rho_i$ | Trajectory-specific polymerase loading rate in promoter state $i$ |
| $\mathrm{CV}_r$ | Coefficient of variation of trajectory-to-trajectory loading-rate variability |
| $L_{ij}$ | Second moment of the trajectory-specific loading rates in states $i$ and $j$ |
| $\pi_i$ | Initial probability of being in promoter state $i$ before the start of the trajectory |
| $T_{\mathrm{Rise}}$ | Rise-time of the MS2 kernel (time to go transcribe through the MS2 array) |
| $T_{\mathrm{Plateau}}$ | Plateau-duration of the MS2 kernel (time to excise the transcript) |
| $I_\phi$ | MS2 intensity scale of a single Pol II transcription event |
| $\Delta t$ | Time between adjacent observations in the discrete transfer-matrix likelihood |
| $m$ | Number of discrete time bins in the finite MS2 memory window |
| $w_\ell$ | Discretized MS2 kernel weight for a Pol II loading event $\ell$ time bins in the past |
| $p_i(t)$ | Probability of being in promoter state $i$ at time $t$ |
| $p_{\mathrm{on}}(t)$ | Probability of being in the ON-state at time $t$ |
| $\gamma_i(t)$ | Variational posterior probability of promoter state $i$ at time $t$ |
| $\gamma_{ij}(t)$ | Variational posterior transition density from state $j$ to state $i$ at time $t$ |
| $N_{ij}$ | Variational posterior expected number of transitions from state $j$ to state $i$ |

| | |
|---|---|
| $\eta(t)$ | Variational posterior density of Pol II loading events at time $t$ |
| $M_{ij}$ | Effective transition rate used in the promoter-state variational posterior |
| $f_i(t)$ | Time-dependent tilt applied to promoter state $i$ in the variational posterior |
| $\bar{r}_t$ | Effective loading rate used in the loading-event variational posterior |
| $\lambda$ | Reference homogeneous Poisson rate used to define loading-event point-process densities |
| $\phi(t)$ | Intensity kernel for the MS2 signal representing the intensity of a single Pol II transcription event |
| $I_0$ | Integrated MS2 kernel, $\int \phi(t)\mathrm{d}t$ |
| $c(t)$ | MS2 kernel auto-correlation, $\int \phi(x)\phi(t+x)\mathrm{d}x$ |
| $\Lambda(t)$ | Loading-rate correlation function |
| $C(x)$ | Clamp function on $[0, T_{\mathrm{Rise}}]$ used in the MS2 autocorrelation expression |
| $\sigma_I^2$ | Variance of the MS2 signal noise |
| $\boldsymbol{\sigma}^2$ | Variance of the E-P separation vector noise |
| $I_{\mathrm{offset}}$ | Constant offset of the MS2 signal |
| $\mathcal{GP}\left(\boldsymbol{\mu}(t), K(t,t')\right)$ | Gaussian Process with mean function $\boldsymbol{\mu}(t)$ and kernel $K(t,t')$ |
| $\mathrm{diag}(\boldsymbol{\sigma}^2)$ | Diagonal matrix with the elements of $\boldsymbol{\sigma}^2$ on the diagonal |
| $k_{\mathrm{on}}$ | ON-rate of the promoter state transition in an E-P-driven two-state promoter model |
| $k_{\mathrm{on}}^{\mathrm{eff}}$ | Apparent ON-rate of the promoter state transition in a two-state promoter model without knowledge of E-P |
| $k_{\mathrm{off}}$ | OFF-rate of the promoter state transition in a two-state promoter model |

#### 10.2 Model overview

We chose a minimal model to enable tractability for trajectory-level inference and to ensure identifiability of as many parameters and processes as possible given the noisy and indirect nature of live-cell locus tracking and nascent MS2 transcriptional reporters. Our model of E-P-driven transcription consists of a two-state promoter with an ON-rate driven by E-P contact. The path of the E-P trajectory was modeled as a Gaussian Process (GP) to capture the non-Rouse behavior of the observed MSD with minimal assumptions about the mechanistic details. Conditionally on a trajectory of the E-P separation vector $\boldsymbol{R}_{0:T}$, the promoter dynamics evolve according to

$$\frac{\mathrm{d}}{\mathrm{d}t}p_{\mathrm{on}}(t) = -(k_{\mathrm{off}} + k_{\mathrm{on}}\theta(R_{\mathrm{E-P}} - R_t))p_{\mathrm{on}}(t) + k_{\mathrm{on}}\theta(R_{\mathrm{E-P}} - R_t)\,, \tag{16}$$

where $\theta$ is the heaviside step function and we have initial condition $p_{\mathrm{on}}(0) = \pi_1$. We modeled polymerase loading times $\tau_i$ as an inhomogeneous Poisson process with a loading rate given by the promoter state $s_t$

$$P(\text{Pol II loading in interval } [t, t+\mathrm{d}t]) = \begin{cases} r_{\mathrm{loading}}\mathrm{d}t & \text{if } s_t = 1 \\ 0 & \text{otherwise} \end{cases}. \tag{17}$$

This choice naturally produces bursty transcription while allowing an E-P drive. MS2 intensity was modeled as a deterministic function of the polymerase loadings

$$I(t) = \sum_i \phi(t - \tau_i)\,, \tag{18}$$

where $\phi(t)$ is the intensity trajectory of a single Pol II loading event and is parameterized by $\theta_\phi = [T_{\mathrm{Rise}}, T_{\mathrm{Plateau}}, I_\phi]$, the time to transcribe the MS2 array, time to transcribe the rest of the gene and splice the transcript, and the intensity of the folded MS2 array. This model of MS2 neglects the randomness in Pol II translocation along the gene and the splicing process. This approximation was essential to keeping the model scalable, as explicit modeling of these processes would require large state-space models. We note that this deterministic approach has been found to be successful in past studies [87, 88].

The measurement model describes how the observed data (e.g., MS2 intensity) relate to the underlying latent variables (e.g., promoter states, E-P proximity). In our dataset, there are two types of simultaneous measurements:

The E-P separation vector and the MS2 intensity. For simplicity, we assume both to be corrupted by Gaussian noise, and the MS2 signal is allowed to have an additional constant offset. The resulting measurement model reads

$$I^{\mathrm{obs}}(t) = I(t) + I_{\mathrm{offset}} + \epsilon_I(t)\,, \tag{19}$$

$$\boldsymbol{R}^{\mathrm{obs}}(t) = \boldsymbol{R}(t) + \boldsymbol{\epsilon}_R(t)\,, \tag{20}$$

where $\epsilon_I(t) \sim \mathcal{GP}(\mathbf{0}, \delta(t-t')\sigma_I^2)$ and $\boldsymbol{\epsilon}_R(t) \sim \mathcal{GP}(\mathbf{0}, \delta(t-t')2\mathrm{diag}(\boldsymbol{\sigma}^2))$ are the Gaussian noise on the MS2 signal and noise for each component of the E-P separation vector, respectively. Defining the transcription processes $\Phi = \{s_{0:T}, \tau_{0:T}\}$ and associated parameters $\vartheta = \{r_{\mathrm{loading}}, k_{\mathrm{on}}, k_{\mathrm{off}}, \boldsymbol{\pi}\}$ we can write the joint distribution of the full model as

$$p(\boldsymbol{R}^{\mathrm{obs}}_{0:T}, I^{\mathrm{obs}}_{0:T}, \boldsymbol{R}_{0:T}, \Phi, \vartheta) = p(I^{\mathrm{obs}}_{0:T}|\Phi, \vartheta, \boldsymbol{R}_{0:T})p(\boldsymbol{R}^{\mathrm{obs}}_{0:T}|\boldsymbol{R}_{0:T})p(\boldsymbol{R}_{0:T})p(\Phi)p(\vartheta)\,, \tag{21}$$

where we defined a shorthand for the conditional MS2 likelihood

$$p(I^{\mathrm{obs}}_{0:T}|\Phi, \vartheta, \boldsymbol{R}_{0:T}) = p(I^{\mathrm{obs}}_{0:T}|\tau_{0:T}, \vartheta_\phi, \sigma_I^2)p(\tau_{0:T}|r_{\mathrm{loading}}, s_{0:T})p(s_{0:T} \mid k_{\mathrm{on}}, k_{\mathrm{off}}, \boldsymbol{\pi}, \boldsymbol{R}_{0:T})\,. \tag{22}$$

To render inference tractable, we found it useful to rewrite this model in a form that utilizes the fact that we measure both the E-P distance and the MS2 intensity. Using Bayes' theorem, we can rewrite the E-P distance part of Equation (21) as a posterior distribution of only the observed E-P data

$$p(\boldsymbol{R}^{\mathrm{obs}}_{0:T}, I^{\mathrm{obs}}_{0:T}, \boldsymbol{R}_{0:T}, \Phi, \vartheta) = p(I^{\mathrm{obs}}_{0:T}|\Phi, \vartheta, \boldsymbol{R}_{0:T})p(\boldsymbol{R}_{0:T} \mid \boldsymbol{R}^{\mathrm{obs}}_{0:T})p(\boldsymbol{R}^{\mathrm{obs}}_{0:T})p(\vartheta)p(\Phi)\,. \tag{23}$$

This is a convenient form as $p(\boldsymbol{R}_{0:T} \mid \boldsymbol{R}^{\mathrm{obs}}_{0:T})$ is directly available using the well-established framework of Gaussian Process Regression (GPR) [89]. By marginalizing over the E-P posterior, we can now define an effective MS2 model which represents the distribution of all transcription-related quantities given the observed MS2 and E-P data

$$p(\boldsymbol{R}^{\mathrm{obs}}_{0:T}, I^{\mathrm{obs}}_{0:T}, \Phi, \vartheta) = p(\boldsymbol{R}^{\mathrm{obs}}_{0:T})\langle p(I^{\mathrm{obs}}_{0:T}|\Phi, \vartheta, \boldsymbol{R}_{0:T})\rangle_{\boldsymbol{R}_{0:T}\sim p(\boldsymbol{R}_{0:T}|\boldsymbol{R}^{\mathrm{obs}}_{0:T})}p(\vartheta)p(\Phi)\,. \tag{24}$$

This is powerful, as it means we can solve the MS2 inference problem separately from the E-P inference problem. It is worth mentioning that this form of the joint distribution is possible because we work with a model where there is no physical feedback from transcription to E-P motion.

Note that the path average in (24) is challenging to calculate in practice. In what follows, we will approximate this average by considering the promoter switching dynamics to be slow on the timescale of posterior E-P distance fluctuations. More concretely, we replace $\theta(R_{\mathrm{E-P}} - R_t)$ by $\langle \theta(R_{\mathrm{E-P}} - R_t \rangle_{\boldsymbol{R}(t)\sim p(\boldsymbol{R}(t)|\boldsymbol{R}^{\mathrm{obs}}_{0:T})}$ in all calculations. This approximation was made for tractability, and we found that we were able to recover the functional relationship between $k_{\mathrm{on}}$ and $R_{\mathrm{E-P}}$ despite it (**Fig. S20**). Future work will explore more sophisticated methods for handling this averaging.

We now show how we evaluated the different objects appearing in Equation (24). The calculation has two parts: evaluation of the E-P posterior and the E-P-marginalized MS2 posterior. The first boils down to a standard Gaussian Process Regression, while the latter required the development of a tailored approach to MS2 inference.

##### 10.3 E-P posteriors from Gaussian Process Regression (GPR)

Since each component of the E-P separation vector evolves independently, the inference for the E-P posteriors can be run for each coordinate separately and then combined to produce trajectories and evaluate averages. In what follows, we therefore consider the one-dimensional case for clarity.

We consider the case where we obtained a noisy time-series as a component of the E-P separation vector at times $\{t_i\}\,, i = 1, 2, \ldots, N$

$$x_i^{\mathrm{obs}} = x_i + \epsilon_i\,, \quad \langle \epsilon_i \rangle = 0\,, \quad \langle \epsilon_i \epsilon_j \rangle = 2\sigma^2 \delta_{ij}\,, \tag{25}$$

where $\sigma^2$ is the single-locus localization error for the component in question. Given a Gaussian process prior, we seek the posterior distribution of the sample process that generated the observations $x_i$ at a different set of times $\{u_i\}, i = 1, 2, \ldots, n$ which may or may not be coincident with $\{t_i\}$. For this, we need the moments

$$\langle x_i x_j \rangle = K(u_i - u_j) \equiv \boldsymbol{\Sigma}_a\,, \qquad (n \times n) \tag{26}$$

$$\langle x_i y_j \rangle = K(u_i - t_j) \equiv \boldsymbol{\Sigma}_b\,, \qquad (n \times N) \tag{27}$$

$$\langle y_i y_j \rangle = K(t_i - t_j) + 2\sigma^2\delta_{ij} \equiv \boldsymbol{\Sigma}_c\,, \qquad (N \times N) \tag{28}$$

from which we can construct the joint distribution of measurement and true process, which may be seen to have the block matrix form

$$p\begin{pmatrix} X \\ Y \end{pmatrix} = \mathcal{N}\left(0, \begin{pmatrix} \boldsymbol{\Sigma}_a & \boldsymbol{\Sigma}_b \\ \boldsymbol{\Sigma}_b^T & \boldsymbol{\Sigma}_c \end{pmatrix}\right)\left[\begin{pmatrix} X \\ Y \end{pmatrix}\right]\,, \tag{29}$$

where we used $X$ and $Y$ to refer to the n-dimensional and N-dimensional vectors of the true and observed trajectories.

We may now obtain the posterior from the formula for the conditional distribution of a multivariate Gaussian

$$p(X|Y) = \mathcal{N}\left(\hat{\mu}, \hat{\mathbf{\Sigma}}\right)[X]\,. \tag{30}$$

with

$$\hat{\mu} = \mathbf{\Sigma}_b \mathbf{\Sigma}_c^{-1} Y\,, \tag{31}$$

$$\hat{\mathbf{\Sigma}} = \mathbf{\Sigma}_a - \mathbf{\Sigma}_b \mathbf{\Sigma}_c^{-1} \mathbf{\Sigma}_b^T\,. \tag{32}$$

This multivariate Gaussian describes the posterior distribution of trajectories that are consistent with the observed trajectory under the noise and process model specified by the MSD. Using this posterior, one can compute confidence intervals for the E-P separation vector at any time of interest, as well as sample possible trajectories that are consistent with the measurements.

For numerical stability, matrix and vector products with the inverse $\mathbf{\Sigma}_c^{-1}$ were solved using triangular solves and the Cholesky decomposition rather than explicitly forming the inverse [89]. Finally, to efficiently do inference on several trajectories, we used `jax` [90] for efficient batch GPU processing. This had the downside of requiring fixed-size inputs and would not be able to handle $\{t_i\}$, which varied between trajectories. We accommodate this by first replacing missing values with placeholder zeros and padding tracks shorter than the longest track with zeros. For a given track, we then set the measurement noise of the missing values to a large constant to project them out from the inference.

#### 10.4 MS2 state-reconstruction with variational inference

While E-P reconstruction relied on standard tools from Gaussian process regression, the MS2 inference is complicated by the nesting of stochastic processes of various kinds. Direct inference on the model was found to be intractable, with the transfer-matrix approach in Section 10.6 being an exception for long intervals between observations and few Pol II loadings. With the goal of obtaining a framework that worked in any regime, we resorted to approximations through variational inference [91].

The derivation below operates on the E-P-marginalized MS2 model introduced in Equation (24); equivalently, all E-P-dependent promoter rates should be understood as the effective rates obtained after averaging over $p(\boldsymbol{R}_{0:T} \mid \boldsymbol{R}_{0:T}^{\text{obs}})$. Section 10.4.1 discusses our model of the MS2 data and briefly introduces its Bayesian network representation. Section 10.4.2 introduces variational inference and mean-field variational inference for Bayesian networks, which is the method we employed in this work. In Section 10.4.3, we derive the necessary averages to implement variational inference for our MS2 model, and in Section 10.5, we describe how to calibrate the observation model to data by fitting the autocorrelation of the MS2 signal. Finally, in Section 10.8 we provide a walk-through of our implementation of the inference scheme as well as benchmarks exploring its validity.

##### 10.4.1 MS2 model specification

We assume that the MS2 data arose from a hierarchical model where promoter state $s_t$ governs Pol II loadings $\tau_i$, which yields observed MS2 $I_k^{\text{obs}}$ through convolution with a signal kernel. For the variational derivation, we allow an arbitrary $K$-state promoter model. The joint probability of the MS2 measurements and the latent processes in the effective model can be written as

$$p(I_{0:T}^{\text{obs}}, \tau_{0:T}, s_{0:T}, J, r, \pi \mid \boldsymbol{R}_{0:T}^{\text{obs}}) = p(I_{0:T}^{\text{obs}} \mid \tau_{0:T}) p(\tau_{0:T} \mid s_{0:T}, r) p(s_{0:T} \mid J, \pi, \boldsymbol{R}_{0:T}^{\text{obs}}) p(J) p(r) p(\pi)\,, \tag{33}$$

where $J$, $r$, and $\pi$ are the promoter transition rates, Pol II loading rates, and initial promoter state probabilities, respectively. In the formulas below, we suppress the explicit conditioning on $\boldsymbol{R}_{0:T}^{\text{obs}}$ and write the effective promoter path density as $p(s_{0:T} \mid J, \pi)$. To provide an intuitive framework for the results to come, we will develop our formulas in the context of Bayesian networks [92]. A Bayesian network is a graphical representation of a joint distribution like Equation (33). In this formalism, each latent random variable ($\tau_{0:T}$ and $s_{0:T}$) gets assigned a node, and an observed node ($I_k^{\text{obs}}$) is colored blue. Parameters ($J$, $r$, $\pi$) get assigned a dot, and each conditional probability term introduces an arrow from the conditioning variables (parents) to the arguments (children). With these rules, the joint probability in Equation (33) becomes the graph shown in **VEPI Fig. 1A**. We will refer to this graph when deriving the variational inference scheme, and it also serves to introduce the variable names employed. We now present the functional form of each conditional probability term in our model.

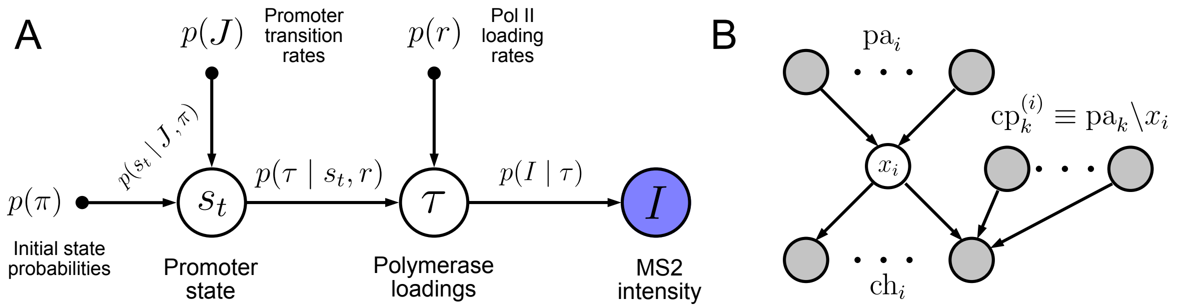

**VEPI figure 1: A**: Bayesian network for our model of the MS2 signal. The promoter state $s_t$ relies on an initial probability $\pi_i$ (Equation (48)) and transition rates $J_{ij}$ (Equation (46)) for its specification through $p(s_t \mid J, \pi)$ (Equation (41)). The promoter state specifies the rate of polymerase loading $\tau$ through $p(\tau \mid s_t, r)$ (Equation (37)) given a loading rate $r_i$ prior (Equation (47)) for each state. Finally, knowing the polymerase loading events, the observed MS2 signal $I_k^{\mathrm{obs}}$ is obtained by convolution with a kernel and addition of Gaussian measurement noise, which yields a Gaussian signal likelihood $p(I_{0:T}^{\mathrm{obs}} \mid \tau_{0:T})$ (Equation (36)). **B**: Definition of parents, children, and co-parents of a node in a Bayesian graph. The set of parents, children, and co-parents forms the Markov blanket of a node $x_i$ (gray nodes). The notation $\mathrm{pa}_k \backslash x_i$ means all parent nodes of node $x_k$ excluding $x_i$.

###### 10.4.1.1 MS2 likelihood $p(I_{0:T}^{\mathrm{obs}} \mid \tau_{0:T})$

We assume observed MS2 $I_k^{\mathrm{obs}}$ at times $t_k \in [0, T]$ to arise from noisy measurement of Pol II loadings at times $\tau_i$ through the deterministic function $\phi(t)$ which gives the signal expected from a single loading event

$$I_k^{\mathrm{obs}} = I_{\mathrm{offset}} + \sum_i \phi(t_k - \tau_i) + \epsilon_k\,, \quad \langle \epsilon_i \epsilon_j \rangle = \sigma_I^2 \delta_{ij}\,. \tag{34}$$

$\phi(t)$ will depend on the position and length of the MS2 array as well as the transcription rate of Pol II and the rate of release of the nascent transcript. For our construct, a natural choice is the rise and plateau form

$$\phi(t) = \begin{cases} I_\phi \frac{t}{T_{\mathrm{Rise}}}\,, & 0 \le t < T_{\mathrm{Rise}} \\ I_\phi\,, & T_{\mathrm{Rise}} \le t \le T_{\mathrm{Plateau}} + T_{\mathrm{Rise}} \\ 0\,, & \text{otherwise} \end{cases}\,, \tag{35}$$

where we expect $T_{\mathrm{Rise}}$ to be approximately the MS2 array length divided by the Pol II transcription speed and $T_{\mathrm{Plateau}}$ to be approximately the length of the gene after the array plus the average time to excise the nascent transcript. We take Gaussian errors on MS2 and thus get a Gaussian likelihood for the observations

$$p(I_{0:T}^{\mathrm{obs}} \mid \tau_{0:T}) = \frac{1}{(2\pi\sigma_I^2)^{N/2}} \exp\left(-\sum_k \frac{\left(I_k^{\mathrm{obs}} - I_{\mathrm{offset}} - \sum_i \phi(t_k - \tau_i)\right)^2}{2\sigma_I^2}\right)\,. \tag{36}$$

We shall take $\sigma_I^2$, $I_{\mathrm{offset}}$, $T_{\mathrm{Rise}}$, $T_{\mathrm{Plateau}}$, and $I_\phi$ to be hyperparameters known a priori, and they are therefore not conditioned on in the likelihood. We show how we obtain them through a fit to the MS2 autocorrelation in Section 10.5.

###### 10.4.1.2 Pol II loading event density $p(\tau \mid s_t, r)$

The loading events are assumed to follow an inhomogeneous Poisson point process with rate $r_k \delta_{k,s_t}$ (We employ Einstein summation convention, so repeated indices are to be summed over) where $s_t$ is the state of the promoter at time $t$ and $r_i$ gives the loading rate of the $i$'th promoter state. The probability for a set of loadings is then given by the path density of an inhomogeneous Poisson point process

$$p(\tau \mid s_t, r) = \exp\left(-\int_0^T [r_{s_t} - \lambda]\mathrm{d}t + \sum_i \log\left(\frac{r_{s_{\tau_i}}}{\lambda}\right)\right)\,, \tag{37}$$

where we normalize with respect to a homogeneous point process with rate $\lambda$ as may be seen from

$$\begin{aligned} \langle p(\tau \mid s_t, r)\rangle_\lambda &= \exp\left(-\int_0^T [r_{s_t} - \lambda]\mathrm{d}t\right) \left\langle \exp\left(\sum_i \log\left(\frac{r_{s_{\tau_i}}}{\lambda}\right)\right)\right\rangle_\lambda\,, \\ &= \exp\left(-\int_0^T [r_{s_t} - \lambda + \lambda - r_{s_t}]\mathrm{d}t\right)\,, \\ &= 1\,. \end{aligned} \tag{38}$$

To arrive at the second line we used the generating functional for a Poisson process

$$\left\langle \exp\left(\xi \sum_i f(t_i)\right)\right\rangle_\lambda = \exp\left(\int_0^T \left(\mathrm{e}^{\xi f(t)} - 1\right) \lambda \mathrm{d}t\right) . \tag{39}$$

###### 10.4.1.3 Promoter state path density $p(s_t \mid J, \pi)$

The $K$ promoter states are assumed to follow a continuous-time Markov chain (CTMC). We use the convention that $J_{ij}$ is the rate of transitioning from state $j \to i$ and $J_i$ is the exit rate from state $i$ defined as

$$J_i = \sum_{j=1}^{K} J_{ji} . \tag{40}$$

For a path with $n$ jumps at ordered times $0 < t_1 < \ldots < t_n < T$, define $t_0 = 0$, $t_{n+1} = T$, and let the state on the interval $[t_m, t_{m+1})$ be $i_m$. At a jump time $t_m$, the notation $s_{t_m^-} = i_{m-1}$ and $s_{t_m^+} = i_m$ denotes the states immediately before and after the jump. The path probability density with respect to the ordered jump times can then be written in the exponential form

$$p(s_{0:T} \mid J, \pi) = \pi_{s_0} \exp\left(-\int_0^T J_{s_t} \mathrm{d}t + \sum_{m=1}^{n} \log J_{s_{t_m^+} s_{t_m^-}}\right) . \tag{41}$$

With this convention, a sum over path space means

$$\sum_{\{s_t\}} F[s] \equiv \sum_{n=0}^{\infty} \sum_{i_0=1}^{K} \sum_{\substack{i_1,\ldots,i_n \\ i_m \neq i_{m-1}}} \int_{0<t_1<\ldots<t_n<T} F(i_0, \ldots, i_n; t_1, \ldots, t_n) \mathrm{d}t_1 \cdots \mathrm{d}t_n . \tag{42}$$

This definition is often most useful through its generating functional. For a dwell-time tilt $f_i(t)$ and a jump tilt $g_{ij}(t)$, define

$$\mathcal{G}_{a,b}[f, g] \equiv \left\langle \exp\left(a \int_0^T f_{s_t}(t)\mathrm{d}t + b \sum_{m=1}^{n} g_{s_{t_m^+} s_{t_m^-}}(t_m)\right)\right\rangle_{p(s_{0:T}\mid J,\pi)} = \mathbf{1}^T \mathcal{T} \exp\left(\int_0^T A^{(a,b)}(t)\mathrm{d}t\right) \boldsymbol{\pi} , \tag{43}$$

where $\mathcal{T}$ denotes time ordering, and the tilted generator is

$$A_{ij}^{(a,b)}(t) = \begin{cases} J_{ij}\mathrm{e}^{b g_{ij}(t)} , & i \neq j , \\ a f_i(t) - J_i , & i = j . \end{cases} \tag{44}$$

This is followed by expanding the time-ordered exponential into ordered products of off-diagonal jump factors separated by diagonal survival factors, which reproduces the path sum in Equation (42). Setting $a = b = 0$ gives the ordinary CTMC generator $A_{ij} = J_{ij} - \delta_{ij} J_j$, whose columns sum to zero. Therefore

$$\sum_{\{s_t\}} p(s_{0:T} \mid J, \pi) = \mathbf{1}^T \mathrm{e}^{AT} \boldsymbol{\pi} = \mathbf{1}^T \boldsymbol{\pi} = 1 . \tag{45}$$

All dwell-time and jump averages used below can be obtained by differentiating Equation (43) with respect to the corresponding tilts.

###### 10.4.1.4 Parameter priors $p(J)$, $p(\pi)$, $p(r)$

We choose priors for $\pi$, $J$, and $r$ which are conjugate in our variational inference scheme. This will turn out to be gamma distributions for $J$ and $r$

$$p(J_{ij}) \propto J_{ij}^{\beta_{ij}-1} \mathrm{e}^{-\theta_{ij} J_{ij}} , \tag{46}$$

$$p(r_i) \propto r_i^{\nu_i - 1} \mathrm{e}^{-\chi_i r_i} , \tag{47}$$

with shape parameters $\beta_{ij}$, $\nu_i$ and rate parameters $\theta_{ij}$, $\chi_i$ and a Dirichlet distribution for $\pi$

$$p(\pi) \propto \pi_i^{\kappa_i - 1} , \tag{48}$$

where $\kappa_i$ is the concentration parameter.

##### 10.4.2 Mean-field variational inference

The previous section introduced the joint distribution of the model. In this section, we will describe the variational inference approach used to take this joint distribution and approximate the posterior distribution. The derivations will closely follow those of [93].

The main goal of Bayesian inference is to obtain the posterior distribution of a set of random variables $X$ in a model conditioned on observations $Y$. This is accomplished through Bayes' rule*

$$P(X \mid Y) = \frac{P(Y, X)}{P(Y)}\,, \tag{49}$$

where $P(Y, X)$ is the model specified through a joint distribution of observed and latent variables, and $P(Y)$ is often referred to as the evidence.

While Equation (49) may be seen to provide the posterior up to a proportionality constant by insertion of Equation (33), this does not imply that the inference problem is solved by specifying a joint distribution $P(X, Y)$. This is because any average of interest would require integration of $P(Y, X)$ over a subset of the latent variables $X$, which is completely intractable for our model. We therefore seek an approximation of $P(X \mid Y)$ from a family of distributions $Q(X)$ that admit tractable calculations of averages and quantities of interest. To find the best approximation within the family $Q$, some metric of proximity between distributions must be invoked. Variational inference arises by choosing the metric to be the Kullback–Leibler divergence (KLD) between the true posterior and the approximation

$$\begin{aligned}
\mathrm{D_{KL}(Q||P)} &= \sum_X Q(X) \log\left(\frac{Q(X)}{P(X \mid Y)}\right)\,, \\
&= -\left\langle \log P(X \mid Y)\right\rangle_Q + \left\langle \log Q(X)\right\rangle_Q\,, \\
&= \log P(Y) - \left\langle \log P(X, Y)\right\rangle_Q + \left\langle \log Q(X)\right\rangle_Q\,, \\
&= \log P(Y) - \mathcal{L}_Q\,,
\end{aligned} \tag{50}$$

where we defined the Evidence Lower Bound (ELBO)

$$\mathcal{L}_Q = \langle \log P(X, Y)\rangle_Q - \langle \log Q(X)\rangle_Q\,. \tag{51}$$

Since $P(Y)$ does not depend on $Q$, it is clear from Equation (50) that minimizing the KLD is the same as maximizing the ELBO. We may thus obtain an optimal approximation to the posterior by maximizing the ELBO. When the approximation is perfect, the ELBO will be equal to the evidence, and for any finite KLD, the ELBO can be seen to bound the evidence through

$$\begin{aligned}
\mathcal{L}_Q &= \log P(Y) - \mathrm{D_{KL}(Q||P)}\,, \\
&\leq \log P(Y)\,,
\end{aligned} \tag{52}$$

which uses the fact that the KLD is non-negative. We therefore see that not only does maximizing the ELBO provide an approximation to the intractable posterior within the family $Q(X)$. The value of the ELBO at the optimum will be a bound on the true evidence of the model, which may be used as an approximation to the evidence for model comparison and goodness of fit.

It is not always straightforward to decide on which $Q$ to use as the approximation to the posterior. Mean-field variational inference addresses this question by providing a flexible choice by assuming that they factorize

$$Q(X) = \prod_{x_i} q(x_i)\,. \tag{53}$$

Note that this need not imply that the various variables in the model have no dependence on each other. As we will see, a given $q(x_i)$ will depend on the other $q$'s through averages, hence the name mean-field. The factorized family therefore provides a rich class of models which can capture many complex dependencies between the variables of interest. It is still an approximation, which is why we test its validity in Section 10.8.

If $Q$ has the factorized form, the ELBO reads

$$\mathcal{L}_Q = \langle \log P(X, Y)\rangle_Q - \sum_i \langle \log q(x_i)\rangle_q\,. \tag{54}$$

Isolating terms that depend on a particular distribution $q(x_i)$, we get

$$\begin{aligned}
\mathcal{L}_Q &= \sum_{x_i} q(x_i) \langle \log P(X, Y)\rangle_{Q\setminus i} - \sum_{x_i} q(x_i) \log q(x_i) - \sum_{i\neq j}\sum_{x_i} q(x_i) \log q(x_i)\,, \\
&= -\, D(q(x_i)||q^*(x_i)) + \text{terms without } q(x_i)\,,
\end{aligned} \tag{55}$$

*We write the version appropriate to latent variable models. The most common version in the literature is to split the joint distribution into a likelihood and prior using the product rule $P(Y, X) = P(Y \mid X)P(X)$, but this form is not so natural when there are multiple latent variables as in our model.

where $\langle\rangle_{Q\backslash i}$ denotes averaging over all variables except $x_i$ under $Q$ and we introduced the distribution

$$q^*(x_i) = \frac{1}{Z}\exp\left(\langle\log P(X,Y)\rangle_{Q\backslash i}\right) \,, \tag{56}$$

where $Z$ is a normalization constant. We see that $\mathcal{L}_Q$ is maximized with respect to $q(x_i)$ when we set $q(x_i) \to q^*(x_i)$, which means that we may directly obtain the optimum distribution if we can compute the average appearing in Equation (56).

For a given $q$, the relevant terms in $\langle\log P(X,Y)\rangle_{Q\backslash i}$ will only contain factors with explicit dependence on $x_i$. If the model is specified using a graphical model, these terms are seen to be the edges from the parents of $x_i$ to $x_i$ as well as all edges from $x_i$ to its children. The edges to the children will contain variables which are the co-parents of $x_i$, meaning the specification of $q^*$ requires averages over the so-called Markov blanket defined by the parents, children, and co-parents of a node (see **Fig. 1B** for a graphical definition of the Markov blanket). Writing Equation (56) in terms of the Markov blanket, $q^*(x_i)$ becomes

$$q^*(x_i) = \frac{1}{Z}\exp\left(\langle\log P(x_i \mid \mathrm{pa_i})\rangle_Q + \sum_{k\in\mathrm{ch}_i}\langle\log P(x_k \mid \mathrm{pa}_k)\rangle_Q\right) \,. \tag{57}$$

###### 10.4.3 Variational inference for MS2

We now seek to implement mean-field variational inference for our model of MS2. Doing so amounts to identifying the form of the distributions $q^*(x_i)$ for each variable, as well as a means to compute the averages required to evaluate the $q^*(x_i)$'s. In the following, we do so for each variable in the MS2 model, starting at the prior nodes and working our way towards $q(\tau)$ in **Fig. 1A**.

###### 10.4.3.1 Initial probability posterior $q(\pi)$

The optimality condition (Equation (57)) for $x_i = \pi_i$ reads

$$q^*(\boldsymbol{\pi}) = \frac{1}{Z}\exp\left(\langle\log p(s_t \mid J,\pi)\rangle_Q\right) p(\boldsymbol{\pi}) \,. \tag{58}$$

The only term in $p(s_t \mid R,\pi)$ containing $\pi$ is the pre-factor (see Equation (41)). Absorbing the rest of the terms into $Z$ and inserting Equation (48) for $p(\pi_i)$, we get the simple expression

$$\begin{aligned} q^*(\boldsymbol{\pi}) =& \frac{1}{Z}\exp\left(\langle\delta_{s_0,i}\rangle_{q(s_t)}\log\pi_i\right) p(\boldsymbol{\pi}) \,, \\ =& \frac{1}{Z}\prod_i \pi_i^{\langle\delta_{s_0,i}\rangle_{q(s_t)}+\kappa_i-1} \,. \end{aligned} \tag{59}$$

Defining the posterior probability of the promoter state

$$\gamma_i(t) = \langle\delta_{s_t,i}\rangle_{q(s_t)} \,, \tag{60}$$

we see that the variational distribution for the initial state probabilities, which maximizes the ELBO is a Dirichlet distribution with concentration parameter

$$\bar\kappa_i^* = \gamma_i(0) + \kappa_i \,. \tag{61}$$

We will also require the following average for the other distributions

$$\langle\log\pi_i\rangle_{q(\pi)} = \psi(\bar\kappa_i) - \psi\left(\bar\kappa_0\right) \,, \tag{62}$$

where $\bar\kappa_0 = \sum_{i=1}^K \bar\kappa_i$ and $\Gamma$ and $\psi$ are the Gamma function and digamma function respectively.

###### 10.4.3.2 Transition rate posterior $q(J)$

The optimality condition (Equation (57)) for $x_i = J_{ij}$ reads

$$q^*(J_{ij}) = \frac{1}{Z}\exp\left(\langle\log p(s_t \mid J,\pi)\rangle_Q\right) p(J_{ij}) \,. \tag{63}$$

$J_{ij}$ appears in two places in $\langle\log p(s_t \mid J,\pi)\rangle_Q$. The first term is straightforward and may be written in terms of $\gamma_i(t)$, the state posterior

$$\gamma_i(t) = \langle\delta_{s_t,i}\rangle_{q(s_t)} \,, \tag{64}$$

to get

$$\langle J_{s_t}\rangle_{q(s_t)} = \sum_j \gamma_j(t) J_j = \sum_j \gamma_j(t) \sum_{i=1}^{K} J_{ij}\,, \tag{65}$$

while the second term may be expressed in terms of the jump average

$$N_{ij} = \left\langle \sum_m \delta_{s_{t_m^+},i} \delta_{s_{t_m^-},j} \right\rangle_{q(s_t)} = \int_0^T \gamma_{ij}(t)\mathrm{d}t\,, \tag{66}$$

which we elaborate on how to compute in the promoter-state posterior below. Assuming them to be known for now, $q^*(J_{ij})$ can be shown to be of the form

$$q^*(J_{ij}) = \frac{1}{Z} J_{ij}^{N_{ij}} \exp\left(-J_{ij}\int_0^T \gamma_j(t)\mathrm{d}t\right) p(J_{ij})\,, \tag{67}$$

which is seen to be a gamma distribution, like the prior, but with updated shape and scale parameters

$$\bar\beta_{ij} = N_{ij} + \beta_{ij}\,, \tag{68}$$

$$\bar\theta_{ij} = \int_0^T \gamma_j(t)\mathrm{d}t + \theta_{ij}\,. \tag{69}$$

Intuitively, $q^*(J_{ij})$ can be thought of as the posterior which arises from having observed $N_{ij}$ state transitions over a duration $\int_0^T \gamma_j(t)\mathrm{d}t$. The formula for the posterior mean $\langle J_{ij}\rangle_{q(J)}$ is

$$\langle J_{ij}\rangle_{q(J)} = \bar\beta_{ij}/\bar\theta_{ij}\,, \tag{70}$$

which conveys the intuition that the best guess for the transition rate of the promoter state from $j \to i$ is obtained by counting the number of transitions of that type and dividing by the time spent in the state $j$ it came from. The prior may also be seen in this light as a set of $\beta_{ij}$ pseudo-observations over a duration $\theta_{ij}$. For the other distributions, we shall also need the average

$$\langle \log J_{ij}\rangle_{q(J)} = \psi(\bar\beta_{ij}) - \log\bar\theta_{ij} \tag{71}$$

###### 10.4.3.3 Loading rate posterior $q(r)$

The optimality condition (Equation (57)) for $x_i = r_i$ reads

$$q^*(r_i) = \frac{1}{Z}\exp\left(\langle \log p(\tau \mid s_t, r)\rangle_{q(s_t),q(\tau)}\right) p(r_i)\,. \tag{72}$$

Inserting the Poisson point-process density from Equation (37), $r_i$ appears in two places in $\langle \log p(\tau \mid s_t, r)\rangle_{q(s_t),q(\tau)}$. The first term is the dwell-time exposure in state $i$ and may be written in terms of $\gamma_i(t)$,

$$\left\langle \int_0^T r_{s_t}\mathrm{d}t \right\rangle_{q(s_t)} = \sum_i r_i \int_0^T \gamma_i(t)\mathrm{d}t\,. \tag{73}$$

The second term contains the loading events and may be expressed using the posterior loading-event density $\eta(t)$

$$\eta(t) = \left\langle \sum_i \delta(t-\tau_i) \right\rangle_{q(\tau)}\,, \tag{74}$$

which we show how to compute below. In terms of $\eta(t)$, we get

$$\left\langle \sum_\ell \log r_{s_{\tau_\ell}} \right\rangle_{q(s_t),q(\tau)} = \sum_i \log r_i \int_0^T \eta(t)\gamma_i(t)\mathrm{d}t\,. \tag{75}$$

Assuming these averages to be known for now, $q^*(r_i)$ can be shown to be of the form

$$\begin{aligned} q^*(r_i) &= \frac{1}{Z}\exp\left(-r_i\int_0^T \gamma_i(t)\mathrm{d}t + \log r_i \int_0^T \eta(t)\gamma_i(t)\mathrm{d}t\right) p(r_i)\,, \\ &= \frac{1}{Z} r_i^{\nu_i + \int_0^T \eta(t)\gamma_i(t)\mathrm{d}t - 1} \exp\left[-r_i\left(\chi_i + \int_0^T \gamma_i(t)\mathrm{d}t\right)\right]\,. \end{aligned} \tag{76}$$

which is seen to be a gamma distribution; like the prior in Equation (47), but with updated shape and rate parameters

$$\bar\nu_i = \nu_i + \int_0^T \eta(t)\gamma_i(t)\mathrm{d}t\,, \tag{77}$$

$$\bar{\chi}_i = \chi_i + \int_0^T \gamma_i(t)\mathrm{d}t\,. \tag{78}$$

The posterior mean is therefore

$$\langle r_i \rangle_{q(r)} = \bar{\nu}_i / \bar{\chi}_i\,, \tag{79}$$

which has the same interpretation as the transition-rate posterior: a number of inferred loading events divided by the inferred exposure time in the relevant state, with the prior contributing pseudo-counts and pseudo-exposure. For the other distributions, we also need

$$\langle \log r_i \rangle_{q(r)} = \psi(\bar{\nu}_i) - \log \bar{\chi}_i\,, \tag{80}$$

###### 10.4.3.4 Promoter state posterior $q(s_t)$

The optimality condition (Equation (57)) for $x_i = s_t$ reads

$$q^*(s_t) = \frac{1}{Z}\exp\left(\langle \log p(\tau \mid s_t, r)\rangle_{q(r),q(\tau)}\right)\exp\left(\langle \log p(s_t \mid J, \pi)\rangle_{q(J),q(\pi)}\right)\,. \tag{81}$$

Inserting Equation (37) and Equation (41), we obtain (absorbing non-$s_t$-dependent terms into $Z$)

$$\begin{aligned}
q^*(s_{0:T}) =& \frac{1}{Z}\exp\left(\langle \log \pi_{s_0}\rangle_{q(\pi)} - \int_0^T \langle r_{s_t}\rangle_{q(r)}\mathrm{d}t + \left\langle \sum_\ell \langle \log(r_{s_{\tau_\ell}})\rangle_{q(r)}\right\rangle_{q(\tau)}\right) \\
&\times \exp\left(-\int_0^T \langle J_{s_t}\rangle_{q(J)}\mathrm{d}t + \sum_m \left\langle \log J_{s_{t_m^+} s_{t_m^-}}\right\rangle_{q(J)}\right)\,, \\
=& \frac{1}{Z}\exp\left(\langle \log \pi_{s_0}\rangle_{q(\pi)} - \int_0^T [\langle r_{s_t}\rangle_{q(r)} - \eta(t)\langle \log(r_{s_t})\rangle_{q(r)}]\mathrm{d}t\right) \\
&\times \exp\left(-\int_0^T \langle J_{s_t}\rangle_{q(J)}\mathrm{d}t + \sum_m \left\langle \log J_{s_{t_m^+} s_{t_m^-}}\right\rangle_{q(J)}\right)\,.
\end{aligned} \tag{82}$$

where we defined the posterior average of the Pol II loadings

$$\left\langle \sum_i f(\tau_i)\right\rangle_{q(\tau)} = \int_0^T \eta(t) f(t)\mathrm{d}t\,. \tag{83}$$

If we now introduce the effective initial probability,

$$\tilde{\pi}_{s_0} = \exp\left(\langle \log \pi_{s_0}\rangle_{q(\pi)}\right)\,, \tag{84}$$

The effective generator

$$M_{ij} = \exp\left(\langle \log J_{ij}\rangle_{q(J)}\right)\,, \quad i \neq j\,, \tag{85}$$

with $M_j = \sum_{i\neq j} M_{ij}$, and the tilt

$$f_j(t) = \langle r_j\rangle_{q(r)} - \eta(t)\langle \log r_j\rangle_{q(r)} + \langle J_j\rangle_{q(J)} - M_j\,, \tag{86}$$

we may rewrite Equation (82) as

$$q^*(s_{0:T}) = \frac{1}{Z}\tilde{\pi}_{s_0}\exp\left(-\int_0^T \left(M_{s_t} + f_{s_t}(t)\right)\mathrm{d}t + \sum_m \log M_{s_{t_m^+} s_{t_m^-}}\right)\,. \tag{87}$$

This is in the same form as the CTMC path weight from Equation (41) with rates $M_{ij}$, multiplied by an additional state-dependent dwell-time tilt. The information from the MS2 data enters through this exponential tilt, which steers the process towards states with loading rates that match the current posterior loading rate. The process described by $q^*(s_{0:T})$ is therefore a continuous-time generalization of a Hidden Markov Model (HMM) with the tilt playing the role of the observation likelihood.

The normalization and required state averages follow directly from the CTMC generating functional in Equation (43) with $J \to M$, $\boldsymbol{\pi} \to \tilde{\boldsymbol{\pi}}$, $a = -1$, and $b = 0$. Define the tilted generator

$$G_{ij}(t) = \begin{cases} M_{ij}\,, & i \neq j\,, \\ -M_j - f_j(t)\,, & i = j\,. \end{cases} \tag{88}$$

The forward and backward filters satisfy

$$\begin{aligned}
\dot{\alpha}_i(t) &= G_{ij}(t)\alpha_j(t)\,, \\
-\dot{\beta}_i(t) &= G_{ji}(t)\beta_j(t)\,,
\end{aligned} \tag{89}$$

with initial and final conditions

$$\alpha_i(0) = \tilde{\pi}_i\,, \quad \beta_i(T) = 1\,. \tag{90}$$

The normalization constant is

$$Z = \sum_{i=1}^{K} \alpha_i(T)\,. \tag{91}$$

Splitting the time-ordered exponential at time $t$ gives the state posterior

$$\gamma_i(t) = \langle \delta_{s_t,i} \rangle_{q(s_t)} = \frac{1}{Z}\alpha_i(t)\beta_i(t)\,. \tag{92}$$

For numerical stability and computational speed, we don't explicitly solve the differential equations for all times. Instead, we choose a dense grid of time-points of interest $z_i$ where we are safe to assume an approximately constant value for the tilt, and we then solve the evolution by matrix exponentiation and recursion

$$\alpha_i(z_i) = \left(\exp\left(G(z_i)\Delta z\right)\right)_{ij} \alpha_j(z_{i-1})\,, \tag{93}$$

$$\beta_i(z_i) = \left(\exp\left(G(z_i)^T \Delta z\right)\right)_{ij} \beta_j(z_{i+1})\,. \tag{94}$$

To account for the possibility of underflow in the probabilities, all calculations are done in log space.

We shall also need the transition density for the computation of the expected number of transitions. This will become an input into the transition rate parameter posterior and also serves as a key quantity for studying the alignment with E-P distance and burst onset. The transition density is obtained by differentiating the jump part of the generating functional. In Equation (43), take $J \to M$, $\boldsymbol{\pi} \to \tilde{\boldsymbol{\pi}}$, $a = -1$, and $b = 1$, and choose the jump tilt $g_{kl}(u) = h_{ij}(u)\delta_{ki}\delta_{lj}$ so that only transitions $j \to i$ are counted. This is equivalently generated by

$$G^{(h)}_{kl}(t) = \begin{cases} M_{kl} \exp\left(h_{ij}(t)\delta_{ki}\delta_{lj}\right)\,, & k \neq l\,, \\ -M_l - f_l(t)\,, & k = l\,, \end{cases} \tag{95}$$

and define

$$Z_{ij}[h] = \mathbf{1}^T \mathcal{T} \exp\left(\int_0^T G^{(h)}(u)\mathrm{d}u\right) \tilde{\boldsymbol{\pi}}\,. \tag{96}$$

Differentiating this generating functional inserts the jump-counting observable,

$$\begin{aligned} \gamma_{ij}(t) =& \left.\frac{\delta \log Z_{ij}[h]}{\delta h_{ij}(t)}\right|_{h=0} \\ =& \left\langle \sum_m \delta(t - t_m)\delta_{s_{t_m^+},i}\delta_{s_{t_m^-},j} \right\rangle_{q(s_t)} \\ =& \frac{1}{Z}\alpha_j(t) M_{ij} \beta_i(t)\,. \end{aligned} \tag{97}$$

The expected number of such transitions is therefore $N_{ij} = \int_0^T \gamma_{ij}(t)\mathrm{d}t$, as used in the transition-rate posterior above.

###### 10.4.3.5 Loading event posterior $q(\tau)$

The optimality condition (Equation (57)) for $x_i = \tau$ reads

$$q^*(\tau) = \frac{1}{Z} p(I^{\mathrm{obs}}_{0:T} \mid \tau_{0:T}) \exp\left(\langle \log p(\tau \mid s_t, r) \rangle_{q(r),q(s_t)}\right)\,. \tag{98}$$

The average can be done straight-forwardly using $\gamma_i(t)$ and we get

$$q^*(\tau) = \frac{1}{Z} p(I^{\mathrm{obs}}_{0:T} \mid \tau_{0:T}) \exp\left(-\int_0^T [\bar{r}_t - \lambda]\mathrm{d}t + \sum_i \log\frac{\bar{r}_{\tau_i}}{\lambda}\right)\,, \tag{99}$$

where we defined the effective rate

$$\bar{r}_t = \exp\left(\sum_i \gamma_i(t) \langle \log r_i \rangle_{q(r)}\right)\,. \tag{100}$$

The distribution in Equation (99) is an inhomogeneous point process weighted by the MS2 likelihood. Such a process is sometimes referred to as a Gibbs or Gibbsian point process [94]. In statistical mechanics, it may be identified with the grand canonical ensemble of a 1D gas of interacting particles with Hamiltonian ($\mathrm{k_B}T = 1$)

$$-H(\tau) = \log p(I^{\mathrm{obs}}_{0:T} \mid \tau_{0:T}) + \sum_i \log(\bar{r}_{\tau_i})\,. \tag{101}$$

The intuition here is that the MS2 data steers the process towards likely values, while $\bar{r}_t$ selects loading event configurations which match the posterior belief of the promoter state and loading rates. For the Gaussian likelihood we employ, the Hamiltonian can be seen to have the pairwise interaction form

$$\begin{aligned} -H(\tau) &= -\sum_k \frac{1}{2\sigma_I^2}\left(I_k^{\text{obs}} - I_{\text{offset}} - \sum_i \phi(t_k - \tau_i)\right)^2 + \sum_i \log \bar{r}_{\tau_i} + \text{constants}\,, \\ &= \sum_i h(\tau_i) + \sum_{i>j} V(\tau_i, \tau_j) + \text{constants}\,, \end{aligned} \tag{102}$$

where we defined the external field $h$ and pair potential $V$

$$h(\tau_i) = \log \bar{r}_{\tau_i} + \sum_k \frac{\left(I_k^{\text{obs}} - I_{\text{offset}}\right)\phi(t_k - \tau_i)}{\sigma_I^2} - \sum_k \frac{\phi(t_k - \tau_i)^2}{2\sigma_I^2}\,, \tag{103}$$

$$V(\tau_i, \tau_j) = -\sum_k \frac{\phi(t_k - \tau_i)\phi(t_k - \tau_j)}{\sigma_I^2}\,, i \neq j\,. \tag{104}$$

Note that the MS2 data only enters through the local external field, while the pair potential just depends on the kernel $\phi$. Writing things in terms of the potentials, we have

$$q^*(\tau) = \frac{1}{Z}\exp\left(\sum_i h(\tau_i) + \sum_{i>j} V(\tau_i, \tau_j)\right)\,. \tag{105}$$

The average of interest is the posterior loading rate density

$$\eta(t) = \left\langle \sum_i \delta(t - \tau_i) \right\rangle_{q(\tau)}\,. \tag{106}$$

There are few analytical results available for such interacting systems, and we therefore resort to sampling to evaluate the needed average. To sample from Equation (105), we employ an optimized version of the methodology described by Møller [95]. The idea is to simulate the loading events as a birth-death process in such a way that the stationary distribution becomes a sample from $q^*(\tau)$. The algorithm relies on the Papangelou conditional intensity

$$\lambda^*(\tau, \xi) = \exp\left(h(\tau) + \sum_i V(\tau, \xi_i)\right)\,, \tag{107}$$

which gives the conditional rate of a new loading event at $\tau$ given the current loading events $\xi$. Given a means to evaluate $\lambda^*$, the algorithm proceeds as follows: With probability $1/2$, either generate a birth event $\tau$ from the rate function $\bar{r}_t / \int_0^T \bar{r}_{t'} \mathrm{d}t'$ or perform a death event by choosing an event $\tau \in \xi$ at random from the $n$ current events in $\xi$ (if $n = 0$ do nothing). Now define the 'Metropolis-Hastings' ratio

$$r(\xi, \tau) = \lambda^*(\xi, \tau)\frac{\int_0^T \bar{r}_t \mathrm{d}t}{(n+1)\bar{r}_\tau}\,. \tag{108}$$

A birth transition is accepted with probability $\min(1, r(x, \tau))$ and a death transition is accepted with probability $\min(1, 1/r(\xi \backslash \tau, \tau))$ where $\xi \backslash \tau$ refers to the set of current events with $\tau$ removed. This ensures that the birth-death Markov chain will converge to a sample from $q^*(\tau)$ once it reaches steady state.

To efficiently simulate the birth-death process, sampling was implemented in `jax` to allow batched GPU computation across many trajectories. We restricted the proposal to the same grid of points $z_i$ used for the forward-backward algorithm to allow for fast sample generation from $\bar{r}_t$ by numerically computing and inverting its CDF on the grid. The finite range of the interaction potential and the outer-product form of the potential allowed for further optimization of the calculation of the Papangelou intensity. The summands over the observation times appearing in Equation (103) are only non-zero over the support of the kernel. Not summing over terms outside the support yielded a significant speedup. Furthermore, since only one particle is added at a time, and only the interaction of one point with the current set is needed to compute $\sum_i V(\tau, \xi_i)$, we can store a vector of $\phi(t_k - \xi_i)/\sigma_I^2$ for the current set of points and evaluation of $\sum_i V(\tau, \xi_i)$ then amounts to a simple dot product over the times $t_k$ where $\phi(t_k - \tau)$ is non-zero.

##### 10.5 Calibration of kernel parameters by fitting the MS2 autocorrelation

To keep the variational scheme identifiable and tractable, we limited the parameters to the promoter state transition matrix, Pol II loading rates, and promoter state initial probability. This left out the parameters of the observation model, which converts Pol II loadings into signal, as well as possible trajectory-to-trajectory variability in the loading rates. Here we describe how we obtain them by fitting the MS2 mean and autocorrelation function to the exact expression predicted by our model. This provided us with good values for the kernel parameters and the MS2 intensity to be used in the variational inference scheme.

To perform the fit, we need the model prediction for the mean and autocorrelation $\langle I^{\mathrm{obs}}(t)I^{\mathrm{obs}}(t')\rangle - \langle I^{\mathrm{obs}}\rangle^2$ of the observed MS2 signal. The mean is straightforward and may immediately be obtained by insertion of the signal model from Pol II loadings from Eq. (34) to get

$$\begin{aligned}\langle I^{\mathrm{obs}}(t)\rangle &= I_{\mathrm{offset}} + \langle I(t)\rangle\,,\\ &= I_{\mathrm{offset}} + \int_{-\infty}^{\infty}\phi(t-t')\langle r_{s_{t'}}\rangle \mathrm{d}t'\,,\\ &= I_{\mathrm{offset}} + \langle r\rangle\int_{-\infty}^{\infty}\phi(x)\mathrm{d}x\,,\\ &\equiv I_{\mathrm{offset}} + \langle r\rangle I_0\,,\end{aligned} \tag{109}$$

where we defined

$$\langle r\rangle = \sum_i p_{\mathrm{ss},i} r_i\,,\quad I_0 = \int_{-\infty}^{\infty}\phi(x)\mathrm{d}x\,, \tag{110}$$

and used the fact that the promoter state average is independent of time and $\phi(t)$ is translationally invariant. $\boldsymbol{p}_{\mathrm{ss}}$ is the steady-state of the promoter model, defined as the normalized eigenstate of the transition rate matrix with eigenvalue $0$

$$\sum_j (J_{ij} - \delta_{ij}J_j)\,p_{\mathrm{ss},j} = 0\,,\quad \sum_i p_{\mathrm{ss},i} = 1 \tag{111}$$

The non-centered correlation function follows a similar line of reasoning. The constant offset will cancel from the centered autocorrelation and therefore only contribute to the mean term above. For the dynamic part of the signal,

$$\begin{aligned}\langle I(t)I(0)\rangle &= \left\langle \sum_{ij}\phi(t-\tau_i)\phi(-\tau_j)\right\rangle\,,\\ &= \int_{-\infty}^{\infty}\int_{-\infty}^{\infty}\phi(t-x)\phi(-y)\langle r_{s_x}r_{s_y}\rangle \mathrm{d}x\mathrm{d}y + \langle r\rangle\int_{-\infty}^{\infty}\phi(t-t')\phi(-t')\mathrm{d}t'\,,\\ &= \int_{-\infty}^{\infty}\int_{-\infty}^{\infty}\phi(x)\phi(y)\langle r_{s_{t-x}}r_{s_{-y}}\rangle \mathrm{d}x\mathrm{d}y + \langle r\rangle\int_{-\infty}^{\infty}\phi(x)\phi(x-t)\mathrm{d}x\,,\\ &= \int_{-\infty}^{\infty}\int_{-\infty}^{\infty}\phi(x)\phi(y)\Lambda(t+y-x)\mathrm{d}x\mathrm{d}y + \langle r\rangle\int_{-\infty}^{\infty}\phi(x)\phi(x-t)\mathrm{d}x\,,\\ &= I_1(t) + I_2(t)\,,\end{aligned} \tag{112}$$

where we defined the loading rate correlation function

$$\Lambda(t) = \langle r_{s_t}r_{s_0}\rangle = \sum_k\sum_j r_k r_j\langle \delta_{s_t,k}\delta_{s_0,j}\rangle\,, \tag{113}$$

and the autocorrelation integrals

$$I_1(t) = \int_{-\infty}^{\infty}\int_{-\infty}^{\infty}\phi(x)\phi(y)\Lambda(t+y-x)\mathrm{d}x\mathrm{d}y\,, \tag{114}$$

$$I_2(t) = \langle r\rangle\int_{-\infty}^{\infty}\phi(x)\phi(t+x)\mathrm{d}x\,. \tag{115}$$

The centered autocorrelation used for fitting is then

$$\mathrm{Cov}\Big[I^{\mathrm{obs}}(t), I^{\mathrm{obs}}(0)\Big] = I_1(t) + I_2(t) + \sigma_I^2\delta(t) - \langle r\rangle^2 I_0^2\,. \tag{116}$$

This expression makes the role of the offset explicit: because the covariance is centered by subtracting $\langle I^{\mathrm{obs}}\rangle^2$, $I_{\mathrm{offset}}$ changes the mean constraint but not the autocorrelation at non-zero lag.

To account for trajectory-to-trajectory variability in loading rates, we let the loading rate in state $i$ be a random variable $\rho_i$ with mean $r_i$ and coefficient of variation $\mathrm{CV}_r$,

$$\langle \rho_i \rangle = r_i \,, \quad \mathrm{Cov}(\rho_i, \rho_j) = \delta_{ij} \left( \mathrm{CV}_r r_i \right)^2 \,. \tag{117}$$

We assume this variability is independent of the promoter state trajectory and constant within each trajectory. The loading-rate second moment entering $\Lambda(t)$ is therefore

$$L_{ij} \equiv \langle \rho_i \rho_j \rangle = r_i r_j + \delta_{ij} \left( \mathrm{CV}_r r_i \right)^2 \,, \tag{118}$$

and loading-rate variability is incorporated by replacing

$$\Lambda(t) \to \Lambda_{\mathrm{CV}}(t) = \sum_k \sum_j L_{kj} \langle \delta_{s_t,k} \delta_{s_0,j} \rangle \,. \tag{119}$$

This is the form used in the fit below; when $\mathrm{CV}_r = 0$, it reduces to the expression with fixed loading rates. For the rise and plateau form for $\phi$ we employ in this study, these integrals can be simplified by analytical evaluation of some of the integrals. Define

$$c(t) = \int_{-\infty}^{\infty} \phi(x)\phi(t+x)\mathrm{d}x \,, \tag{120}$$

in terms of which

$$I_1(t) = \int_{-\infty}^{\infty} c(x)\Lambda_{\mathrm{CV}}(t+x)\mathrm{d}x \,, \tag{121}$$

$$I_2(t) = \langle r \rangle c(t) \,. \tag{122}$$

$c(t)$ may be evaluated analytically and comes out to be

$$\begin{aligned} c(t) = & I_\phi^2 \Bigg[ (T_{\mathrm{Plateau}} - \mid t \mid)_+ \\ & + \frac{1}{2T_{\mathrm{Rise}}} \left[ C(T_{\mathrm{Rise}} + T_{\mathrm{Plateau}} - t)^2 - C(T_{\mathrm{Rise}} - t)^2 + C(T_{\mathrm{Rise}} + T_{\mathrm{Plateau}} + t)^2 - C(T_{\mathrm{Rise}} + t)^2 \right] \\ & + \frac{1}{T_{\mathrm{Rise}}^2} \left\{ \frac{t}{2} \left[ C(T_{\mathrm{Rise}} - t)^2 - C(-t)^2 \right] + \frac{1}{3} \left[ C(T_{\mathrm{Rise}} - t)^3 - C(-t)^3 \right] \right\} \Bigg] \,, \end{aligned} \tag{123}$$

where $C(x) = \min(\max(x, 0), T_{\mathrm{Rise}})$ is the clamp function on $[0, T_{\mathrm{Rise}}]$ and $(x)_+$ is RELU function which is zero for $x < 0$ and $x$ otherwise. The only remaining integral is the one in $I_1$, which requires the autocorrelation of a CTMC $\langle \delta_{s_t,k} \delta_{s_0,j} \rangle$. This result follows straightforwardly from the propagator

$$p(s_t = i, s_0 = j) = \left( \mathrm{e}^{At} \right)_{ij} p_{\mathrm{ss},j} \,, \quad A_{ij} = J_{ij} - \delta_{ij} J_j \,, \tag{124}$$

yielding

$$\langle \delta_{s_t,i} \delta_{s_0,j} \rangle = \begin{cases} \left( \mathrm{e}^{At} \right)_{ij} p_{\mathrm{ss},j} & t \geq 0 \\ p_{\mathrm{ss},j} \left( \mathrm{e}^{A^T(-t)} \right)_{ji} & \text{otherwise} \end{cases} \,. \tag{125}$$

From this result, we have all we need to construct the fit function if we perform the last integral in $I_1(t)$ numerically.

To fit the curves to data, we estimated the covariance of the sample mean and autocorrelation by bootstrapping to get a likelihood which we maximized with respect to $J_{ij}$, $r_i$, $\sigma_I^2$, $T_{\mathrm{Rise}}$, $T_{\mathrm{Plateau}}$, $I_\phi$, $I_{\mathrm{offset}}$, and $\mathrm{CV}_r$ using the `L-BFGS-B` method in `scipy.optimize.minimize` using `jax` to obtain Jacobians. The fit was done jointly between 1.5noC, 339CECP, 87noC, 170noC, 339CE, 253noC, 339CP, 339noC sharing all parameters but $k_{\mathrm{on}}^{\mathrm{eff}}$ and $\sigma_I^2$. The results of the calibration can be found in (**Fig. S21**, **Table S9**). We note that while we fit $\sigma_I^2$, this was primarily done to allow the fit to not focus too much on the zero-lag part of the covariance function. When running the inference, we estimated per-track noises as the mean absolute value of single-frame changes in the MS2 trajectory.

#### 10.6 Calibration of $k_{\mathrm{on}}$ and $R_{\mathrm{E-P}}$ using an exact transfer-matrix evidence for a discretized effective-rate model

The variational inference scheme described above provides promoter-state and loading-rate estimates for each condition, while the MS2 autocorrelation fit fixes the observation parameters $T_{\mathrm{Rise}}$, $T_{\mathrm{Plateau}}$, $I_\phi$, $I_{\mathrm{offset}}$, and $\sigma_I^2$. However, it does not allow us to learn the parameters connecting E-P distance and transcription directly. To ensure we obtain a robust likelihood for $k_{\mathrm{on}}$ and $R_{\mathrm{E-P}}$, we employed an exact finite-state transfer-matrix evidence for the observed MS2 traces, conditional on the GPR posterior over E-P separation trajectories. The exactness comes at the cost that we must restrict Pol II loadings to be binary events at the observation times, meaning that we can only process the 30 s data due to state-spaces becoming too large for 5 s data, and this method would not work for genes with very high Pol II loading rates. The approach is very similar to the compound-state hidden Markov model (cpHMM) introduced by Lammers et al. [88] but differs in an important way. In cpHMM, one only keeps track of the promoter state, and to compute MS2 it is assumed that Pol II has no randomness such that MS2 is a deterministic function of the promoter state history. This is only a good approximation when transcription levels are high, and can fail when only a few Pol II are present in the window. For our data, we find ourselves in this regime and therefore had to adapt it to account for the stochastic nature of Pol II dynamics.

For each trajectory $a$ and observation interval ending at time $t_n$, we first compute the posterior contact probability

$$p_{c,a,n}(R_{\mathrm{E-P}}) = \langle \theta(R_{\mathrm{E-P}} - \|\boldsymbol{R}_a(t_n)\|) \rangle_{\boldsymbol{R}_a(t_n)\sim p(\boldsymbol{R}_a(t_n)|\boldsymbol{R}^{\mathrm{obs}}_{a,0:T})} \,. \tag{126}$$

In practice, this average is evaluated by numerical quadrature over the Gaussian posterior for $\boldsymbol{R}_a(t_n)$. The contact probability enters the two-state promoter transition over one observation interval through (neglecting multiple transitions in an interval)

$$u_{a,n} = 1 - \mathrm{e}^{-k_{\mathrm{on}} p_{c,a,n}(R_{\mathrm{E-P}})\Delta t}\,, \quad v = 1 - \mathrm{e}^{-k_{\mathrm{off}}\Delta t}\,, \tag{127}$$

where $u_{a,n}$ is the probability of an OFF-to-ON transition in the interval and $v$ is the probability of an ON-to-OFF transition. With columns indexing the previous state and rows indexing the next state, the promoter transition matrix is

$$P_{a,n} = \begin{pmatrix} 1-u_{a,n} & v \\ u_{a,n} & 1-v \end{pmatrix} . \tag{128}$$

The MS2 signal has a finite memory because a Pol II loading event contributes signal only over the support of $\phi$. We therefore augment the promoter state with a binary loading-history vector $\mathbf{b}_n \in \{0,1\}^m$, where

$$m = \left\lfloor \frac{T_{\mathrm{Rise}} + T_{\mathrm{Plateau}}}{\Delta t} \right\rfloor + 1\,. \tag{129}$$

The element $b_{n,\ell}$ indicates whether a loading event occurred $\ell$ frames before the observation at $t_n$. Conditional on promoter state $i$, the probability of a loading event in the next time step is

$$q_i = 1 - \mathrm{e}^{-r_i \Delta t}\,. \tag{130}$$

The loading-history update matrix $B_i(\mathbf{b}' \mid \mathbf{b})$ shifts the history by one frame and appends a new binary loading event. Equivalently, if $\ell_n \in \{0,1\}$ denotes the newly appended loading event,

$$B_i(\mathbf{b}' \mid \mathbf{b}) = q_i^{\ell_n}(1-q_i)^{1-\ell_n} \tag{131}$$

when $\mathbf{b}'$ is obtained from $\mathbf{b}$ by dropping the oldest entry and appending $\ell_n$, and $B_i(\mathbf{b}' \mid \mathbf{b}) = 0$ otherwise.

For a loading-history state $\mathbf{b}$, the expected MS2 signal is the discrete convolution with the MS2 kernel,

$$\mu(\mathbf{b}) = I_{\mathrm{offset}} + \sum_{\ell=0}^{m-1} w_\ell b_\ell\,, \tag{132}$$

where $w_\ell$ is the discretized rise-and-plateau kernel with scale $I_\phi$. If the offset has already been subtracted from the fitted MS2 trace, this is equivalent to setting $I_{\mathrm{offset}} = 0$ in the emission model. For an observed frame, the Gaussian emission likelihood is

$$G_{a,n}(\mathbf{b}) = \frac{1}{\sqrt{2\pi\sigma_{I,a}^2}} \exp\left( -\frac{\left(I^{\mathrm{obs}}_{a,n} - \mu(\mathbf{b})\right)^2}{2\sigma_{I,a}^2} \right) . \tag{133}$$

For missing or padded observations, we set $G_{a,n}(\mathbf{b}) = 1$ for all $\mathbf{b}$, so those frames propagate the hidden state without contributing to the evidence. This padding allows Pol II loading events before the first measured MS2 frame to contribute to later signal through the finite MS2 memory.

Let $\alpha_{a,n}(i,\mathbf{b})$ be the normalized forward probability over the joint promoter and loading-history state after observing frame $n$. We initialize the recursion with the promoter off and an empty loading history, $\alpha_{a,0}(0,\mathbf{0}) = 1$, followed by the emission likelihood for the first frame. For each subsequent frame,

$$\tilde{\alpha}_{a,n}(i,\mathbf{b}') = \sum_{j,\mathbf{b}} B_i(\mathbf{b}' \mid \mathbf{b}) P_{a,n}(i \mid j) \alpha_{a,n-1}(j,\mathbf{b})\,, \tag{134}$$

$$Z_{a,n} = \sum_{i,\mathbf{b}} G_{a,n}(\mathbf{b}) \tilde{\alpha}_{a,n}(i,\mathbf{b})\,, \tag{135}$$

$$\alpha_{a,n}(i,\mathbf{b}) = \frac{G_{a,n}(\mathbf{b})\tilde{\alpha}_{a,n}(i,\mathbf{b})}{Z_{a,n}}\,. \tag{136}$$

The normalization constants $Z_{a,n}$ are accumulated to obtain the exact evidence for the discretized hidden-state model,

$$\log \mathcal{Z}_a(k_{\rm on}, R_{\rm E-P}) = \sum_n \log Z_{a,n} \,. \tag{137}$$

This recursion is exact because it sums over all promoter paths and all binary loading histories compatible with the finite MS2 memory, rather than sampling or assigning a single burst history.

Assuming independent trajectories and conditions, the joint evidence is

$$\log \mathcal{Z}(k_{\rm on}, R_{\rm E-P}) = \sum_a \log \mathcal{Z}_a(k_{\rm on}, R_{\rm E-P}) \,. \tag{138}$$

We evaluate this quantity on a grid of $k_{\rm on}$ and $R_{\rm E-P}$. When reporting evidence for $R_{\rm E-P}$ alone, we marginalize over $k_{\rm on}$ by numerical quadrature,

$$\log p(R_{\rm E-P} \mid \mathcal{D}) = \text{const.} + \log \sum_q \exp\left(\log \mathcal{Z}(k_{{\rm on},q}, R_{\rm E-P})\right) \Delta k_{{\rm on},q} \,, \tag{139}$$

corresponding to a flat prior over the range of $k_{\rm on}$ included in the grid.

###### 10.6.1 VEPI implementation

We implemented coordinate ascent variational inference (CAVI) to optimize the posteriors [96]. We chose initializations to force the model into a basin of promoter state largely driven by MS2, (high loading rate and low transition rate) since we found that random initializations led to convergence to a suboptimal homogeneous state. Following initialization, we cycled through a schedule of updates of the posteriors where, for each variable, we updated its variational distribution using the averages of the others. Since each update maximizes the ELBO with respect to the current averages, each step increases the ELBO. The iteration was stopped when the relative change in the transition rate averages between iterations reached 0.5%.

Before fitting, the data was subtracted by the offset inferred from the MS2 autocorrelation fits. This helped ensure that any residual in MS2 wasn't being interpreted as Pol II loading, possibly leading to under-estimated loading rates or even bursting. For a similar reason, we set a rather strong prior on the ON-state loading rate to the inferred value from the MS2 autocorrelation fits. This was done as initial runs showed the loading rate to drift low for tracks without bursts, leading to an erroneously inferred long ON-period when there really was a burst in the track.

##### 10.7 Validation

We first validated the MS2 inference without an E-P drive. For this, we generated MS2 simulated data from the true model and assessed the ability of the inference to recover the hidden states and parameters when given correct values for the MS2 kernel.

We explored the following grid of conditions:

- Signal-to-Noise Ratio (SNR) $I_\phi/\sigma_\phi$ : `0.5 , 0.875, 1.25 , 1.625, 2`
- $k_{\rm on}^{\rm eff}$: `1/15 min. , 1/67.5 min., 1/120 min. , 1/172.5 min., 1/225 min.`
- $k_{\rm off}$: `1/ min. , 1/4.5 min., 1/8 min. , 1/11.5 min., 1/15 min.`
- Time step: `0.1 min., 0.5 min.`
- Number of traces: `50, 200`
- Track maximum duration: `500 min.`

When simulating, we also dropped 10% of measurements to emulate the missing frames in our experimental data and shortened the trajectory by a random amount up to half the trajectory length. Note that this is a different ON-rate than we have referred to before, as this model does not contain an E-P distance. In a mean-field sense, they differ roughly by the contact probability

$$k_{\rm on}^{\rm eff} \approx k_{\rm on} P(\text{contact}) \,, \tag{140}$$

but this need not always be the case. We assessed the ability of the model to reconstruct promoter state and Pol II loadings as well as loading rate and transition rates. **Fig. S18A**-**C** shows the posterior metrics used for validation on an example trajectory. For visual examples of the type of MS2 trajectories in these different conditions, see **Fig. S19** for a set of trajectories drawn randomly.

We found that MS2 was always well reconstructed with little bias (**Fig. S18D**). This was further elucidated when looking at the error in estimation of the number of Pol II loaded at a given time point (**Fig. S18E**). This is a stringent metric as any error early in the trajectory will give you an offset throughout the trajectory. Despite this, VEPI consistently estimated Pol II counts within less than one polymerase on average, with a few difficult simulation conditions yielding an average over-estimation of two polymerases.

Promoter states were also well-reconstructed, with most points being correctly predicted (**Fig. S18F**). We found that, for most simulated conditions, we could predict the ON-state with an AUC of 0.9 or higher from thresholding $P(ON)$ (**Fig. S18G**). The True Positive Rate (TPR) and False Positive Rate (FPR) for each condition were consistently above 0.8 and below 0.2, respectively (**Fig. S18H**). Furthermore, we found that $P(ON)$ never steered much further than 10% away from the true value on average (**Fig. S18I**).

Transition rates got close to their true values, but were generally slightly under-estimated and were around 70%-90% of their true value independently of SNR (**Fig. S18J**, **K**). The main determinant of this under-estimation seemed to be bursts with few loadings, as the simulations with only a few loadings per burst formed a tail with an under-estimated rate of as low as 50% of the true value. Loading rates were similarly under-estimated but generally fell closer to the true value with a tighter scatter of at most 80% of bursts below the true value. The few-burst outliers fell much lower, and we suspect this is due to many bursts not having any loadings, biasing the estimate down (**Fig. S18L**).

Having validated MS2 inference, we then assessed whether the VEPI could accurately infer the parameters connecting E-P distance and MS2. For this, we simulated Rouse polymer trajectories and drove a promoter with $R_{\mathrm{E-P}}$ of 30 nm. The polymer parameters were adapted from [8], and promoter parameters were chosen in a noisy regime with transition rates comparable to those ultimately found in experiments.

- $k_{\mathrm{off}}$: `1/60 min.`
- $r_{\mathrm{loading}}$: `0.5 min.`$^{-1}$
- $T_{\mathrm{Rise}}$: `2.0 min.`
- $T_{\mathrm{Plateau}}$: `4.0 min.`
- SNR: `0.5`
- Track duration: `5 hours`
- Number of trajectories: `1000`
- Bond spring constant: `1/0.177 s`
- Bead diffusion coefficient: `2810 nm`$^2$`/s`
- Polymer length: `334`
- Enhancer position: `122`
- Promoter position: `212`

We then evaluated our transfer-matrix solution for the likelihood of $k_{\mathrm{on}}$ and $R_{\mathrm{E-P}}$ and found that $R_{\mathrm{E-P}}$ is overestimated with the current model (**Fig. S20**). The true pair of $k_{\mathrm{on}}$ and $R_{\mathrm{E-P}}$ was on the line of the most likely $k_{\mathrm{on}}$ for a given $R_{\mathrm{E-P}}$, but the combination was not the most likely. In data, we similarly saw that the peak in the likelihood landscape was for very large $R_{\mathrm{E-P}}$, which prompted us to only use the ridge of the most likely $k_{\mathrm{on}}$ for a given $R_{\mathrm{E-P}}$ in the main text. We suspect that the over-estimation is due to the time-scale separation approximation employed when averaging over the E-P posterior for the tractability of the inference scheme.

##### 10.8 Results

The application of VEPI on the data proceeded as follows:

1. Fit the MSD to obtain the parameters for the GP.
2. Fit the MS2 autocorrelation to obtain the MS2 kernel parameters.
3. Run the likelihood computation to get the $k_{\mathrm{on}}$ which matches our estimates of $R_{\mathrm{E-P}}$ (already described in main text).
4. Run VEPI without an E-P drive to get per-trajectory loading rates, initial probabilities, $r_{\mathrm{loading}}$, and $k_{\mathrm{off}}$
5. Run VEPI with E-P drive to infer full E-P driven MS2 dynamics.

MSD-fitting was described earlier, and the result of the MS2 autocorrelation fits can be found in **Table S9**. The fit results were found to be in surprisingly good agreement with the data, given the simplicity of the model employed. The mean tracked well across conditions (**Fig. S21B**), and we found satisfactory agreement between the measured and fitted correlation functions (**Fig. S21C**). There were deviations in individual conditions, but given the small number of fittable parameters and the fact that most were shared globally, fits were deemed appropriate for calibration.

Given calibrated values, we ran VEPI without E-P drive to obtain per-trajectory calibration parameters and to study the bursting behavior and transcription of the system. The loading rates were consistently scattered around a mean of 0.4 $min.^{-1}$ and the value of 0.7 $min.^{-1}$ from the autocorrelation fits was well within the spread of per-track fitted loading rates (**Fig. S22A**). The average per-track loading rate (removing the ones close to the prior) was well in agreement with the one from the autocorrelation fits. There was a slight trend of higher loading rates being fitted for the two highly transcribing conditions: 1.5noC and 339CECP. We do not know if this is real or an artifact of the simple two-state model employed, not having the dynamic range to capture the full spectrum of transcriptional activity in the data.

Moving on to burst-calls, we found the average probability of bursting to correlate well with the average MS2 (**Table S9**, **Fig. S23A**). Generally, tracks displayed quite long bursts which often lasted a significant fraction of the entire trajectory (**Fig. S23B**). This is in agreement with the inferred ON-times, which were generally in the hour range regardless of condition (**Table S9**, **Fig. S22C**). These are long durations compared to previous findings from MS2 imaging in mESCs [97, 98]. There may be two main reasons for this discrepancy. First, previous analysis methods did not have an explicit model for Pol II loading, and therefore might be more likely to break up bursts when seeing small dips in MS2, which may be explainable with fluctuations in loading rather than promoter state switching. Second, our system may display more RNA retention at the gene or delayed excision of the transcript, leading to extended periods of MS2 per Pol II loading, effectively bridging bursts, making the burst appear longer. Regardless of the mechanism, this does not limit our ability to infer properties of E-P activation, as these happen at burst onset, and while missing some burst onsets might make the dataset smaller, it won't inhibit our ability to study the E-P proximity and dynamics around the onsets that we do resolve.

Finally, similarly to the simulated data, we find that the MS2 is predicted with very little bias, suggesting that the inference of Pol II is properly deconvolving the trajectory (**Fig. S22D**). This gave us confidence that the inference made for a good baseline onto which to introduce the E-P drive. The results of that analysis are presented in the main text. We include here a final analysis on the consistency of the inferred model for the E-P-driven promoter. By simulating from the prior model, one can check the agreement with the fitted model parameters. We do not expect perfect agreement in such an analysis, as there is a great deal of heterogeneity, possibility of multiple promoter states, loop extrusion, E-P attraction, etc. in the data which we do not explicitly model in our simple model. Despite this, we found that both the mean (**Fig. S24A**) and most of the MS2 intensity histograms (**Fig. S24B**-**H**) were largely in agreement with predictions. The main difference seemed to be the appearance of a right low-probability tail in the data that our model was not able to recapitulate. This might also explain the somewhat larger dynamic range seen in the mean of the real data compared to our predictions.

### 11 Supplementary Note 1: experimental and computational considerations for uncovering E-P proximity-transcription correlations

#### 11.1 Introduction

Prior imaging studies attempting to understand Enhancer-Promoter (E-P) interactions have reported either a lack of correlation between E-P 3D proximity and transcription, or even an anti-correlation between E-P 3D proximity and transcription [84, 99, 100] as further discussed in [49, 101–104]. This observation is seemingly at odds with studies using genomic approaches (such as Hi-C and Micro-C) that consistently demonstrate a strong positive correlation between physical E-P "contacts" and transcriptional output [105–108], including this study. Below we show that this seeming discrepancy is likely technical rather than biological and due to noise and biases in imaging data.

Here, we study and discuss how a correlation between E-P 3D proximity and transcription in live cells might be missed due to experimental and analytical limitations, and we examine which experimental and analytical aspects are key to observing such a correlation if it exists. Aspects of this have also been discussed previously [49, 103, 109, 110]. In the following, we will assume and simulate a "hit-and-run dynamic contact model" for functional E-P interactions, where the enhancer transiently contacts the promoter and activates it when their 3D distance is below 40 nm. We will then show how this causal relationship between E-P contact and transcription can easily be missed due to several experimental and analytic limitations including the label-locus separation of fluorescent labels, localization uncertainty, delay between E-P interactions and transcription signal, and the temporal smearing introduced by burst-calling algorithms. In this note, we apply 3D polymer simulations and stochastic transcription modeling to systematically demonstrate how these factors can obscure the true E-P distance "dip" at the onset of transcription bursts, and review the different aspects one must consider to resolve this conflict.

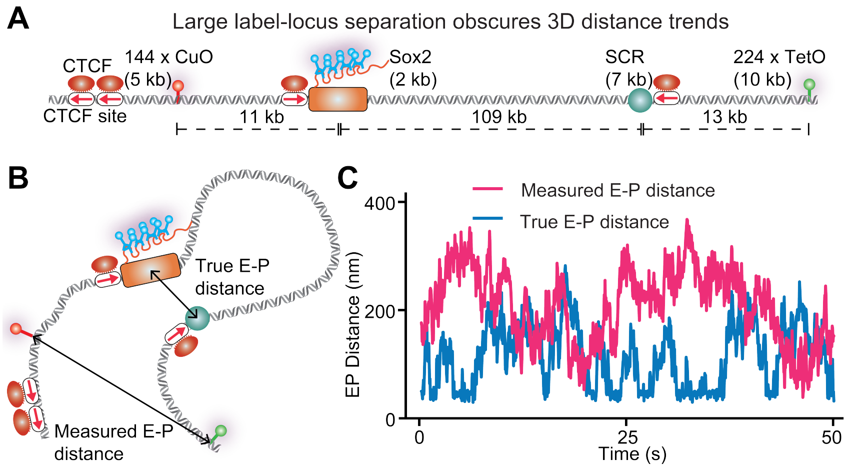

**Supplementary Note 1 Fig. 1: Comparison of true E-P distance vs. measured E-P distance** (**A**) Illustration of the E-P pair of interest, fluorescent labels and their genomic separations used by Alexander *et al.* [84]. (**B**) Label-locus separation leads to large deviation of measured E-P distance from true E-P distance. (**C**) Example trajectories of measured E-P distance versus true E-P distance for the E-P configuration in (A), from polymer simulations.

#### 11.2 The reliability of a fluorescent label to serve as a positional reporter depends strongly on the label-locus separation

To read out E-P 3D distance, we need to know the precise 3D position of both the enhancer and promoter. This currently is not possible via live-cell imaging, and the field instead uses positional proxies by inserting fluorescently labeled arrays near the enhancer or promoter of interest (e.g., LacO, TetO, CuO, or MS2 cassettes). The ability of the fluorescent label to serve as a reliable reporter depends strongly on its size and distance from the enhancer or promoter of interest. If we seek to know the 3D position of the center of an enhancer and we assume we measure the center of the fluorescent label, the separation is half the length of the fluorescent label plus the separation between the fluorescent label and the enhancer, plus half the length of the enhancer (**Supplementary Note 1 Fig. 1A**).

Alexander *et al.* [84] achieved a major breakthrough and reported the first simultaneous live-cell imaging of E-P interactions and nascent transcription in mammalian cells using a similar approach to that of Chen *et al.* [111], who reported such imaging of a synthetic E-P system in *Drosophila* the year prior. Alexander *et al.* studied the *Sox2* E-P loop using a fluorescent labeling approach shown in **Supplementary Note 1 Fig. 1A**. Alexander *et al.* observed no correlation between E-P proximity and transcription and concluded that their data "*supports an unexpected* *mechanism for enhancer control of Sox2 expression that uncouples transcription from enhancer proximity*". In this note, we will use the groundbreaking study of Alexander *et al.* as an illustration to investigate whether direct contact can be ruled out. Indeed, this was also discussed by Alexander *et al.* "*It is important to note that we cannot exclude the importance of direct Sox2/SCR contacts in Sox2 activation*".

In the *Sox2* study, the label separation was 11 kb and 13 kb (**Supplementary Note 1 Fig. 1A**), respectively, which can strongly reduce the reliability of the inserted fluorescent label to serve as a positional reporter (**Supplementary Note 1 Fig.1B,C**, see also [49, 103] where this is discussed).

##### 11.2.1 Mathematical derivation of measured distance

Let $\mathbf{r}_E(t)$ and $\mathbf{r}_P(t)$ denote the time-dependent position vectors of the functional enhancer and promoter. The biologically relevant variable driving the burst is the functional Euclidean distance:

$$d_{\text{func}}(t) = \|\mathbf{r}_E(t) - \mathbf{r}_P(t)\| \tag{141}$$

In our simulations, the promoter transitions to the ON state only when $d_{\text{func}}(t) < R_{\text{contact}}$ (set to ~40 nm). However, the fluorescent labels introduce spatial offset vectors $\boldsymbol{\delta}_E$ and $\boldsymbol{\delta}_P$ relative to the functional sites. These offsets depend on the genomic distance (in base pairs) between the label insertion site and the functional element. The measured positions are:

$$\mathbf{R}_E(t) = \mathbf{r}_E(t) + \boldsymbol{\delta}_E(t) \tag{142}$$

$$\mathbf{R}_P(t) = \mathbf{r}_P(t) + \boldsymbol{\delta}_P(t) \tag{143}$$

The measured distance $d_{\text{meas}}(t)$ is:

$$d_{\text{meas}}(t) = \|\mathbf{R}_E(t) - \mathbf{R}_P(t)\| = \|(\mathbf{r}_E(t) - \mathbf{r}_P(t)) + (\boldsymbol{\delta}_E(t) - \boldsymbol{\delta}_P(t))\| \tag{144}$$

Let $\boldsymbol{\Delta}(t) = \boldsymbol{\delta}_E(t) - \boldsymbol{\delta}_P(t)$ be the total offset vector. At the precise moment of functional contact, the functional distance is small but non-zero ($d_{\text{func}} \leq R_{\text{contact}} \approx 40$ nm). The measured distance is the magnitude of the vector sum of this contact separation and the label offset:

$$d_{\text{meas}}(t_{\text{contact}}) = \|(\mathbf{r}_E - \mathbf{r}_P) + \boldsymbol{\Delta}(t)\| \tag{145}$$

By the triangle inequality, the measured distance at contact is bounded by $\|\boldsymbol{\Delta}\| - R_{\text{contact}} \leq d_{\text{meas}} \leq \|\boldsymbol{\Delta}\| + R_{\text{contact}}$. Since the label offset $\|\boldsymbol{\Delta}\|$ often significantly exceeds the contact radius (e.g., hundreds of nanometers for separations of several kilobases), Equation 145 establishes a "Noise Floor". Even when the enhancer and promoter are physically touching ($d_{\text{func}} = 0$), the microscope records a distance distributed around $\|\boldsymbol{\Delta}\|$.

To quantify this effect, we simulated the expected distribution of offset magnitudes $\|\boldsymbol{\Delta}\|$ for the experimental configuration (**Supplementary Note 1 Fig. 1A**) that reported no correlation between E-P spatial proximity and transcription [84]. Briefly, we performed 3D polymer simulations using the same methods and parameters as those in the main text, where we defined a single chromosome consisting of 70,000 lattice sites with each site corresponding to 1 kb of DNA. We repeated the 109 kb E-P pair along with the surrounding CTCF binding sites (**Supplementary Note 1 Fig. 1A**) 7 times on the chromosome. The initial conformation of the polymer was taken from previous simulation that was run for more than 15 million simulation time steps [112], and then normally distributed velocities and harmonic bonds between cohesins were added, and another 100,000 simulation time steps were run to allow the polymer to relax into a steady-state conformation before recording 3D distances between the E-P pair and the probes placed at different genomic distances.

In scenarios where labels are placed far from the functional sites, such as the 11–13 kb separation reported previously [84], the offset vector does not average to zero but instead creates a substantial baseline distance. (**Supplementary Note 1 Fig.2**) illustrates this distribution obtained from our 3D polymer simulations, showing that even in the case of perfect functional contact, the measured distance would typically register as nearly 200 nm, effectively masking the interaction event from detection.

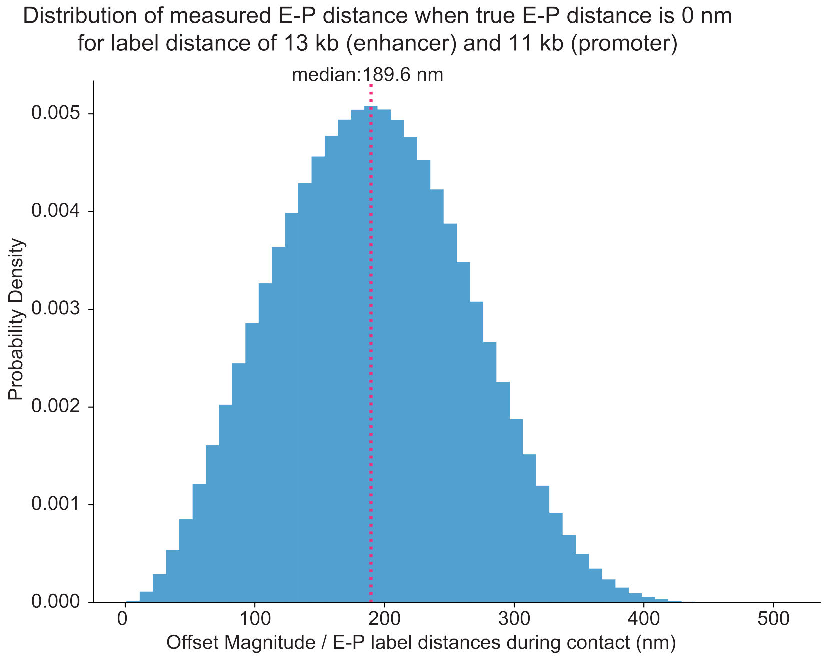

**Supplementary Note 1 Fig. 2: The measured 3D distance can be very large even when the true 3D distance is 0 nm.** Simulation of the offset vector magnitude $\|\boldsymbol{\Delta}\|$ for the experimental configuration used in Alexander et al. 2019 [84], where the center-to-center distance for label-enhancer and the label-promoter was 13 kb and 11 kb respectively. Even with direct contact ($d_{\text{func}} \approx 0$), the measured distance would center around a median offset of $\sim$190 nm, obscuring the interaction event.

#### 11.3 Stochastic transcription modeling and MS2 signal generation

To assess the impact of label positioning and temporal dynamics on transcription, we next coupled the 3D E-P polymer simulations to nascent transcription for a "hit-and-run" contact model, where the enhancer can activate the promoter whenever $d_{\text{func}} \leq R_{\text{contact}} \approx 40$nm. Specifically, we coupled our polymer simulations to a stochastic telegraph model of transcription. This model causally links physical E-P contact to transcriptional bursting and simulates the resulting fluorescent signals from nascent RNAs. Thus, it allows us to assess whether experimental and analytical limitations can obscure the causal relationship between E-P contact and nascent transcription.

##### 11.3.1 Two-state promoter model with distance-dependent switching

We modeled the promoter as a two-state system, switching between an inactive (OFF) state, $P_{\text{OFF}}$, and an active (ON) state, $P_{\text{ON}}$. The promoter can only produce mRNA in the ON state.

$$\mathrm{P_{OFF}} \underset{k_r}{\overset{k_f}{\rightleftarrows}} \mathrm{P_{ON}} \xrightarrow{k_{\text{load}}} \mathrm{mRNA}$$

Here we assume the promoter does not transcribe in the OFF state for simplicity, since the basal transcription activity of the *Sox2* promoter simulated here was reported to be negligible [84]. However, most endogenous promoters exhibit some level of basal transcription activity. This basal activity would make detecting a dip in E-P distance at transcription burst onsets even more challenging, as not all observed bursts would be attributable to E-P interaction events.

The promoter state transition probabilities are governed by the instantaneous functional distance $d_{\text{func}}(t)$ between the enhancer and promoter loci in the simulation. Let $S(t) \in \{0, 1\}$ represent the promoter state at time $t$. The probability of switching from OFF to ON in a time step $\Delta t$ is defined as:

$$P(S(t+\Delta t) = 1 | S(t) = 0) = \begin{cases} 1 - e^{-k_{\text{on}}\Delta t} & \text{if } d_{\text{func}}(t) < R_{\text{contact}} \\ 0 & \text{otherwise} \end{cases} \tag{146}$$

where $k_{\text{on}}$ is the activation rate and $R_{\text{contact}}$ is the contact threshold (1.5 monomers $\approx$ 40 nm). Conversely, the promoter turns OFF with a constant rate $k_{\text{off}}$:

$$P(S(t+\Delta t) = 0 | S(t) = 1) = 1 - e^{-k_{\text{off}}\Delta t} \tag{147}$$

In our simulations, we used the following parameters, constrained by prior studies: $k_{\text{off}} = 0.5\text{min}^{-1}$ [97] and $k_{\text{on}} = 5\text{min}^{-1}$ [110].

##### 11.3.2 Polymerase II loading and signal convolution

When the promoter is in the ON state ($S(t) = 1$), productive Polymerase II (Pol II) initiation events were modeled as a Poisson process with rate $k_{\text{load}}$:

$$N_{\text{pols}}(t) \sim \text{Poisson}(k_{\text{load}}\Delta t) \tag{148}$$

where $N_{\text{pols}}(t)$ is the number of polymerases loaded at time $t$. We set $k_{\text{load}} = 10.0$ min$^{-1}$ based on prior findings [113]. The observed MS2 fluorescence signal $I(t)$ is the convolution of the Pol II loading history $N_{\text{pols}}(t)$ with a gene-specific kernel $K(x)$ that describes the fluorescence intensity of a single polymerase as it traverses the gene:

$$I(t) = \sum_{\tau=0}^{T_{\text{elongation}}} N_{\text{pols}}(t - \tau) \cdot K(v \cdot \tau) \tag{149}$$

where $v$ is the elongation speed (2000 bp/min [114,115]) and $K(x)$ represents the number of MS2 loops transcribed at position $x$ along the gene. Here, $T_{\text{elongation}}$ represents the total residence time of a polymerase on the gene, defined as $T_{\text{elongation}} = L_{\text{gene}}/v$. This variable sets the temporal memory of the system; the observed signal at any instant is the cumulative sum of all initiation events that occurred within the window $[t - T_{\text{elongation}}, t]$.

##### 11.3.3 Impact of 5' UTR / first-intron vs. 3' UTR labeling

The shape of the kernel $K(x)$ dictates how quickly a transcriptional event is converted into an observable signal (see figure below). For 5' UTR or first-intron labeling, $K(x)$ rises early, providing an almost immediate readout of promoter activity. In contrast, 3' labeling introduces a gap where $K(x) = 0$ for the majority of the elongation time, effectively delaying the signal relative to the physical E-P contact event (**Supplementary Note 1 Fig.3**). We modeled two distinct scenarios:

- **First intron MS2 labeling near the 5'UTR:** The MS2 cassette is placed in the first intron of the gene. The kernel $K(x)$ rises rapidly as the polymerase transcribes the loops but drops to zero immediately after the cassette is fully transcribed ($x > L_{\text{cassette}}$), creating a transient "shark fin" profile:

$$K(x) = \begin{cases} \min\left(\left\lfloor \frac{x}{L_{\text{loop}}} \right\rfloor + 1, N_{\text{loops}}\right) & \text{for } x \le L_{\text{cassette}} \\ 0 & \text{for } x > L_{\text{cassette}} \end{cases} \tag{150}$$

  We implemented this transient profile because, in our study, the MS2 cassette is encoded within the first intron; since splicing frequently occurs co-transcriptionally in mammalian cells [81–83], the kernel is modeled to drop to zero once the MS2 sequence is transcribed to represent the excision and release ("falling off") of the intron while the polymerase continues elongation. We note that the precise timing and fidelity of co-transcriptional splicing is uncertain and may be gene specific [81–83]. However, for simplicity we model it as instantaneous in this note.

- **3' UTR labeling:** The MS2 cassette is placed at the end of the gene (length $L_{\text{gene}}$). The polymerase must traverse the entire gene body before the signal appears. The kernel $K_{3'}(x)$ remains zero until $x > L_{\text{gene}} - L_{\text{cassette}}$:

$$K_{3'}(x) = \begin{cases} 0 & \text{if } x < L_{\text{gene}} - L_{\text{cassette}} \\ \min\left(\left\lfloor \frac{x-(L_{\text{gene}}-L_{\text{cassette}})}{L_{\text{loop}}} \right\rfloor + 1, N_{loops}\right) & \text{otherwise} \end{cases} \tag{151}$$

  This introduces a significant temporal delay $\tau_{\text{delay}} = (L_{\text{gene}} - L_{\text{cassette}})/v$.

During this delay $\tau_{\text{delay}}$, the chromatin polymer continues to diffuse until the moment of signal detection ($t_{\text{detect}} = t_{\text{contact}} + \tau_{\text{delay}}$). Consequently, 3' labeling obscures the 3D distance dip significantly more than 5' labeling or first-intron labeling, providing a mechanistic explanation for why studies utilizing 3' labeling [84], especially if the gene is long, may fail to detect correlations between E-P spatial proximity and transcription.

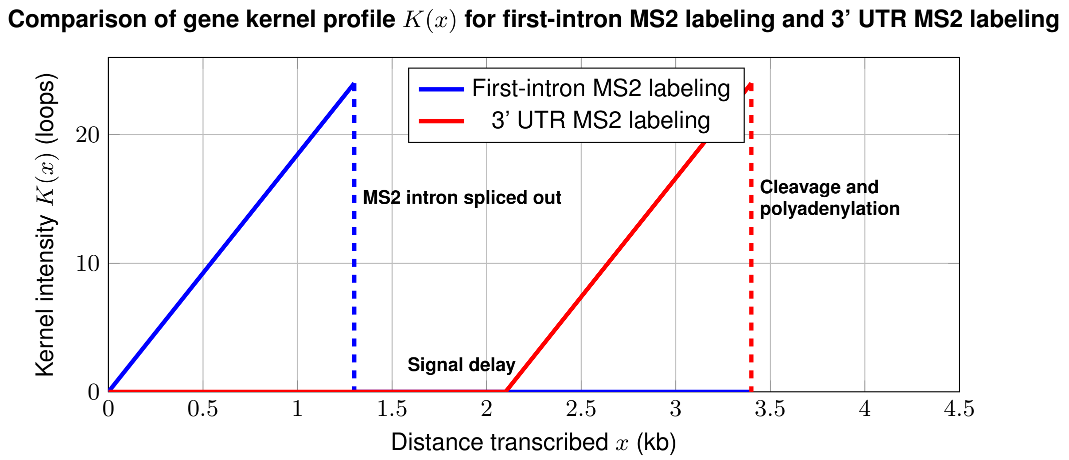

**Supplementary Note 1 Fig. 3: Fluorescence kernels for different MS2 label positions.** The plot reflects the specific kernel logic used in the simulation. With labeling in the first intron (Blue), the kernel provides an immediate signal but is transient, dropping to zero once the cassette is transcribed ($x > 1.3$ kb) to mimic the co-transcriptional splicing of the MS2-labeled intron. With 3' labeling (Red), the kernel remains at zero for the majority of transcription, introducing a delay that decouples the observed signal from the initial E-P contact event.

##### 11.3.4 Modeling stochastic elongation and jitter

While Equation (150)–(151) describes the deterministic signal generation, *in vivo* elongation is stochastic due to polymerase pausing and traffic. To account for this in our analysis, we model the effective delay $\Delta t$ between transcription initiation and the burst onset as a random variable drawn from a Gamma distribution, where we define burst onsets as the MS2 signal appearance from the first Pol II loading event in a continuous window when the promoter is the ON state:

$$\Delta t \sim \Gamma(k, \theta) \tag{152}$$

Mechanistically, a gamma distribution is chosen because transcription elongation consists of a sequence of discrete, stochastic steps; the total time to traverse the probe region is therefore the sum of many independent exponential dwell times, which can be described by a Gamma distribution [116]. While the Gamma distribution mathematically approximates a Gaussian distribution in the limit of a large number of steps, we utilize the Gamma distribution here for two reasons. First, it accommodates the potential for rate-limiting pauses that reduce the effective number of stochastic steps, resulting in a skewed delay profile. Second, unlike a Gaussian distribution, the Gamma distribution has positive support, enforcing the constraint that temporal delays cannot be negative.

The shape ($k$) and scale ($\theta$) parameters are derived from the mean delay $\mu = L_{probe}/v$ and the coefficient of variation ($CV$), which we set to 0.6 to model elongation jitter:

$$k = \frac{1}{CV^2}, \quad \theta = \mu \cdot CV^2 \tag{153}$$

This stochastic treatment ensures that the "Ground Truth" burst onset times used for pileup analysis reflect the realistic temporal dispersion of polymerases arriving at the MS2 probe site. We note that this treatment is optimistic, as e.g. stochastic binding of elongation factors would introduce additional variation in delays beyond what is modeled here.

##### 11.3.5 The Consequence of delays and sampling

(**Supplementary Note 1 Fig.4**) illustrates how these multiple factors—label offset, elongation delay, and sparse sampling—combine to obscure the E-P interaction event. In the simulated trajectory shown, the "trigger" interaction occurs at $t = 0$ (green vertical line, top panel), where the true E-P distance (blue line, top panel) drops below 40 nm for a sufficient duration that the promoter switched to the ON state. In the ON state, Pol II can load, but this incorporates another delay between the promoter switching ON and the first Pol II loading event. Further, because the MS2 labels are at the 3' end, the fluorescence signal (gray line, bottom panel) does not begin to rise until ~80 seconds later (the elongation delay). Because the distance is measured between labels separated by 13 kb and 11 kb (pink line, top panel), even with unrealistically fast sampling, the measured E-P distance was above 200 nm even when the true E-P distance is around 40 nm. By the time an increase in MS2 signal is detected at around $t = 100$s (vertical orange line, bottom panel), the polymer has diffused away, and the measured 3D distance has now relaxed back to ~300 nm. Note that we have not added localization error in the simulated E-P distance measurements here, and typical localization error on the order of 10s of nanometers in each dimension would make it even more challenging to detect the dip in E-P distance [49, 103]. Thus, a naive correlation analysis would associate the high MS2 signal with a large E-P distance, completely missing the causal contact event that occurred two minutes earlier, which we explore further below. This example highlights how challenging it is to detect E-P contact mediated promoter activation.

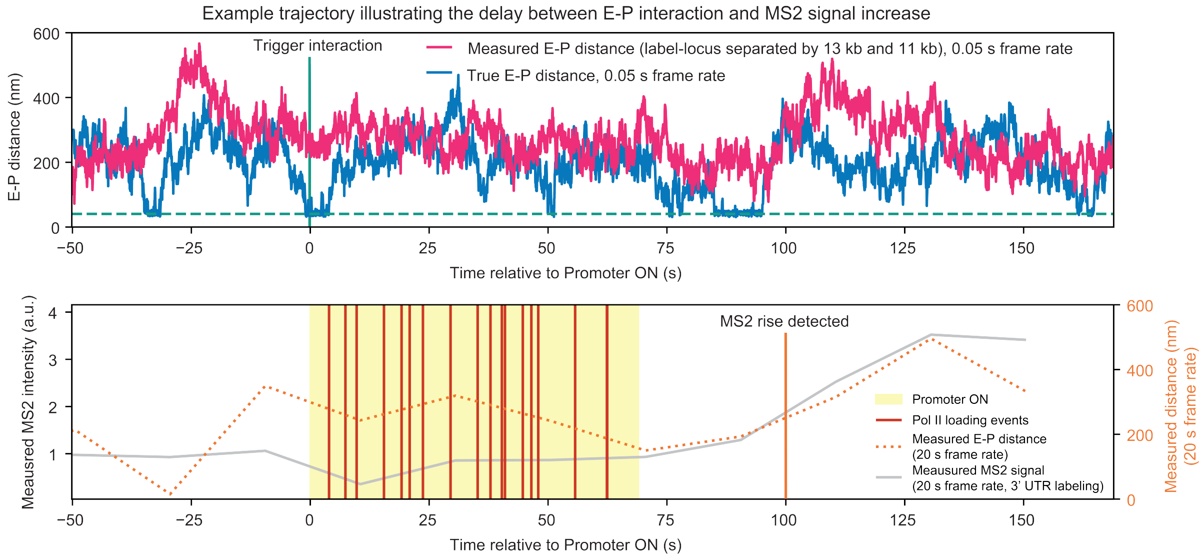

**Supplementary Note 1 Fig. 4: Temporal decoupling of E-P interaction and transcription signal.** Representative simulation trajectories showing the sequence of events: (1) Physical E-P contact at $t = 0$ (blue line drops to the green dashed line, $< 40$ nm), with the label offset (pink vs blue line) further compressing the dynamic range of the distance measurement and obscuring E-P interactions; (2) Promoter turns ON (the yellow window) and Pol II loads (red vertical lines); (3) Elongation delay of ~80s before 3' MS2 signal increases; (4) Sparse sampling (dotted orange line) captures the distance long after the E-P interaction has vanished.

#### 11.4 The impact of conventional signal processing (burst calling)

Even if the experimental aspects are optimized by considering all factors discussed above, another important dimension that can obscure the correlation of E-P 3D proximity and transcription is data analysis. Since the only experimental measurement for transcription is MS2 signal, a common and intuitive approach of correlating E-P distance and transcription is to perform an MS2 burst pile-up analysis [84, 111]: 1) identify windows of transcription by processing the MS2 signal to call bursts that correspond to increased transcription activity 2) pile up E-P distance trajectories at the beginning of the called bursts and see whether we can detect a dip in E-P distances right before the called bursts. As we showed above, due to the significant delay between the E-P interaction event that triggers promoter state switching and the rise of MS2 signal, such an approach would obscure the dip in E-P distance due to the variable delay composed of several stochastic steps.

Moreover, we illustrate below that the "memory" of the promoter state means one E-P interaction event can lead to several called bursts, further obscuring the correlation between E-P spatial proximity and transcription.

##### 11.4.1 Burst definition and burst calling algorithm

The precise definition of what constitutes a burst varies among studies [117–119] and often two definitions are used. First, a burst can be observationally defined as a "continuous period of MS2 signal". This has the advantage of being directly related to the observed experimental MS2 trajectories, but it is less well mechanistically defined and the resulting parameters (e.g. burst duration) depend strongly on whether e.g. the MS2 array is placed in the 5'UTR, an intron, or the 3'UTR. Second, a burst can be defined as a "continuous period of the promoter being in the ON state". This has the advantage of being mechanistically interpretable and, in this case, the beginning of a promoter ON state is directly related to the initiating E-P contact event. However, since promoter state cannot be directly observed in live cells, this has the disadvantage of requiring more complex analysis methodology and assumptions. Because the first definition is most commonly used including in prior studies relating E-P interactions to transcription [84, 111], we proceed with this "MS2 signal" based definition here.

We employed a change point detection algorithm to call MS2 burst: bursts are identified as peaks in the fluorescence signal in the MS2 channel that exceed a minimum height and a minimum vertical prominence (quantifying vertical drop on either side of the peak). This ensures that only distinct events, separated by a significant drop in signal, are counted as separate bursts, reducing false positives from small fluctuations on top of a plateau. The ”called burst onsets” are therefore defined as the start time of each called MS2 burst.

###### 11.4.2 Promoter memory and multiple called bursts

In the example below (**Supplementary Note 1 Fig. 5A**, left), the promoter enters a sustained ON state (yellow shaded region) due to an E-P interaction event. During this window when the promoter is in the ON state, multiple Pol II loading events occur leading to several called bursts. The first called burst aligns reasonably well with the start of promoter ON state (though delayed by elongation), whereas the subsequent called bursts do not correspond to distinct windows where the promoter is in the ON state. This effect is even more pronounced when simulating with smaller $k_{\mathrm{off}}$ and smaller $k_{\mathrm{load}}$ values that are still biologically plausible [83] (**Supplementary Note 1 Fig. 5A**, right). When aligning distance traces to these later bursts, the E-P interaction that triggered the state has long since dissolved. Including these bursts in a MS2 pileup analysis dilutes the signal, as they contribute distance trajectories that are far less correlated with the initial contact event (**Supplementary Note 1 Fig. 5B**). This phenomenon further explains why common and intuitive MS2 pile-up analyses may fail to recover the sharp decrease in E-P 3D distance at burst onsets.

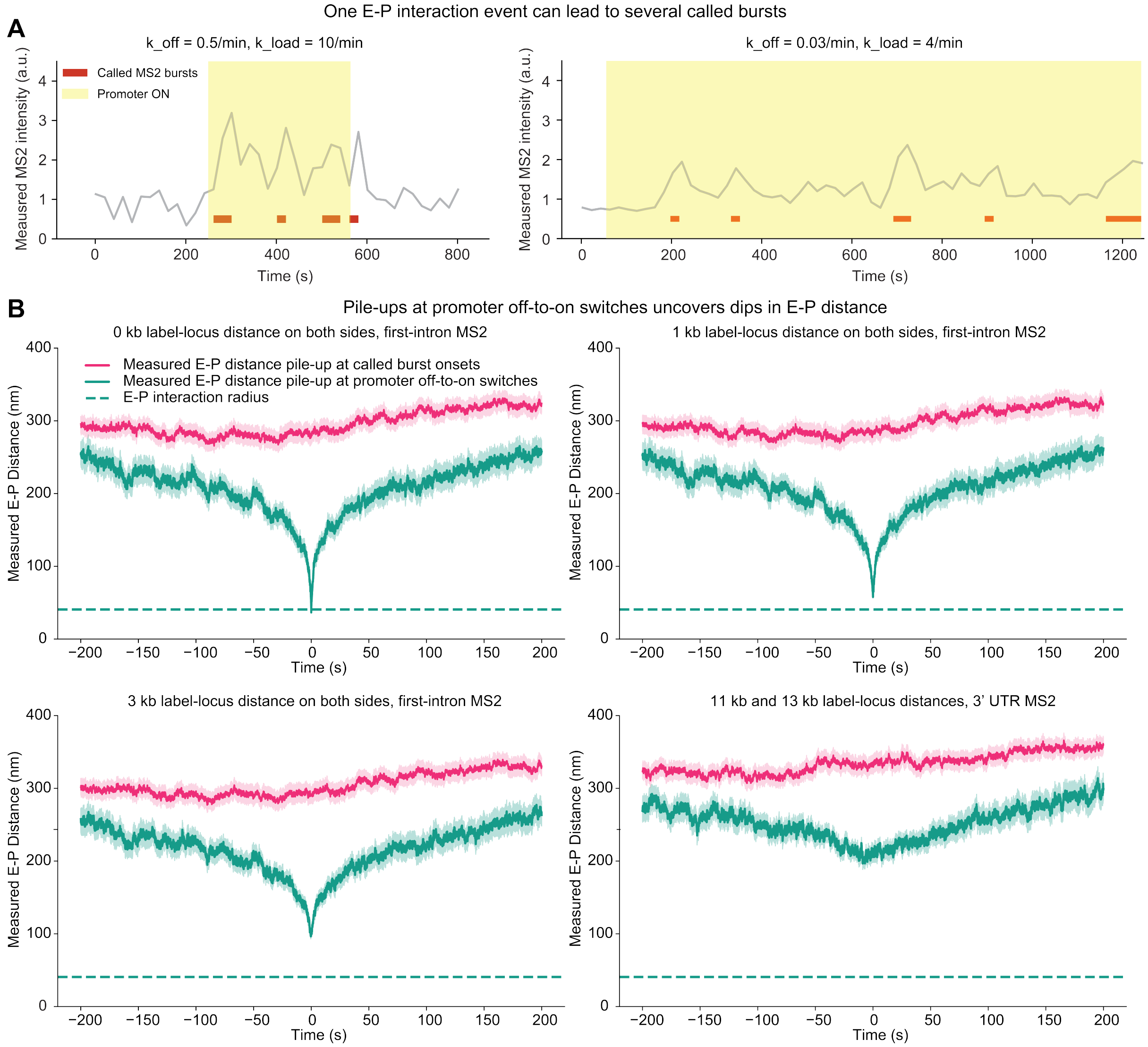

**Supplementary Note 1 Fig. 5: Calling bursts via MS2 signal processing.** (**A**) A single, continuous promoter ON state (yellow shading) can produce a fluctuating MS2 signal that is interpreted as multiple distinct bursts (red bars) by peak-calling algorithms. Aligning E-P distance trajectories to these called bursts dilutes the correlation signal, as the original E-P contact event occurred minutes earlier. (**B**) Piling up E-P distances at called bursts obscures the dip in E-P distance (magenta). In contrast, one can uncover the dip in E-P distance if piling up at promoter OFF-to-ON switches instead (green). The solid lines are average E-P distance of the pile-up and the shaded area around them shows the standard error of the mean.

##### 11.4.3 The need for advanced analysis frameworks

The discrepancies highlighted above demonstrate that intuitive and common analysis methods, which rely on simple thresholding or peak detection of the MS2 signal to define transcriptional "events," are insufficient for capturing the underlying causal interactions. The temporal stochasticity of steps leading to MS2 signal increase and the "memory" of the promoter state mean that the observed MS2 signal is a delayed and distorted convolution of the true promoter activity. To resolve this, in our study we aim to move beyond identifying phenomenological "bursts" and instead adopt more sophisticated analysis frameworks capable of inferring the underlying system states:

- **Machine learning:** We developed a Long Short-Term Memory (LSTM) based deep learning framework to map the complex temporal relationship between 3D spatial dynamics and transcriptional output. The LSTM was trained using 3D E-P distances as the sole input to predict the onset of transcription bursts extracted from the MS2 signal (**Fig. 3**). This approach allowed us to isolate and uncover the signatures of 3D E-P distance dynamics that precede burst onsets.

- **Statistical inference:** We developed a comprehensive inference framework to recover the underlying dynamics of both transcription and E-P interactions from noisy experimental data (**Fig. 4**). First, we employed Bayesian inference to deconvolve stochastic MS2 signals, estimating the posterior probability of promoter states to pinpoint moments of gene activation. Concurrently, we utilized Gaussian Process Regression to model the E-P distance trajectories, allowing us to infer continuous, probabilistic E-P distance functions from discrete, noisy tracking data. Bridging these inferred activation moments with the modeled E-P distance allows us to robustly detect E-P interactions driving transcription.

These approaches were informed by the simulations and analyses outlined above and provide a handle for identifying the true burst onsets—defined as the first Pol II loading event in a continuous ON window—and, importantly, the precise moment the promoter switches to the ON state. By aligning 3D distance measurements to these causative events, we were able to robustly detect the transient E-P interactions that drive transcription.

#### 11.5 Recipe for correlating E-P spatial proximity and transcription

In summary, to reconcile the seemingly contradictory observations between imaging and genomics approaches, we highlight several important factors one must consider to uncover the temporal correlation between E-P 3D proximity and transcription (several of which have also been previously discussed [49, 103]):

- **Small fluorescent labels:** Avoid large arrays to label E-P pairs. Use small fluorogenic systems (e.g., ParB/*parS*) to label functional elements to minimize $\|\mathbf{\Delta}\|$.
- **Minimize separation between E,P and fluorescent labels:** To serve as a good positional reporter, the fluorescent array should be placed as close to the functional element as possible to minimize $\|\mathbf{\Delta}\|$.
- **Small enhancers and promoters:** Large enhancers and promoters will effectively increase the separation between E,P and fluorescent labels.
- **5' labeling or first-intron labeling:** Place the RNA labeling array in the 5' UTR or in the first intron to minimize the time-delay ($\tau$) between physical proximity and RNA detection.
- **Insulation:** Ensure the E-P pair is insulated from other regulatory elements and choose a promoter with minimal basal activity so that the observed transcription is entirely dependent on, and can be attributed to, interactions with a single enhancer.
- **Maximize E-P separation:** To distinguish "E-P interaction" from "lack of E-P interaction", it helps to choose E-P pairs with large genomic separation ($> 100$ kb).
- **High frequency/long duration:** High frequency prevents aliasing of transient contacts; long duration accumulates enough events ($N$) to recover the dip from statistical noise ($\sigma \propto 1/\sqrt{N}$).
- **Alternative analysis accounting for noise and delays:** Be careful with binary MS2 burst calling. Use statistical inference or machine learning approaches to account for noise and time-delays, and to uncover the underlying promoter states.

#### 11.6 Conclusion

Two arguments have commonly been used to rule out E-P contact mediated promoter activation. First, the large measured E-P distances during burst onsets (~100–300 nm) intuitively seem incompatible with E-P contact models. Second, the lack of an E-P distance "dip" in MS2 pile-up analysis appears to rule out E-P contact [84]. We show here that neither argument rules out E-P contact models: large measured E-P 3D distances and a lack of an E-P distance "dip" in MS2 pile-up plots are fully consistent with a dynamic E-P hit-and-run contact model. Thus, our findings here are not inconsistent with prior studies. Because detecting transient E-P contact mediated promoter activation is at the very edge of current technical capabilities, extremely careful experimental design and analysis is necessary. However, we stress that this does not mean that a transient "hit-and-run" E-P contact model applies universally and to other E-P pairs studied previously, such as e.g. *Sox2* [84]. It simply means that transient E-P contact cannot be ruled out on the basis of prior studies, as also previously discussed [49, 103]. Determining to what extent spatiotemporal E-P interaction mechanisms are universal or E-P pair specific is an important area for future studies. Finally, we acknowledge we have not exhausted the simulation parameter space here. Nevertheless, the conclusions generally hold when varying different parameters and our simulations highlight important experimental and computational considerations essential for uncovering the correlation between E-P 3D proximity and transcription, if it exists.

Our mathematical and computational analysis demonstrates that label-locus separation and conventional burst-calling can easily mask the signature of E-P interactions.

#### 12 Supplementary Note 2: mathematical derivation for RCMC vs flow cytometry comparison

##### 12.1 Introduction and problem statement

The primary objective of this derivation is to establish a theoretical framework to interpret the relationship between protein expression levels ($F$), measured by flow cytometry, and physical interaction probabilities of chromosome conformation capture ($f$), measured by Region Capture Micro-C (RCMC) [56], a type of 3C assay. Empirically, we observed a linear relationship between these two metrics across various cell lines with different E-P configurations. However, the interpretation of this linearity is not immediately obvious. Specifically, we aim to understand what this linearity implies about the relationship between two critical length scales:

- **The functional interaction radius ($R_{\mathrm{E-P}}$):** The effective 3D spatial distance threshold required for an enhancer to physically engage a promoter and activate transcription.
- **The experimental capture radius ($R_{\mathrm{RCMC}}$):** The 3D distance threshold within which two genomic loci must reside to be successfully crosslinked, ligated and sequenced in the RCMC assay.

In this Supplementary Note, we investigate whether the observation of linearity ($F \propto f$) provides evidence that the functional interaction radius and the experimental capture radius are equivalent ($R_{\mathrm{E-P}} \approx R_{\mathrm{RCMC}}$). We structure the argument in three parts:

1. **Part 1 (The simple polymer baseline):** We first examine the relationship for simple polymers with broad distance distributions. We demonstrate that in the limit of small contact radii, linearity is a geometric consequence of volume scaling, expected regardless of whether the functional interaction radius matches the experimental capture radius. We also define the boundaries of this approximation, showing that outside the small-radius limit, linearity imposes stricter constraints on the relationship between $R_{\mathrm{E-P}}$ and $R_{\mathrm{RCMC}}$.
2. **Part 2 (Chromatin scaling):** While Part 1 establishes the baseline, Part 2 is necessary to define the explicit mathematical form of the "background" polymer state. By deriving the power-law scaling, we establish how the contact probability of the background chromatin scales with capture volume ($r_i^3$) and genomic distance ($s^{-3\nu}$). This specific scaling is what allows us to mathematically distinguish the "background" component from the "stabilized" component in Part 3.
3. **Part 3 (The critical deviation):** We introduce the biological reality by considering protein-mediated transient interactions, such as CTCF-mediated loops or transient bonds due to affinity ("stickiness") between the E-P pair. We show that these interactions disrupt the simple volume scaling derived in Part 1. Importantly, we demonstrate that in the presence of such transient interactions, a linear relationship is *only* mathematically possible if the experimental capture radius is effectively equal to the functional radius ($R_{\mathrm{RCMC}} \approx R_{\mathrm{E-P}}$).

**Summary of theoretical findings:** Overall, we show that the linear relationship between expression (as measured by flow cytometry) and interaction probability (as measured by RCMC) that we observe can only exist if $R_{\mathrm{E-P}} \approx R_{\mathrm{RCMC}}$, that is, if the functional E-P interaction radius $R_{\mathrm{E-P}}$ is approximately equal to the RCMC capture radius $R_{\mathrm{RCMC}}$. Specifically, for stabilized interactions (i.e. those that show enrichment over background and appear as a 'dot' in RCMC maps and which may be stabilized by CTCF sites and/or affinity between CREs), a non-linear relationship between expression and RCMC interaction probability would be expected unless $R_{\mathrm{E-P}} \approx R_{\mathrm{RCMC}}$. Thus, the fact that we observe a linear relationship implies that $R_{\mathrm{E-P}} \approx R_{\mathrm{RCMC}}$ and since $R_{\mathrm{RCMC}}$ has been estimated to be on the order of 42 nm [60], the observed linearity can to our knowledge only be explained by a contact-like functional E-P interaction radius. Outside this small-radius limit, the curvature of the distribution also implies that linearity requires $R_{\mathrm{E-P}} \approx R_{\mathrm{RCMC}}$. Furthermore, for chromatin containing stabilized interactions, visible as "dots" in RCMC maps (Part 3), the presence of a "stabilized" state breaks the trivial volume scaling entirely. Consequently, observing linearity in real biological data—which likely contains transient bonds—provides the necessary constraint to conclude that the functional interaction radius $R_{\mathrm{E-P}}$ must be approximately equal to the RCMC capture radius $R_{\mathrm{RCMC}}$.

##### 12.2 Part 1: General derivation for simple polymers

Before assuming a specific polymer model for chromatin, we consider the general conditions under which a plot of protein expression against contact probability yields a linear relationship. This section serves as a "null model"—a theoretical baseline for a polymer lacking specific structural features such as CTCF mediated loops. We define $c_i$ as a specific E-P configuration (genomic distance and CTCF binding site arrangement), and note that in this study, the same enhancer and the same promoter were used in these different E-P configurations. We make two fundamental assumptions:

1. The RCMC signal is proportional to the cumulative contact probability at the experimental radius $R_{\mathrm{RCMC}}$ [103].
2. The protein expression level is proportional to the cumulative contact probability at the functional radius $R_{\mathrm{E-P}}$ [105].

###### 12.2.1 Taylor expansion at small contact radii

For small contact radii, we can approximate the probability of contact by examining the probability density function (PDF) of the E-P distance vector, $p(\boldsymbol{r}|c_i)$, near the origin. The cumulative contact probability $p_c$ is defined as the integral of this PDF over the spherical capture volume defined by radius $R_{\mathrm{RCMC}}$:

$$p_c(c_i, R_{\mathrm{RCMC}}) = \int_{|\boldsymbol{r}|\leq R_{\mathrm{RCMC}}} p(\boldsymbol{r}|c_i)\, d\boldsymbol{r} = 4\pi \int_0^{R_{\mathrm{RCMC}}} r^2 p(\boldsymbol{r}|c_i)\, dr \tag{154}$$

Assuming the distribution $p(\boldsymbol{r}|c_i)$ is continuous and varies slowly near the origin (a valid assumption for broad polymer distributions where the mean separation is much larger than $R_{\mathrm{RCMC}}$), we can perform a Taylor expansion of the density around $\boldsymbol{r} = 0$:

$$p(\boldsymbol{r}|c_i) = p(\mathbf{0}|c_i) + \nabla p(\mathbf{0}|c_i) \cdot \boldsymbol{r} + \mathcal{O}(r^2) \tag{155}$$

Substituting the leading order term ($p(\mathbf{0}|c_i)$) back into the integral:

$$p_c(c_i, R_{\mathrm{RCMC}}) \approx 4\pi \int_0^{R_{\mathrm{RCMC}}} r^2 p(\mathbf{0}|c_i)\, dr = p(\mathbf{0}|c_i) \left[\frac{4\pi}{3} r^3\right]_0^{R_{\mathrm{RCMC}}} \tag{156}$$

$$p_c(c_i, R_{\mathrm{RCMC}}) \approx p(\mathbf{0}|c_i) \frac{4\pi}{3} R_{\mathrm{RCMC}}^3 \tag{157}$$

###### 12.2.2 Separation of variables: two multiplicative terms

Equation (157) yields an important result: the contact probability separates into two distinct multiplicative terms.

- **The condition-dependent term ($h$):** Let $h(c_i) = p(\mathbf{0}|c_i)$. This term represents the local density at the origin. It encapsulates all the biology and physics of the specific E-P configuration $c_i$ (e.g., genomic distance and CTCF arrangement).
- **The geometric term ($g$):** Let $g(R_{\mathrm{RCMC}}) = \frac{4\pi}{3} R_{\mathrm{RCMC}}^3$. This term depends solely on the volume of the capture sphere and is independent of the E-P configuration.

###### 12.2.3 Implication for linearity in simple polymers

We now examine the ratio of the measured flow cytometry signal ($F$) to the RCMC signal ($f$) for a set of different E-P configurations $\{c_i\}$. These different E-P configurations have different E-P genomic distances and different CTCF binding site arrangements, but have the same enhancer and the same promoter. Due to the fact that the same enhancer and the same promoter were used, we can expect these E-P configurations' expression to have the same dependency on contact probability at functional radius $R_{\mathrm{E-P}}$:

$$F(c_i) \approx C_{\mathrm{E-P}} \cdot p_c(c_i, R_{\mathrm{E-P}}) \tag{158}$$

where $C_{\mathrm{E-P}}$ is a constant factor for this specific choice of E-P sequence. For these different constructs, the RCMC assay followed the same protocol and analysis, and thus we can expect the RCMC signal to have the same dependency on the contact probability at capture radius $R_{\mathrm{RCMC}}$:

$$f(c_i) \approx C_{\mathrm{RCMC}} \cdot p_c(c_i, R_{\mathrm{RCMC}}) \tag{159}$$

Thus we can simplify the ratio of the measured flow cytometry signal ($F$) to the RCMC signal ($f$):

$$\frac{F(c_i)}{f(c_i)} \approx \frac{C_{\mathrm{E-P}} \cdot p_c(c_i, R_{\mathrm{E-P}})}{C_{\mathrm{RCMC}} \cdot p_c(c_i, R_{\mathrm{RCMC}})} \approx \frac{C_{\mathrm{E-P}} \cdot h(c_i) g(R_{\mathrm{E-P}})}{C_{\mathrm{RCMC}} \cdot h(c_i) g(R_{\mathrm{RCMC}})} = \frac{C_{\mathrm{E-P}} \cdot g(R_{\mathrm{E-P}})}{C_{\mathrm{RCMC}} \cdot g(R_{\mathrm{RCMC}})} \tag{160}$$

Substituting in the geometric term from above, we get:

$$\frac{F(c_i)}{f(c_i)} \approx \frac{C_{\mathrm{E-P}} \cdot p(\mathbf{0}|c_i) \frac{4\pi}{3} R_{\mathrm{E-P}}^3}{C_{\mathrm{RCMC}} \cdot p(\mathbf{0}|c_i) \frac{4\pi}{3} R_{\mathrm{RCMC}}^3} = \frac{C_{\mathrm{E-P}} \cdot R_{\mathrm{E-P}}^3}{C_{\mathrm{RCMC}} \cdot R_{\mathrm{RCMC}}^3} \tag{161}$$

Thus, in the case of unstabilized chromosomal interactions, e.g. E-P interactions that do not result in a 'dot' on RCMC maps, a general linear relationship can be expected to hold regardless of the specific values of $R_{\mathrm{E-P}}$ and $R_{\mathrm{RCMC}}$.

**Intuitive interpretation:** imagine the enhancer as a "cloud" of probability surrounding the promoter. The density of this cloud at the center depends on the specific chromatin structure $c_i$ (e.g., a short genomic distance creates a dense cloud; a long distance creates a sparse cloud). $p_c(c_i, R_{\mathrm{RCMC}})$ measures the amount of this cloud captured within a radius $R_{\mathrm{RCMC}}$. As long as the radius is small enough that the cloud density is roughly uniform inside them, both measurements are simply equal to the local density multiplied by the spherical volume. When we take the ratio $F/f$, the density (which varies between different E-P configurations) cancels out completely. We are left with only the ratio of volumes multiplied by some constants ($\frac{C_{\mathrm{E-P}} \cdot R^3_{\mathrm{E-P}}}{C_{\mathrm{RCMC}} \cdot R^3_{\mathrm{RCMC}}}$). Consequently, for simple polymers in the small-radius limit, a linear relationship is guaranteed by geometry alone. It does not imply that the functional and experimental radii are equal, only that they are both probing the same local density environment. Nevertheless, as we shall see below, the expectation of a linear relationship breaks down if chromosomal interactions are stabilized.

###### 12.2.4 Generalization to finite contact radii

The Taylor expansion derived above holds strictly in the limit where the capture radius is much smaller than the mean spatial separation of the loci ($R_{\mathrm{RCMC}} \ll \sqrt{\langle R^2 \rangle}$). However, if the contact radii are not small, we cannot assume that the probability density is constant near the origin. We must consider the exact form of the distribution.

Assuming a Gaussian distribution for the E-P separation vector (as defined later in Equation (167)), the cumulative probability of contact within a radius $r$ for a polymer with mean squared separation $\sigma_i^2 = \langle R^2(c_i) \rangle$ is given by the exact integration:

$$p_c(c_i, r) = 4\pi \int_0^r \rho^2 \left( \frac{3}{2\pi\sigma_i^2} \right)^{3/2} \exp\left( -\frac{3\rho^2}{2\sigma_i^2} \right) d\rho \tag{162}$$

This integral can be solved analytically using the error function (erf):

$$p_c(c_i, r) = \mathrm{erf}\left( \sqrt{\frac{3}{2}} \frac{r}{\sigma_i} \right) - \sqrt{\frac{6}{\pi}} \frac{r}{\sigma_i} \exp\left( -\frac{3r^2}{2\sigma_i^2} \right) \tag{163}$$

Let us define the scaling function $\Psi(x) = \mathrm{erf}(x) - \frac{2}{\sqrt{\pi}} x e^{-x^2}$. The contact probability becomes a function of the ratio between the capture radius and the mean spatial separation between the E-P: $p_c \propto \Psi\left( r\sqrt{3/2}/\sigma_i \right)$. We can now re-examine the ratio of expression ($F$) to RCMC signal ($f$) without the small-radius approximation:

$$\frac{F(c_i)}{f(c_i)} \approx \frac{C_{\mathrm{E-P}} \cdot \Psi\left( \frac{R_{\mathrm{E-P}}}{\sigma_i} \sqrt{\frac{3}{2}} \right)}{C_{\mathrm{RCMC}} \cdot \Psi\left( \frac{R_{\mathrm{RCMC}}}{\sigma_i} \sqrt{\frac{3}{2}} \right)} \tag{164}$$

**Consequence:** Unlike the Taylor expansion result in Equation (161), the polymer size $\sigma_i$ (which depends on genomic distance) does not automatically cancel out in this ratio. The function $\Psi(x)$ is non-linear. Consequently, the ratio $F/f$ will vary as a function of genomic distance ($\sigma_i$) unless the arguments of the numerator and denominator are identical. Therefore, for finite contact radii where the Taylor approximation does not apply, observing a linear relationship across constructs with varying genomic distances ($c_i$) implies that the numerator and denominator must be scaling identically. This is only possible if:

$$\Psi\left( \frac{R_{\mathrm{E-P}}}{\sigma_i} \sqrt{\frac{3}{2}} \right) \propto \Psi\left( \frac{R_{\mathrm{RCMC}}}{\sigma_i} \sqrt{\frac{3}{2}} \right) \implies R_{\mathrm{E-P}} \approx R_{\mathrm{RCMC}} \tag{165}$$

Thus, outside the very small radius limit, the observation of linearity provides even stronger evidence that the functional radius, $R_{\mathrm{E-P}}$, matches the experimental RCMC capture radius, $R_{\mathrm{RCMC}}$.

**Intuitive interpretation:** Why does the cancellation fail here? Recall the "cloud" analogy from the small-radius case.

- **Small radius case:** The volumes are so small that the cloud density inside them is effectively constant. Measuring a larger volume just means capturing more volume at that same density.

- **Finite radius case:** The volumes are now large enough to capture the *curvature* of the cloud (the density drops off when moving away from the center).

Importantly, the "shape" of this curvature depends on the polymer size $\sigma_i$ (genomic distance). A short genomic distance creates a sharp peak; a longer genomic distance creates a broad, gentle hill. If $R_{\mathrm{E-P}}$ and $R_{\mathrm{RCMC}}$ are different sizes, they sample different parts of this curve. For a sharp peak (short distance), the difference between sampling the tip ($R_{\mathrm{E-P}}$) and the shoulder ($R_{\mathrm{RCMC}}$) is drastic. For a broad hill (long distance), the difference is subtle. Because the relative proportion of the cloud captured changes depending on its shape (the genomic distance), the ratio $F/f$ is no longer constant across E-P configurations. The only way to get a constant ratio is if the two volumes are identical in size ($R_{\mathrm{E-P}} \approx R_{\mathrm{RCMC}}$), ensuring they always sample the exact same portion of the curve regardless of its shape.

**Summary for Part 1:** We have examined the baseline behavior for simple polymers (without specific loops) in two regimes:

1. **Small contact radii ($R_{\mathrm{RCMC}} \ll \sigma$):** The Taylor expansion shows that the condition-dependent term $p(\mathbf{0}|c_i)$ cancels out. In this limit, linearity is a trivial result expected regardless of the difference between the functional interaction radius $R_{\mathrm{E-P}}$ and experimental RCMC capture radius $R_{\mathrm{RCMC}}$.

2. **Finite contact radii:** When the capture radius is comparable to the polymer fluctuation size, the cancellation is no longer exact due to the curvature of the probability distribution. In this more general regime, observing a linear relationship across constructs with varying genomic distances ($c_i$) strictly implies that the functional and experimental radii must be similar ($R_{\mathrm{E-P}} \approx R_{\mathrm{RCMC}}$).

Thus, under the null model of a simple polymer, linearity is either a trivial consequence of geometry (if radii are small) or a specific consequence of matching radii (if radii are large). Given estimations of RCMC capture radius $R_{\mathrm{RCMC}}$ ~42 nm by prior work [60] and this work, as well as estimation of functional radius $R_{\mathrm{E-P}}$ both point to the direction of small contact radii ($R_{\mathrm{RCMC}} \ll \sigma$), we will focus on the case of small contact radii and explore additional factors that might break the linearity below.

#### 12.3 Part 2: Specific case for chromatin scaling

While Part 1 establishes that linearity might be trivial for simple polymers when the radius is small, we must determine the explicit form of the "background" polymer state to contrast it with the "stabilized" state in Part 3 below. Here, we apply the logic from Part 1 to a chromatin fiber following a general power-law scaling relationship.

##### 12.3.1 Relationship between spatial and genomic distance

We define the relationship between the genomic separation $s$ (in base pairs) and the mean squared spatial distance $\langle R^2 \rangle$ using a power law with a general scaling exponent $\nu$:

$$\langle R^2(s) \rangle = b^2 s^{2\nu} \tag{166}$$

where $b$ is a pre-factor related to the Kuhn length. The exponent $\nu$ varies by model: $\nu = 1/2$ for an Ideal Chain (random walk), while $\nu \approx 1/3$ for a Fractal Globule (a crumbled, unknotted polymer often used to model chromatin [120]).

##### 12.3.2 The probability density function (PDF)

We assume the separation vector $\boldsymbol{r}$ between the enhancer and promoter follows a Gaussian distribution, scaled by the variance $\langle R^2 \rangle$. This is the standard propagator for polymer physics [121]:

$$G(\boldsymbol{r}; s) = \left( \frac{3}{2\pi \langle R^2 \rangle} \right)^{3/2} \exp\left( -\frac{3|\boldsymbol{r}|^2}{2\langle R^2 \rangle} \right) \tag{167}$$

##### 12.3.3 Cumulative probability of contact

To find the probability of interaction $P_{\text{interaction}}(s)$, we integrate this density over the spherical volume $V$ defined by the radius $r_i$:

$$P_{\text{interaction}}(s) = \int_0^{r_i} \left( \frac{3}{2\pi \langle R^2 \rangle} \right)^{3/2} \exp\left( -\frac{3r^2}{2\langle R^2 \rangle} \right) 4\pi r^2 \, dr \tag{168}$$

##### 12.3.4 The scaling approximation

Consistent with the Taylor expansion in Part 1, we consider the regime of distal interactions where the mean spatial separation is much larger than the radius of the spherical volume ($\sqrt{\langle R^2 \rangle} \gg r_i$). In this limit, the argument of the exponential is small ($r_i^2/\langle R^2 \rangle \approx 0$), and the term $\exp\left(-\frac{3r^2}{2\langle R^2 \rangle}\right) \approx 1$. The integral simplifies to the density at the origin multiplied by the sphere volume:

$$P_{\text{interaction}}(s) \approx \underbrace{\left(\frac{3}{2\pi\langle R^2 \rangle}\right)^{3/2}}_{\text{Density at } r=0} \times \underbrace{\frac{4}{3}\pi r_i^3}_{\text{Sphere volume}} \tag{169}$$

Now, substituting the general scaling $\langle R^2 \rangle = b^2 s^{2\nu}$:

$$P_{\text{interaction}}(s) \approx \frac{4}{3}\pi r_i^3 \left(\frac{3}{2\pi b^2 s^{2\nu}}\right)^{3/2} = \frac{4}{3}\pi r_i^3 \left(\frac{3}{2\pi b^2}\right)^{3/2} (s^{2\nu})^{-3/2} \tag{170}$$

Simplifying the exponent $(-3/2 \cdot 2\nu = -3\nu)$, we arrive at the scaling law for the background polymer:

$$\boxed{P_{\text{background}}(s, r_i) \propto r_i^3 \cdot s^{-3\nu}} \tag{171}$$

This confirms that for the background chromatin motion, contact probability scales cubically with radius ($r_i^3$) and follows a power law with genomic distance.

#### 12.4 Part 3: The case with transient bonds (stickiness)

The assumption of a purely broad, continuous distance distribution fails if the E-P pair is able to "stick" together for a sustained period. This behavior is characteristic of specific biological regulation, such as:

- **CTCF-CTCF interactions:** Cohesin actively stabilizes loops, holding anchors in close proximity.
- **E-P affinity ('stickiness'):** Inherent biochemical affinity (e.g., formation of microcompartments due to block co-polymer microphase separation of the anchors [122]) keeps the pair in proximity above and beyond the background polymer random collisions.

##### 12.4.1 Mixture model formulation

We approximate this scenario as a superposition of two populations. Let $\alpha(c_i)$ be the fraction of time (or fraction of the population) the E-P pair $c_i$ spends in the **stabilized state**. The total probability density is a weighted sum:

$$P_{\text{total}}(r) = (1 - \alpha(c_i))P_{\text{background}}(r) + \alpha(c_i)P_{\text{stabilized}}(r) \tag{172}$$

Here, $P_{\text{background}}(r)$ is the broad polymer distribution derived in Part 2, while $P_{\text{stabilized}}(r)$ represents tight spatial confinement due to stabilized interactions (e.g. CTCF-CTCF loops and E-P affinity) above background polymer interactions.

##### 12.4.2 Approximation of the stabilized state

While the stabilized E-P complex is often idealized as a point source using a Dirac delta function, physically, the stabilized state has a finite spatial extent defined by the size of the crosslinked protein complex, $\sigma_{\text{bond}}$. We model the probability density of this state as a narrow Gaussian distribution with variance $\sigma_{\text{bond}}^2$, which is significantly smaller than the polymer variance ($\sigma_{\text{bond}}^2 \ll \langle R^2 \rangle$). The cumulative probability of capturing this bound state within a radius $r$ is given by the error function:

$$p_c^{\text{stabilized}}(r) = \int_{|\boldsymbol{r}| \leq r} P_{\text{stabilized}}(\boldsymbol{r})\, d\boldsymbol{r} \approx \operatorname{erf}\left(\frac{r}{\sigma_{\text{bond}}\sqrt{2}}\right) \tag{173}$$

We assume the regime where the measurement radius is larger than the physical size of the protein complex ($R_{\text{RCMC}}, R_{\text{E-P}} \gg \sigma_{\text{bond}}$). In this limit, the error function approaches unity. For example, if $r = R_{\text{RCMC}} \approx$ 42 nm [60] and the complex size $\sigma_{\text{bond}} \approx$ 10 nm, $\operatorname{erf}(4) \approx 1$. Thus, the expression simplifies to:

$$p_c^{\text{total}}(c_i, R_{\text{RCMC}}) \approx \alpha(c_i) + (1 - \alpha(c_i))p_c^{\text{background}}(R_{\text{RCMC}}) \tag{174}$$

##### 12.4.3 Volume scaling interpretation

Equation (21) reveals two competing scaling laws within the same measurement:

1. **Background polymer component:** Scales with volume ($\propto r^3$). As the capture radius increases, we capture more of the broad polymer cloud. This term is weighted by the background fraction $(1-\alpha)$.
2. **Stabilized component:** Scales as a constant ($\approx 1$). Unlike the background polymer, the stabilized state is spatially confined within a small volume ($\sigma_{\text{bond}}$). Once the capture radius exceeds this physical size ($r \gg \sigma_{\text{bond}}$), the entire stabilized population is captured, making this term independent of further increases in volume. This term is weighted by the fraction $\alpha$.

##### 12.4.4 The breakdown of linearity

We now re-evaluate the ratio of Expression ($F$) to Contact ($f$) using different radii ($R_{\mathrm{E-P}}$ and $R_{\mathrm{RCMC}}$). Substituting the full expression from Eq. (15):

$$\frac{F(c_i)}{f(c_i)} \approx \frac{C_{\mathrm{E-P}}}{C_{\mathrm{RCMC}}} \cdot \frac{\alpha(c_i) + (1-\alpha(c_i))p_c^{\text{background}}(R_{\mathrm{E-P}})}{\alpha(c_i) + (1-\alpha(c_i))p_c^{\text{background}}(R_{\mathrm{RCMC}})} \tag{175}$$

To see the dependency on $\alpha(c_i)$ more clearly, we can rearrange the terms to group by $\alpha$:

$$\frac{F(c_i)}{f(c_i)} \approx \frac{C_{\mathrm{E-P}}}{C_{\mathrm{RCMC}}} \cdot \frac{p_c^{\text{background}}(R_{\mathrm{E-P}}) + \alpha(c_i)[1 - p_c^{\text{background}}(R_{\mathrm{E-P}})]}{p_c^{\text{background}}(R_{\mathrm{RCMC}}) + \alpha(c_i)[1 - p_c^{\text{background}}(R_{\mathrm{RCMC}})]} \tag{176}$$

**Analysis of the limit $R_{\mathrm{RCMC}} \to 0$:** If the experimental radius is very small while the functional radius is larger ($R_{\mathrm{RCMC}} \ll R_{\mathrm{E-P}}$), the polymer probability in the denominator vanishes ($p_c(R_{\mathrm{RCMC}}) \to 0$) while the numerator remains significant. The ratio becomes:

$$\lim_{R_{\mathrm{RCMC}} \to 0} \frac{F}{f} \approx \frac{C_{\mathrm{E-P}}}{C_{\mathrm{RCMC}}} \cdot \frac{p_c^{\text{background}}(R_{\mathrm{E-P}}) + \alpha(c_i)[1 - p_c^{\text{background}}(R_{\mathrm{E-P}})]}{\alpha(c_i)} \tag{177}$$

As $\alpha(c_i)$ approaches 0 (E-P without CTCF and stickiness), this ratio diverges; as $\alpha(c_i)$ increases, the ratio drops. Because $\alpha(c_i)$ varies biologically between constructs, the data points will not lie on a single linear trend.

**Restoring linearity:** We ask under what condition the ratio in Equation (176) becomes a constant $K$ independent of the loop fraction $\alpha(c_i)$. By inspection, if $p_c^{\text{background}}(R_{\mathrm{E-P}}) = p_c^{\text{background}}(R_{\mathrm{RCMC}})$, the numerator and denominator of the second fraction are identical, and the ratio is exactly 1 regardless of $\alpha$. Mathematically, for the ratio $\frac{A+\alpha(1-A)}{B+\alpha(1-B)}$ to be constant for all $\alpha$, we strictly require $A = B$. Thus, linearity is restored if and only if:

$$p_c^{\text{background}}(R_{\mathrm{E-P}}) \approx p_c^{\text{background}}(R_{\mathrm{RCMC}}) \implies R_{\mathrm{E-P}} \approx R_{\mathrm{RCMC}} \tag{178}$$

**Intuitive interpretation:** Consider the stabilized state as the "signal" and the random polymer collisions as the "background noise."

- **The stabilized state (signal):** This is a point source. Once the radius is non-zero, you catch 100% of the signal. The "signal" amount is constant regardless of volume size.
- **The background state (background):** This is a diffuse cloud. A larger volume captures significantly more background noise than a smaller volume. The "noise" amount scales with volume ($r^3$).

If expression is determined by a larger functional interaction radius ($R_{\mathrm{E-P}}$) but RCMC with a smaller experimental capture radius ($R_{\mathrm{RCMC}}$), the expression measurement will have a much lower "signal-to-noise" ratio than the RCMC measurement, where we define 'signal' as the stabilized interactions and 'noise' as the random background collisions inherent to the polymer. Importantly, different E-P configurations have different intrinsic signal strengths ($\alpha$). A construct with a strong loop (high signal) will be less affected by the background noise than a construct with no loop (pure background collisions). Because the impact of the mismatched volumes varies depending on the signal strength of the E-P configuration, the relationship is no longer linear. The only way to ensure that all E-P configurations—whether strong loops or weak loops—fall on the same line is if both measurements have the same "signal-to-noise" ratio. This requires the volumes to be the same size ($R_{\mathrm{E-P}} \approx R_{\mathrm{RCMC}}$).

#### 12.5 Visualization of volume ratios

The plot below visualizes the theoretical divergence derived in Part 3. We simulate a dataset containing both "background" polymer-only constructs and "stabilized" constructs with varying loop fractions ($\alpha = 5\%, 20\%$).

**Explanation of the plot:**

- We now plot the capture probability at the RCMC capture radius $P(R_{\text{RCMC}})$ on the x-axis vs. the interaction probability at the functional interaction radius $P(R_{\text{E-P}})$ on the y-axis.
- The **dashed lines** represent the linear relationship expected for simple polymers.
- **Scenario 1 (blue series, mismatch):** Here, $R_{\text{RCMC}} > R_{\text{E-P}}$, specifically $(R_{\text{RCMC}}/R_{\text{E-P}})^3 = 3$. The simple polymers follow the line $x = 3y$ (or $y = x/3$). The looped constructs (solid blue dots) are shifted off this line because adding the constant $\alpha$ to both axes does not preserve this slope.
- **Scenario 2 (red series, match):** Here, $R_{\text{RCMC}} = R_{\text{E-P}}$. The simple polymers follow the line $y = x$. The looped constructs (solid red dots) also fall exactly on this line, because adding the constant $\alpha$ to both axes preserves the identity slope.
- **Scenario 3 (teal series, mismatch):** Here $R_{\text{RCMC}} < R_{\text{E-P}}$, specifically $(R_{\text{RCMC}}/R_{\text{E-P}})^3 = 1/3$. The simple polymers follow the line $x = y/3$ (or $y = 3x$). The looped constructs are shifted off this line.

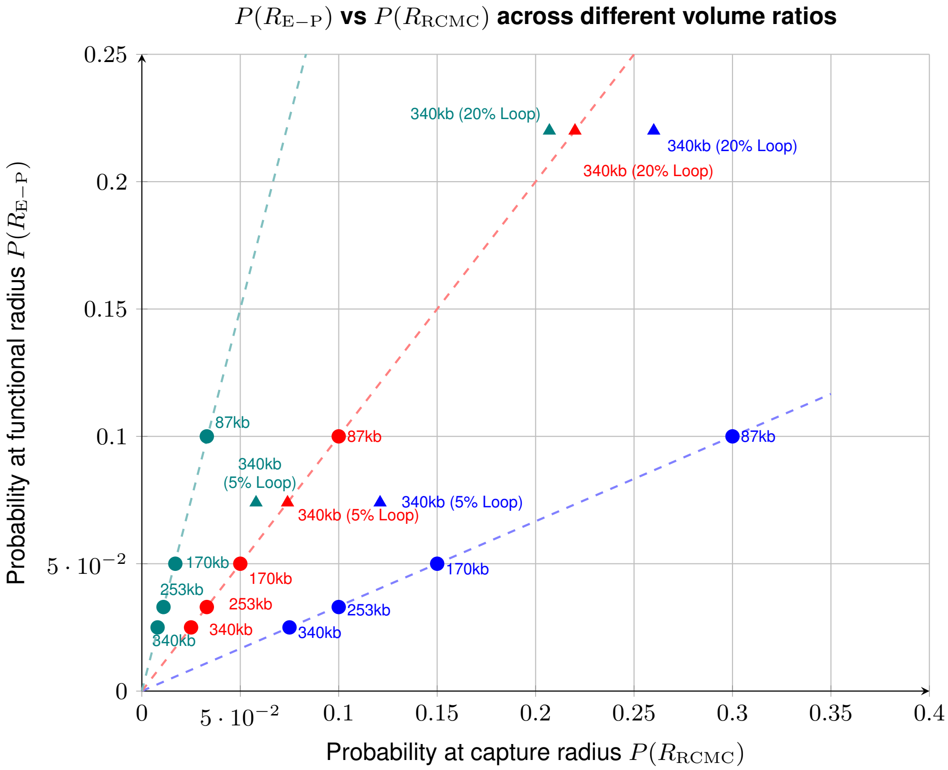

**Supplementary Note 2 Fig. 1: Impact of loop fraction ($\alpha$) on linearity**. Conditions with non-zero stabilized fractions ($\alpha > 0$, triangles) only align with conditions with zero stabilized fractions (circles on polymer trend line) when $R_{\text{RCMC}} = R_{\text{E-P}}$ (red series).

#### 12.6 Part 4: Analytical validation of the $R_{\text{E-P}}$ estimate via CTCF-facilitation

While 3D polymer simulations enables the estimation of $R_{\text{E-P}}$ by accounting for excluded volume, stickiness between the E-P pair, and the exact 3D chromosome organization, the ~20-fold expression increase conferred by convergent CBSs (**Fig. 2E-G**) can also be modeled using a direct analytical approximation. This provides an orthogonal validation of our simulation-derived estimate of $R_{\text{E-P}} \approx 29$ nm.

For the 339 kb E-P pair without CBSs (339noC), the probability of the E-P pair residing within a sphere with an interaction radius $R_{\text{E-P}}$ can be evaluated using the small-radius scaling approximation for a 3D Gaussian chain derived earlier in Part 2. By evaluating the Gaussian probability density function (Equation 167) over a small spherical capture volume of radius $R_{\mathrm{E-P}}$, we substitute $R_{\mathrm{E-P}}$ as the capture volume radius and $R^2_{\mathrm{RMS}}$ for $\langle R^2 \rangle$ to yield:

$$P_{339\mathrm{noC}}(r < R_{\mathrm{E-P}}) \approx \underbrace{\frac{4\pi}{3} R^3_{\mathrm{E-P}}}_{\text{Sphere volume}} \times \underbrace{\left(\frac{3}{2\pi R^2_{\mathrm{RMS}}}\right)^{3/2}}_{\text{Density at origin}} = \sqrt{\frac{54}{\pi}} \left(\frac{R_{\mathrm{E-P}}}{R_{\mathrm{RMS}}}\right)^3 \tag{179}$$

where $R_{\mathrm{RMS}}$ is the root-mean-square 3D distance of the E-P pair. From our Bayesian MSD fitting (**Table S6**), $R_{\mathrm{RMS}} \approx 280$ nm for 339noC.

For the 339 kb E-P pair with convergent CBSs (339CECP), transcription can be approximated to scale with looping probability of the CTCF-CTCF loop. Based on our BILD analysis (**Fig. S13E**), this looping probability is $P_{\mathrm{loop}} \approx 8.9\%$. When looped, the E-P pair is effectively held in contact by cohesin, making the functional interaction probability $P_{339\mathrm{CECP}}(r < R_{\mathrm{E-P}}) \approx P_{\mathrm{loop}}$.

Given the experimentally observed 19.5-fold increase in BFP expression, and assuming expression is proportional to the functional interaction probability, we obtain the following ratio:

$$\frac{P_{339\mathrm{CECP}}}{P_{339\mathrm{noC}}} = \frac{P_{\mathrm{loop}}}{\sqrt{\frac{54}{\pi}} \left(\frac{R_{\mathrm{E-P}}}{R_{\mathrm{RMS}}}\right)^3} = 19.5 \tag{180}$$

Substituting $P_{\mathrm{loop}} = 0.089$ (**Fig. S13E**) and $R_{\mathrm{RMS}} = 280$ nm (**Table S6**) and solving for $R_{\mathrm{E-P}}$:

$$R_{\mathrm{E-P}} = 280 \times \left(\frac{0.089}{19.5 \times \sqrt{\frac{54}{\pi}}}\right)^{1/3} \approx 280 \times (0.0011)^{1/3} \approx 28.9 \text{ nm} \tag{181}$$

This simple analytical calculation yields an estimate of 28.9 nm, further supporting the $\sim$29 nm $R_{\mathrm{E-P}}$ estimate derived independently from comparing BFP expression and interaction probability calculated from polymer simulations (**Fig. 2F**).

#### 12.7 Conclusion

This mathematical derivation provides a theoretical framework for interpreting the linear relationship between flow cytometry expression data and RCMC contact data.

- **Ambiguity in simple polymers:** We demonstrated that for simple polymers (Part 1), linearity could be a trivial consequence of the volume scaling and does not provide information about the functional radius, provided the radius is small. Practically, the appearance of a 'dot' on RCMC maps show stabilized interactions above the background interactions expected for a simple polymer. We observe dots in our RCMC maps both with and without CTCF sites, which rules out the simple polymer model for the experimental E-P pairs studied here.
- **Resolution via loops:** However, the introduction of transient bonds or CTCF-CTCF loops (Part 3) breaks this trivial scaling. The "stabilized loop" state introduces a constant term ($\alpha$) that does not scale with volume. Here, this corresponds to the appearance of a dot on our RCMC maps indicative of a stabilized loop.
- **Final deduction:** Consequently, if linearity is observed in a dataset that includes E-P configurations with different CTCF site arrangement/occupancy or just different genomic distance with the same CTCF arrangement (which all have different $\alpha$), it is mathematically impossible for the capture radius $R_{\mathrm{RCMC}}$ to differ significantly from the functional radius $R_{\mathrm{E-P}}$.

Therefore, the robust linearity observed in our experimental data implies that the **functional interaction radius** for transcription is approximately the same as the **capture radius** of the RCMC assay, i.e. that $R_{\mathrm{E-P}} \approx R_{\mathrm{RCMC}}$. Since the experimental RCMC capture radius has been estimated to be $\approx 42$nm, the observed linearity rules out action-at-a-distance models and argues that functional E-P interactions occur on the "contact-like" scale.

#### 13 Supplementary Note 3: MS2 intensity correction

##### 13.1 Observation and hypothesis of MS2 signal decay

Across cell lines and conditions, we observed a decay of the ensemble mean MS2 intensity over the imaging time course (**Supplementary Note 3 Fig. 1**). The magnitude of such decay is much larger than the less than 20% decay due to photobleaching, which has already been corrected for.

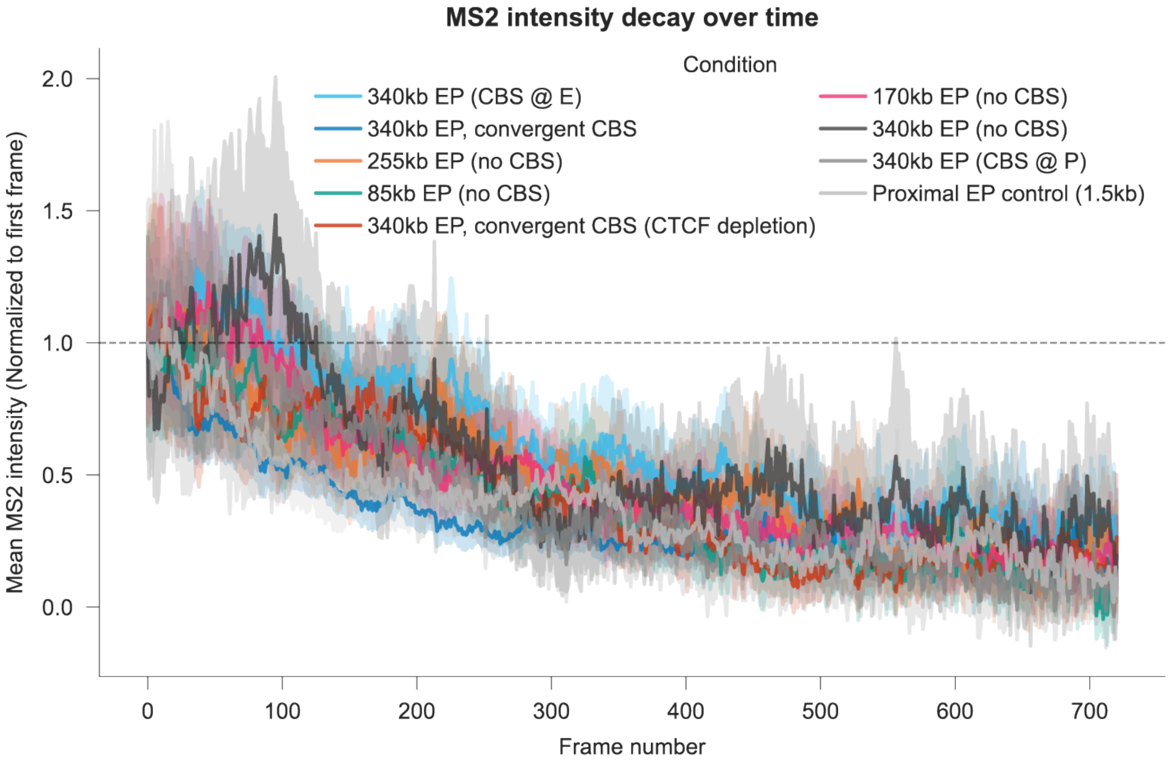

**Supplementary Note 3 Fig. 1:** Normalized MS2 intensity decay over time across cell lines and conditions.

We hypothesized that this post-photobleaching correction MS2 signal decay results from the sequestration of the finite nuclear pool of MCP-2x-Halo proteins by transcribed RNAs. As transcription proceeds, newly synthesized RNAs continuously sequester MCP-2x-Halo, progressively leading to a reduced pool of MCP-2x-Halo available to bind nascent RNAs at the synthetic promoter and consequently the reduced fluorescence per RNA molecule transcribed later in time.

##### 13.2 Evidence supporting the MCP-2x-Halo sequestration hypothesis

The sequestration of MCP protein by RNAs is a well-known phenomenon [37–39]. Consistent with prior observations, we frequently observe cells accumulating bright, distinct nuclear foci in the MCP channel of our live-cell movies (**Supplementary Note 3 Fig. 2**). We can readily distinguish foci and ”floating RNAs” from true nascent RNAs due to our fluorescent DNA promoter label (via ANCH3) as well as the apparent diffusion coefficient (DNA-associated nascent RNAs diffuse very slow and subdiffusively; ”floating RNAs” diffuse much more rapidly and less subdiffusively).

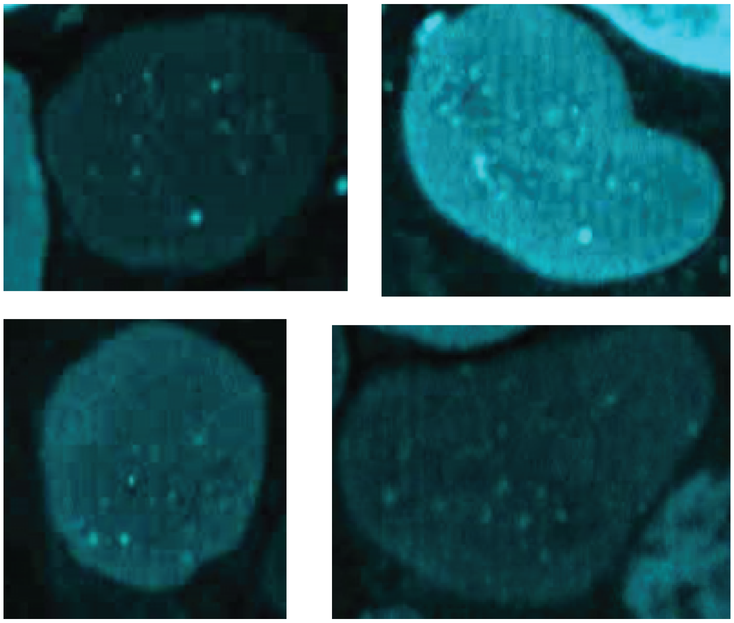

**Supplementary Note 3 Fig. 2:** Representative live-cell images demonstrating the accumulation of bright MCP-2x-Halo nuclear foci.

To mathematically disentangle potentially other purely imaging time-dependent effects from the effect of MCP-2x-Halo sequestration, we performed a track-level partial correlation analysis. We found that MS2 signal decay is significantly more correlated with cumulative transcription history than with imaging time. By holding one variable constant, we confirmed that signal attenuation is fundamentally driven by cumulative transcriptional history rather than the imaging time (**Supplementary Note 3 Fig. 3**).

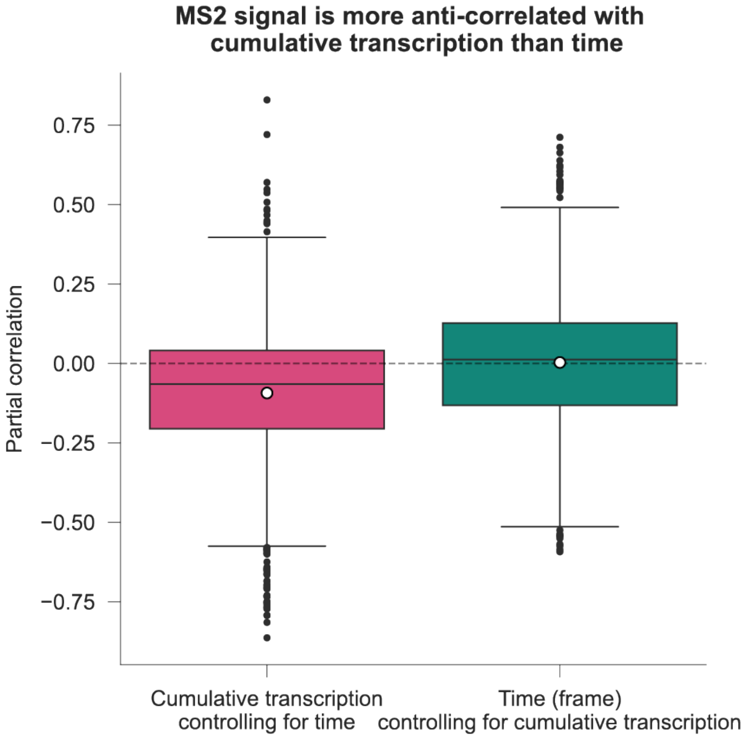

**Supplementary Note 3 Fig. 3:** Track-level partial correlation demonstrating that cumulative transcription history is a vastly superior predictor of MS2 decay than chronological time.

To further visualize and validate that MS2 decay is fundamentally driven by cumulative transcription history rather than imaging time, we compared orthogonal grouping strategies. When the tracks were chronologically grouped by the acquisition frame number, and the ensemble mean MS2 intensity of each frame was mapped against its corresponding ensemble mean cumulative transcription history, the data resolved into a highly correlated decay envelope (**Supplementary Note 3 Fig. 4A**). Conversely, when grouping the exact same dataset by their cumulative transcription history and plotting the mean intensity of those bins against their corresponding chronological frame—the relationship dissolved into an uncoordinated scatter (**Supplementary Note 3 Fig. 4B**). If the MS2 decay was primarily a function of time, synchronizing the observations by their cumulative transcription and plotting them against time would yield a smooth decay curve, whereas mapping chronological frames against unsynchronized transcriptional histories would produce scatter. The fact that the observed data exhibit the exact opposite behavior mathematically isolates cumulative transcriptional history as the driver of MS2 signal decay.

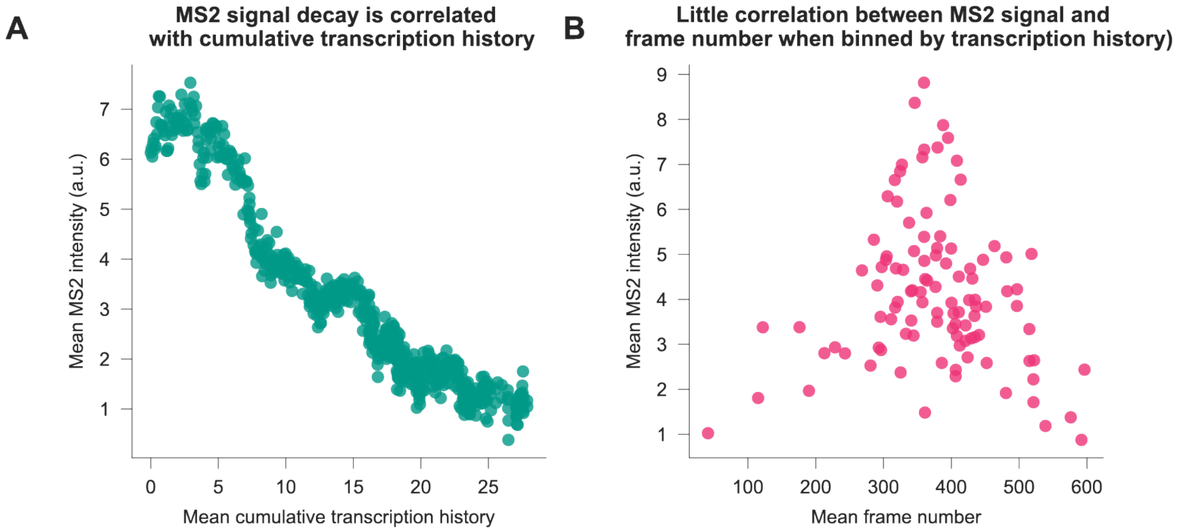

**Supplementary Note 3 Fig. 4:** Track-level partial correlation demonstrating that cumulative transcription history is a vastly superior predictor of MS2 decay than chronological time.

Another corollary of the MCP-2x-Halo sequestration hypothesis is that one allele's transcription activity will reduce the MS2 intensity of the other allele since they share the same pool of MCP-2x-Halo. For nuclei where tracks were available for both alleles, we observed that the MS2 signal is significantly lower when the other allele is actively transcribing, consistent with the expectation (**Supplementary Note 3 Fig. 5**).

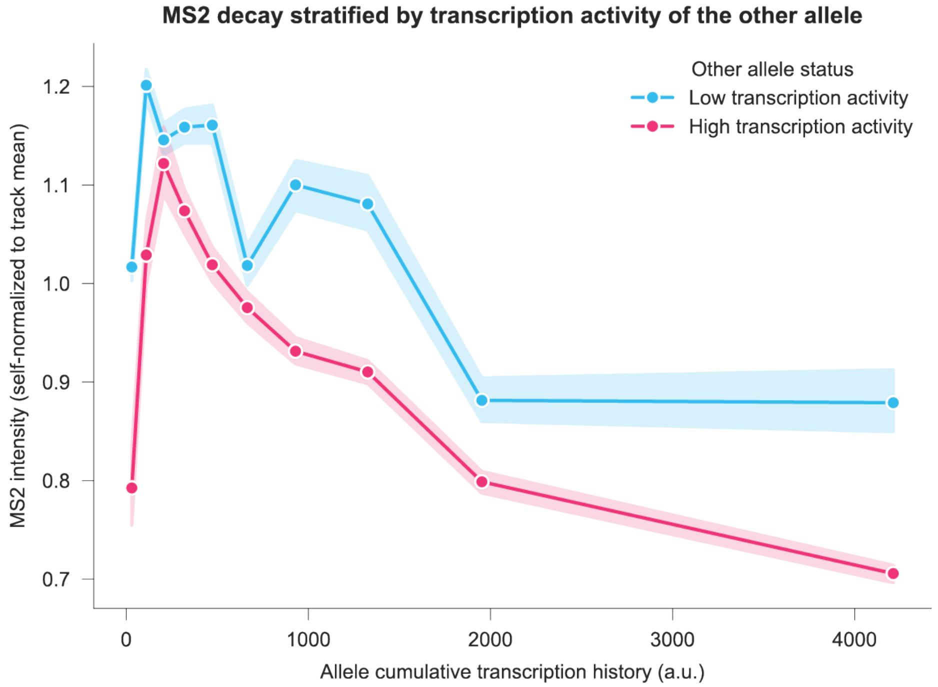

**Supplementary Note 3 Fig. 5:** Effect of transcription from the other allele on the MS2 intensity of the allele of interest. Shaded area around the curves corresponds to 95% confidence interval.

#### 13.3 Empirical correction of MS2 signal decay

To accurately quantify transcription dynamics over extended imaging periods, we implemented a data-driven, empirical correction pipeline to reverse the time-dependent decay of the MS2 signal caused by sequestration of MCP-2x-Halo by RNAs. This approach extracts the macroscopic decay envelope directly from the ensemble behavior of the experimental data, ensuring the correction is intrinsically calibrated to the specific imaging conditions, fluorophore kinetics, and heterogeneity of each experimental group. The code for running MS2 decay correction is available at https://github.com/ahansenlab/synEP/tree/main/MS2DecayCorrection.

##### 13.3.1 Resolving biological heterogeneity: fractional transcription history and archetype stratification

To successfully extract a representative decay envelope, the correction model must account for the cell-to-cell heterogeneity inherent to live-cell imaging—specifically, variations in the temporal synchronization of transcription and the absolute capacity of the MCP-2x-Halo pool. We resolve these intertwined challenges prior to ensemble averaging through a two-step normalization and stratification approach. To build intuition for the necessity of this approach, consider the mathematical challenges of averaging live-cell imaging tracks from an unsynchronized, heterogeneous cell population. We face two distinct ambiguities:

- **Temporal ambiguity (the synchronization problem):** Imagine two highly active alleles from different nuclei. Allele A began transcribing well before the imaging acquisition started, while Allele B just activated transcription at frame 1. By frame 50, Allele A has already severely depleted its available MCP-2x-Halo pool, causing its newly transcribed RNAs to appear dimmer. Allele B's pool is still full, so its bursts appear brighter. If we average these tracks using absolute chronological time, we erroneously fuse a late-stage decayed signal with an early-stage baseline.

- **Amplitude ambiguity (the scale problem):** Now imagine two alleles that activate at the exact same time, but transcribe at different levels due to the stochastic nature of transcription. Allele C is highly productive and generates massive bursts, while Allele D generates a slow, weak stream of transcripts. Even if their timelines are perfectly synchronized, averaging their absolute intensities (the y-axis) will mash a high-amplitude signal and a low-amplitude signal into an artificial, flattened middle-ground that represents neither decay curves accurately.

First, to resolve the **temporal ambiguity** of unsynchronized transcription lifecycles, the decay axis (x-axis) is normalized into a fractional transcription history ($H_{frac}$). For a given track, the cumulative transcription output at frame $t$ is calculated and normalized to the total transcription output observed for that specific track by the end of the acquisition, bounding the progression axis between 0 and 1. Mapping the decay envelope to $H_{frac}$ synchronizes tracks by their relative stage of MCP-2x-Halo depletion. This aligns Allele A and Allele B not by the arbitrary clock of the microscope, but by their progression through their respective observed transcription capacity. Furthermore, because MCP-2x-Halo expression level varies widely from cell to cell, normalizing the transcription history to the cell's own observed maximum transcription output allows $H_{frac}$ to serve as a mathematical proxy for the relative exhaustion of that specific cell's MCP-2x-Halo reservoir. Second, to resolve the **amplitude ambiguity** between high- and low-transcribing alleles, tracks must be stratified prior to averaging. While $H_{frac}$ synchronizes the timeline of MCP-2x-Halo depletion, it does not correct the y-axis disparity between the highly transcribing alleles and quiet alleles. Because distinct track populations initiate at vastly differing absolute MS2 intensities, global ensemble averaging of unstratified data would still create a flattened, unrepresentative decay envelope. To prevent this amplitude artifact, tracks are stratified into distinct transcriptional archetypes using K-Means clustering based on their total transcription output. To avoid arbitrary thresholding, the optimal number of clusters ($k$) was determined dynamically for each condition. The pipeline iteratively tested $k = 1$ through $k = 5$ and automatically selected the $k$ that maximized the Silhouette Score, ensuring that tracks were objectively grouped into mathematically natural, well-separated subpopulations. Notably, across all experimental conditions analyzed, this dynamic optimization consistently converged on $k = 2$, effectively bifurcating tracks into distinct "low transcribers" and "high transcribers" archetypes prior to binning (**Supplementary Note 3 Fig. 6A**).

##### 13.3.2 Quantile binning and ensemble averaging

Within each identified K-Means archetype, the MS2 intensity of each frame across tracks were pooled to reconstruct the macroscopic signal decay along the $H_{frac}$ axis. Because live-cell locus tracking is subject to progressive dropout (where many trajectories terminate early due to replications etc.), traditional uniform binning (e.g., binning strictly by intervals of 0.1 $H_{frac}$) can result in statistically noisy tails. To resolve this, the data was grouped using dynamic quantile binning. The continuous $H_{frac}$ axis was sliced into discrete bins (up to a maximum of 20), adjusting the bin edges such that every bin contained an approximately equal number of empirical observations. For each bin $b$, the ensemble average of the $H_{frac}$ position and the mean MS2 intensity ($\langle I_b \rangle$) were calculated, producing an evenly weighted scatter of the decay envelope (**Supplementary Note 3 Fig. 6A**).

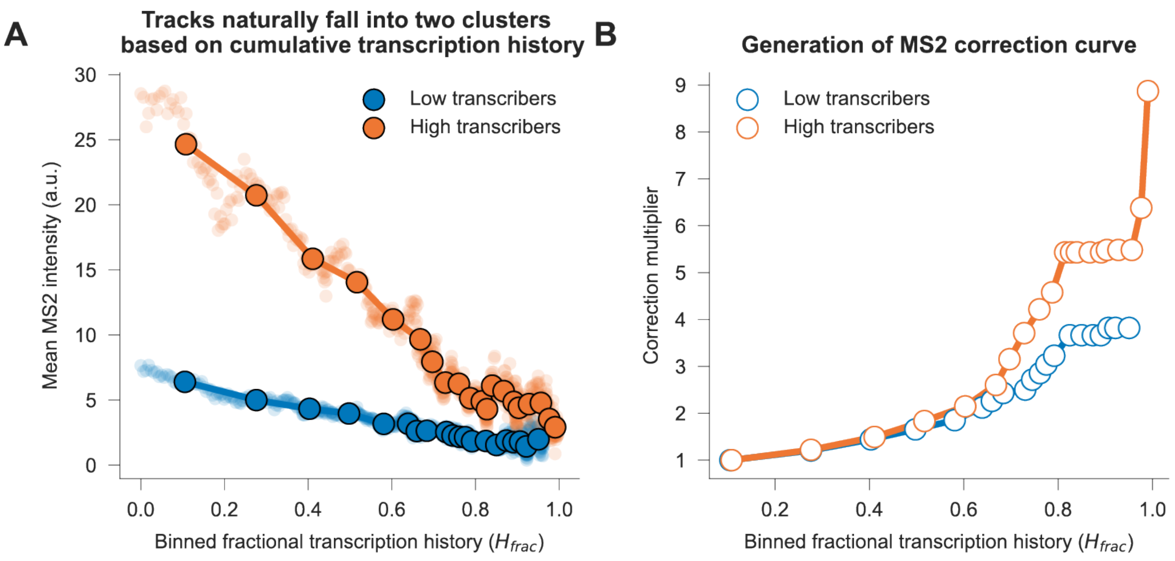

**Supplementary Note 3 Fig. 6:** Derivation of the MS2 signal correction curve.

##### 13.3.3 Multiplier derivation and linear interpolation

To isolate the fractional signal loss, the discrete decay envelope was then converted into a correction map. We observed that early stages of transcription sometimes exhibit a brief "rising phase" before the MS2 signal decay kicks in. To avoid artificially over-correcting these early, pre-peak fluctuations, the reference intensity ($I_{peak}$) was defined as the maximum smoothed intensity of the envelope, and the peak index ($b_{peak}$) was identified. The raw correction multiplier for each bin, $M_{raw}(b)$, was calculated as the ratio of the peak intensity to the bin's smoothed mean intensity ($\langle I_b \rangle$). For all bins preceding the peak ($b \le b_{peak}$), the multiplier was strictly fixed at 1.0 to ignore the initial rising phase:

$$M_{raw}(b) = \begin{cases} 1.0, & \text{if } b \le b_{peak} \\ \frac{I_{peak}}{\langle I_b \rangle}, & \text{if } b > b_{peak} \end{cases}$$

Furthermore, because MCP-2x-Halo depletion is presumably a one-way physical process driven by sequestration, the theoretical correction factor required to restore the signal should continuously increase. To prevent local statistical noise from causing non-biological dips in the correction curve, we enforced strict monotonicity. The final discrete multiplier map $M(b)$ ensures that the correction factor is only allowed to increase or plateau as fractional history increases:

$$M(b) = \max_{j \le b} M_{raw}(j)$$

To apply this discrete bin-level correction map back to each track, we utilized linear interpolation. For any given data point in a track, the pipeline looked up its current $H_{frac}$ value, located the two nearest quantile bin averages, and interpolated the precise multiplier $M(t)$. The raw intensity of that specific frame was then multiplied by the interpolated factor:

$$I_{corr}(t) = I(t) \cdot M(t)$$

This approach dynamically reverses the MS2 signal decay (**Supplementary Note 3 Fig. 7**).

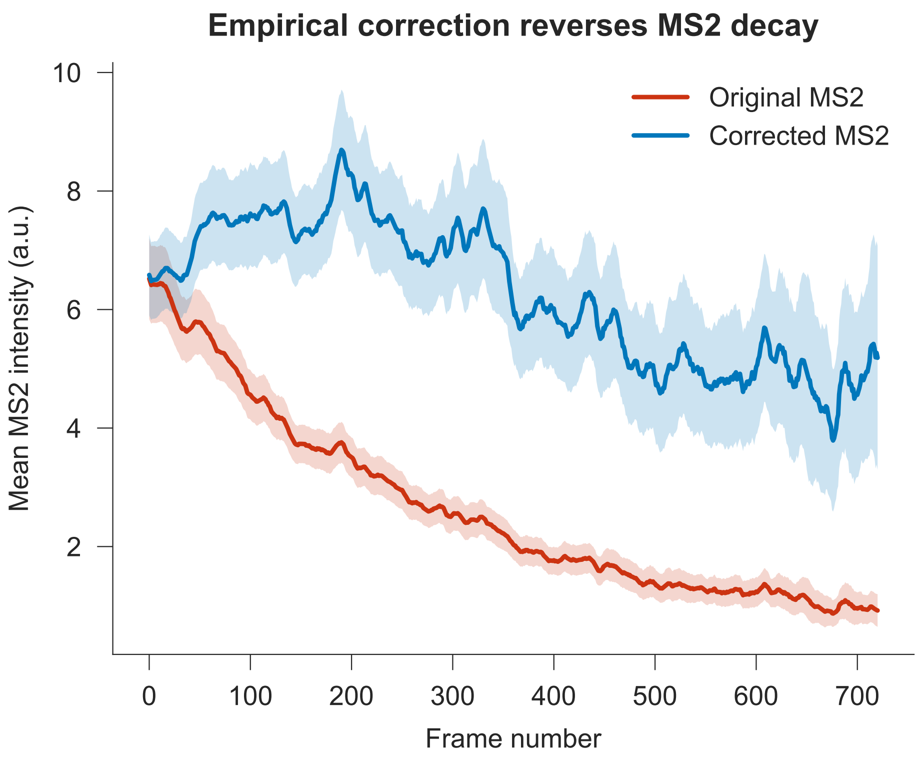

**Supplementary Note 3 Fig. 7:** Empirical correction reverses MS2 signal decay. Shaded area around the curves corresponds to 95% confidence interval.

##### 13.3.4 Application of the empirically corrected MS2 signal in this study

To ensure analytical rigor, for all aggregate quantification in this study, such as calculating the ensemble mean or median MS2 intensities, we utilized the fully corrected MS2 signal to accurately account for the observed MS2 signal decay.

However, because the multiplicative correction process inevitably amplifies baseline measurement noise alongside the true burst signal, analysis sensitive to local signal-to-noise ratios (such as change-point detection and statistical inference for burst calling) were performed using the raw uncorrected MS2 signal. Finally, to transparently present the data exactly as captured, all representative single-trajectory plots of MS2 signals display the raw uncorrected MS2 intensities.

#### 14 Supplementary Figures

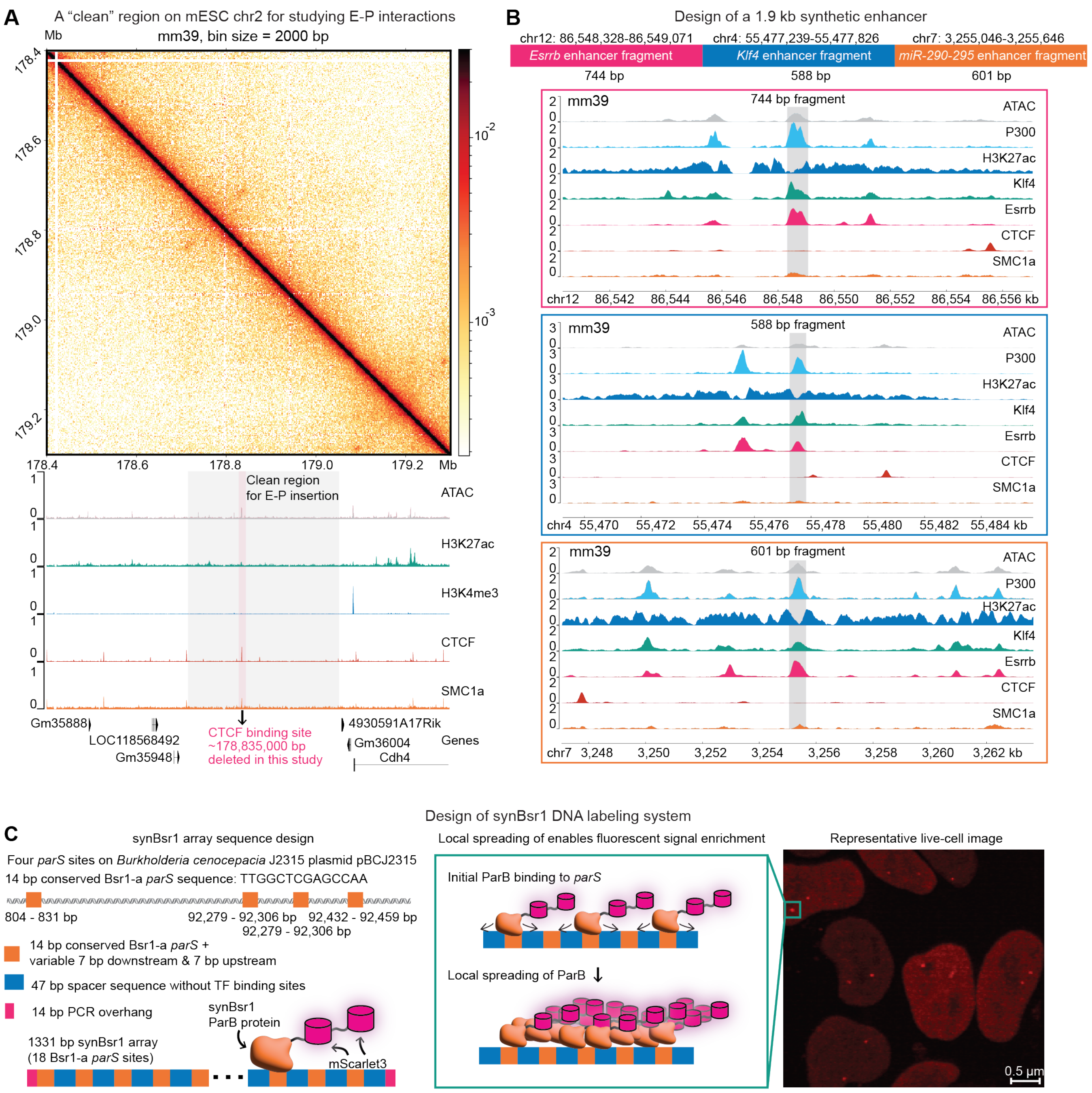

**Fig. S1: Genomic region selection and the bottom-up design of the synthetic enhancer and the synBsr1 labeling system.** (**A**) Micro-C contact map [12] and epigenetic tracks of the selected 'clean' region on mouse embryonic stem cell (mESC) chromosome 2 used for synthetic E-P insertion. The ChIP-seq tracks are from previously published datasets: ATAC (GSE90892) [123], H3K27ac (GSE90893) [123], H3K4me3 (GSE90893) [123], CTCF (GSM5668637) [8], SMC1a (GSM5668638) [8]. The y-axis limit for all tracks have been adjusted to accommodate the maximum signal observed across all tracks within the visualized window. (**B**) Design of the 1.9 kb synthetic enhancer, combining fragments from the *Esrrb*, *Klf4*, and *miR-290-295* enhancers [28]. Shown at the bottom are the ChIP-seq tracks of these constituent fragments in their endogenous loci from previously published datasets: P300 (GSE90893), Klf4 (GSE90893), Esrrb (GSE90893) [123]. ATAC, H3K27ac, CTCF, and SMC1a tracks are from the same sources in (A). The y-axis limit for all tracks within a panel have been adjusted to accommodate the maximum signal observed across all tracks within the visualized window. (**C**) Schematic detailing the sequence design and local spreading mechanism of the synBsr1 DNA labeling system. A representative live-cell image demonstrating fluorescent signal enrichment at the labeled synthetic enhancer is shown on the right.

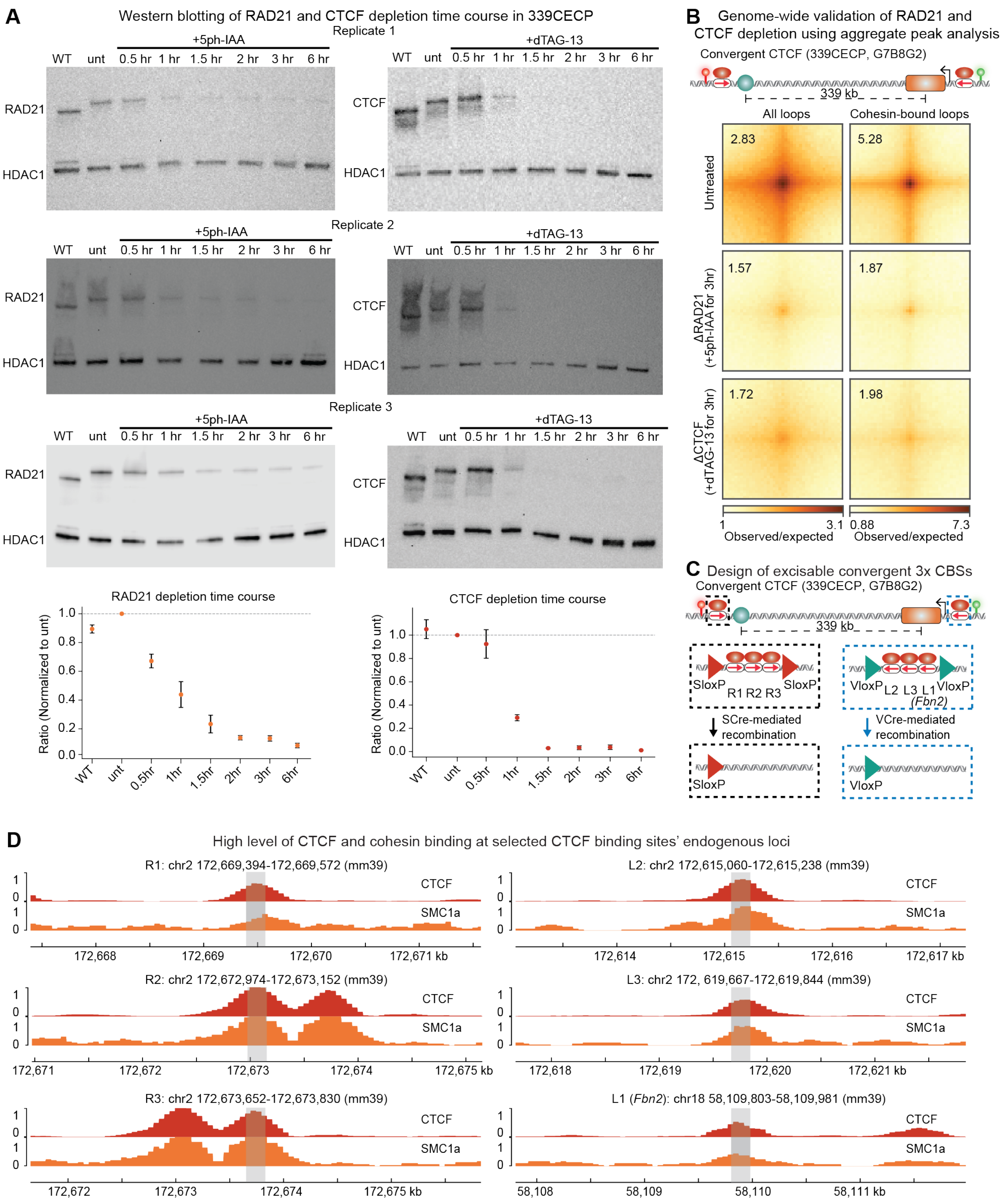

**Fig. S2: Acute depletion of RAD21 and CTCF, and the design of convergent 3x CBSs.** (**A**) Western blot time-course analyses quantifying the degradation of RAD21 and CTCF following treatment with 5ph-IAA and dTAG-13, respectively, across three biological replicates in the 339CECP cell line. (**B**) Genome-wide validation of RAD21 and CTCF depletion using aggregate peak analysis (APA) on Micro-C data, validating the loss of structural loops following respective degron-targeted treatments.(**C**) Design of the convergent 3x CBSs used in this study. R1, R2, R3, L2, L3 were from [22], and L1 was from [8]. R1, R2, R3 near the synthetic enhancer are flanked by SloxP sites so that they can be deleted upon the expression of SCre recombinase [26]. L2, L3, *Fbn2* L1 near the synthetic promoter are flanked by VloxP sites so that they can be deleted upon the expression of VCre recombinase [26]. (**D**) The six selected CBSs had high level of CTCF and cohesin binding, as seen in the ChIP-seq tracks of CTCF (GSM3508478) and SMC1a (GSM3508477) [8]. The y-axis limit for all tracks within a panel are adjusted to accommodate the maximum signal observed across all tracks within the visualized window.

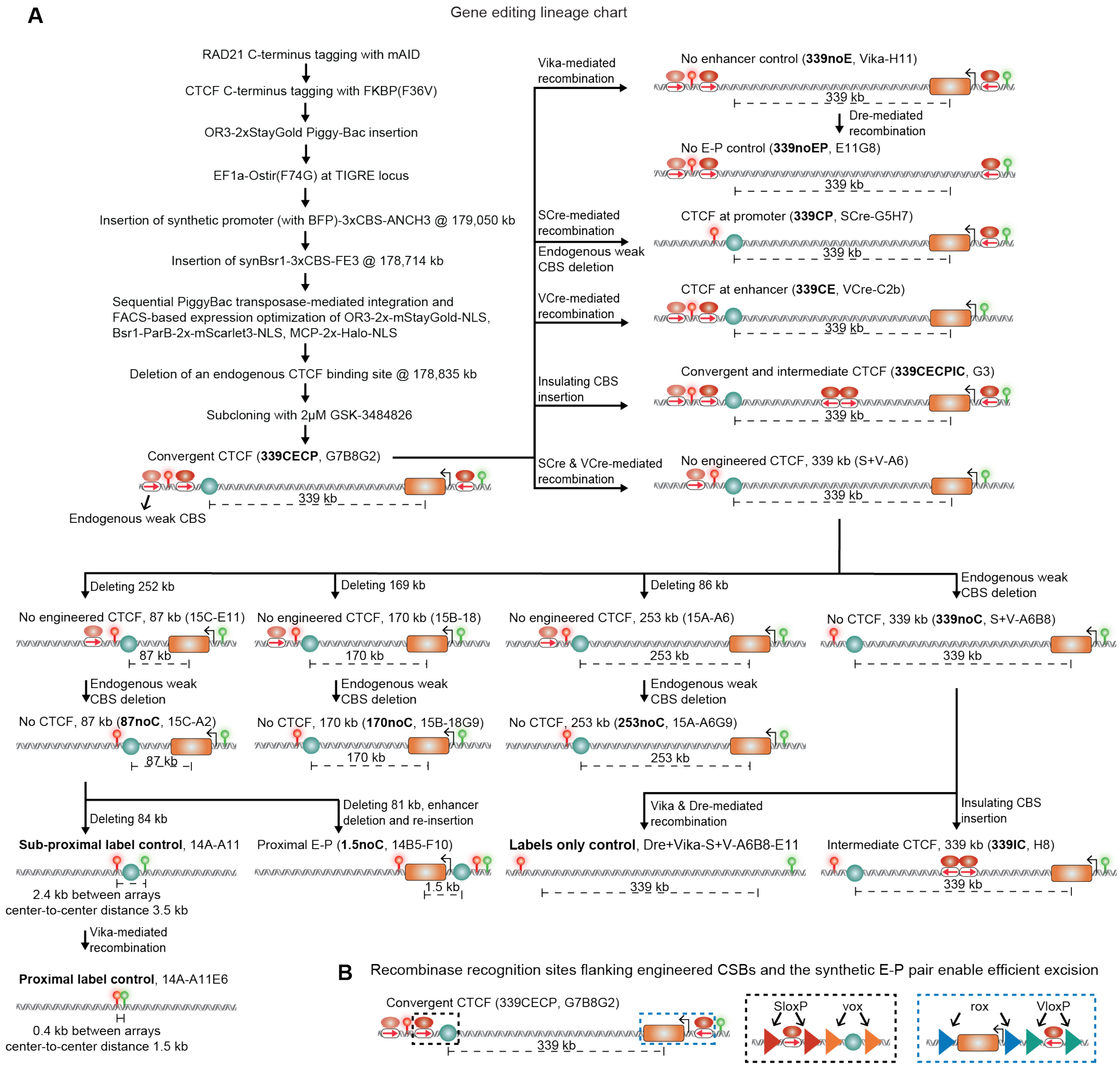

**Fig. S3: Gene editing lineage chart.** (**A**) A flowchart detailing the sequential genome editing steps—including CRISPR/Cas9-mediated insertions/deletions, degron tagging, and recombinase-mediated excisions (using Vika, Dre, SCre, and VCre)—used to generate the suite of cell lines with different E-P distances and CBS configurations, as well as the proximal label control cell line used in this study. (**B**) Schematic illustrating the orthogonal recombinase recognition sites flanking engineered CBSs and the synthetic E-P pair, which allows efficient excision of engineered CBSs, the synthetic enhancer, or the synthetic promoter.

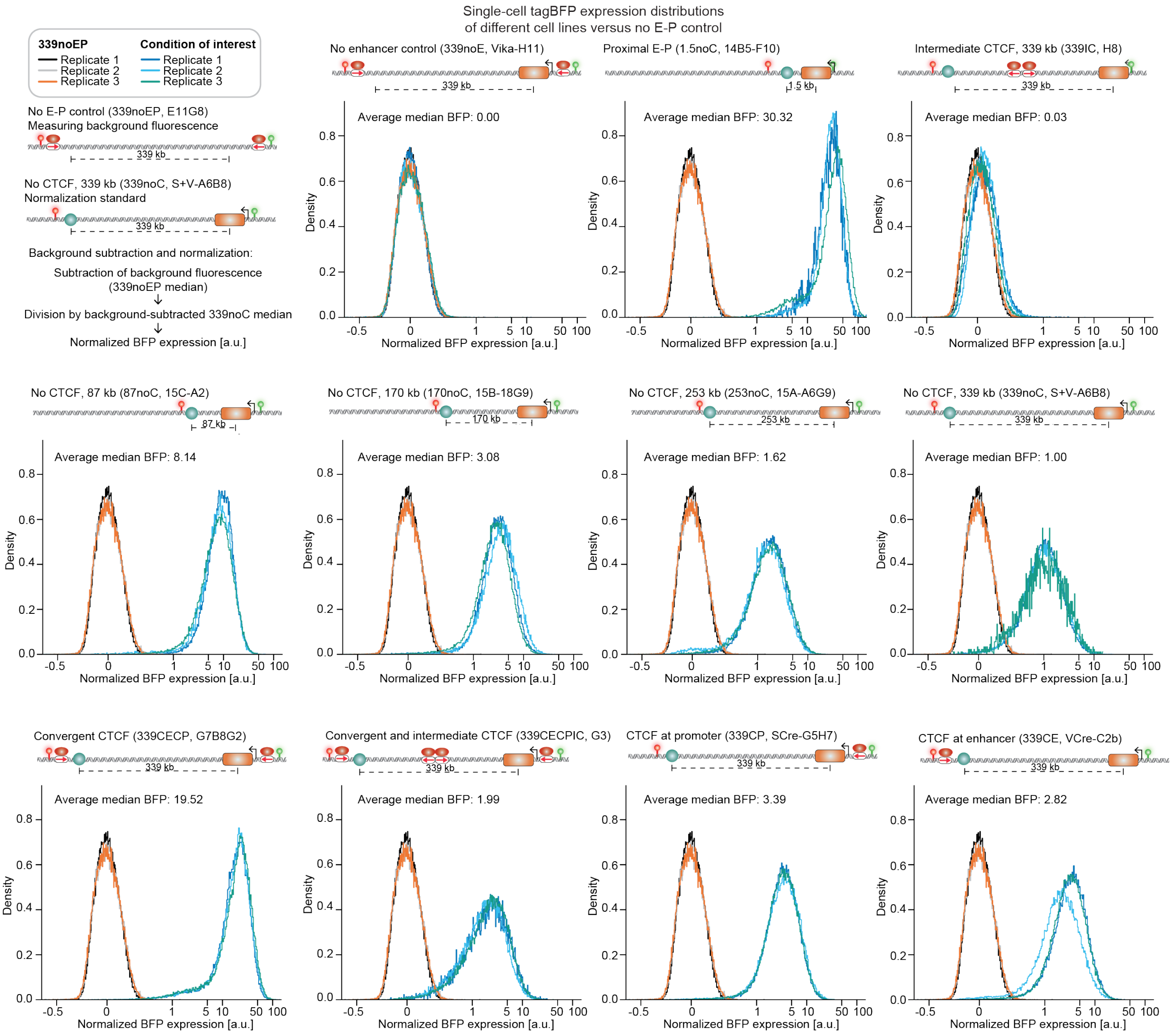

**Fig. S4: Flow cytometry quantification of tagBFP expression across cell lines.** Single-cell tagBFP expression distributions for the different engineered cell lines compared against the no E-P control (339noEP) measuring background fluorescence. The background subtraction and normalization strategy is shown on the top left. Briefly, for every single cell, the raw fluorescence was background-subtracted using the median of the 339noEP control population. This value was then divided by the background-subtracted median of the 339noC population (used as the normalization standard). To properly visualize single-cell populations containing near-zero and negative fluorescence values alongside highly expressing cells, the $x$-axis is plotted using a standard flow cytometry biexponential (inverse hyperbolic sine) transformation.

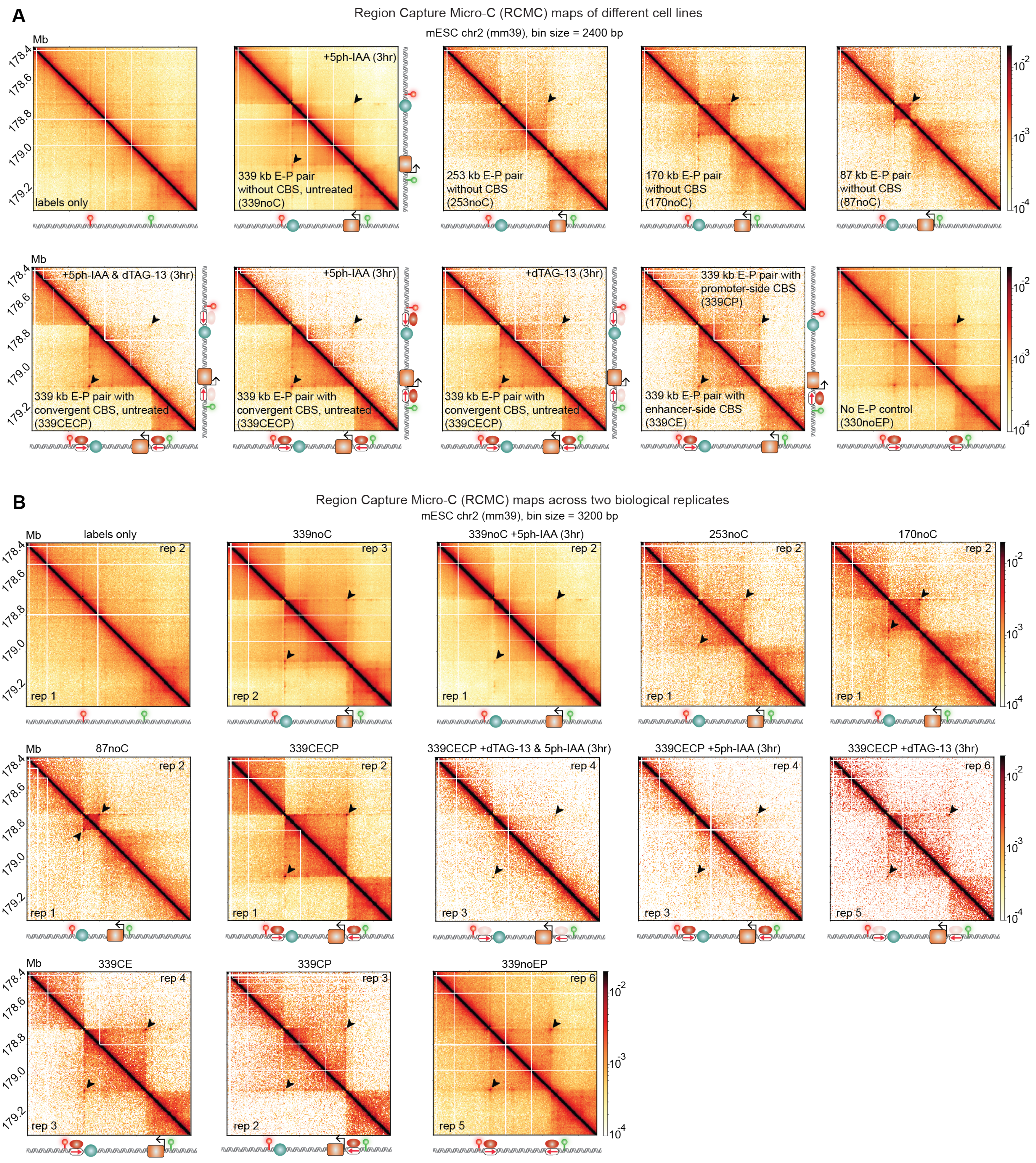

**Fig. S5: RCMC quantified E-P interaction strength.** (**A**) Region Capture Micro-C (RCMC) [56, 57] contact maps across the different engineered cell lines and degron-depletion conditions, highlighting focal interaction enrichments.(**B**) Comparison of contact maps between two biological replicates for different engineered cell lines and degron-depletion conditions.

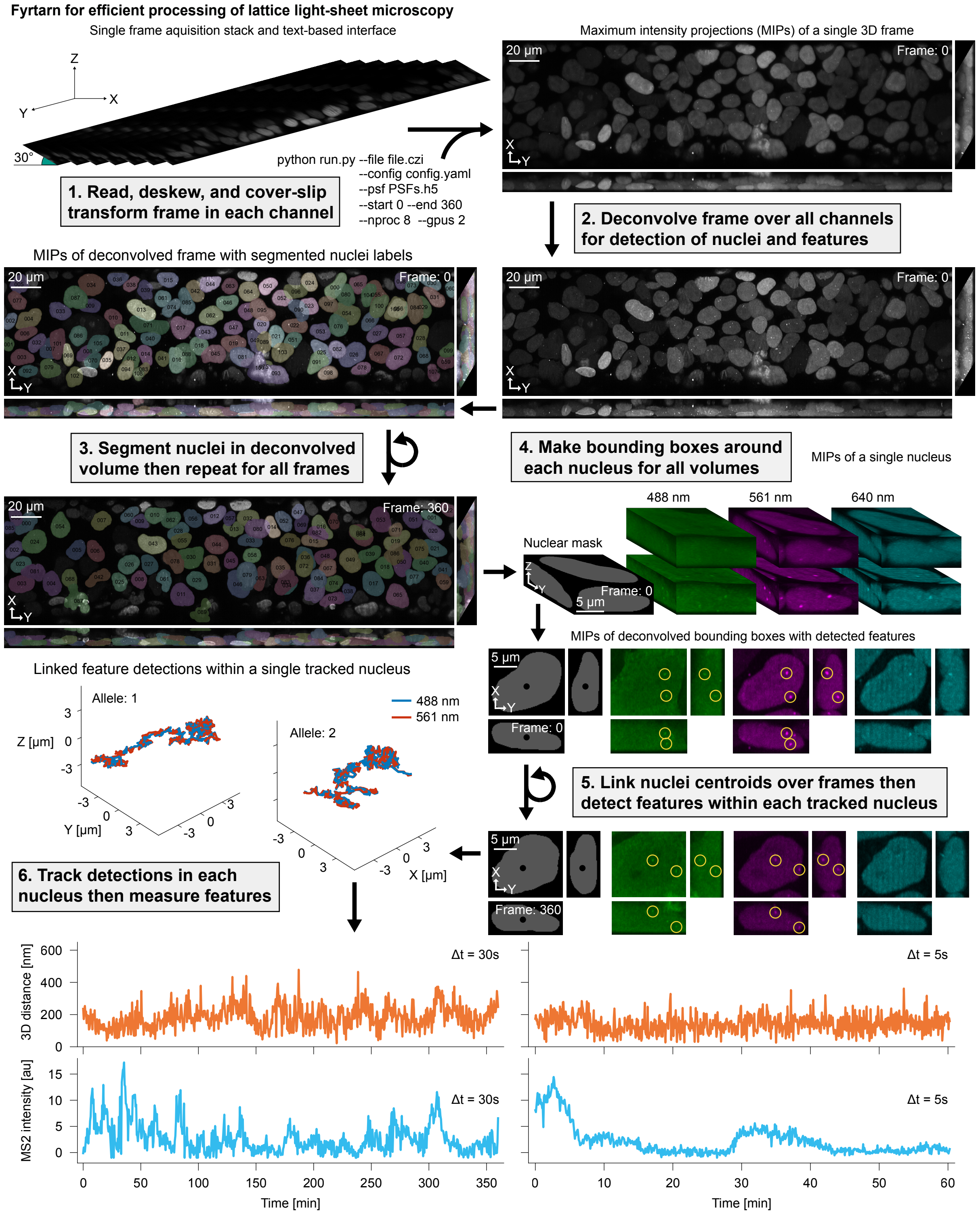

**Fig. S6: Fyrtarn for efficient quantification of live-cell dynamics in lattice light-sheet microscopy. 1.** Acquired stacks are read then deskewed with a cover-slip transformation to yield isotropic volumes. **2.** Deskewed volumes are deconvolved to facilitate nuclear segmentation and feature detection. **3-4.** Deconvolved nuclear-localized signal is first used to generate segmentation masks then deskewed volumes are subsetted to create bounding boxes around each nucleus. **5.** Detection, localization, and tracking of features within each centroid-tracked nucleus. **6.** Representative 3D distances and MS2 intensities trajectories after measuring tracked features.

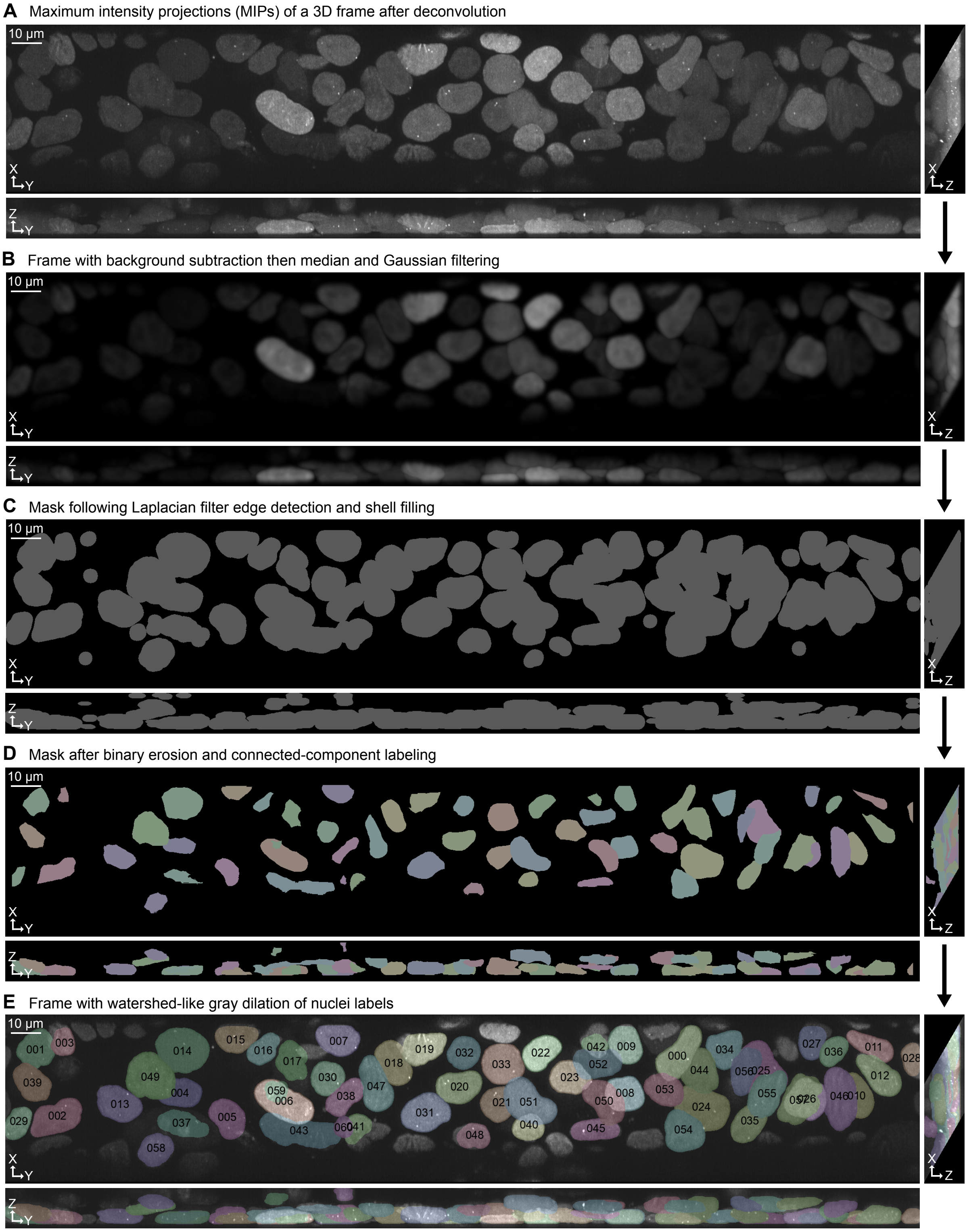

**Fig. S7: Overview of method used to segment nuclei. (A)** Representative 3D frame of nuclear-localized signal after deconvolution shown with maximum intensity projections (MIPs) along each axis. **(B)** Noise reduction with median and Gaussian filtering prior to edge detection. **(C)** Binary mask following flood-filling of Laplacian of Gaussian (LoG) zero-crossing detections. **(D)** Labeled nuclei seeds after morphological erosion of binary mask for watershed-like separation of nearby nuclei. **(E)** Morphological gray dilation of labeled nuclei masks overlaid on the input frame.

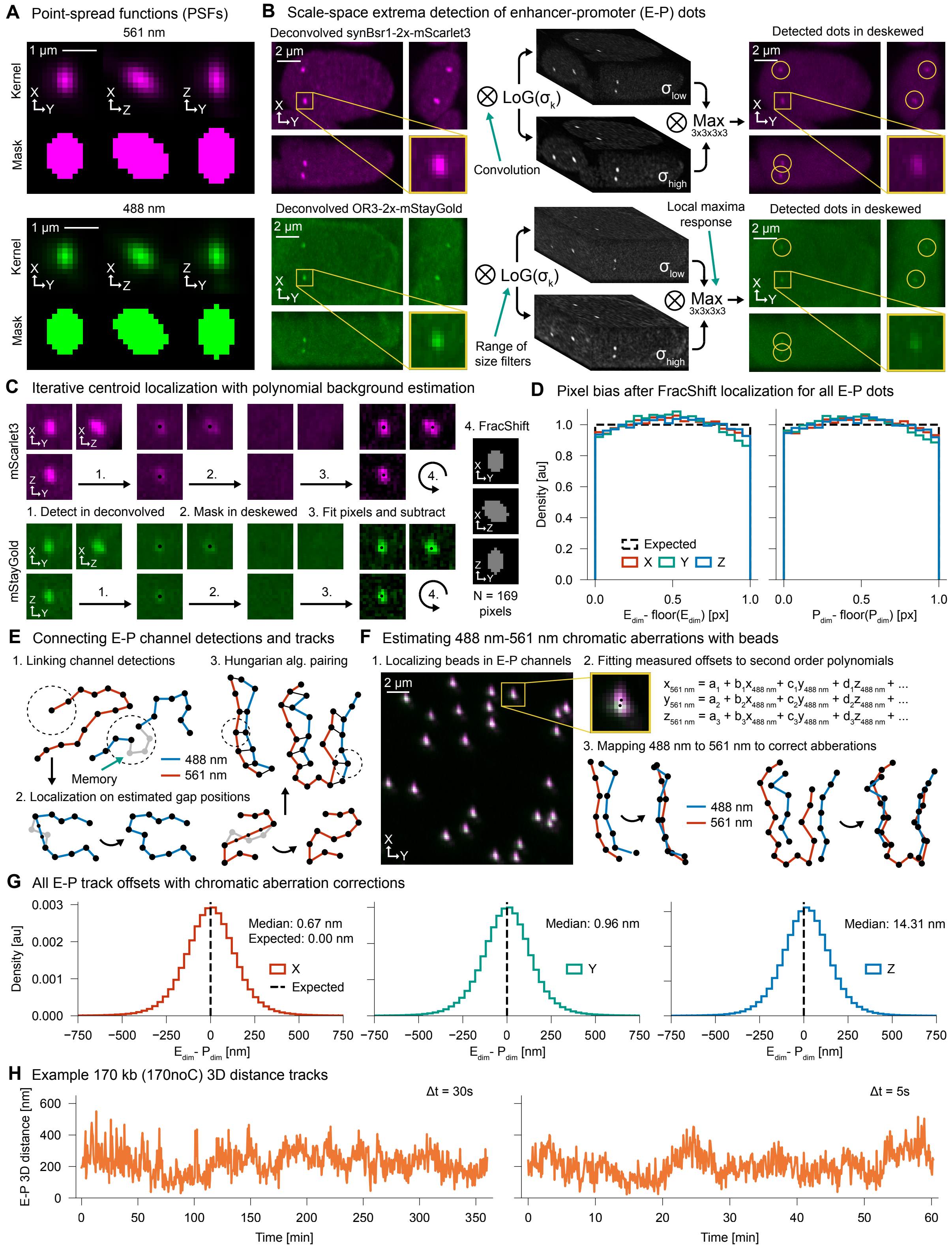

**Fig. S8: Localization and tracking of enhancer-promoter (E-P) locus pairs for measuring 3D distances. (A)** Kernels and signal masks for bead-measured 561 nm (enhancer) and 488 nm (promoter) point-spread functions (PSFs). **(B)** Schematic for normalized Laplacian of Gaussian detection over scale and space in deconvolved signal. **(C-D)** Background corrected FracShift localization of E-P dots in deskewed signal after detection with deconvolution and residual pixel bias computed for all localizations. **(E)** Schematic of E-P channel linking and track pairing after localizing positions. **(F-G)** Fitting of chromatic aberrations on bead localizations and evaluation of corrections with all E-P localizations. **(H)** Example 170kb cell line 3D distance trajectories from 30 sec (left) and 5 sec (right) time intervals.

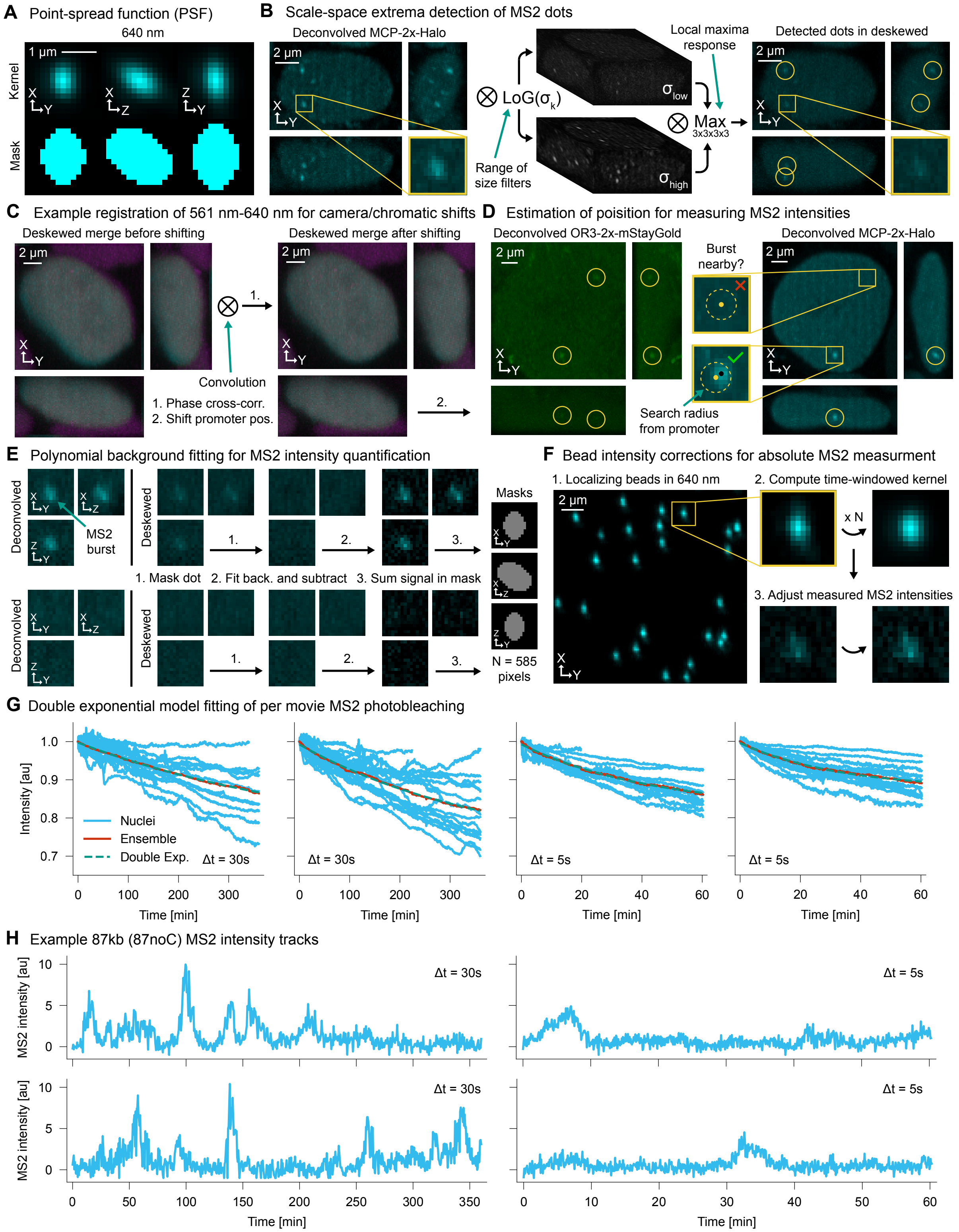

**Fig. S9: Quantification of MS2 intensities for measuring nascent transcription.** **(A)** Kernel and signal mask for bead-measured 640 nm (MS2) point-spread function (PSF). **(B)** Schematic for normalized Laplacian of Gaussian detection over scale and space in deconvolved signal. **(C)** Example registration of 561 nm and 640 nm channels using phase cross-correlation to estimate chromatic and camera shifts. **(D)** Method for localization of MS2 intensity measurement position based on detection of bursts and estimated promoter positions. **(E)** Intensity background correction procedure with second order polynomials. **(F)** Representative correction of MS2 intensities based on a two-week window of bead intensities. **(G)** Example photobleaching curves from 30 sec and 5 sec movies. A double-exponential model is fit to a curve averaged from tracked nuclei. **(H)** Example MS2 intensity tracks.

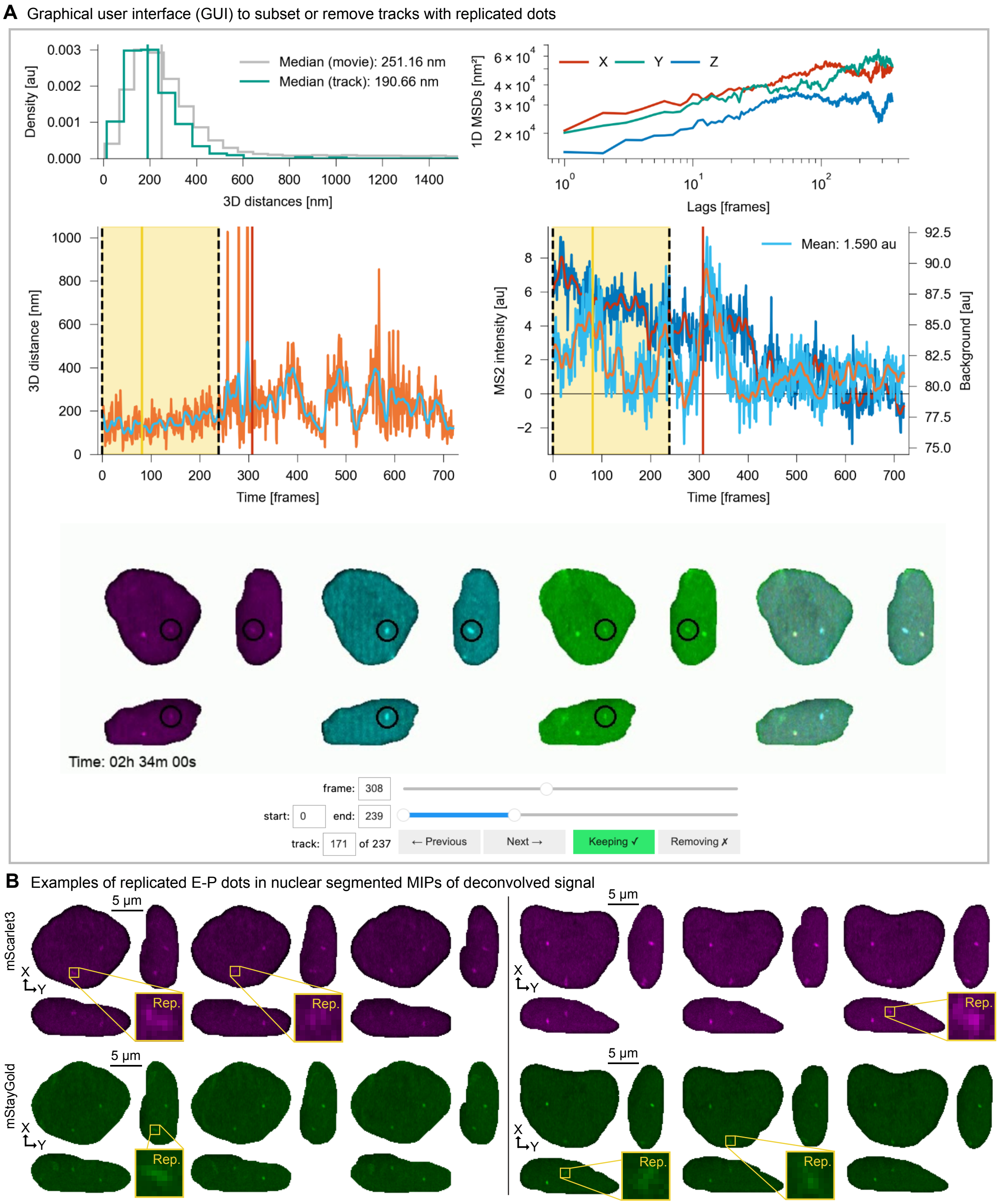

**Fig. S10: Overview of quality control (QC) to subset or remove tracks with replicated E-P loci.** **(A)** Jupyter-based graphical user interface (GUI) to view 3D distances and MS2 intensities. Sliders and buttons allow a user to subset or to remove tracks with replicated E-P loci. **(B)** Example replicated nuclei (left and right) with maximum intensity projections (MIPs) after nuclear segmentation and visualizations of sister chromatid dots.

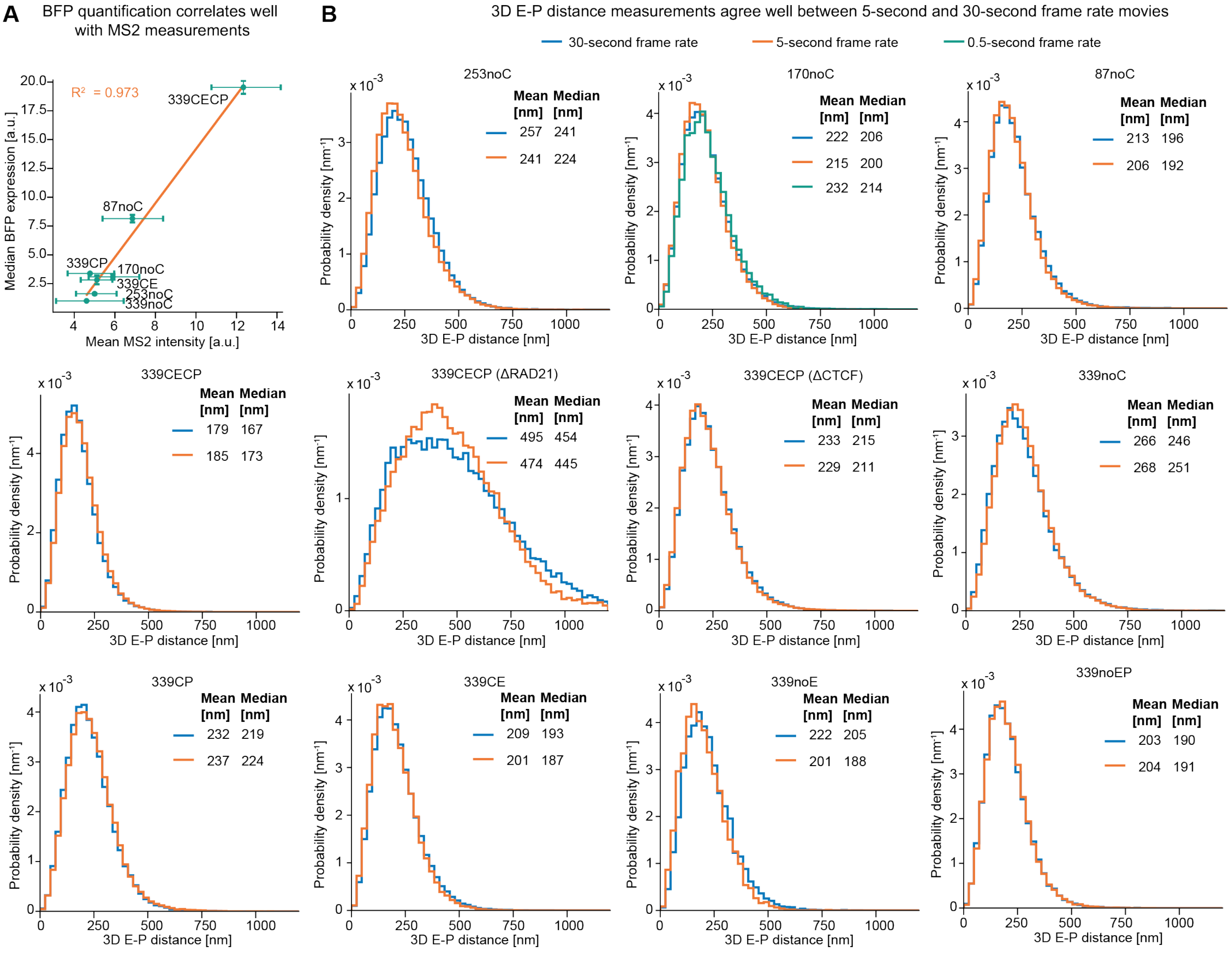

**Fig. S11: BFP-measured expression correlates well with MS2 intensity and 3D E-P distance distributions across cell lines and conditions are similar between 5-second and 30-second trajectory data.** (**A**) Quantification tagBFP expression via flow cytometry correlates well with MS2 intensity measured by live-cell imaging. (**B**) Probability density functions of 3D E-P distances demonstrating agreement between measurements derived from 0.5-second, 5-second, and 30-second frame rate live-cell movies across different cell lines.

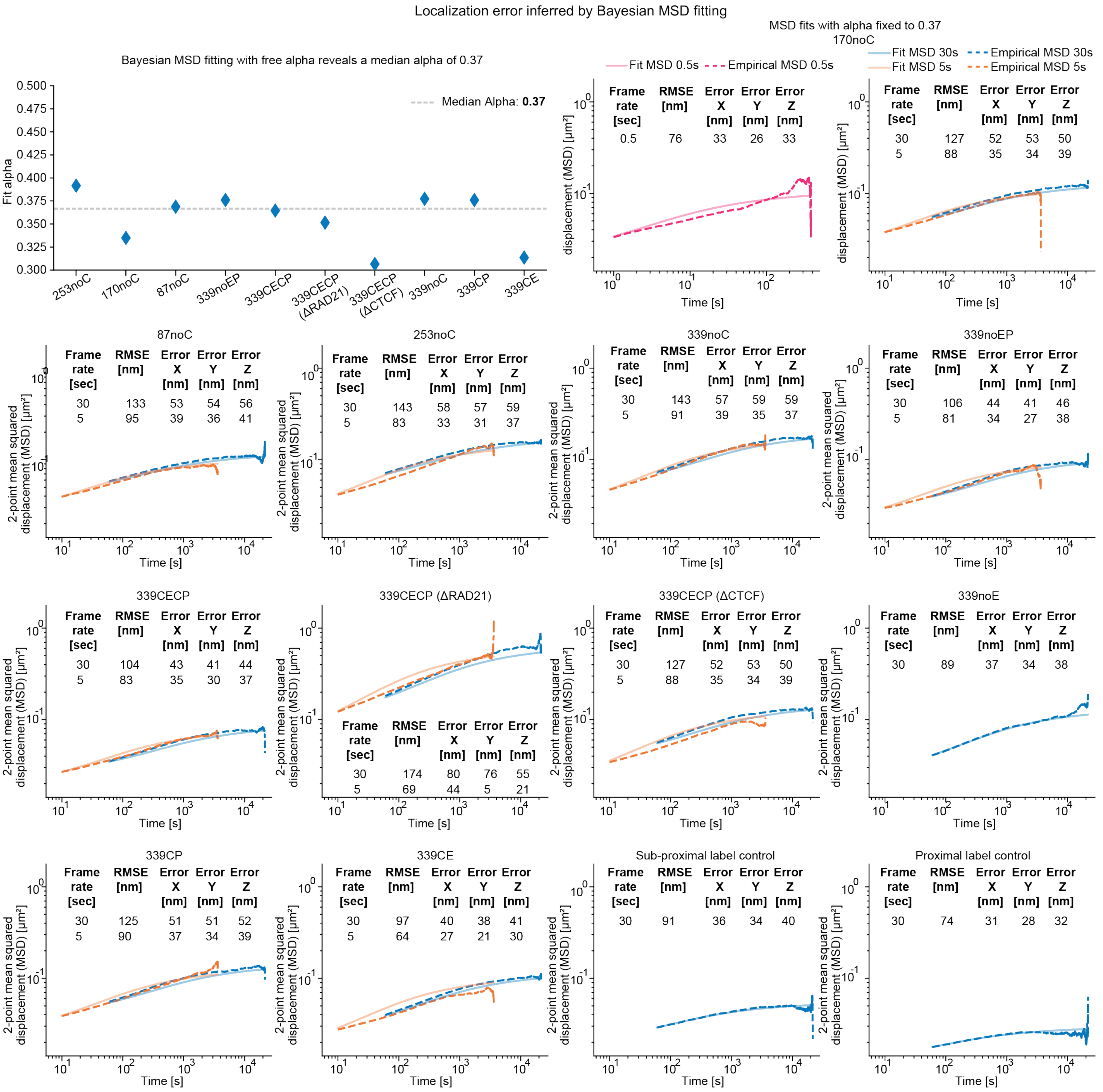

**Fig. S12: Localization errors inferred by Bayesian MSD fitting.** Bayesian MSD fitting [52] with free alpha was first performed to uncover a median alpha of 0.37. Subsequent Bayesian MSD fitting with alpha fixed 0.37 was then carried out to infer localization error along with spatial dimension. This value is consistent with previous MINFLUX tracking in mESCs, though we emphasize that chromatin dynamics does not appear to follow a power law across time in mESCs, such that the apparent alpha appears to depend on the timescale under study [53] For each condition, if 0.5-second and/or 5-second frame rate data were available and have over 40 trajectories, Bayesian MSD fit was performed jointly across data from different frame rates; otherwise Bayesian MSD fit was performed on 30-second data alone.

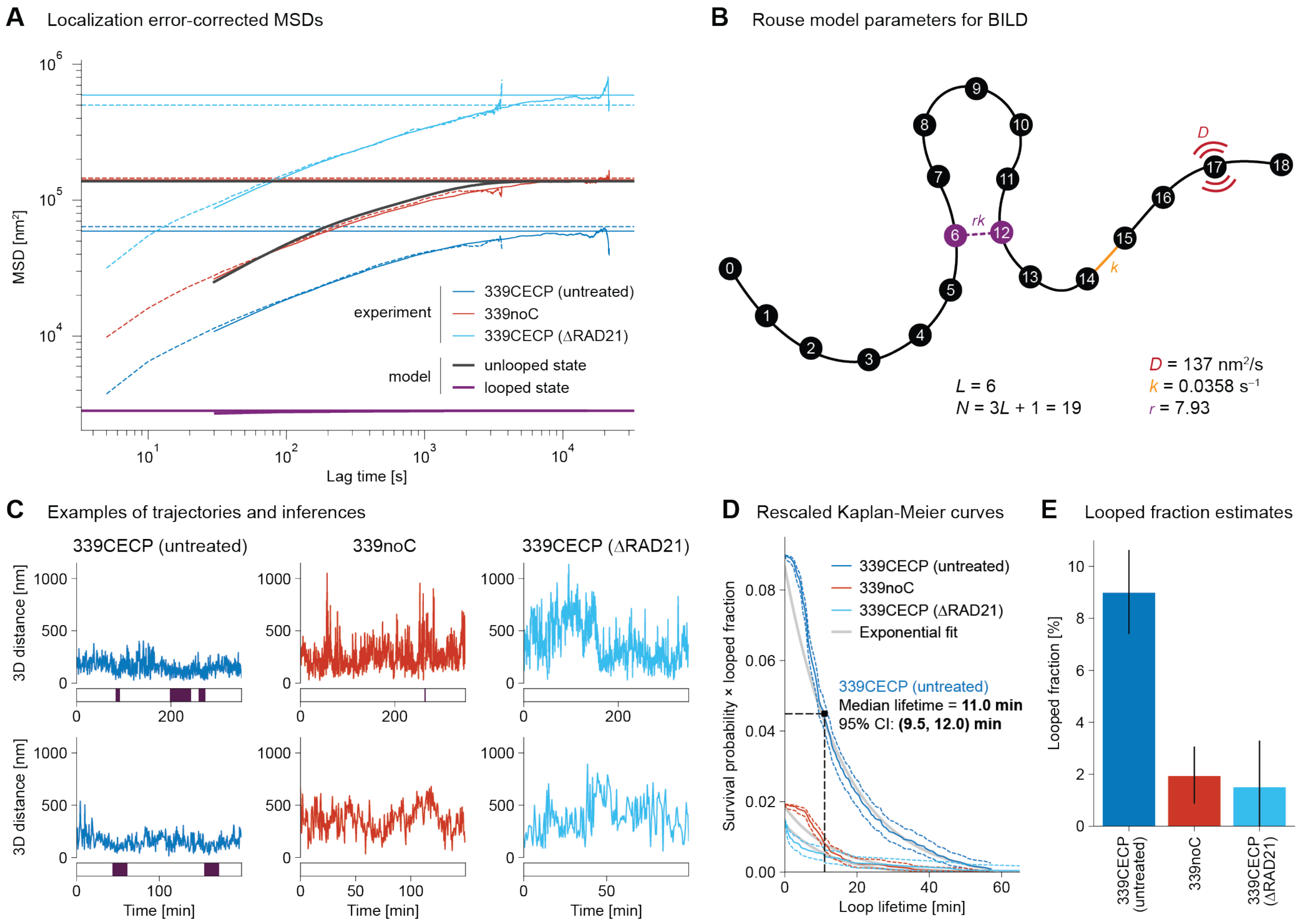

**Fig. S13: Bayesian Inference of Looping Dynamics (BILD) estimates looped fraction and loop lifetime of 339 kb E–P loop with convergent CBS (339CECP).** (**A**) Localization error-corrected two-point MSDs between fluorescent markers at enhancer and promoter. For experimental condition, dashed lines indicate MSDs of tracks collected with $\Delta t = 5\,\mathrm{s}$ between frames, and solid lines indicate MSDs of tracks collected with $\Delta t = 30\,\mathrm{s}$ between frames. Horizontal lines indicate the theoretical MSD plateau at $2\langle R^2\rangle$, where $\langle R^2\rangle$ is the mean squared distance between loop anchors. (**B**) Schematic of underlying Rouse model used for BILD [8], whose parameters are derived from the MSD fit results. (**C**) Example trajectories and their resulting BILD inferences. Inferred looping events are indicated by purple shading. (**D**) Kaplan-Meier survival curves of looping events rescaled by looped fraction. Dashed lines indicate bounds of 95% confidence intervals from Greenwood's formula. Gray lines indicate the survival curves of the maximum likelihood models assuming exponentially distributed lifetimes. (**E**) Mean looped fraction in all three conditions (false-positive corrected). Black lines indicate 95% bootstrapped confidence intervals.

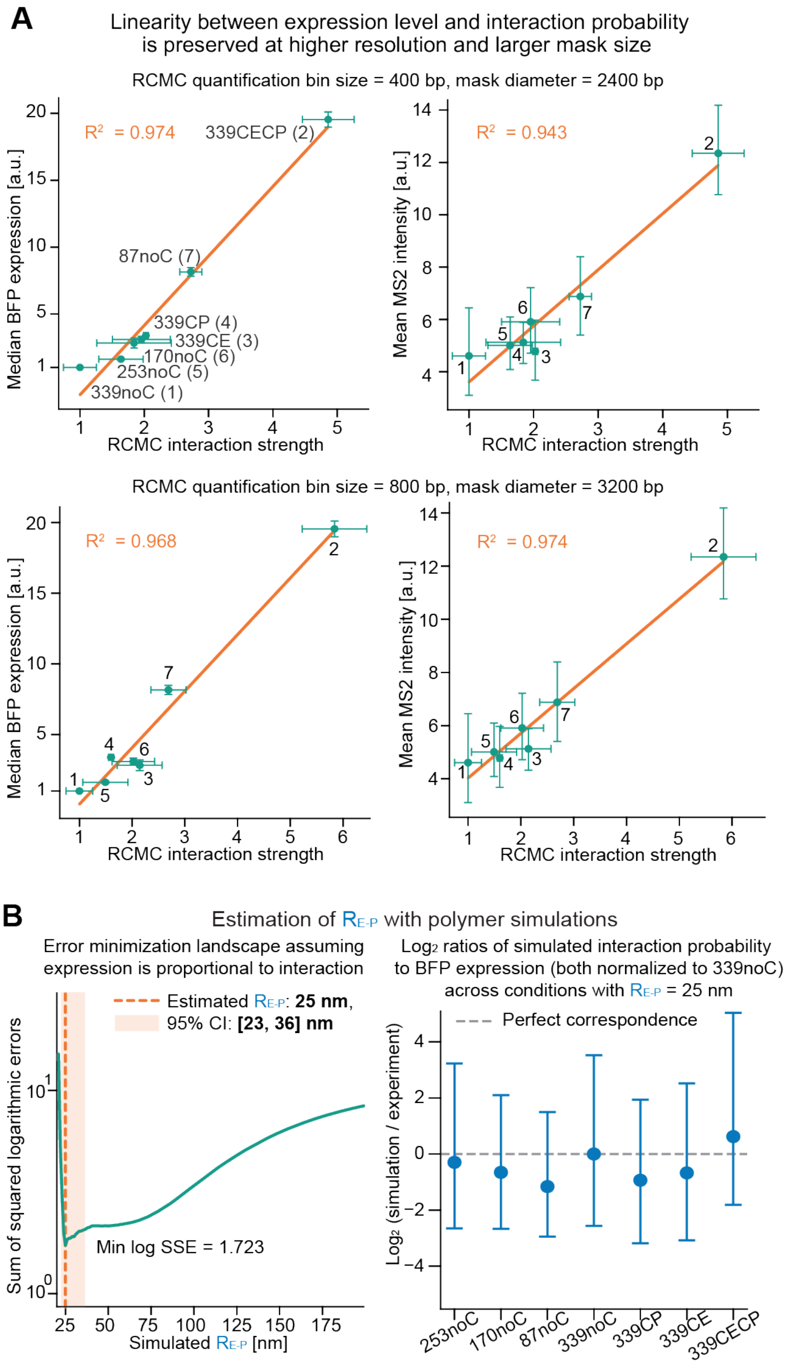

**Fig. S14: Linearity between expression and RCMC-quantified interaction strength with different bin sizes and mask sizes, and estimation of RCMC capture radius.** (**A**) Linear regression analysis demonstrating that the strong linear correlation between transcription levels (normalized median tagBFP expression or mean MS2 intensity) and RCMC interaction strength is preserved across different RCMC matrix resolutions (400 bp vs 800 bp) and quantification mask sizes (2400 bp vs 3200 bp diameters). (**B**) Estimation of $R_{\mathrm{E-P}}$ using 3D polymer simulations, under the assumption that gene expression is proportional to E-P interaction probability. The left panel plots the error minimization landscape (Sum of Squared Errors, SSE) comparing experimental median tagBFP expression against normalized simulated interaction probabilities across different theoretical interaction radii. Minimizing the sum of squared errors (SSE) between the experimental and simulated datasets yields an optimal $R_{\mathrm{E-P}}$ of 25 nm (95% CI: 23–36 nm). The right panel displays the $\log_2$ ratio of the simulated interaction probability to the observed BFP expression at the optimal $R_{\mathrm{E-P}}$ of 25 nm across seven E-P configurations (both metrics normalized to the 339noC reference). Central data points indicate the ratio of the population means and the error bars represent the combined 95% bounds of simulations and experiments.

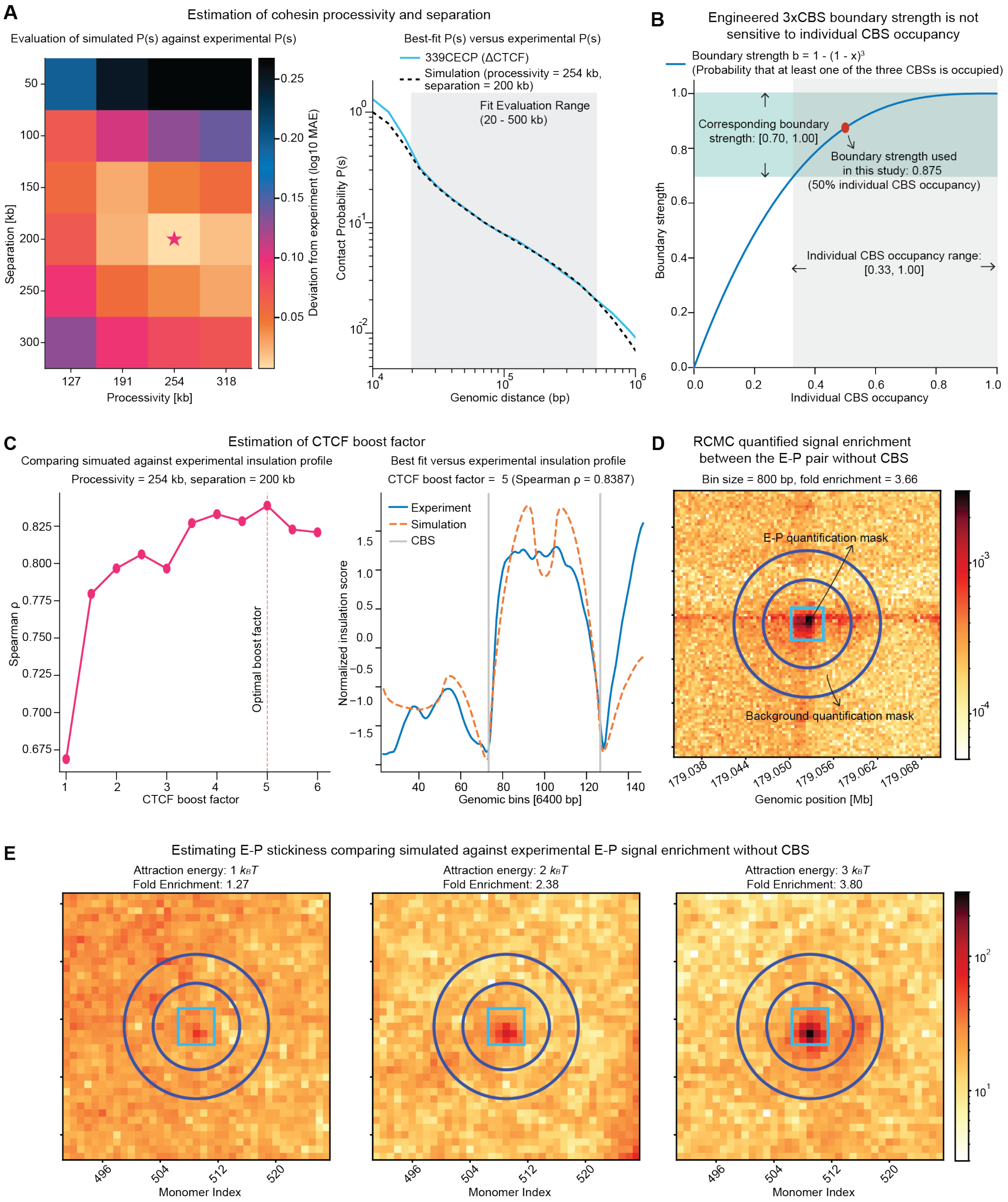

**Fig. S15: Estimation of parameters for polymer simulations.** (**A**) Heatmap evaluating simulated versus experimental contact probability decay ($P(s)$) to estimate cohesin processivity and separation. (**B**) Boundary strength of the engineered 3x CBS used in this study is robust to variations in individual CBS occupancy. (**C**) Estimation of the CTCF boost factor (the fold increase in cohesin residence time when stalled by CTCF) by comparing simulated against experimental insulation profiles. (**D**) RCMC-quantified focal signal enrichment between the E-P pair without CBSs. (**E**) Estimation of intrinsic E-P stickiness by comparing simulated versus experimental focal E-P signal enrichment, identifying an optimal attraction energy of $3k_BT$ used in polymer simulations.

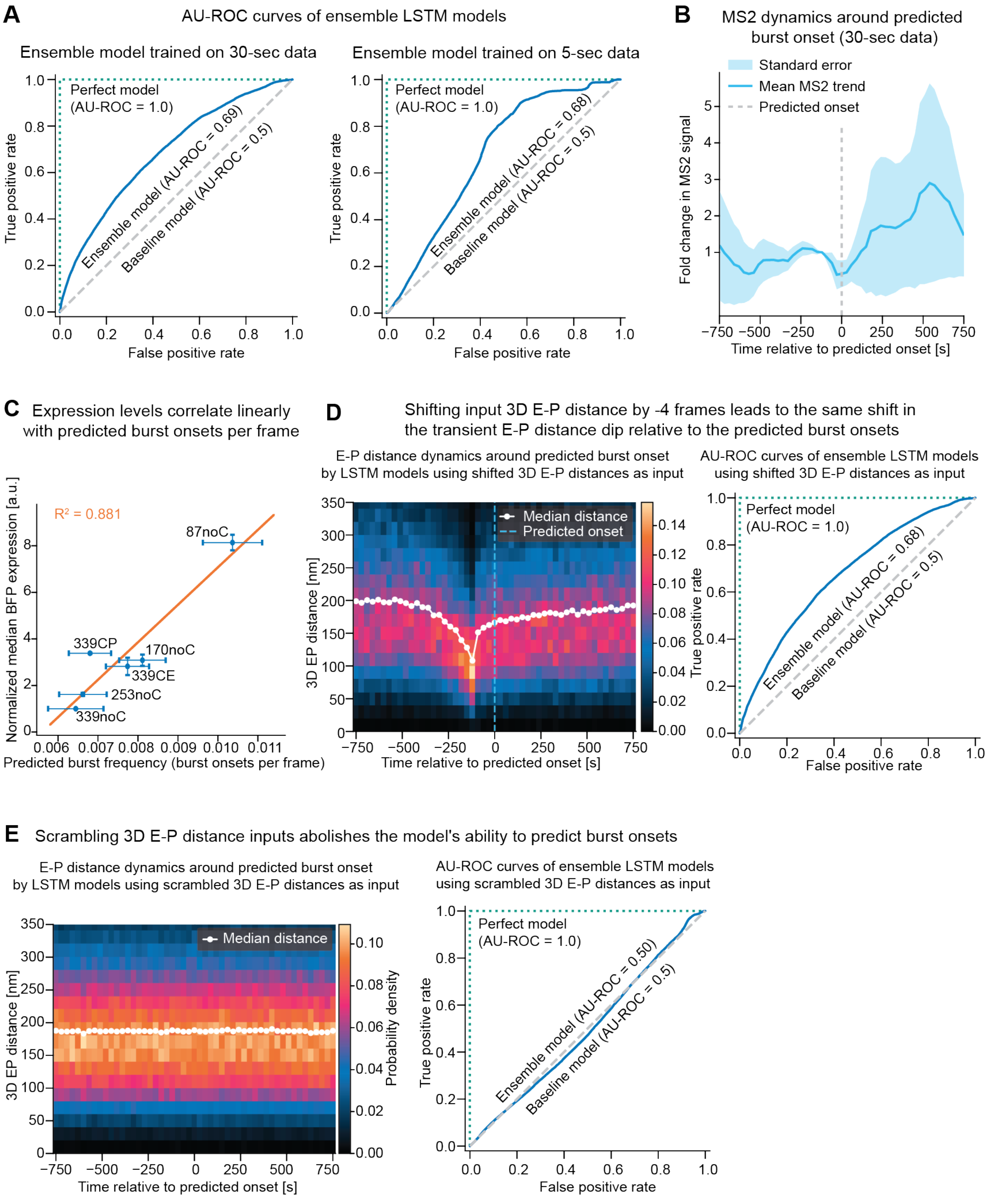

**Fig. S16: Evaluations and negative controls of the Long Short-Term Memory (LSTM) model.** (**A**) Receiver Operating Characteristic (AU-ROC) curves evaluating the performance of the ensemble LSTM models trained on 30-second (left) and 5-second (right) frame rate data. (**B**) Mean fold-change in MS2 signal aligned to the predicted burst onset time in the 30-second dataset. (**C**) Positive linear correlation between the frequency of predicted burst onsets and the normalized median tagBFP expression across conditions. (**D**) Positive control showing that shifting the input 3D E-P distance by -4 frames leads to the same shift in the transient E-P distance dip relative to the predicted burst onsets, while retaining $\sim$0.70 AU-ROC. (**E**) Negative control demonstrating that scrambling the 3D E-P distance inputs abolishes the model's predictive power (AU-ROC drops to 0.50) and eliminates the characteristic dip in 3D E-P distance prior to burst onsets.

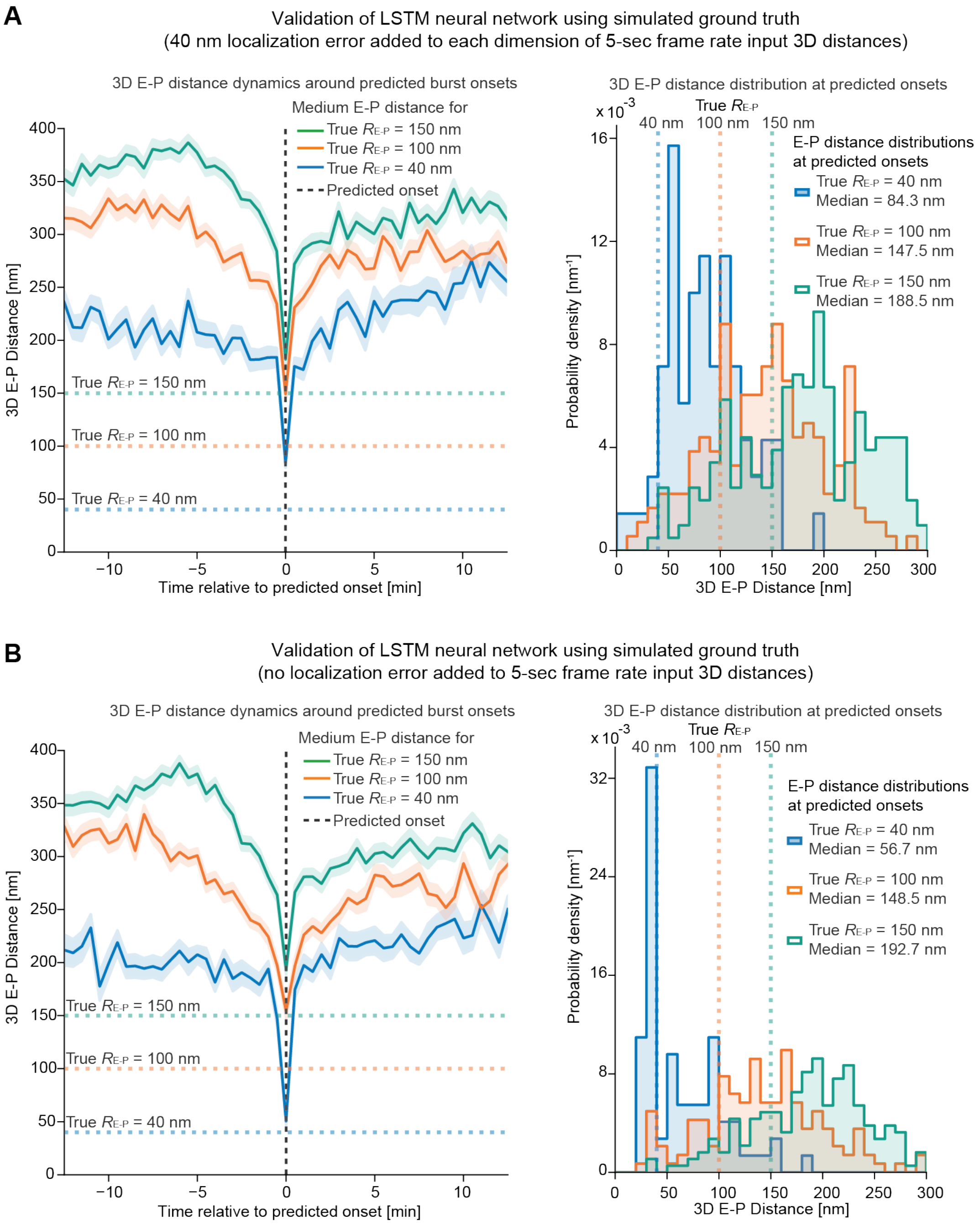

**Fig. S17: Validation of LSTM neural network using simulated ground truth.** (**A**) Validation with 40 nm localization error added to each dimension of the input 3D E-P distances to the LSTM. The input 3D distances from 3D polymer simulations described above were downsampled to a frame rate of 5 seconds to reflect the acquisition rate from our live-cell imaging experiments. Pile-up of 3D E-P distances at predicted burst onsets mimics the pile-up of 3D E-P distances from live-cell imaging experiments (**Fig.3B**) with a transient 'dip' at the predicted onsets. The 'dip' from LSTM trained on a ground truth $R_{\mathrm{E-P}}$ of 40 nm is significantly lower than those from LSTM neural networks trained with ground truth $R_{\mathrm{E-P}}$ of 100 nm and 150 nm, respectively (left panel). The shaded region around the median line represents the standard error of the mean (SEM). Similarly, the distribution of 3D E-P distances at predicted onsets from the LSTM trained with 40 nm ground truth $R_{\mathrm{E-P}}$ is significantly left-shifted from those using ground truth $R_{\mathrm{E-P}}$ of 100 nm and 150 nm (right panel). (**B**) The same validation with no localization error was added to the input 3D E-P distances. Pile-up of 3D E-P distances at predicted burst onsets using the 40 nm ground truth $R_{\mathrm{E-P}}$ reveals a transient 'dip' in E-P distance at onsets close to the ground truth $R_{\mathrm{E-P}}$ (left panel). The distribution of 3D E-P distances at predicted onsets from the LSTM trained with 40 nm ground truth $R_{\mathrm{E-P}}$ is roughly centered around the ground truth $R_{\mathrm{E-P}}$, and significantly left-shifted from those using ground truth $R_{\mathrm{E-P}}$ of 100 nm and 150 nm (right panel).

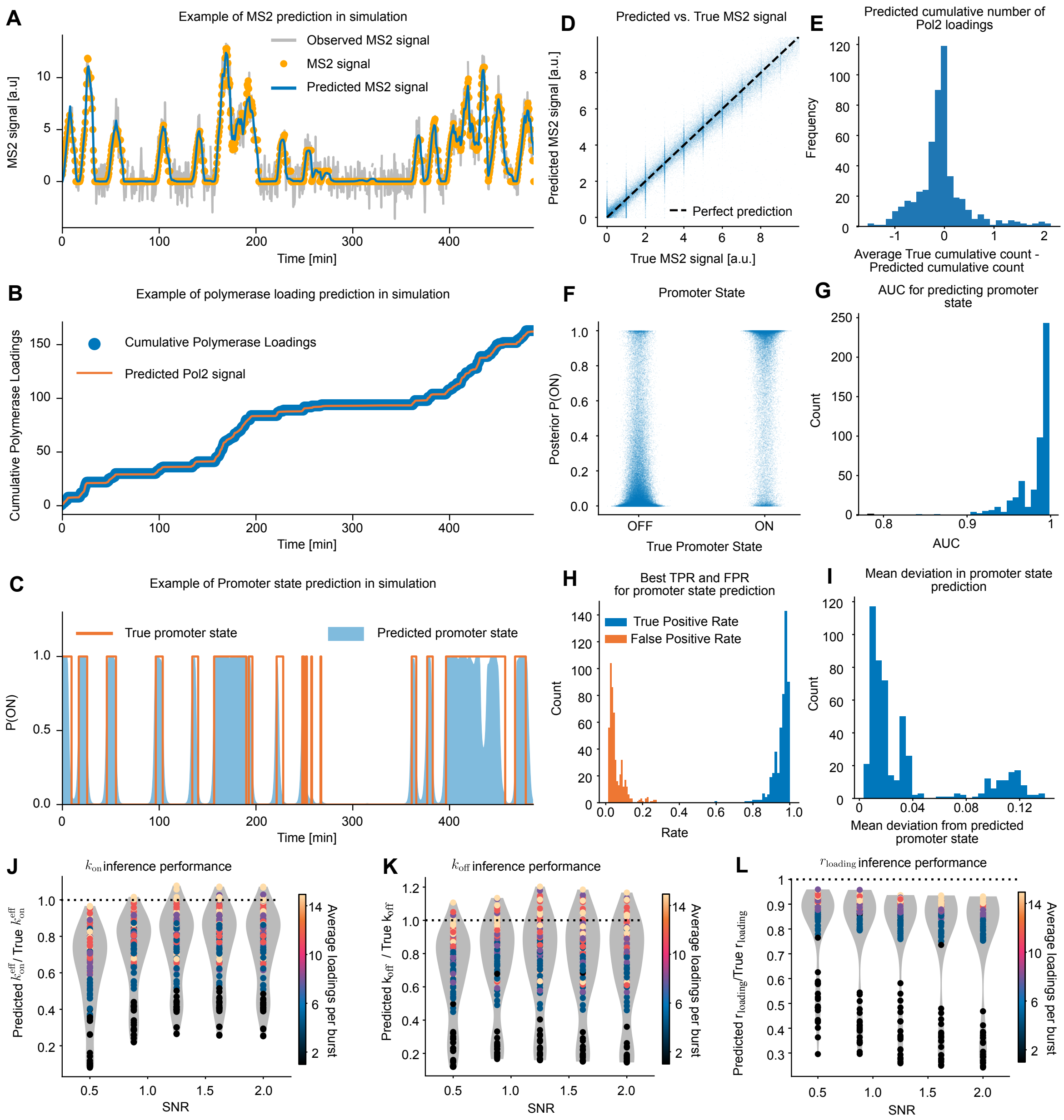

**Fig. S18: Validation of VEPI for MS2 inference.** (**A**) MS2 reconstruction on an example trajectory using Variational Enhancer-Promoter Inference (VEPI). Prediction is the mean of the predicted MS2 values across sampled Pol II loadings from the posterior. (**B**) Pol II reconstruction for the track in (A), displaying the cumulative count of loaded polymerases and the integral of the posterior average rate of Pol II loadings. (**C**) Promoter state reconstruction for the track in (A), showing the fraction of posterior trajectories that are on shaded blue against the true promoter state. (**D**) Predicted MS2 signal against true. (**E**) Average deviation between true and predicted cumulative loading counts across the entire trajectory which is between 250 min. and 500 min. (example shown in (B)). (**F**) Agreement between P(ON) and the true promoter state. (**G**) AUC for predicting promoter state by thresholding P(on) for each dataset. (**H**) True Positive Rate (TPR) and False Positive Rate (FPR), which maximized $\mathrm{TPR} - \mathrm{FPR}$ for each dataset. (**I**) Average absolute deviation between P(on) and the true promoter state per dataset. (**J**) Predicted $k_{\mathrm{on}}^{\mathrm{eff}}$ / True $k_{\mathrm{on}}^{\mathrm{eff}}$ versus Signal-to-Noise Ratio (SNR). (**K**) Predicted $k_{\mathrm{off}}$ / True $k_{\mathrm{off}}$ versus Signal-to-Noise Ratio (SNR). (**L**) Predicted loading rate over true loading rate versus Signal-to-Noise Ratio (SNR).

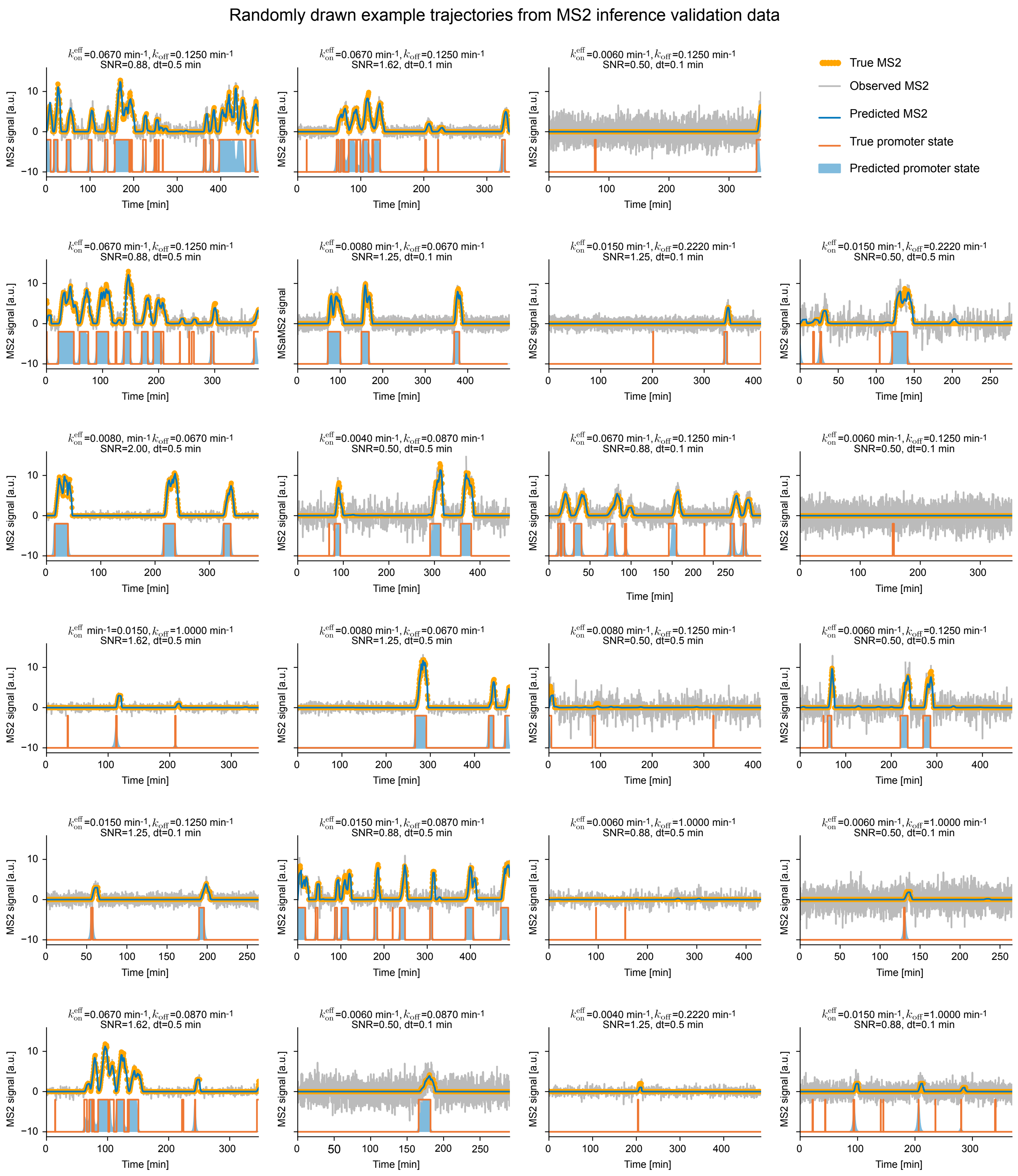

**Fig. S19: Example trajectories and predictions from the simulation data for MS2 validation**

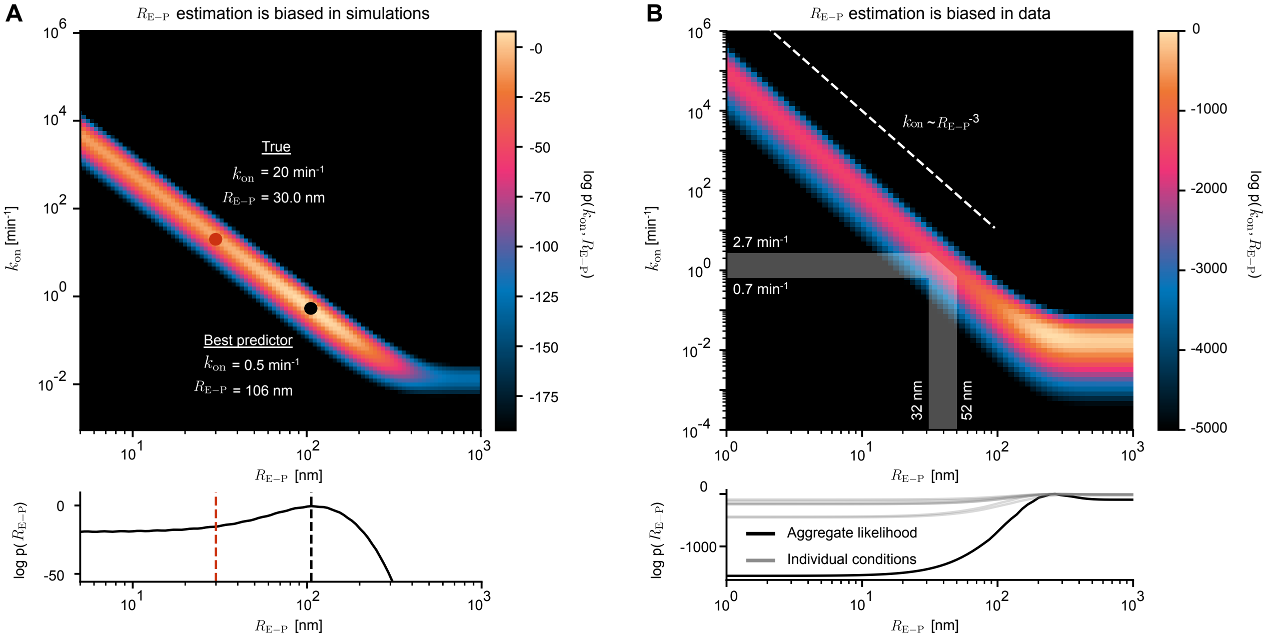

**Fig. S20: VEPI identifies a ridge relating $k_{\mathrm{on}}$ and $R_{\mathrm{E-P}}$ but does not independently identify $R_{\mathrm{E-P}}$** (**A**) $k_{\mathrm{on}}$, $R_{\mathrm{E-P}}$ likelihood in simulations. Log $p(R_{\mathrm{E-P}})$ was obtained by marginalizing over $k_{\mathrm{on}}$ by exponentiation and integrating over the column. (**B**) likelihood grid for 30 s data including the cell lines 87noC, 339CECP, 170noC, 339CE, 253noC, 339CP, 339noC.

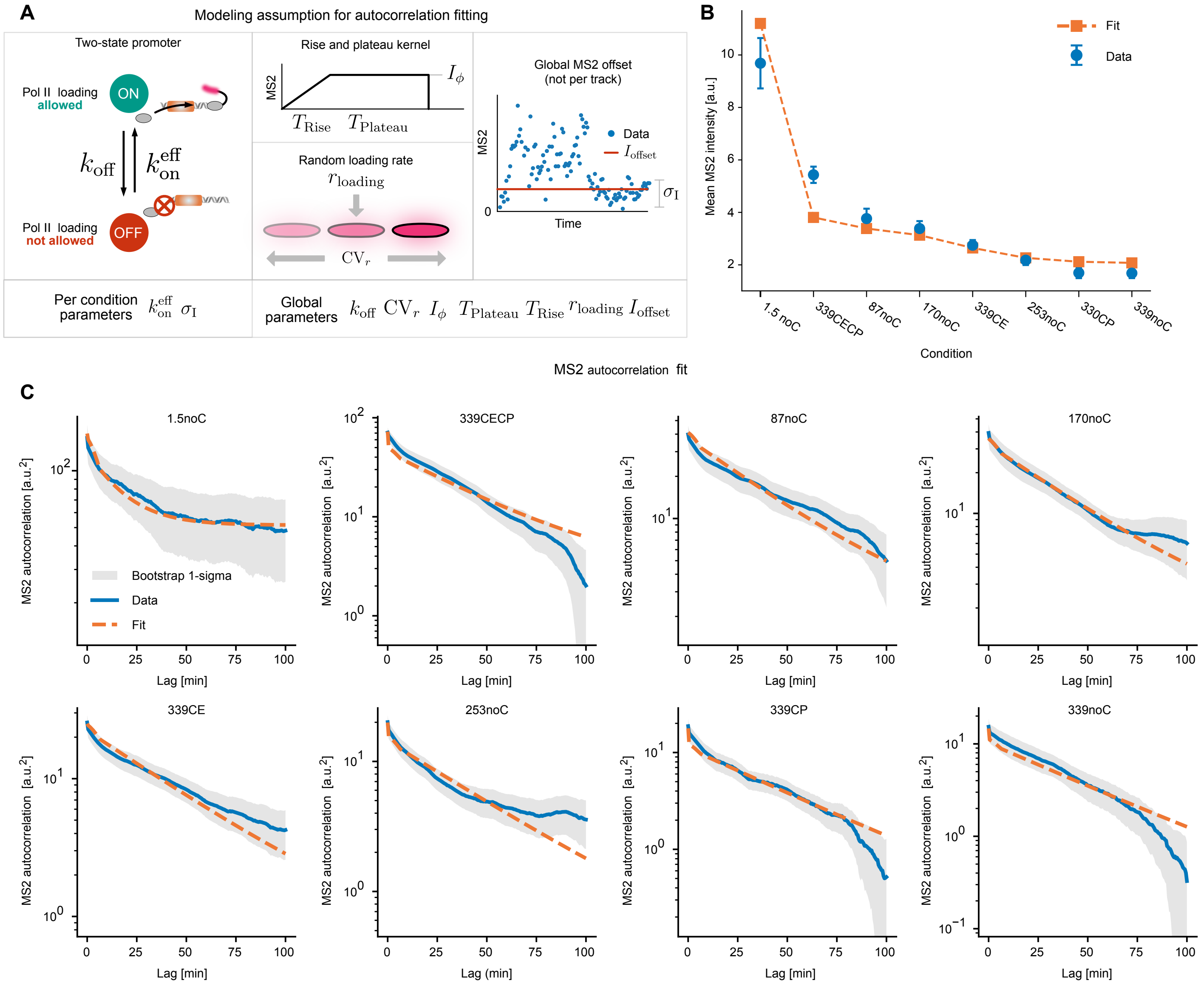

**Fig. S21: MS2 Autocorrelation Function Fits** (**A**) Assumptions of the model used to fit the MS2 autocorrelation function. We employed a two-state promoter model where loading was only allowed in the ON-state (left). Each Pol II loading was assumed to add the same fittable rise-and-plateau kernel to the MS2 signal (center top). Cell-to-cell heterogeneity was modeled as a gamma-distributed loading rate with a fittable coefficient of variation around the mean (center bottom). Finally, the MS2 was allowed to have a non-zero offset, which was taken to be a global value shared between all conditions and a per-condition noise value. (**B**) Comparison between mean MS2 and predicted values from the fit. Note that these were jointly fit with the covariance functions, so perfect agreement is not expected. (**C**) MS2 autocorrelation with function fits for each condition. Bootstrapped standard deviations were obtained from 10,000 bootstrap samples where tracks were redrawn with replacement from the dataset.

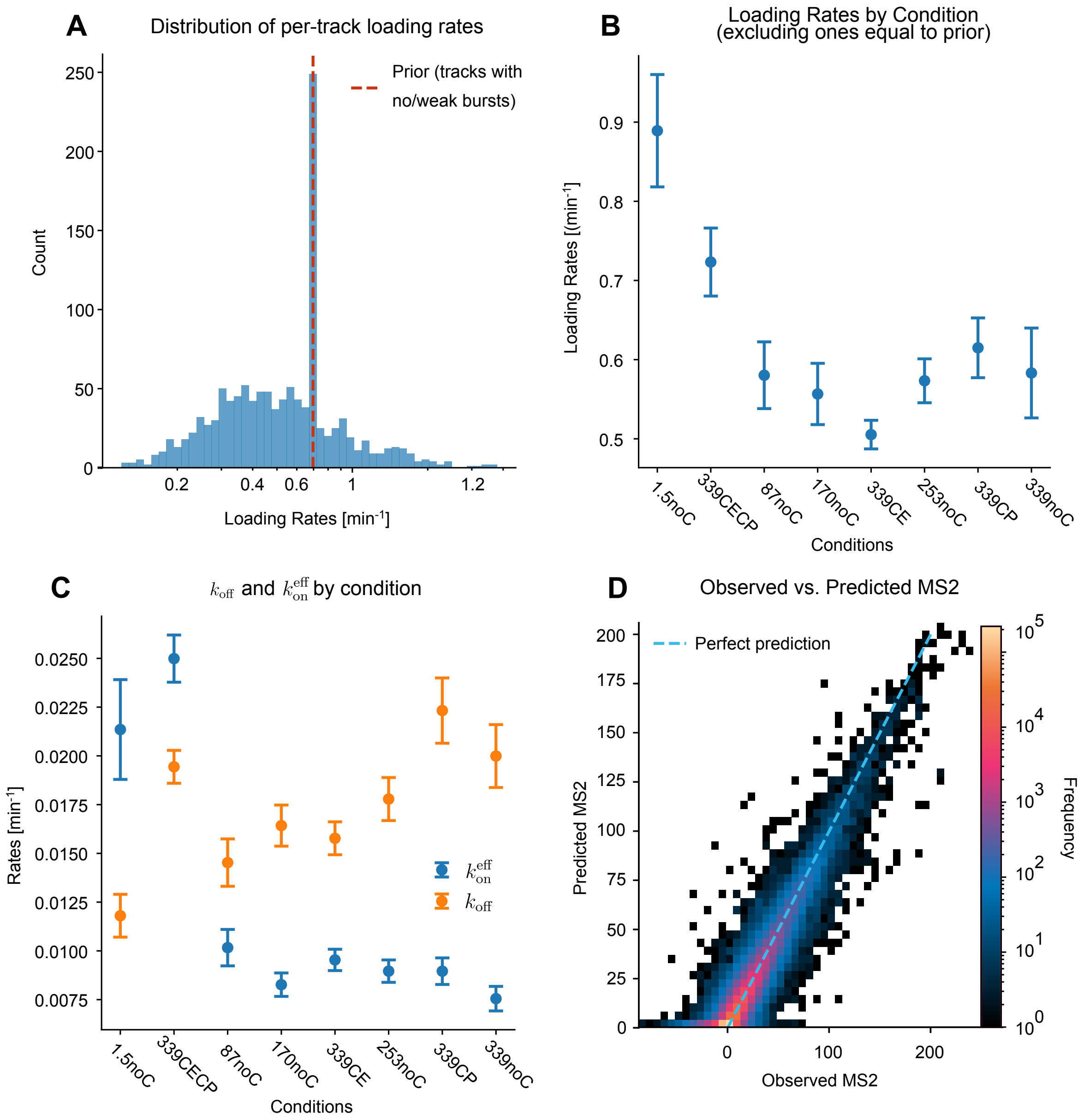

**Fig. S22: VEPI inference using MS2 alone** (**A**) Distribution of per-track loading rates across all conditions. Prior is highlighted as a significant fraction of trajectories did not leave this value. This is primarily due to few or no loading events in the trajectory. (**B**) Average $\pm$ standard error on the mean for the loading rate across conditions. (**C**) Inferred $k_{\mathrm{on}}^{\mathrm{eff}}$ and $k_{\mathrm{off}}$ across conditions. Data is posterior mean $\pm$ standard deviation (not standard error). (**D**) Predicted vs. observed MS2. All data were acquired at 30 s interval.

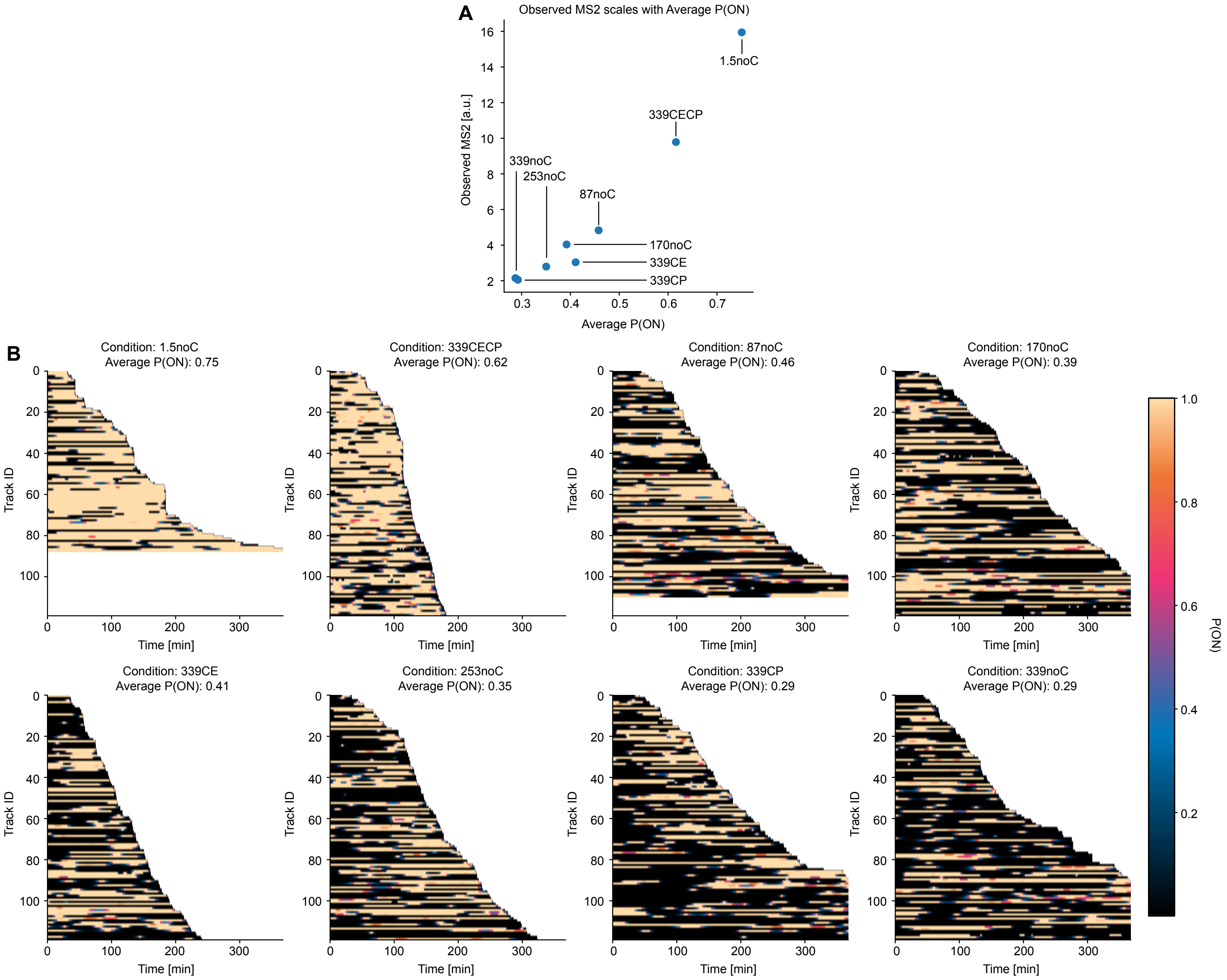

**Fig. S23: Burst predictions based on MS2 alone** (**A**) Plot of per-condition averaged MS2 versus average probability of the promoter being on. (**B**) Heatmaps showing the per-track inferred trajectories of the probability of the promoter being on. All data were acquired at 30 s interval.

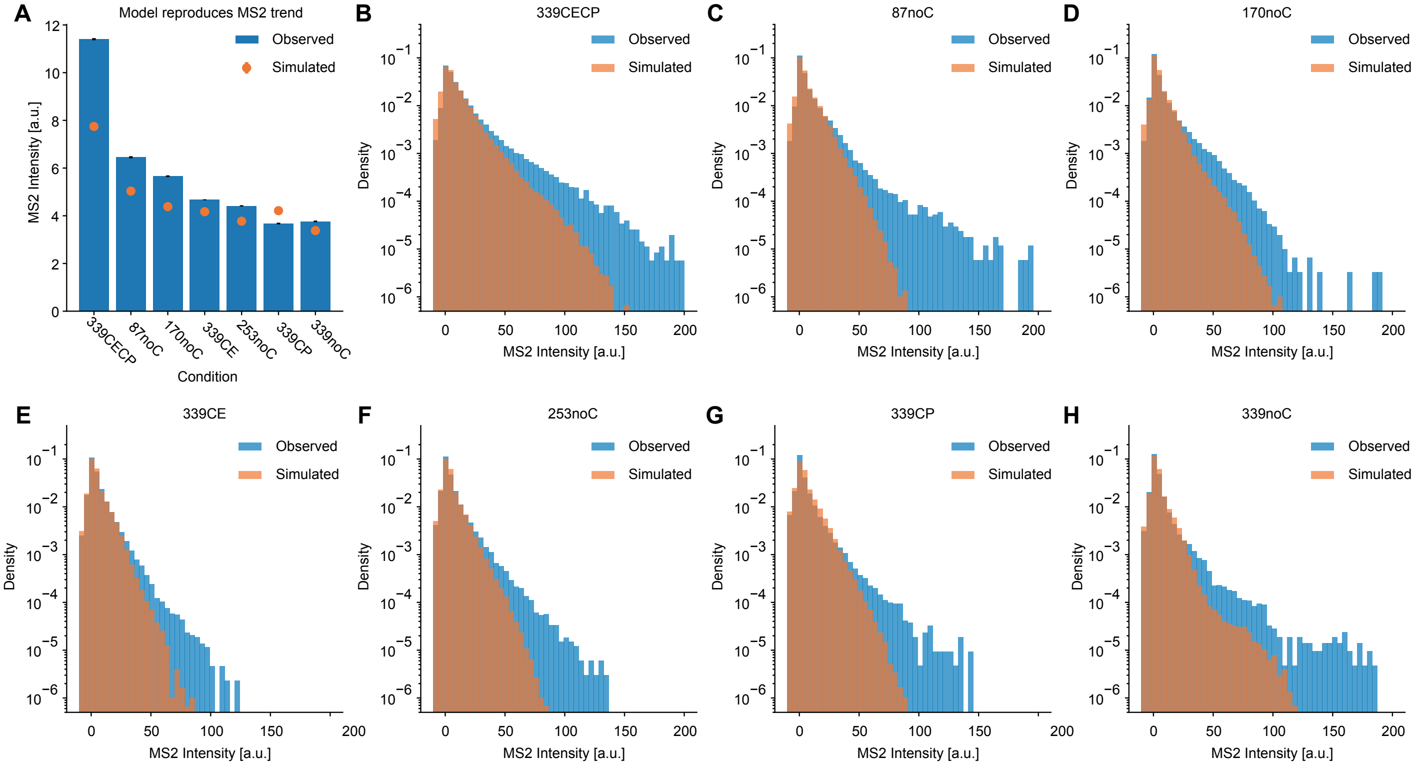

**Fig. S24: Predicted MS2 from the fitted E-P driven two-state model** (**A**) Mean for the simulated and observed MS2 trajectories. (**B**-**H**) Observed and predicted MS2 intensity distribution for the various conditions. All data were acquired at 30 s interval.

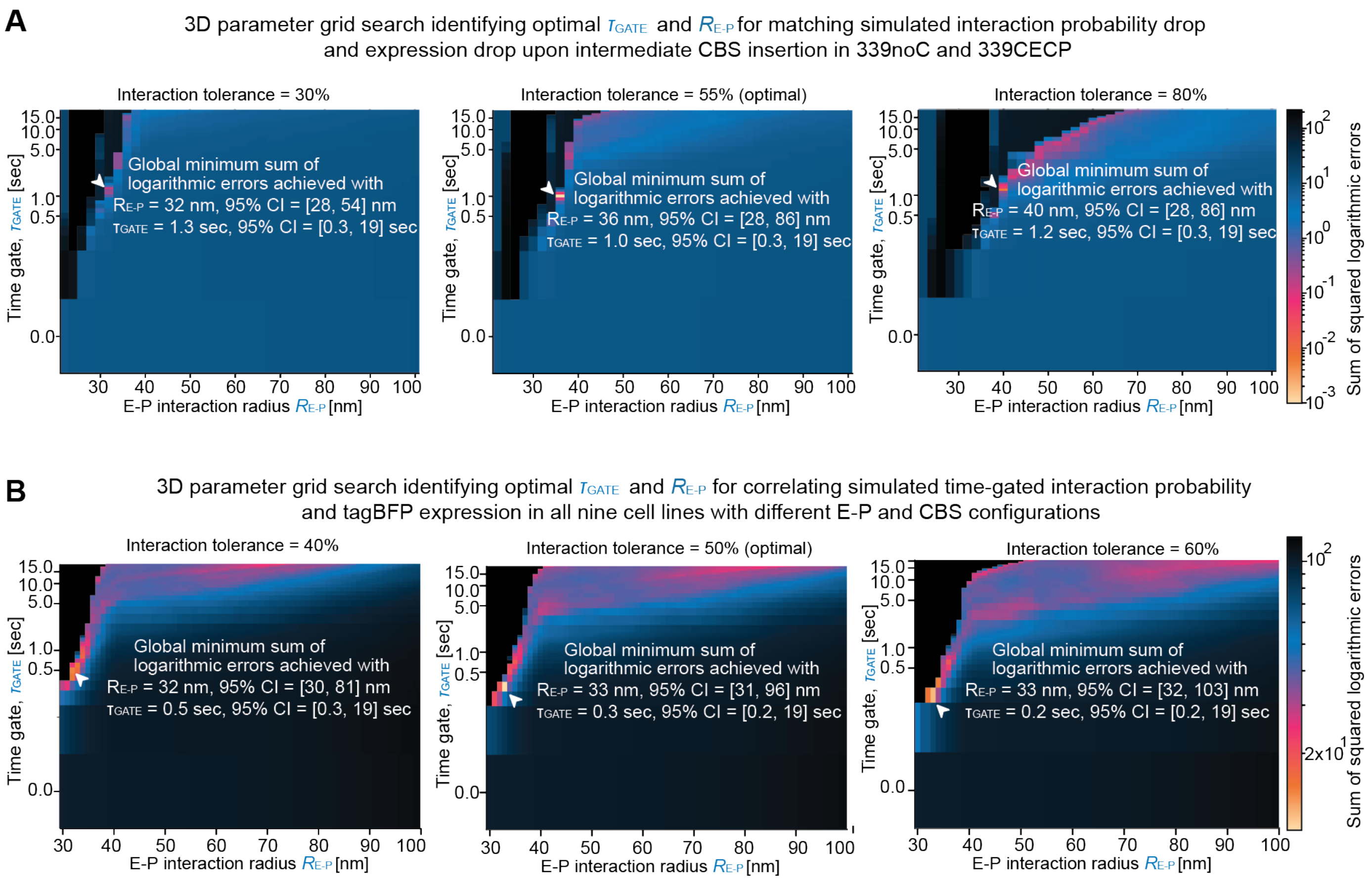

**Fig. S25: 3D parameter scanning identified an optimal time gate of ∼0.3–1.0 second and a corresponding $R_{\mathrm{E-P}}$ of ∼33–36 nm.** (**A**) A three-dimensional grid search (scanning interaction radius, time gate, and interaction tolerance) identified an optimal time gate ($\tau_{\mathrm{GATE}}$) of 1.0 second, an optimal $R_{\mathrm{E-P}}$ of 36 nm, and an optimal interaction tolerance of 55% to recapitulate the experimental reduction in tagBFP expression upon the insertion of a pair of 3xCBSs in the halfway between E-P pair in 339noC and 339CECP (middle panel). The interaction tolerance accounts for dynamic polymer flickering, stipulating that the E-P pair must remain within the 36 nm threshold for at least 55% of the 1.0-second window to count as productive interactions. The error minimization landscapes at interaction tolerances of 30% (left panel) and 80% (right panel) demonstrate that the optimal $R_{\mathrm{E-P}}$ and $\tau_{\mathrm{GATE}}$ estimates are robust to interaction tolerance values, yielding an optimal $R_{\mathrm{E-P}} \approx 32–40$ nm and an optimal $\tau_{\mathrm{GATE}} \approx 1.0–1.3$ seconds. (**B**) A three-dimensional grid search expanding the optimization to the correlation between simulated time-gated interaction probability and tagBFP expression in all nine cell lines with different E-P and CBS configuration (339noC, 339IC, 339CE, 339CP, 339CECP, 339CECPIC, 253noC, 170noC, 87noC) identified an optimal time gate ($\tau_{\mathrm{GATE}}$) of 0.3 second, an optimal $R_{\mathrm{E-P}}$ of 33 nm, and an optimal interaction tolerance of 50% (middle panel). Similar optimal $R_{\mathrm{E-P}}$ and $\tau_{\mathrm{GATE}}$ estimates were identified in other interaction tolerance levels (examples are shown for 40% and 60% in the left and right panels, respectively).

### 15 Supplementary Tables

**Table S1:** List of plasmids used in this study

| Name | Purpose | Associated sgRNA sequence |
|---|---|---|
| pASH267 | Repair plasmid used for editing the mAID degron tag (with a GDGAGLIN linker and V5 tag) onto the C-terminus of RAD21 | CCTCAGATAATATGGAACCG |
| pASH326 | Repair plasmid used for editing the FKBP(F36V) degron tag onto the C-terminus of CTCF | CTGGGGCCTTGCTCGGCACC |
| pASH323 | PiggyBac gene expression vector encoding L30-OR3-(GS)3-mStayGoldx2-P2A-T2A-BSD-SV40polyA | N/A |
| pASH329 | Repair plasmid used for inserting EF1$\alpha$-OsTIR(F74G) into the TIGRE safe harbor locus | ACTGCCATAACACCTAACTT |
| pASH449 | Repair plasmid used for inserting the synthetic promoter, 3x CTCF binding sites, and ANCH3 array | CGCCATACGGGTGACCCACC |
| pASH450 | Repair plasmid used for inserting the synthetic enhancer, 3x CTCF binding sites, and synBsr1 array | GATGCAATCAATTCGGGACA |
| pASH349 | PiggyBac gene expression vector encoding L30-OR3-NLS-2x-mStayGold-NLS | N/A |
| pASH400 | PiggyBac gene expression vector encoding L30-synBsr1-ParB-2x-mScarlet3-NLS | N/A |
| pASH380 | PiggyBac gene expression vector encoding EF1$\alpha$-MCP-2x-Halo | N/A |
| pASH375 | Plasmid encoding Cas9 from S. pyogenes, puromycin resistance gene, and two guide RNAs, modified from [7] | N/A |
| pASH470 | Repair plasmid used for deleting a 2331 bp segment containing an endogenous CTCF binding site $\sim$178,835,000 bp (mm39) on chr2 | GTGCTAAAGGTGGTCCGTGC,<br>GAGTTCAGTTCTGCGCGTTG |
| pASH447 | Plasmid encoding Vika recombinase and puromycin resistance gene, with an internal ribosome entry site (IRES) between the two genes, under the control of a pCAG promoter | N/A |
| pASH471 | Repair plasmid used for deleting $\sim$84 kb DNA between the two fluorescent labels in 15C-A2 | ATTGTGTTCAAGTCCCGATC,<br>ACCGTGAAGGCGCGAGAGTC |
| pASH448 | Plasmid encoding Dre recombinase and puromycin resistance gene, with an internal ribosome entry site (IRES) between the two genes, under the control of a pCAG promoter | N/A |
| pASH445 | Plasmid encoding SCre recombinase and puromycin resistance gene, with an internal ribosome entry site (IRES) between the two genes, under the control of a pCAG promoter | N/A |
| pASH446 | Plasmid encoding VCre recombinase and puromycin resistance gene, with an internal ribosome entry site (IRES) between the two genes, under the control of a pCAG promoter | N/A |
| pASH472 | Repair plasmid used for deleting $\sim$84 kb DNA between the 339 kb synthetic E-P pair to generate the 253 kb E-P pair | CACTCCAGGTGTCCCGTCAT,<br>TATCGATCACCTCCAGCCAG |
| pASH473 | Repair plasmid used for deleting $\sim$169 kb DNA between the 339 kb synthetic E-P pair to generate the 170 kb E-P pair | CACTCCAGGTGTCCCGTCAT,<br>TAGTCCCTTGTAGAATCGAA |
| pASH474 | Repair plasmid used for deleting $\sim$254 kb DNA between the 339 kb synthetic E-P pair to generate the 87 kb E-P pair | CACTCCAGGTGTCCCGTCAT,<br>CTCCTACAGAAGTCGATGTG |
| pASH475 | Repair plasmid used for deleting $\sim$81 kb DNA between the 87 kb synthetic E-P pair in 15C-A2 | ATTGTGTTCAAGTCCCGATC,<br>CAGGTGGCACTCCCGTATGG |
| pASH476 | Repair plasmid used for inserting the synthetic enhancer downstream and right next to the synthetic promoter | CGCCTTCACGGTCTAGGTTG |
| pASH477 | Repair plasmid used for inserting the insulating CTCF binding sites between the 339 kb E-P pair | GCAGATGCGGGCGTATACAT |

**Table S2:** List of primers used in this study

| Name | Sequence | Purpose |
|---|---|---|
| P779 | GGGCTGAAGCAACGAATAGA | Primer for genotyping FKBP(F36V) tagging of CTCF |
| P780 | CACAGTAAACCCTCAGAGACTTAG | Primer for genotyping FKBP(F36V) tagging of CTCF |
| P978 | AACCATTAGATGAAGGCTTTGGAG | Primer for genotyping insertion of OsTIR(F74G) expression cassette at the TIGRE safe harbor locus |
| P981 | AGGTTTTAGGGGAACTCGGATG | Primer for genotyping insertion of Ostir(F74G) expression cassette at the TIGRE safe harbor locus |
| P1444 | TGCATCCTCTCATTGCTCCG | Primer for genotyping insertion of the synthetic promoter construct along with 3x CTCF binding sites and ANCH3 array |
| P1445 | TGCATTCTAGTTGTGGTTTGTCC | Primer for genotyping insertion of the synthetic promoter construct along with 3x CTCF binding sites and ANCH3 array |
| P1490 | GACCAGGGCTCAGCAGATTT | Primer for genotyping insertion of the synthetic enhancer construct along with 3x CTCF binding sites and synBsr1 array |
| P1491 | TCTGGCCTGCAGGAGATAGT | Primer for genotyping insertion of the synthetic enhancer construct along with 3x CTCF binding sites and synBsr1 array |
| P1591 | GGGACTAGCCATTGCACACT | Primer for genotyping deletion of a 2331 bp segment containing an endogenous CTCF binding site ∼178,835,000 bp (mm39) on chr2 |
| P1592 | GAGATGTGGGTTCTGCACCA | Primer for genotyping deletion of a 2331 bp segment containing an endogenous CTCF binding site ∼178,835,000 bp on chr2 |
| P1858 | ATTCCACAACCTAGCAATTCTGAAG | Primer for genotyping excision of the synthetic enhancer by the Vika recombinase |
| P1859 | CTTGGTCATTGCCCTACGCT | Primer for genotyping excision of the synthetic enhancer by the Vika recombinase |
| P1911 | ACACGTTTGATTTGGGCAGTG | Primer for genotyping excision of the synthetic enhancer by the Dre recombinase |
| P1912 | CAAGCCGAGTCTGGCGCAATG | Primer for genotyping excision of the synthetic enhancer by the Dre recombinase |
| P1860 | TCCAAACCTCATCAAAGTGGG | Primer for genotyping excision of the engineered 3x CTCF binding sites upstream of the synthetic enhancer by the SCre recombinase |
| P1861 | TTGGTCATTGCCCTACGCTG | Primer for genotyping excision of the engineered 3x CTCF binding sites upstream of the synthetic enhancer by the SCre recombinase |
| P1862 | TCCAAACCTCATCAAAGTGGG | Primer for genotyping excision of the engineered 3x CTCF binding sites downstream of the synthetic promoter by the VCre recombinase |
| P1863 | TTGGTCATTGCCCTACGCTG | Primer for genotyping excision of the engineered 3x CTCF binding sites downstream of the synthetic promoter by the VCre recombinase |
| P1964 | TCTTGCCCTCACACCTGAAC | Primer for genotyping deletion of a weak endogenous CTCF binding site upstream of the synthetic enhancer |
| P1965 | GCTGGAAAAGCCCTGGAGTA | Primer for genotyping deletion of a weak endogenous CTCF binding site upstream of the synthetic enhancer |
| P1897 | GTATGACCGGCACCTCATGT | Primer for genotyping deletion of ∼84 kb DNA between the 339 kb E-P pair in S+V-A6 to generate the 253 kb E-P pair |
| P1898 | GGCACCTGGAGATCTTGCAT | Primer for genotyping deletion of ∼84 kb DNA between the 339 kb E-P pair in S+V-A6 to generate the 253 kb E-P pair |
| P1903 | GGAGGGATGATGTAGCCACG | Primer for genotyping deletion of ∼169 kb DNA between the 339 kb E-P pair in S+V-A6 to generate the 170 kb E-P pair |
| P1904 | CACCGTGCATTTTTGCTGGA | Primer for genotyping deletion of ∼169 kb DNA between the 339 kb E-P pair in S+V-A6 to generate the 170 kb E-P pair |
| P1907 | CTCTGGGGGAAATGCGAAGT | Primer for genotyping deletion of ∼254 kb DNA between the 339 kb E-P pair in S+V-A6 to generate the 87 kb E-P pair |
| P1908 | CAGAGGTTGGGCGTCTTCTT | Primer for genotyping deletion of ∼254 kb DNA between the 339 kb E-P pair in S+V-A6 to generate the 87 kb E-P pair |
| P1864 | TAGAACCCACAACGTGACCG | Primer for genotyping deletion of ∼84 kb DNA between the fluorescent labels in 15C-A2 which will have 2362 bp remaining between the fluorescent labels |
| P1865 | AGCGCTCACAATTTGCCAAG | Primer for genotyping deletion of ∼84 kb DNA between the fluorescent labels in 15C-A2 which will have 2362 bp remaining between the fluorescent labels |
| P1866 | GCTGACGGCAGGGAGTATAG | Primer for genotyping deletion of ∼81 kb DNA between the synthetic E-P pair in 15C-A2 |
| P1867 | TGGGGCTGTCACAATCTTCAAT | Primer for genotyping deletion of ∼81 kb DNA between the synthetic E-P pair in 15C-A2 |
| P2001 | ATTCTCATGGTCTGGGTGCC | Primer for genotyping insertion of the synthetic enhancer downstream and right next to the synthetic promoter |
| P2002 | TGGATGACCATTTGAGGGGC | Primer for genotyping insertion of the synthetic enhancer downstream and right next to the synthetic promoter |
| P2031 | GTTGGCCAGCTGTGTGAATG | Primer for genotyping insertion of the insulating CTCF binding sites between the 339 kb E-P pair |
| P2032 | GAAGTCTCAAGTCTCCGCCC | Primer for genotyping insertion of the insulating CTCF binding sites between the 339 kb E-P pair |

**Table S3:** Trajectory summary for 30s Dataset

| Cell Line (condition) | Number of trajectories | Average trajectory length | Total frame count |
|---|---|---|---|
| 339CECP | 226 | 377 | 85312 |
| 339CECP (△CTCF) | 192 | 468 | 89893 |
| 339CECP (△RAD21) | 78 | 355 | 27690 |
| 339CE | 241 | 432 | 104180 |
| 339CP | 121 | 435 | 52586 |
| 339noEP | 114 | 374 | 42608 |
| 339noE | 45 | 491 | 22097 |
| 339noC | 121 | 427 | 51648 |
| 253noC | 182 | 444 | 80845 |
| 170noC | 147 | 497 | 73066 |
| 87noC | 111 | 371 | 41155 |
| 1.5noC | 101 | 247 | 24963 |
| Proximal label control | 43 | 299 | 12843 |
| Sub-proximal label control | 70 | 356 | 24905 |
| 339noC (△RAD21) | 11 | 281 | 3093 |
| 253noC (△RAD21) | 14 | 335 | 4692 |
| 170noC (△RAD21) | 23 | 305 | 7016 |
| 87noC (△RAD21) | 36 | 211 | 7608 |
| 1.5noC (△RAD21) | 9 | 307 | 2762 |
| 339CECP (△RAD21 & △CTCF) | 6 | 429 | 2574 |

**Table S4:** Trajectory summary for 5s Dataset

| Cell Line (condition) | Number of trajectories | Average trajectory length | Total frame count |
|---|---|---|---|
| 339CECP | 233 | 513 | 119633 |
| 339CECP (△CTCF) | 115 | 507 | 58310 |
| 339CECP (△RAD21) | 74 | 382 | 28248 |
| 339CE | 90 | 435 | 39162 |
| 339CP | 155 | 429 | 66553 |
| 339noEP | 90 | 443 | 39892 |
| 339noE | 15 | 307 | 4609 |
| 339noC | 168 | 413 | 69421 |
| 253noC | 153 | 413 | 63246 |
| 170noC | 153 | 481 | 73564 |
| 87noC | 125 | 413 | 51571 |

**Table S5:** Trajectory summary for 0.5s Dataset

| Cell Line (condition) | Number of trajectories | Average trajectory length | Total frame count |
|---|---|---|---|
| 170noC | 48 | 420 | 20174 |

**Table S6:** Parameters inferred by Bayesian MSD fitting (30s Dataset)

| Cell Line (condition) | Root-mean-square error [nm] | Error X [nm] | Error Y [nm] | Error Z [nm] | Prefactor $\Gamma$ [$\mu$m$^2$/s$^{0.37}$] | Root-mean-square distance [nm] |
|---|---|---|---|---|---|---|
| 339CECP | 104 | 43 | 41 | 44 | 0.0017 | 182 |
| 339CECP ($\triangle$CTCF) | 121 | 50 | 50 | 49 | 0.0033 | 243 |
| 339CECP ($\triangle$RAD21) | 174 | 80 | 76 | 55 | 0.0148 | 543 |
| 339CE | 97 | 40 | 38 | 41 | 0.0027 | 222 |
| 339CP | 125 | 51 | 51 | 52 | 0.0030 | 240 |
| 339noEP | 106 | 44 | 41 | 46 | 0.0022 | 205 |
| 339noE | 89 | 37 | 34 | 38 | 0.0032 | 242 |
| 339noC | 143 | 57 | 59 | 59 | 0.0040 | 280 |
| 253noC | 143 | 58 | 57 | 59 | 0.0037 | 260 |
| 170noC | 127 | 52 | 53 | 50 | 0.0029 | 222 |
| 87noC | 133 | 53 | 54 | 56 | 0.0028 | 206 |
| Proximal label control | 74 | 31 | 28 | 32 | 0.0009 | 97 |
| Sub-proximal label control | 91 | 36 | 34 | 40 | 0.0016 | 139 |

**Table S7:** Parameters inferred by Bayesian MSD fitting (5s Dataset)

| Cell Line (condition) | Root-mean-square error [nm] | Error X [nm] | Error Y [nm] | Error Z [nm] | Prefactor $\Gamma$ [$\mu$m$^2$/s$^{0.37}$] | Root-mean-square distance [nm] |
|---|---|---|---|---|---|---|
| 339CECP | 83 | 35 | 30 | 37 | 0.0017 | 182 |
| 339CECP ($\triangle$CTCF) | 71 | 30 | 24 | 33 | 0.0033 | 243 |
| 339CECP ($\triangle$RAD21) | 69 | 44 | 5 | 21 | 0.0148 | 543 |
| 339CE | 64 | 27 | 21 | 30 | 0.0027 | 222 |
| 339CP | 90 | 37 | 34 | 39 | 0.0030 | 240 |
| 339noEP | 81 | 34 | 27 | 38 | 0.0022 | 205 |
| 339noC | 91 | 39 | 35 | 37 | 0.0040 | 280 |
| 253noC | 83 | 33 | 31 | 37 | 0.0037 | 260 |
| 170noC | 88 | 35 | 34 | 39 | 0.0029 | 222 |
| 87noC | 95 | 39 | 36 | 41 | 0.0028 | 206 |

**Table S8:** Parameters inferred by Bayesian MSD fitting (0.5s Dataset)

| Cell Line (condition) | Root-mean-square error [nm] | Error X [nm] | Error Y [nm] | Error Z [nm] | Prefactor $\Gamma$ [$\mu$m$^2$/s$^{0.37}$] | Root-mean-square distance [nm] |
|---|---|---|---|---|---|---|
| 170noC | 76 | 33 | 26 | 33 | 0.0029 | 222 |

**Table S9:** Point estimate results from VEPI and MS2 autocorrelation fitting (30s Dataset)

| **Global parameters** | Meaning | Corrfit | Inference |
|---|---|---|---|
| $k_{\text{off}}$ | off rate [min$^{-1}$] | 0.02098 | 0.01747 |
| $r_{\text{loading}}$ | active-state loading rate [min$^{-1}$] | 0.6967 | 0.6142 |
| $\text{CV}_r$ | loading-rate coefficient of variation | 0.7488 | 0.6599 |
| $T_{\text{Rise}}$ | MS2 kernel rise time [min] | 0.4558 | — |
| $T_{\text{Plateau}}$ | MS2 kernel plateau time [min] | 6.055 | — |
| $I_{\phi}$ | intensity per loaded polymerase [a.u.] | 3.319 | — |
| $I_{\text{offset}}$ | mean-intensity offset [a.u.] | 1.620 | — |

| | $k_{\text{on}}^{\text{eff}}$ [min$^{-1}$] | | $\sigma_{\text{I}}$ [a.u] | |
|---|---|---|---|---|
| Condition | Corrfit | Inference | Corrfit | Inference |
| 1.5noC | 0.04063 | 0.02135 | 3.032 | 6.122 |
| 339CECP | 0.003720 | 0.02499 | 4.762 | 4.562 |
| 87noC | 0.002901 | 0.01017 | 0.06430 | 3.134 |
| 170noC | 0.002436 | 0.008266 | 0.6398 | 3.028 |
| 339CE | 0.001592 | 0.009537 | 0.7089 | 2.933 |
| 253noC | 0.0009771 | 0.008961 | 2.048 | 3.511 |
| 339CP | 0.0007465 | 0.008958 | 2.359 | 4.711 |
| 339noC | 0.0006795 | 0.007550 | 1.873 | 3.065 |

The inferred global $r_{\text{loading}}$ and $\text{CV}_r$ pools the per-track active-state loading rates across all listed conditions; $\text{CV}_r = \text{sd}(r_{\text{loading}})/\text{mean}(r_{\text{loading}})$. The inferred global $k_{\text{off}}$ is the track-count-weighted mean of condition-specific off rates. The inferred condition-specific $\sigma_{\text{I}}$ values are mean per-track noise estimates.

**Table S10:** List of software

| Name | Link | Source |
|---|---|---|
| python | https://www.python.org/ | [124] |
| notebook | https://jupyter.org/ | [125] |
| ipympl | https://matplotlib.org/ipympl/ | [126] |
| numpy | https://numpy.org/ | [127] |
| scipy | https://www.scipy.org/ | [54] |
| numba | https://numba.pydata.org/ | [40] |
| tqdm | https://tqdm.github.io/ | [128] |
| pyyaml | https://pyyaml.org/ | [129] |
| tifffile | https://pypi.org/project/tifffile/ | [130] |
| matplotlib | https://matplotlib.org/ | [131] |
| opencv-python | https://docs.opencv.org/4.12.0/ | [132] |
| aicspylibczi | https://pypi.org/project/aicspylibczi/ | [133] |
| trackpy | http://soft-matter.github.io/trackpy/v0.5.0/introduction.html | [134] |
| h5py | https://www.h5py.org/ | [135] |
| cupy-cuda12x | https://cupy.dev/ | [41] |
| bayesmsd | https://github.com/OpenTrajectoryAnalysis/bayesmsd | [136] |
| bioframe | https://github.com/open2c/bioframe | [137] |
| biopython | https://biopython.org/ | [138] |
| cooler | https://github.com/open2c/cooler | [61] |
| cooltools | https://github.com/open2c/cooltools | [139] |
| cython | https://cython.org/ | [140] |
| ipywidgets | https://ipywidgets.readthedocs.io/ | [141] |
| noctiluca | https://github.com/OpenTrajectoryAnalysis/noctiluca | [142] |
| pandas | https://pandas.pydata.org/ | [143] |
| polychrom | https://github.com/open2c/polychrom | [72] |
| pybbi | https://github.com/nvictus/pybbi | [144] |
| pyfaidx | https://github.com/mdshw5/pyfaidx | [145] |
| pysam | https://github.com/pysam-developers/pysam | [146] |
| ruptures | https://centre-borelli.github.io/ruptures-docs/ | [86] |
| scikit-learn | https://scikit-learn.org/ | [147] |
| seaborn | https://seaborn.pydata.org/ | [148] |
| openmm (simtk) | https://openmm.org/ | [73] |
| statsmodels | https://www.statsmodels.org/ | [149] |
| tensorflow | https://www.tensorflow.org/ | [150] |
| tracklib | https://github.com/OpenTrajectoryAnalysis/tracklib | [151] |
| jax | https://docs.jax.dev/en/latest/index.html | [90] |

#### 16 Movie Captions

**Movie S1**: Maximum intensity projections (MIPs) along each dimension (bottom) of the corresponding bounding boxes after applying nuclear segmentation masks for a representative 339CECP($\Delta$RAD21) cell line trajectory. Enhancer (synBsr1-ParB-2x-mScarlet3, magenta) and promoter (OR3-2x-mStayGold, green) loci were tracked for 721 frames with a 30 second time interval and 3D distance was measured between the localizations (top). Nascent transcription (MCP-2x-Halo, cyan) was measured based on the tracked promoter position.

**Movie S2**: Maximum intensity projections (MIPs) along each dimension (bottom) of the corresponding bounding boxes after applying nuclear segmentation masks for a representative 339CECP cell line trajectory. Enhancer (synBsr1-ParB-2x-mScarlet3, magenta) and promoter (OR3-2x-mStayGold, green) loci were tracked for 641 frames with a 30s second time interval and 3D distance was measured between the localizations (top). Nascent transcription (MCP-2x-Halo, cyan) was measured based on the tracked promoter position.

**Movie S3**: Maximum intensity projections (MIPs) along each dimension (bottom) of the corresponding bounding boxes after applying nuclear segmentation masks for a 87noC cell line representative trajectory. Enhancer (synBsr1-ParB-2x-mScarlet3, magenta) and promoter (OR3-2x-mStayGold, green) loci were tracked for 721 frames with a 30 second time interval and 3D distance was measured between the localizations (top). Nascent transcription (MCP-2x-Halo, cyan) was measured based on the tracked promoter position.

**Movie S4**: Maximum intensity projections (MIPs) along each dimension (bottom) of the corresponding bounding boxes after applying nuclear segmentation masks for a representative 339CECP cell line trajectory. Enhancer (synBsr1-ParB-2x-mScarlet3, magenta) and promoter (OR3-2x-mStayGold, green) loci were tracked for 721 frames with a 30 second time interval and 3D distance was measured between the localizations (top). Nascent transcription (MCP-2x-Halo, cyan) was measured based on the tracked promoter position.

**Movie S5**: Maximum intensity projections (MIPs) along each dimension (bottom) of the corresponding bounding boxes after applying nuclear segmentation masks for a representative 87noC cell line trajectory. Enhancer (synBsr1-ParB-2x-mScarlet3, magenta) and promoter (OR3-2x-mStayGold, green) loci were tracked for 724 frames with a 5 second time interval and 3D distance was measured between the localizations (top). Nascent transcription (MCP-2x-Halo, cyan) was measured based on the tracked promoter position.

**Movie S6**: Maximum intensity projections (MIPs) along each dimension (bottom) of the corresponding bounding boxes after applying nuclear segmentation masks for a representative 339CECP cell line trajectory. Enhancer (synBsr1-ParB-2x-mScarlet3, magenta) and promoter (OR3-2x-mStayGold, green) loci were tracked for 724 frames with a 5 second time interval and 3D distance was measured between the localizations (top). Nascent transcription (MCP-2x-Halo, cyan) was measured based on the tracked promoter position.

**Movie S7**: Maximum intensity projections (MIPs) along each dimension (bottom) of the corresponding bounding boxes after applying nuclear segmentation masks for a representative 170noC cell line trajectory. Enhancer (synBsr1-ParB-2x-mScarlet3, magenta) and promoter (OR3-2x-mStayGold, green) loci were tracked for 763 frames with a 0.5 second time interval and 3D distance was measured between the localizations (top). Nascent transcription (MCP-2x-Halo, cyan) was measured based on the tracked promoter position.

[33] M. B. Pappalardi, K. Keenan, M. Cockerill, W. A. Kellner, A. Stowell, C. Sherk, K. Wong, S. Pathuri, J. Briand, M. Steidel, P. Chapman, A. Groy, A. K. Wiseman, C. F. McHugh, N. Campobasso, A. P. Graves, E. Fairweather, T. Werner, A. Raoof, R. J. Butlin, L. Rueda, J. R. Horton, D. T. Fosbenner, C. Zhang, J. L. Handler, M. Muliaditan, M. Mebrahtu, J.-P. Jaworski, D. E. McNulty, C. Burt, H. C. Eberl, A. N. Taylor, T. Ho, S. Merrihew, S. W. Foley, A. Rutkowska, M. Li, S. P. Romeril, K. Goldberg, X. Zhang, C. S. Kershaw, M. Bantscheff, A. J. Jurewicz, E. Minthorn, P. Grandi, M. Patel, A. B. Benowitz, H. P. Mohammad, A. G. Gilmartin, R. K. Prinjha,

D. Ogilvie, C. Carpenter, D. Heerding, S. B. Baylin, P. A. Jones, X. Cheng, B. W. King, J. I. Luengo, A. M. Jordan, I. Waddell, R. G. Kruger, M. T. McCabe, Discovery of a first-in-class reversible DNMT1-selective inhibitor with improved tolerability and efficacy in acute myeloid leukemia. *Nat. Cancer* **2** (10), 1002–1017 (2021). doi: 10.1038/s43018-021-00249-x.

[57] M. K. Huseyin, C. K. Y. Hong, M. Nagano, J. A. Drayton, J. M. Jusuf, J. H. Yang, V. Ramanathan, V. Y. Goel, S. Kim, A. S. Hansen, Mapping 3D genome organization at nucleosome-scale with Micro-C and Region Capture Micro-C (RCMC). *Nat. Protoc.* (in press). doi: 10.1038/s41596-026-01393-3.

[92] C. M. Bishop, *Pattern recognition and machine learning*, Information science and statistics (Springer, New York) (2006).

[101] W. Bialek, T. Gregor, G. Tkačik, Action at a distance in transcriptional regulation. arXiv:1912.08579 [q-bio.SC] (2019).

[108] J. E. Moore, H. E. Pratt, K. Fan, N. Phalke, J. Fisher, S. I. Elhajjajy, G. Andrews, M. Gao, N. Shedd, Y. Fu, M. C. Lacadie, J. Meza, M. Khandpekar, M. Ganna, E. Choudhury, R. Swofford, H. Phan, C. C. Ramirez, M. Campbell, M. Likhite, N. P. Farrell, A. K. Weimer, A. Pampari, V. Ramalingam, F. Reese, B. Borsari, X. Yu, E. Wattenberg, M. Ruiz-Romero, M. Razavi-Mohseni, J. Xu, T. Galeev, A. Colubri, M. A. Beer, R. Guigó, M. B. Gerstein, J. M. Engreitz, M. Ljungman, T. E. Reddy, M. P. Snyder, C. B. Epstein, E. Gaskell, B. E. Bernstein, D. E. Dickel, A. Visel, L. A. Pennacchio, A. Mortazavi, A. Kundaje, Z. Weng, An expanded registry of candidate *cis*-regulatory elements. *Nature* (2026). doi: 10.1038/s41586-025-09909-9.

[127] C. R. Harris, K. J. Millman, S. J. van der Walt, R. Gommers, P. Virtanen, D. Cournapeau, E. Wieser, J. Taylor, S. Berg, N. J. Smith, R. Kern, M. Picus, S. Hoyer, M. H. van Kerkwijk, M. Brett, A. Haldane, J. F. del Río, M. Wiebe, P. Peterson, P. Gérard-Marchant, K. Sheppard, T. Reddy, W. Weckesser, H. Abbasi, C. Gohlke, T. E. Oliphant, Array programming with NumPy. arXiv:2006.10256 [cs.MS] (2020).

[128] C. da Costa-Luis, S. K. Larroque, K. Altendorf, H. Mary, richardsheridan, M. Korobov, N. Yorav-Raphael, I. Ivanov, M. Bargull, N. Rodrigues, Shawn, M. Dektyarev, M. Górny, mjstevens777, M. D. Pagel, M. Zugnoni, JC, CrazyPython, C. Newey, A. Lee, pgajdos, Todd, S. Malmgren, redbug312, O. Desh, N. Nechaev, M. Boyle, M. Nordlund, MapleCCC, J. McCracken, tqdm: A fast, extensible progress bar for Python and CLI (2024). doi: 10.5281/zenodo.14231923.

[129] K. Simonov, PyYAML: YAML parser and emitter for python (2025); https://pyyaml.org/.

[130] C. Gohlke, cgohlke/tifffile: v2025.10.16 (2025). doi: 10.5281/zenodo.17373292.

[131] J. D. Hunter, Matplotlib: A 2D graphics environment. *Comput. Sci. Eng.* **9** (3), 90–95 (2007). doi: 10.1109/MCSE.2007.55.

[132] G. Bradski, The OpenCV library. *Dr. Dobb's Journal of Software Tools* (2000).
